# Measuring thermodynamic preferences to form non-native conformations in nucleic acids using melting experiments reveals a rich sequence-specific DNA conformational landscape

**DOI:** 10.1101/2020.12.26.424438

**Authors:** Atul Rangadurai, Honglue Shi, Yu Xu, Bei Liu, Hala Abou Assi, John D. Boom, Huiqing Zhou, Isaac J. Kimsey, Hashim M. Al-Hashimi

## Abstract

Thermodynamic preferences to form non-native conformations are crucial for understanding how nucleic acids fold and function. However, they are difficult to measure experimentally because this requires accurately determining the population of minor low-abundance (<10%) conformations in a sea of other conformations. Here we show that melting experiments enable facile measurements of thermodynamic preferences to adopt non-native conformations in DNA and RNA. The key to this ‘delta-melt’ approach is to use chemical modifications to render specific minor non-native conformations the major state. The validity and robustness of delta-melt is established for four different non-native conformations under various physiological conditions and sequence contexts through independent measurements of thermodynamic preferences using NMR. delta-melt is fast, simple, cost-effective, and enables thermodynamic preferences to be measured for exceptionally low-populated conformations. Using delta-melt, we obtained rare insights into conformational cooperativity, obtaining evidence for significant cooperativity (1.0-2.5 kcal/mol) when simultaneously forming two adjacent Hoogsteen base pairs. We also measured the thermodynamic preferences to form G-C^+^ and A-T Hoogsteen and A-T base open states for nearly all sixteen trinucleotide sequence contexts and found distinct sequence-specific variations on the order of 2-3 kcal/mol. This rich landscape of sequence-specific non-native minor conformations in the DNA double helix may help shape the sequence-specificity of DNA biochemistry. Thus, melting experiments can now be used to access thermodynamic information regarding regions of the free energy landscape of biomolecules beyond the native folded and unfolded conformations.

**Significance Statement:** Thermodynamic preferences of nucleic acids to adopt non-native conformations are crucial for understanding how they function but prove difficult to measure experimentally. As a result, little is known about how these thermodynamic preferences vary with sequence and structural contexts, physiological conditions, and chemical modifications. Here, we show that modifications stabilizing non-native conformations and rendering them the major state, in conjunction with melting experiments, enable facile measurements of thermodynamic preferences to form various non-native conformations in DNA and RNA. delta-melt provided rare insights into the cooperativity of forming tandem Hoogsteen base pairs and revealed large and distinct sequence-specific preferences to form G-C^+^ and A-T Hoogsteen and A-T base open conformations in DNA, which may contribute to sequence-specific DNA biochemistry.

## Introduction

Biomolecules do not fold into a single structure but rather form dynamic ensembles of many interconverting conformations (1–3). Low populated non-native conformational states in the dynamic ensemble can become the major state when a molecule binds a partner molecule and forms a complex (Fig. 1a, b). In these cases, energetically favorable intermolecular contacts must cover the energetic cost, or conformational penalty 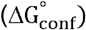, associated with stabilizing the minor non-native conformation relative to the major native state (Fig. 1a) (4–6). Therefore, the thermodynamic preferences to form non-native conformations can be essential determinants of binding affinity and specificity, and catalytic efficiency.

**Figure 1.**
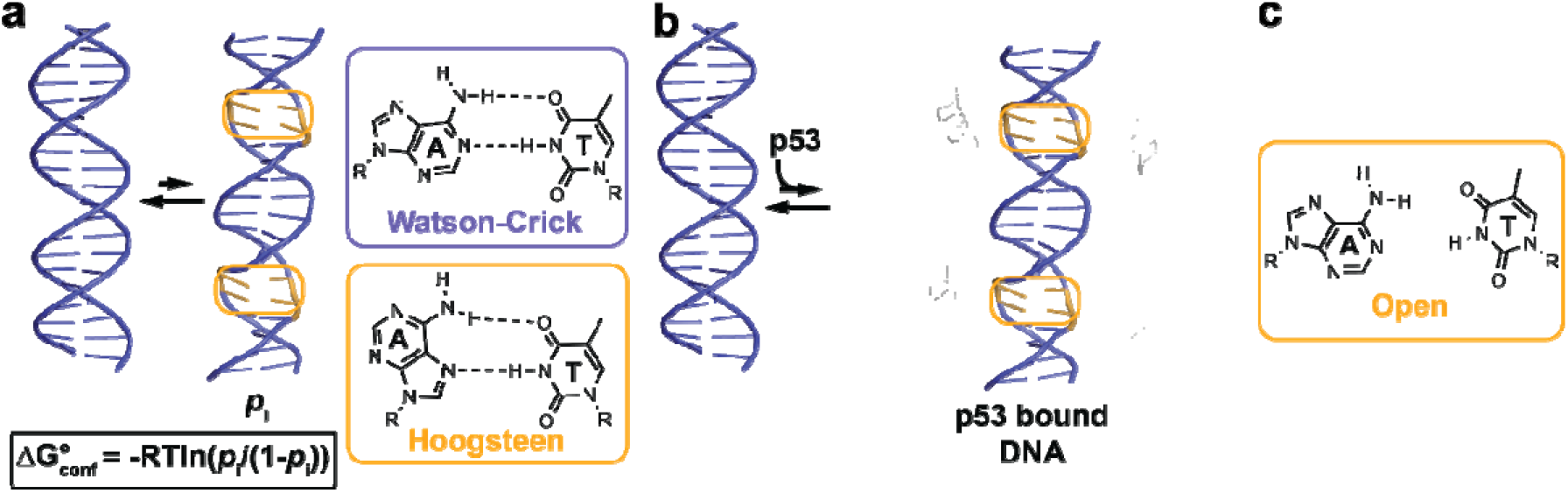
Thermodynamic preferences to form minor non-native conformations in DNA can influence binding. (a) The p53-bound conformation of a DNA duplex with Hoogsteen bps (in orange) exists as a minor conformation with population *p*_i_ in the unbound DNA ensemble. The free energy cost to stabilize the minor conformation on binding of p53 depends on *p*_i_ and is given by, in which *R* is the universal gas constant (units of kcal/mol/K) and T is the temperature (in Kelvin). (b) Binding of p53 to duplex DNA (PDB ID: 3KZ8) (9) results in a change in the conformation of bps from Watson-Crick (purple) in the free DNA to Hoogsteen (orange) in the DNA-p53 complex. The free DNA structure corresponds to an idealized B-form DNA double helix generated using 3DNA (112). (c) Schematic representation of the base open state of A-T bps, which is implicated in DNA-protein recognition and damage repair.

For example, in the DNA double helix, Watson-Crick G-C and A-T base pairs (bps) exist in dynamic equilibrium with a minor Hoogsteen conformation, with populations on the order of 0.1-1.0% (7, 8), corresponding to an energetic cost of 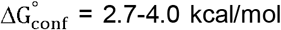. However, the Hoogsteen bp can become the major conformation when duplex DNA is bound by proteins such as p53 (9)(Fig. 1b). Here, energetically favorable intermolecular contacts make up for the conformational penalty to convert the bp from Watson-Crick to the minor Hoogsteen conformation. Indeed, DNA modifications that increase the thermodynamic propensities to form Hoogsteen bps have been shown to improve p53-DNA binding affinity (4, 10). Changes in the structures of RNA and DNA are ubiquitously observed upon binding to proteins and other partners (11, 12). In all cases, the binding energetics will have to make up for the relevant conformational penalties.

Conformational preferences, and therefore conformational penalties, can vary depending on sequence and structural contexts, due to epigenetic and post-transcriptional modifications as well as damage, and due to changes in physiological conditions such as pH, temperature, salt composition and concentration. This can, in turn, help shape the specificities of biochemical processes.

For example, base pairs in DNA can spontaneously open (Fig. 1c)(13, 14), and when bound to damage repair proteins, can adopt extra-helical conformations (15). The thermodynamic preferences to form the base open and extra-helical conformations have been shown to vary with base modifications and sequence context, in turn determining the binding affinity (15) and catalytic efficiency (16) of glycosylase enzymes, which excise damaged bases from DNA, as well as the processivity of helicases, which unwind DNA (17). DNA bps near mismatches have been shown to have increased propensities to form Hoogsteen bps, and Hoogsteen bps are often observed in non-canonical regions of DNA in crystal structures of DNA-protein complexes (18, 19). In addition, wobble G•T/U mismatches can form Watson-Crick-like conformations through tautomerization and ionization of the bases (20, 21). The preferences to form Watson-Crick-like conformations can vary with pH and chemical modifications, in turn shaping the probability of misincorporating G•T bps during DNA replication (20), and providing a basis for recoding translation (22–24).

In RNA, sequence-specific preferences to adopt alternative conformations determine the energetics of RNA-RNA tertiary assembly (25, 26) and RNA-protein binding affinities (27). In addition, pH and temperature-dependent preferences to adopt alternative secondary structures can help control mRNA translation (28, 29) and the efficiency of enzymatic processing of micro RNAs (30). By increasing the thermodynamic preferences to form single-stranded RNA, the epitranscriptomic modification *N*^6^-methyladenosine (31) has been shown to improve the binding of single-stranded RNA binding proteins to modified RNAs (32). Aberrant modulation of RNA conformational preferences due to mutations have also been linked to disease (reviewed in (5)).

Despite their importance in shaping the specificity of biochemical reactions involving nucleic acids, measuring conformational preferences in DNA and RNA and how they vary with sequence, structural and chemical contexts, as well as physiological conditions, remains a significant challenge in biophysics and structural biology. This is because measuring the energetic cost to form non-native conformations requires accurately determining their low abundance in a sea of other conformations in the apo-ensemble. A variety of approaches have been used to measure such conformational equilibria, including Nuclear Magnetic Resonance (NMR) (33), Electron Paramagnetic Resonance (EPR) (34) and Infrared (IR) spectroscopy (35), Fluorescence Resonance Energy Transfer (FRET) (36), Cryo-EM and X-ray crystallography (37), and chemical probing (38). However, with the exception of NMR, most of these techniques are not applicable to conformations such as Hoogsteen bps and the base open states, which have populations <1%, are short-lived with lifetimes shorter than a few milliseconds, and which involve localized bp rearrangements. Most of these approaches are also technically demanding, often requiring synthesis of molecules with specialized labels, with NMR in particular requiring large sample quantities. Thus, they do not lend themselves to comprehensively exploring the conformational preferences of nucleic acids. As a result, the thermodynamic preferences to form Hoogsteen or base open states have yet to be measured for all sixteen trinucleotide sequence contexts. Therefore, there is a need for alternative approaches that can more readily measure conformational preferences in nucleic acids as well as proteins.

Prior NMR studies have shown that chemical modifications and mutations can render minor non-native conformations (Fig. 2a) the major state (population >90%) in proteins (39) and nucleic acids (40), thereby enabling their in-depth structural as well as functional characterization (41). Here, we show that by combining such chemical modifications (Fig. 2b), with melting experiments that measure folding energetics, it is feasible to measure the thermodynamic preferences to form minor non-native conformations in RNA and DNA. We used UV spectroscopy to measure melting energetics (42, 43)(Fig. 2c) given its simplicity, lack of requirement for specific labeling, and reduced sample requirements relative to other approaches (44–46).

**Figure 2.**
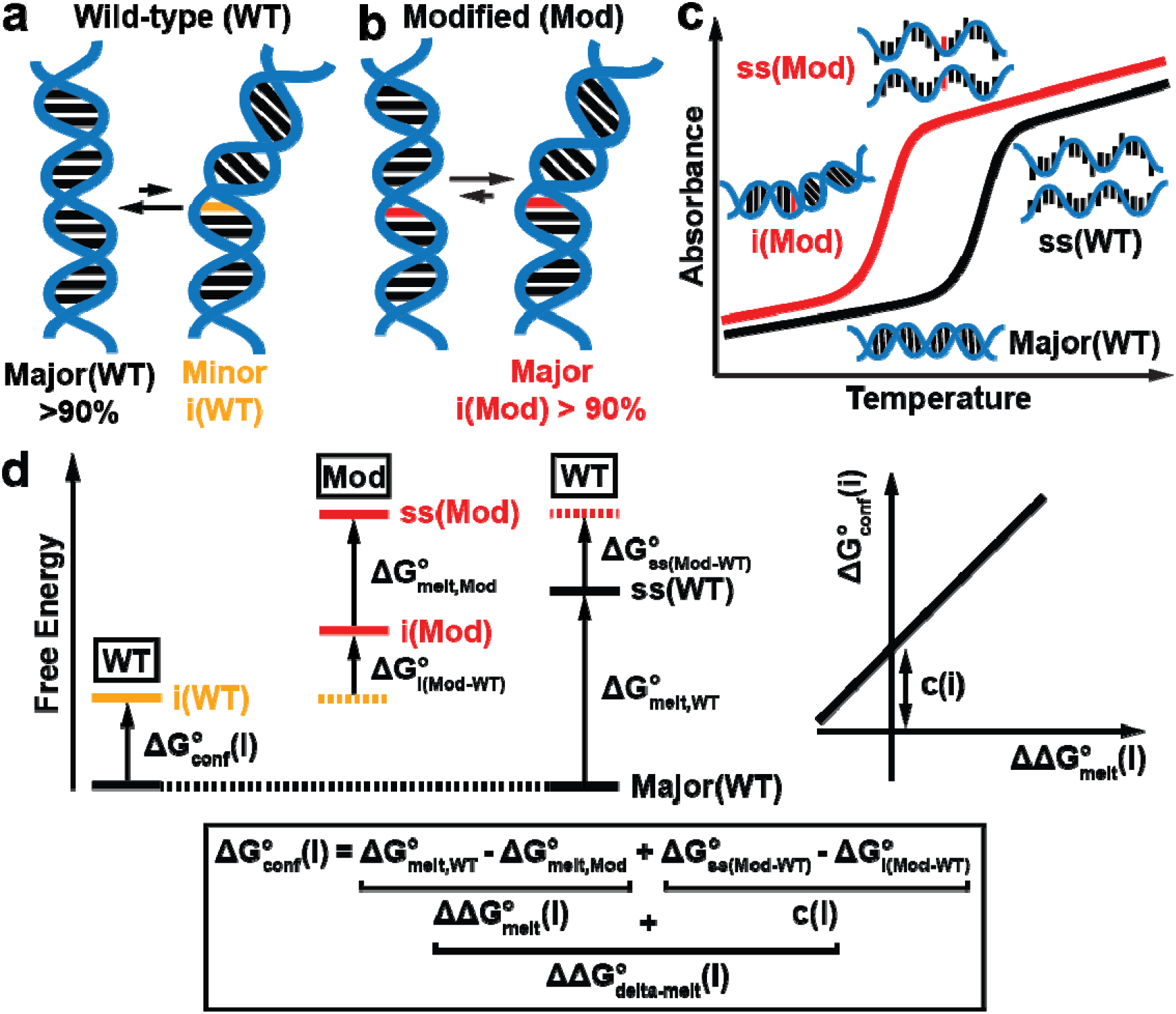
Conceptual underpinnings of delta-melt for measuring thermodynamic preferences to adopt non-native conformational states in nucleic acids. (a) Dynamic equilibrium in the unmodified wild-type (WT) nucleic acid molecule involving a major and the i^th^ minor non-native conformation (i(WT)) involving a change in the conformation of a bp (in orange). (b) Chemical modifications (red, Mod) are introduced to bias the conformational ensemble towards the i^th^ non-native conformational state i(WT) of the WT nucleic acid. (c) Schematic diagram showing optical melting experiments on unmodified WT (black) and modified Mod (red) nucleic acids. (d) Free energy diagram of delta-melt (left), relationship between the free energies in delta-melt (bottom) and expected correlation between and (right).

The approach termed ‘delta-melt’ builds on prior studies employing differences in melting energetics to estimate the contribution of motifs (44, 47) or specific functional groups (48, 49) to nucleic acid stability. Such studies provided the basis for nearest neighbor rules and secondary structure prediction algorithms (50, 51). However, in contrast to these prior studies, delta-melt measures the difference in melting energetics between an unmodified (WT) molecule and a molecule chemically modified (Mod) to shift the ensemble toward the desired non-native state (Fig. 2a, b). Converting differences in melting energetics into differences between the thermodynamic stability of the major and minor conformational states requires calibration against independent measurements of conformational preferences using other biophysical techniques. In this study, we extensively verified the robustness of delta-melt for four different non-native conformational states under a variety of physiological conditions and sequence contexts through independent measurements of thermodynamic preferences using NMR.

We obtained rare insights into conformational cooperativity using delta-melt, estimating that tandem Hoogsteen bps form cooperatively by 1.0-2.5 kcal/mol in DNA. We also measured the thermodynamic preferences to form G-C^+^ and A-T Hoogsteen, and A-T base open states for nearly all sixteen trinucleotide sequence contexts, and found distinct sequence-specific variations on the order of 2-3 kcal/mol, corresponding to ~10-100 fold variation in the population of the minor conformational state. These results suggest that the DNA double helix codes for a rich layer of sequence-specific conformational preferences, which could in turn tune the specificity of DNA biochemical transactions. Thus, melting experiments can be used to access thermodynamic information regarding regions of the free energy landscape of biomolecules beyond the native folded and unfolded conformations, opening the door to more systematic explorations of how the ensemble varies with sequence and structural contexts, chemical modifications as well as physiological conditions.

## Results

### Conceptual underpinnings of delta-melt

The conceptual underpinnings of delta-melt can be understood by using a free energy diagram (Fig. 2d). The desired thermodynamic preference, 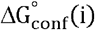, in the wild-type molecule to form the i^th^ non-native conformational state is given by the free energy difference between the i^th^ non-native (i(WT)) and major (Major(WT)) conformations (Fig. 2a),

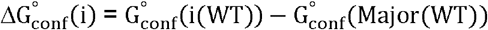

In principle, 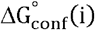 could be obtained from the difference in the melting energetics 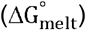 between the Major(WT) and i(WT) conformational states in the wild-type molecule:

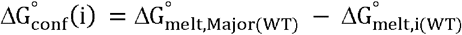

In practice however, because the minor i(WT) conformation is lowly populated, its melting energetics are difficult to measure experimentally (52, 53). To address this limitation, we introduce a chemical modification that renders i(WT) the dominant conformation in the modified (Mod) molecule, typically by destabilizing the major state relative to the minor state (Fig. 2b). The degree to which the modification will affect the melting energetics relative to the WT molecule will depend on the desired thermodynamic preference to form the minor conformation 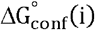 and also on how the modification affects the energetics of the minor conformation and single-strand (ss)(Fig. 2d). Therefore, the desired 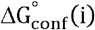 can be obtained from the difference between the melting energetics of WT and Mod molecules, 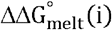, plus an offset, c(i), which takes into account the difference in free energy to modify ss relative to i(WT)(Fig. 2d):

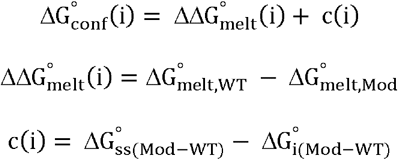

Thus, 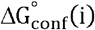 can be estimated using a pair of melting experiments, provided that the offset c(i) is known. The value of c(i) can be determined experimentally provided that it does not vary across different sequence contexts and/or environmental conditions (changes in temperature, pH, and salt) of interest, by calibrating 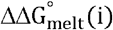 data against 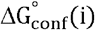 values independently measured via biophysical techniques such as NMR for a few experimental conditions and/or sequence contexts (Fig. 2d). c(i) can be obtained as the intercept from a linear fit of 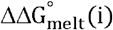 versus 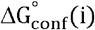 (Fig. 2d). Such calibrations with resource intensive measurements are routinely performed in other contexts such as micro-array-based binding measurements, which are calibrated against ITC/EMSA experiments to enable measurement of binding affinities in high throughput (4, 54). The calibration enables us to take into consideration the effect of the modification on the energetics of the minor and ss states. This is in contrast to prior studies employing melting experiments to estimate the energetic contribution of specific motifs to nucleic acid stability (44, 47–49, 55), in which it is typically assumed that modifications do not impact the ss energetics (i.e. 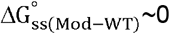). Following calibration to estimate c(i), 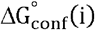 for a new sample can be estimated using a pair of melting experiments using,

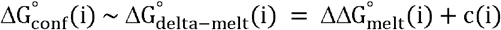

One of the advantages of delta-melt is that in theory, there is no limit on the population and lifetime of the minor conformation that can be studied as long as it can be stabilized using a modification. It should be applicable to conformational states that fall outside NMR detection, and we will encounter such an example in this study (*vide infra*). Likewise, in theory, there is no size limit provided that the site-specifically modified nucleic acid can be synthesized (56). However, deviations from two-state melting behavior are more likely in large nucleic acid molecules, and this can complicate analysis of the UV melting data, as observed even for some of the short duplexes studied here (*vide infra*). In such cases, the melting experiments have to be supplemented with information from other techniques to understand the structural basis behind changes in absorbance (reviewed in (57)). In addition, delta-melt does not provide any information regarding the kinetics of interconversion or structure of the non-native state. It is also assumed that the modification robustly renders the desired i^th^ minor conformation the major state (>90%) robustly across different contexts and experimental conditions. If the modification confers a weaker bias, delta-melt may underestimate the true value of 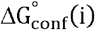.

### Testing delta-melt on A-T Hoogsteen base pairs in duplex DNA

We initially performed a set of experiments to test the underlying assumptions and general utility of delta-melt to measure conformational preferences to form four different minor non-native conformations in DNA and RNA. As a first test, we used delta-melt to measure the preferences to form A-T Hoogsteen bps (7) in duplex DNA. The thermodynamic preferences to form A-T Hoogsteen bps in duplex DNA are of interest because they are proposed to determine DNA-protein binding affinities (4, 10) and the propensity of DNA damage induction (58).

To form a Hoogsteen bp, the purine base in a Watson-Crick bp has to flip 180° about the glycosidic bond to adopt a *syn* conformation, and the DNA backbone has to constrict by ~2 Å to allow hydrogen bonding between the bases (Fig. 3a)(59, 60). We used *N*^1^-methylated adenine (m^1^A^+^), a positively charged naturally occurring form of alkylation damage, to substantially bias (> 90%) the conformation of the A-T bp toward Hoogsteen, as shown previously (7, 61). The modification destabilizes the Watson-Crick A-T bp both by disrupting a hydrogen bond and through steric collisions (Fig. 3a)(7, 61). Prior NMR studies showed m^1^A^+^-T to be an excellent structural mimic of the transient A-T Hoogsteen bps in unmodified DNA (62). As m^1^A^+^-T is a good structural mimic of the transient A-T Hoogsteen bp, it may also serve as a good energetic mimic for delta-melt experiments.

**Figure 3.**
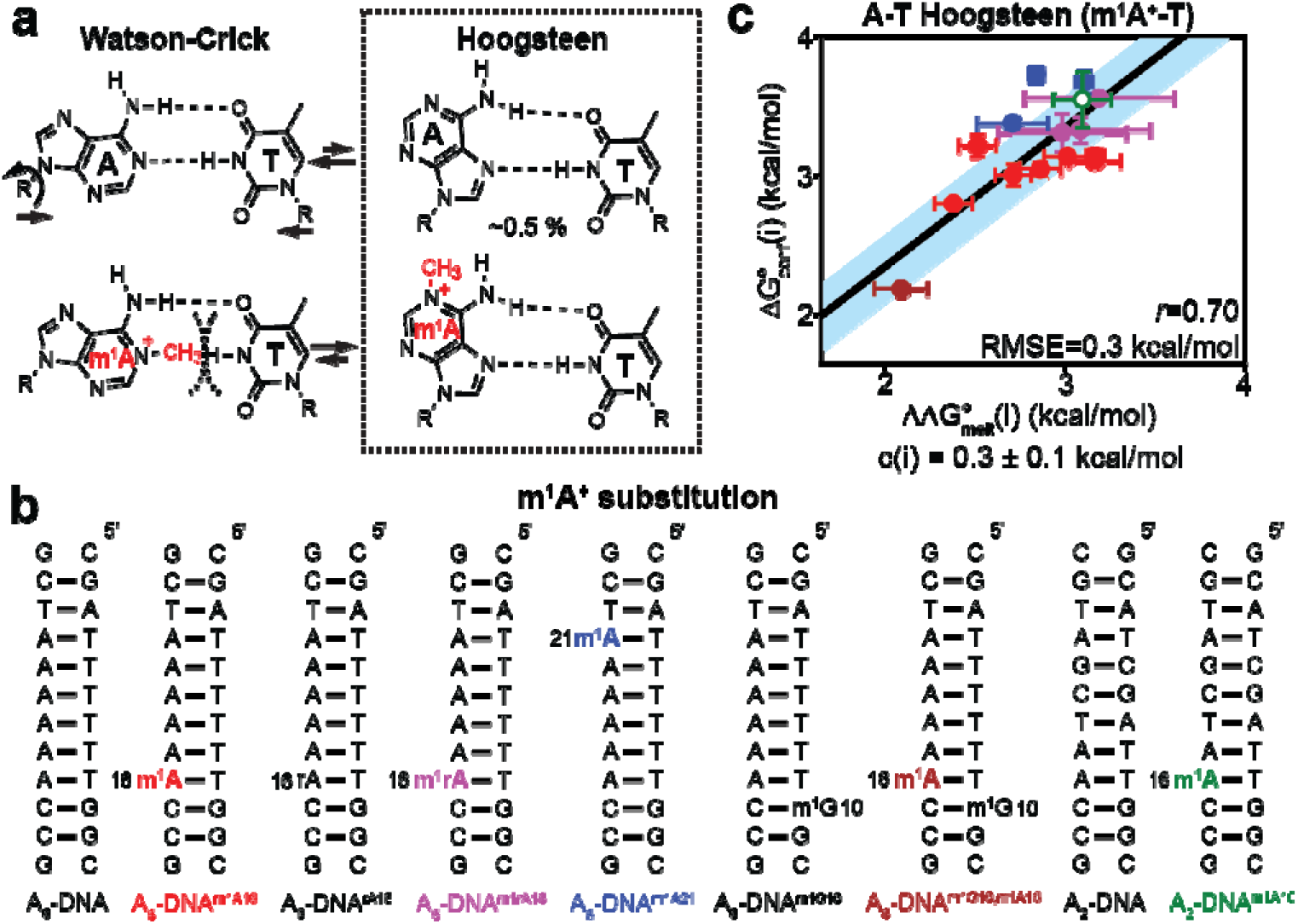
Testing delta-melt on A-T Hoogsteen bps in DNA using the *N*^1^-methyladenosine mimic. (a) In duplex DNA, A-T Watson-Crick bps exist in dynamic equilibrium with minor A-T Hoogsteen bps. Population of the Hoogsteen bp was obtained as described previously (8). *N*^1^-methylation of adenine (m^1^A^+^, red) biases the equilibrium to favor the Hoogsteen bp. Steric clashes are indicated using dashed lines. (b) DNA duplexes used in UV melting experiments with and without m^1^A^+^ substitution (in bold). rA corresponds to ribo adenosine. (c) Correlation plot between for formation of A-T Hoogsteen bps (Table S5) and obtained from melting measurements on m^1^A^+^-T/A-T bp containing DNA (Tables S1-S2). Points are color-coded according to duplex (panel b). Error bars for NMR and delta-melt measurements were obtained using a Monte-Carlo scheme (106), and by propagating the uncertainties from UV melts, respectively, as described in Methods. Shown are the Pearson’s correlation coefficient (*r*) and root mean square error (RMSE)(Methods). Blue shaded region denotes the estimate of the error of linear regression obtained using Monte-Carlo simulations, while open symbols denote data derived from weak RD profiles (Methods).

We analyzed sequences shown previously to form stable duplexes in solution and bp positions for which the Watson-Crick to Hoogsteen exchange had previously been characterized using NMR relaxation dispersion (RD)(7, 62, 63)(Fig. 3b). This included A-T bps in two different tri-nucleotide sequences (5’-CAA-3’/5’-AAT-3’) embedded in three different duplexes in two different positional contexts. These data were supplemented with additional NMR RD and corresponding delta-melt experiments under varying buffer and temperature conditions (pH = 4.4-6.9, T = 10-30°C, and 25-150 mM NaCl)(Fig. 3b, Fig. S1-S2 and S4-S6, Tables S1-S5). Additional measurements were performed when chemically modifying the Hoogsteen bp with ribo-guanosine or its neighbor using *N*^1^-methyl guanosine, both of which are known to perturb the thermodynamic propensity to form Hoogsteen bps (63)(Fig. 3b). This yielded a total of fourteen datapoints which could be used to test delta-melt.

All delta-melt experiments were performed using the same buffer conditions used in the NMR RD experiments (Methods). The ‘curve-fitting’ approach (64, 65) was used to extract the free energy of melting 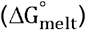 from the UV melting curves for all samples (64, 65). The approach assumes two-state melting behavior and is commonly used in the literature (64, 65). A more rigorous approach evaluates the two-state assumption using concentration-dependent melting experiments to extract thermodynamic parameters (42, 43, 66). Although concentrationdependent measurements help gauge the two-state behavior and are also required to determine the enthalpy 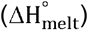 and entropy 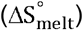 of melting more accurately, this substantially increases the sample requirements and reduces the throughput of the experiment. In addition, the concentration-dependence is a necessary but insufficient condition for two-state melting (44). As we were primarily interested in free energy of melting 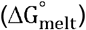, which can be measured accurately using curve-fitting since the errors in the enthalpy and entropy typically cancel out (42), and since we have already previously verified using NMR that some of the duplexes studied here (A_2_ and A_6_-DNA)(67) exhibit two-state melting behavior, we favored using the curve-fitting approach in the application of delta-melt. For a more elaborate discussion regarding the UV melting experiments and the assumptions used in the analysis, the reader is referred to Discussion S1.

Indeed, a reasonable correlation (*r*=0.70) was observed between 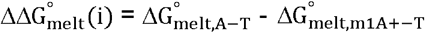 (Tables S1-S2) and 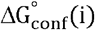 measured using NMR RD (Fig. 3c and Table S5). The correlation could be linearly fit (*r*=0.7, Fig. 3c) with an RMSE of 0.3 kcal/mol and with c(i) ~ 0.3 kcal/mol. The quantitative agreement between the delta-melt energetics and those derived using NMR RD suggest that the offset c(i) corresponding to the difference in energetic cost to methylate adenine-N1 in the ss versus the A-T Hoogsteen bp (Discussion S2) does not vary substantially across the sequence and chemically modified contexts and buffer conditions examined, and that any variations remain small relative to the differences in 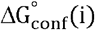 (~ 2 kcal/mol). These results therefore also indicate that delta-melt can be used to measure the thermodynamic preferences to form A-T Hoogsteen bps as a function of experimental conditions.

Nevertheless, the correlation was not ideal and m^1^A^+^-T is an imperfect mimic of the A-T Hoogsteen bp as the methyl group introduces a positive charge to the adenine base. We therefore also tested neutral modifications such as C-T and T-T mismatches, which were recently shown to mimic the A-T Hoogsteen bp based on NMR chemical shifts analysis (4)(Fig. 4b). As in the A-T Hoogsteen bp, the two bases in C-T and T-T mismatches also have to come into closer proximity relative to Watson-Crick bps to hydrogen bond thus mimicking this key feature of the Hoogsteen conformation (62)(Fig. 4a). Mismatches also have the advantage that they can be readily installed when synthesizing DNA. These experiments also assessed the robustness of delta-melt when using different modifications to mimic the minor conformation.

**Figure 4.**
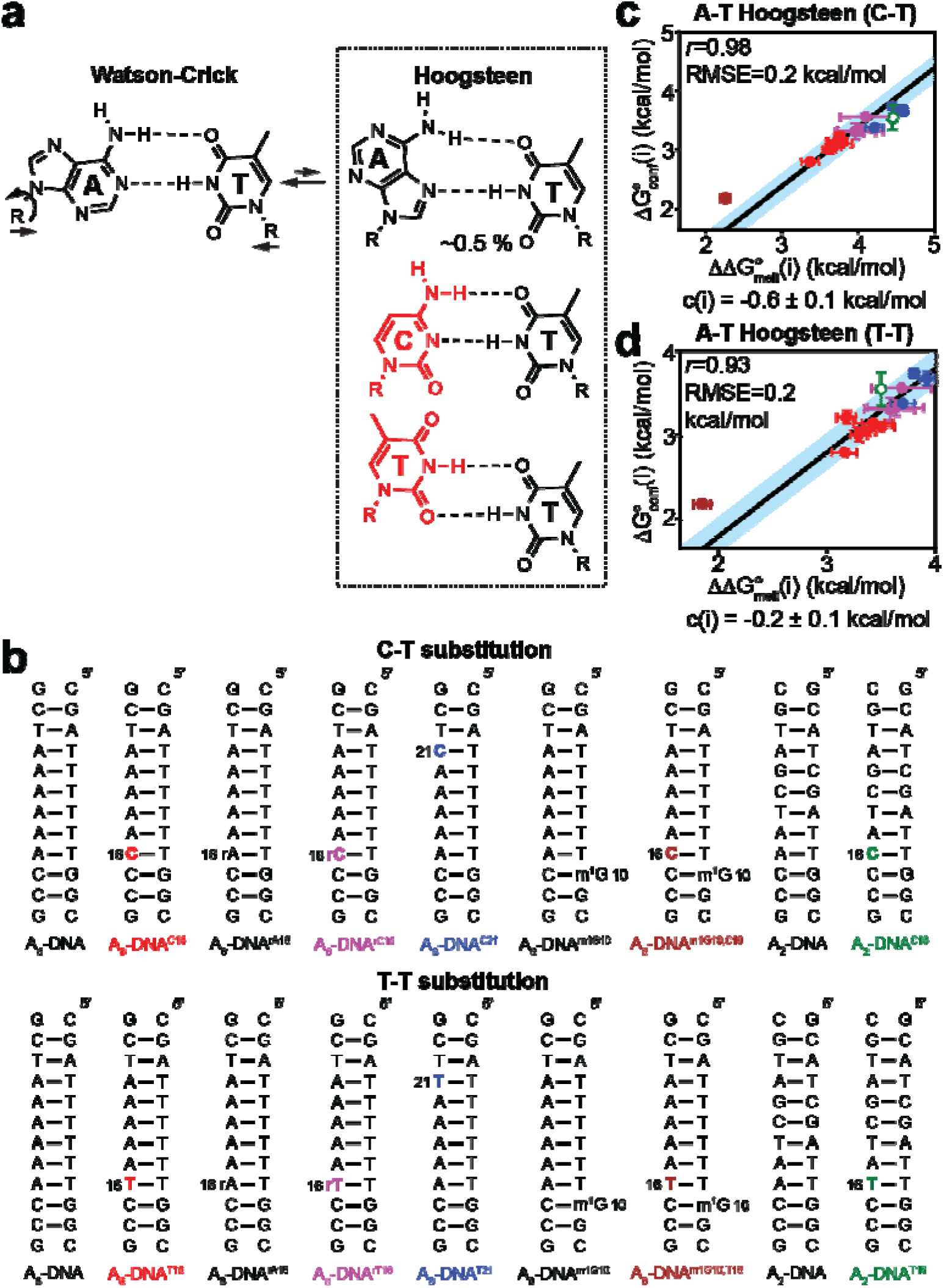
Testing delta-melt on A-T Hoogsteen bps in DNA using C-T and T-T mismatches as Hoogsteen mimics. (a) C-T and T-T mismatches mimic the conformation of the A-T Hoogsteen bp. (b) Duplexes used in UV melting experiments with and without C-T and T-T mismatches (in bold). Correlation plot between obtained from melting experiments on C-T/A-T (c) and T-T/A-T bp containing DNA (Table S2) and for formation of A-T Hoogsteen bps as determined using NMR RD (Table S5). Points in panels c and d are color-coded according to the corresponding duplex in panel b. Error bars for NMR and delta-melt measurements were obtained using a Monte-Carlo scheme (106), and by propagating the uncertainties from UV melts, respectively, as described in Methods. Shown are the Pearson’s correlation coefficient (*r*) and root mean square error (RMSE)(Methods). Blue shaded region denotes the estimate of the error of linear regression obtained using Monte-Carlo simulations, while open symbols denote data derived from weak RD profiles (Methods).

Indeed, an improved correlation (r=0.98, and 0.93) was observed between 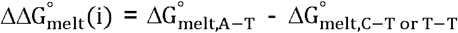 for both C-T and T-T mismatches (Fig. 4c and d, Table S2) and 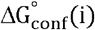 to form A-T Hoogsteen bps as measured using NMR RD (Table S5). The data could be fit with RMSE = 0.2 kcal/mol for both C-T and T-T (Fig. 4c and d, Discussion S2). Nevertheless, the deviation seen for the A16-T9 bp near the m^1^G10-C15^+^ Hoogsteen bp in A_6_-DNA^m1G10^ (brown point in Fig. 4c,d), indicates that C-T/T-T mismatches may not be good energetic mimics of A-T Hoogsteen bp when neighboring another Hoogsteen bp, possibly because they do not capture unique stacking interactions involving adjacent *syn* purine base. Indeed, this deviation was not observed when using m^1^A^+^ (Fig. 3c).

Taken together, the above results establish m^1^A^+^-T, C-T and T-T bps to be good energetic mimics of A-T Hoogsteen bps and support the underlying assumptions used in delta-melt to measure the thermodynamic preferences to form A-T Hoogsteen bps.

### Testing delta-melt on G-C^+^ Hoogsteen base pairs in duplex DNA

Next, we tested the utility of delta-melt to measure the thermodynamic preferences to form protonated G-C^+^ Hoogsteen bps in duplex DNA. Like A-T Hoogsteen bps, G-C^+^ Hoogsteen bps can form through rotation of the guanine base about the glycosidic bond and constriction of the two partner bases to form Hoogsteen hydrogen bonds (Fig. 5a). As G-C^+^ Hoogsteen bps entail the protonation of cytosine, their thermodynamic preferences strongly vary with pH (68), providing a unique opportunity to test the utility of delta-melt in measuring pH-dependent conformational preferences. G-C^+^ Hoogsteen bps have been proposed to contribute to DNA recognition by proteins such as TBP (69) and to DNA damage induction (70).

**Figure 5.**
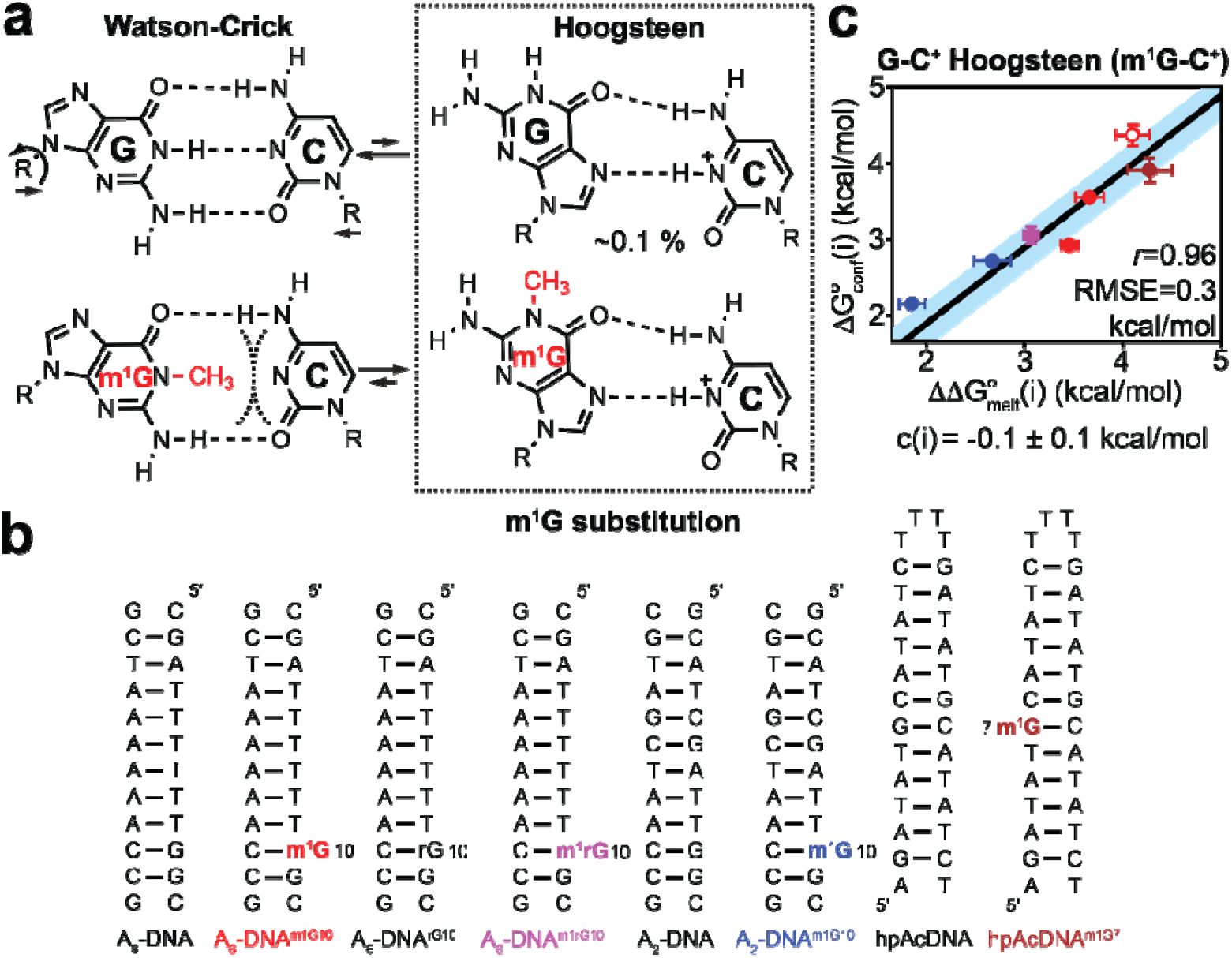
Testing delta-melt on G-C^+^ Hoogsteen bps. (a) In duplex DNA, G-C Watson-Crick bps exist in dynamic equilibrium with a minor G-C^+^ Hoogsteen bp. Population of the minor Hoogsteen bp was obtained as described previously (8). m^1^G (red) biases the Watson-Crick to Hoogsteen equilibrium of G-C bps towards the Hoogsteen conformation via loss of a hydrogen bond donor (G-N1), and due to steric clashes (curved black dashes). (b) Duplexes used in UV melting experiments with and without m^1^G substitution (indicated in bold). (c) Correlation plot between obtained from melting experiments on m^1^G-C/G-C bp containing DNA (Fig. S1, Table S1) and measured using NMR for formation of G-C^+^ Hoogsteen bps (Table S5). Points are color-coded according to the duplexes in panel b. Error bars for NMR and delta-melt measurements were obtained using a Monte-Carlo scheme, and by propagating the uncertainties from UV melts, respectively, as described in Methods. Shown are the Pearson’s correlation coefficient (*r*) and root mean square error (RMSE)(Methods). Blue shaded region denotes the estimate of the error of linear regression obtained using Monte-Carlo simulations, while open symbols denote data derived from weak RD profiles (Methods).

As for the A-T Hoogsteen bp, we used *N*^1^-methyl guanosine (m^1^G), another naturally occurring form of alkylation damage, to bias the conformation of the G-C bp towards Hoogsteen (7)(Fig. 5a). Prior NMR studies showed m^1^G-C^+^ to be an excellent structural mimic of the transient G-C Hoogsteen bps in unmodified DNA (62). As m^1^G is a neutral modification, it is potentially a better Hoogsteen mimic relative to m^1^A^+^.

To test delta-melt on G-C^+^ Hoogsteen bps, we chose sequences shown previously to form stable duplexes in solution and bp positions for which transient Hoogsteen bps have been previously characterized using NMR RD (7, 62, 63)(Fig. 5b). We also performed additional NMR RD and corresponding delta-melt experiments under different salt and pH conditions (pH 4.4-6.9, T = 25 °C, and 25-150 mM NaCl), yielding in total seven points for comparison, corresponding to two different tri-nucleotide sequence contexts (5’-TGG-3’/5’-TGC-3’) embedded in three different duplexes, at two positional contexts (Fig. 5b, Fig. S1-S2 and S4-S6 and Tables S1-S5).

UV melting experiments were used to measure the energetics of duplex melting with and without m^1^G substitution at these positions under the same buffer conditions used for NMR. An excellent correlation (*r*=0.96) was observed between the difference in melting energetics 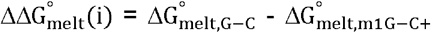 and 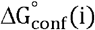 measured using NMR RD (Fig. 5c, Tables S1 and S5). The correlation could be fit with an RMSE of 0.3 kcal/mol (Fig. 5c) and c(i) ~ −0.1 kcal/mol (Discussion S2). The improved delta-melt correlation observed with m^1^G-C^+^ (*r*=0.96) relative to the m^1^A^+^-T bp (*r*=0.70, Fig. 3c) could be because m^1^G does not introduce a positive charge.

These results support the assumption that c(i) (Discussion S2) does not vary substantially across the sequence and chemically modified contexts, and buffer conditions examined and indicate that delta-melt can be used to measure the thermodynamic preferences to form G-C^+^ Hoogsteen bps as well.

### Testing delta-melt on base open A-T states in duplex DNA

One of the advantages of delta-melt is that it can potentially be applied to low-populated and short-lived states falling below detection methods of conventional biophysical methods. We therefore tested its utility on the A-T base open state (Fig. 1c), which based on NMR measurements of imino proton exchange rates (14), has an exceptionally low population of ~0.001% and short lifetime on the order of ~0.1 μs (Fig. 6a). Compared to G-C^+^ and A-T Hoogsteen bps, the base open state is disfavored by ~4 kcal/mol, providing an interesting test for the general applicability of delta-melt on an exceptionally low-populated conformational state that cannot be detected even by NMR RD methods.

**Figure 6.**
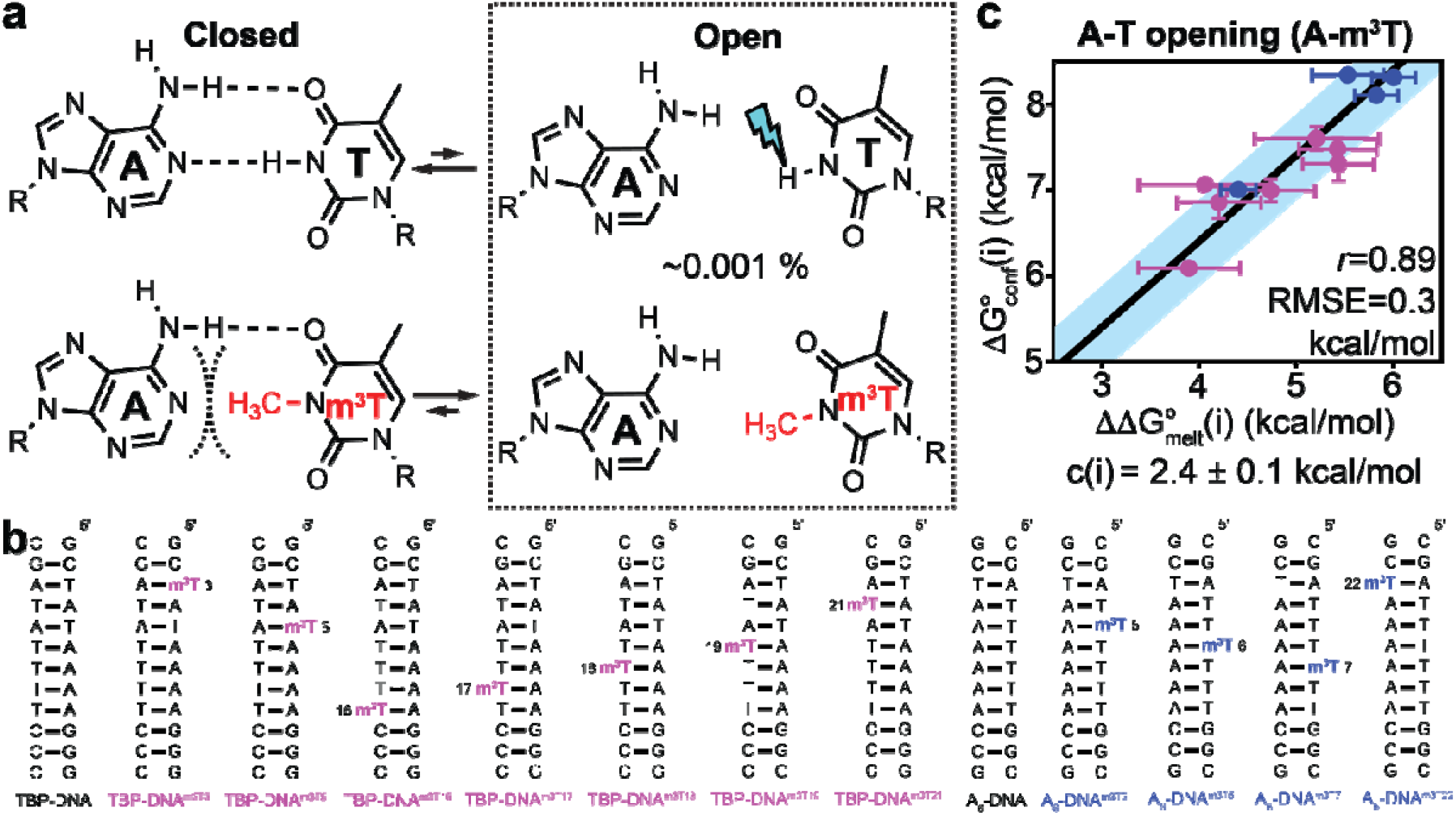
Testing delta-melt on the open state of A-T bps. (a) Exchange between closed and open A-T Watson-Crick bps in DNA. The T-H3 imino proton in the open state is susceptible to solvent exchange (blue thunder). m^3^T is hypothesized to bias the ensemble to mimic the base open state by disrupting the A(N1)---H3-(N3)T hydrogen bond, and by sterically (curved dashed lines) disfavoring the closed bp. (b) Duplexes used in UV melting experiments with and without m^3^T substitution (in bold). (c) Correlation plot between for formation of open A-T bps (Table S8) and obtained from melting experiments on A-m^3^T/A-T bp containing DNA (Tables S1-S2). Points are color-coded according to the duplex that they correspond to (panel b). Error bars for NMR and delta-melt measurements were obtained using a Monte-Carlo scheme, and by propagating the uncertainties from UV melts, respectively, as described in Methods. Shown are the Pearson’s correlation coefficient (*r*) and root mean square error (RMSE)(Methods). Blue shaded region denotes the estimate of the error of linear regression obtained using Monte-Carlo simulations (Methods).

Based on imino proton solvent exchange experiments (71), the base open state entails the loss of the A(N1)---H3-(N3)T hydrogen bond. Thus, we hypothesized that substituting T by *N*^3^-methyl thymine (m^3^T), so that the thymine imino proton is replaced by a methyl group, should disrupt the A(N1)---H3-(N3)T hydrogen bond, and bias the ensemble towards the base open state (71). Indeed, NMR spectra (Fig. S7) revealed that the modification disrupted the A(N1)---H3-(N3)T hydrogen bond in the modified bp with m^3^T adopting an intra-helical conformation.

We analyzed sequences shown previously to form stable duplexes in solution and bp positions for which A-T base opening has been previously characterized using NMR (TBP-DNA, Fig. 6b)(72). We also performed additional imino proton exchange NMR and corresponding delta-melt experiments for another duplex (A_6_-DNA) under different buffer and temperature conditions, yielding in total eleven points for comparison, corresponding to six different trinucleotide sequence contexts (5’-TAG-3’/5’-TAT-3’/5’-AAG-3’/5’-AAA-3’/5’-TAA-3’/5’-GAT-3’) in two different duplexes and a range of buffer conditions (pH 8.0-8.8, T = 15-25 °C, 100 mM NaCl)(Fig. 6b, Fig. S1-S2, S4 and S8, and Tables S1-S2 and S6-S8).

UV melting experiments were used to measure the energetics of duplex melting with and without m^3^T substitutions at these positions under the NMR buffer conditions (Fig. S1-S2 and Tables S1-S2). Interestingly, a very good correlation (*r*=0.89) was observed between the difference in melting energetics 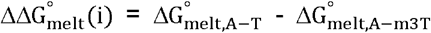 and 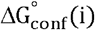 for base opening measured using NMR imino proton exchange measurements (Fig. 6c, Table S1-S2 and S8)(72). The correlation could be fit with RMSE of 0.3 kcal/mol and c(i) ~2.4 kcal/mol (Fig. 6c)(Discussion S2). These results demonstrate the utility of delta-melt to measure the thermodynamic preferences for A-T base opening across different sequence contexts and buffer conditions, and also establish A-m^3^T as a mimic of the base open state which has been recalcitrant to experimental structural characterization.

### Testing delta-melt on isomerization of the *N*^6^-methylamino group in duplex RNA

We also examined the utility of delta-melt to address more subtle conformational rearrangements involving rotation of a single methylamino group. *N*^6^-methyladenosine (m^6^A) is a highly abundant RNA modification that plays roles in virtually all aspects of mRNA metabolism (31, 73). When paired with uridine, the methylamino group in m^6^A isomerizes between *anti* (major) and *syn* (minor) conformations resulting in the loss of a hydrogen-bond (Fig. 7a)(74). This conformational change has been proposed to play roles slowing a variety of biochemical processes that involve duplex melting and annealing (67).

**Figure 7.**
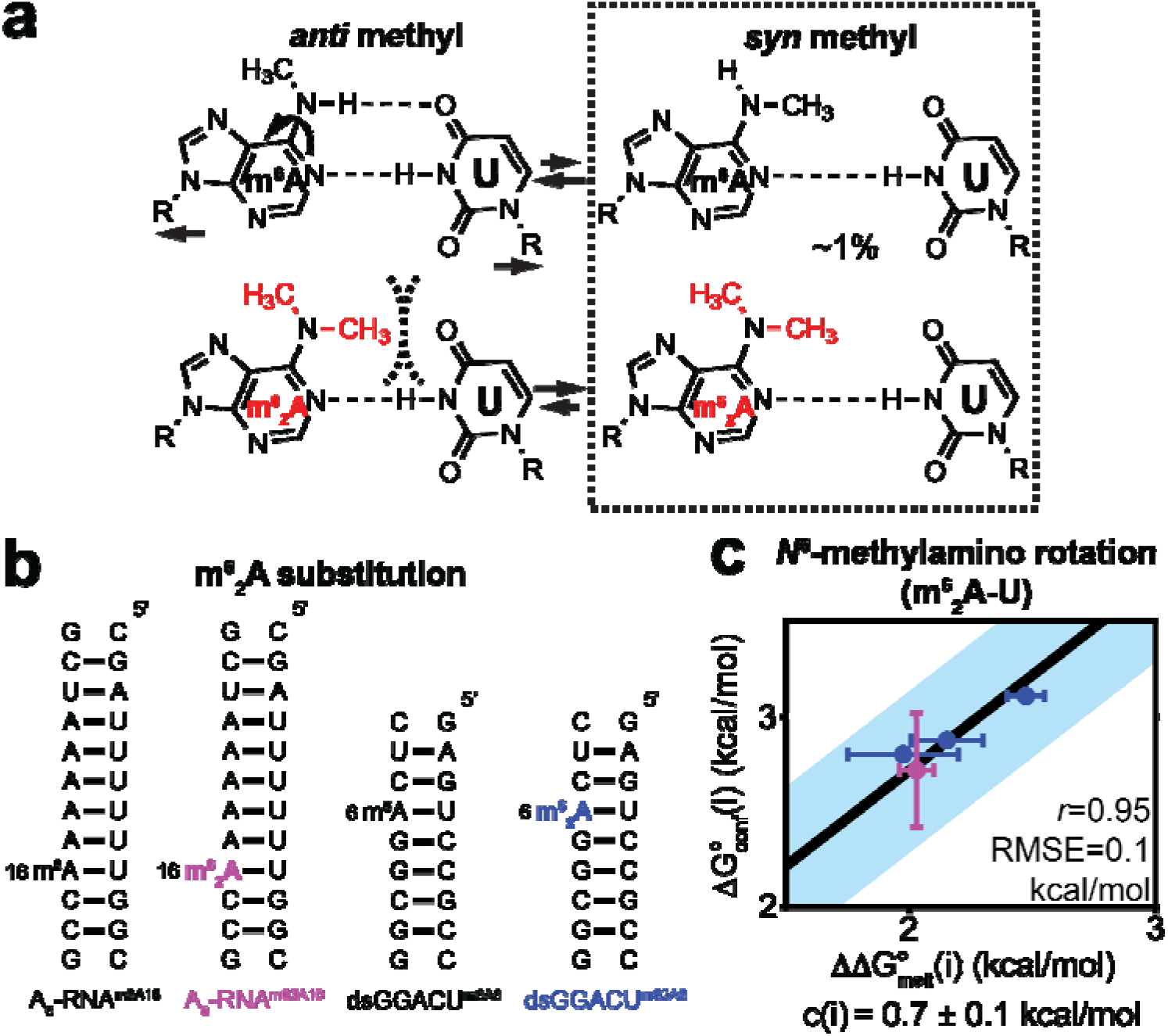
Testing delta-melt on the isomerization of the *N*^6^-methylamino group in m^6^A-U base pairs in RNA. (a) (Top) Equilibrium between major *anti* and minor *syn* conformations of the *N*^6^-methylamino group in m^6^A-U bps in RNA. (Bottom) m^6^_2_A mimics the *syn* methylamino group by sterically disfavoring (black dashes) the conformation with the *anti* methyl. (b) Duplexes used in UV melting experiments with m^6^A and m^6^_2_A (indicated in bold) modifications. (c) Correlation plot between obtained from melting experiments on m^6^A-U/m^6^_2_A-U bp containing RNA (Table S1) and measured using NMR RD for formation of m^6^A(*syn*)-U bps (Table S5). Points are color-coded according to the duplexes in panel b. Error bars for NMR and delta-melt measurements were obtained using a Monte-Carlo scheme (106), and by propagating the uncertainties from UV melts, respectively, as described in Methods. Shown are the Pearson’s correlation coefficient (*r*) and root mean square error (RMSE) (Methods). Blue shaded region denotes the estimate of the error of linear regression obtained using Monte-Carlo simulations.

Using NMR CEST experiments (75), it was recently shown that the minor non-native m^6^A(*syn*)-U bp transiently forms with a population of ~1% (74). It was also shown that *N*^6^,*N*^6^-dimethyladenosine (m^6^_2_A), a post transcriptional modification in tRNA (76), structurally mimics this non-native m^6^A(*syn*)-U bp (74). This modification destabilizes the major m^6^A(*anti*)-U conformation both by disrupting the A(N6)-H6---(O4)T hydrogen bond and through steric collisions (curved dashed lines, Fig. 7a).

We tested delta-melt on two sequences shown previously to form stable duplexes in solution, at bp positions for which the non-native m^6^A(*syn*)-U bp has been previously characterized using NMR, and for which the m^6^_2_A modification has been shown to recapitulate the NMR chemical shifts of the non-native m^6^A(*syn*)-U bp (74)(Fig. S4). Delta-melt was used to measure temperature-dependent (pH 6.8, T = 37-65 °C, 25 mM NaCl) thermodynamic preferences to form the non-native m^6^A(*syn*)-U bp at two positions in two duplexes in two different tri-nucleotide sequence and position contexts (5’-CAA-3’/5’-GAC-3’)(Fig. 7b); UV melting experiments were used to measure the energetics of duplex melting with m^6^A and m^6^_2_A modifications at these positions under the NMR buffer conditions (Fig. S1 and S4-S6, Tables S1 and S3-S5).

An excellent correlation (*r*=0.95) was observed between 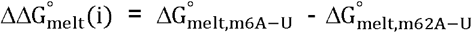 and 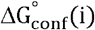 for *N*^6^-methylamino isomerization in m^6^A measured using NMR RD (Fig. 7c)(74). The correlation could be fit with an RMSE of 0.1 kcal/mol and c(i) ~ 0.7 kcal/mol (Fig. 7c)(Discussion S2). These results show that delta-melt can be applied to measure thermodynamic preferences involving isomerization of small chemical groups, as well as broadens its applicability to RNA in addition to DNA.

### Using delta-melt to measure the cooperativity when forming tandem Hoogsteen base pairs in duplex DNA

Having benchmarked delta-melt and tested its underlying assumptions, we applied the approach to gain new insights into the biophysical properties of duplex DNA, which might not be feasible to obtain using more conventional approaches. In this first application, we used delta-melt to examine the cooperativity of forming two adjacent Hoogsteen bps. Tandem Hoogsteen bps are frequently observed in crystal structures of DNA in complex with proteins and drugs, and in the context of DNA lesions that bias the bp conformation to Hoogsteen (reviewed in (77)). Furthermore, crystal structures of certain A-T repeat sequences form helices in which all bps are Hoogsteen (78), again pointing to Hoogsteen bps forming cooperatively. Thus, the thermodynamic preference to form a Hoogsteen bp could be modified by the presence of a nearby Hoogsteen bp, in turn impacting conformational penalties integral to recognition of proteins and other molecules. Such cooperativity could extend to other states such as the base open conformation.

Measuring conformational cooperativity is especially challenging as it requires accurately determining the abundance of exceptionally low-populated conformational states in which two or more bp positions are simultaneously in a minor conformation. The population of tandem Hoogsteen bps is expected to be on the order of the product of the probabilities for forming each Hoogsteen bp (~0.01%). Moreover, such conformations must be distinguished from states in which only one bp is Hoogsteen. Owing to these unique challenges, cooperativity has not been measured to date for any bp rearrangement in nucleic acids.

In theory, delta-melt can be used to measure conformational cooperativity for any arbitrary number of conformational transitions by performing experiments employing the appropriate number of modifications in a duplex. We tested this approach and measured the cooperativity of forming tandem Hoogsteen bps using the validated m^1^A^+^ and m^1^G substitutions to induce Hoogsteen bps at specific positions. In what follows, it will be assumed that tandem *N*^1^-methylated bps are good energetic mimics of their unmodified tandem Hoogsteen counterparts, and that the offset c(i) for tandem Hoogsteen bps is small, similar to isolated Hoogsteen bps (Fig. 3c, 5c)(Discussion S2), or is equal to the sum of the offsets for forming the individual Hoogsteen bps (Methods).

If tandem Hoogsteen bps do form cooperatively in duplex DNA, the energetic cost to form two Hoogsteen bps simultaneously 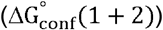 should be smaller than the sum of the energetic cost to form each Hoogsteen bp at each position 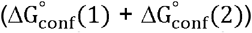 i.e. 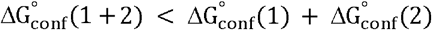. Conversely, 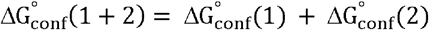 for non-cooperativity and 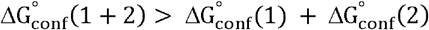 for the case of negative cooperativity.

Thus, subject to the assumptions outlined above, positive cooperativity predicts that the sum of energetic destabilization of single *N*^1^-methylations 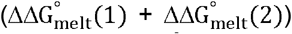 is greater than the energetic destabilization due to tandem *N*^1^-methylation 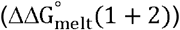 with the difference between the two quantities 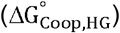 providing a measure of the extent of cooperativity (Fig. 8a),

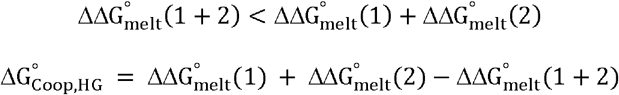

**Figure 8.**
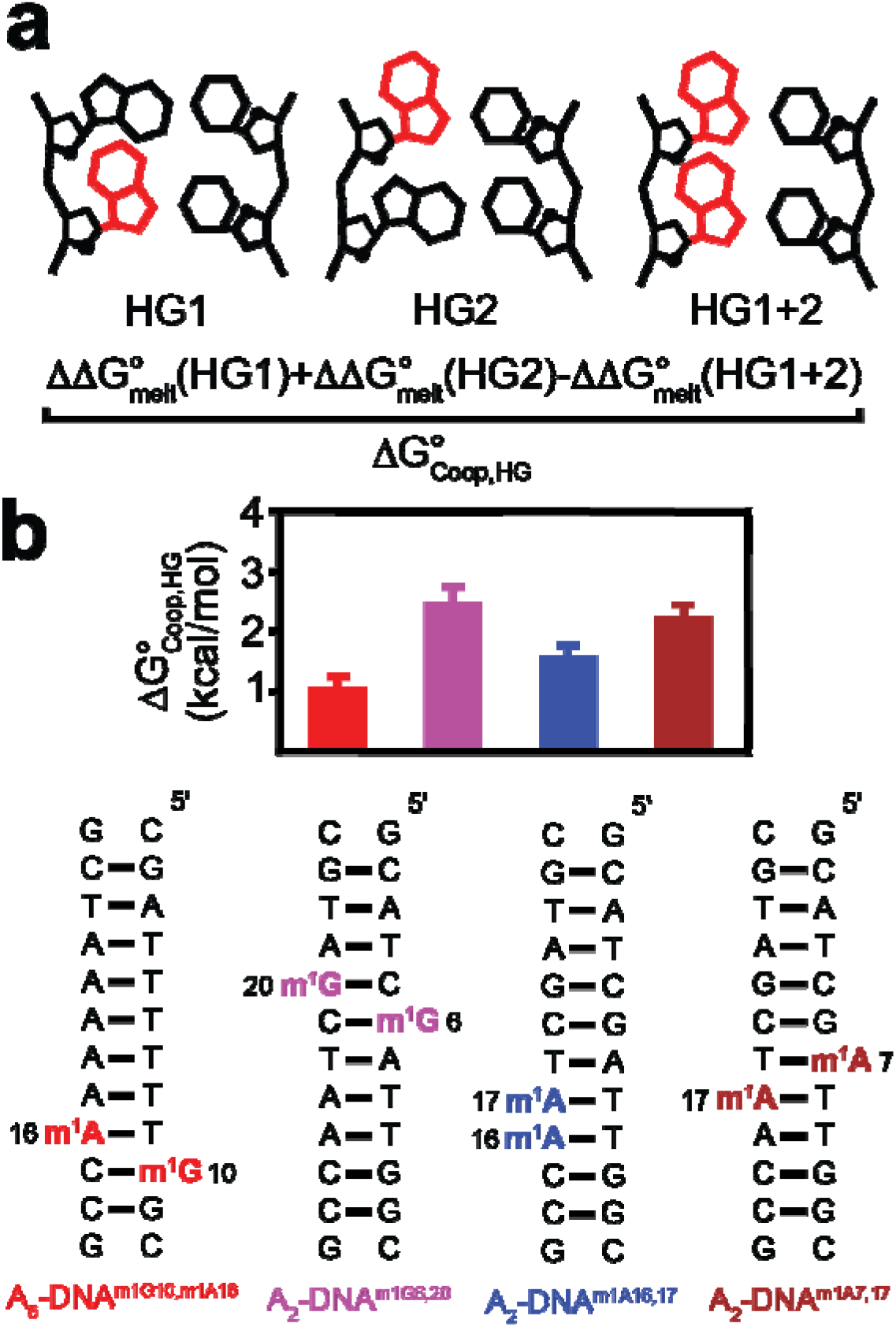
Measuring cooperativity of Hoogsteen base pair formation in DNA. (a) Estimating the cooperativity of Hoogsteen bp formation () using delta-melt (Methods). Red bases denote modified *N*^1^-methylated nucleotides (m^1^A^+^ and m^1^G) that bias the bp conformation towards Hoogsteen. (b) The value of over four different bp steps (Methods, Fig. S9). Bp steps are color-coded according to the duplexes shown below. Errors in were determined by propagating the uncertainties from the UV melts as described in Methods. Buffer conditions were 15 mM sodium phosphate, 150mM sodium chloride and 0.1mM EDTA and pH 5.4 or 6.8 for cooperativity involving G-C and only A-T bps, respectively. All values are reported at 25 °C.

We used delta-melt to assess the cooperativity of forming tandem T-A/G-C, C-G/G-C, T-A/T-A and A-T/T-A Hoogsteen bps corresponding to four different bp steps (5’-TG-3’/5’-CG-3’/5’-TT-3’/5’-AT-3’) embedded within two different duplexes under different buffer conditions (pH 5.4-6.8, 25 °C, 150 mM NaCl)(Fig. S9). We used the same DNA sequences employed in the experiments used to measure thermodynamic preferences for forming single Hoogsteen bps (Fig. 3c, 5c). Melting experiments were performed on tandem *N*^1-^ methylated DNA duplexes in which both bps are expected to be Hoogsteen, and their singly methylated counterparts, in which only one bp should be Hoogsteen (Fig. 8b and Fig. S9).

We first performed NMR experiments on the tandem methylated duplexes and their singly methylated counterparts to verify the presence of stably formed Hoogsteen bps. As expected, the singly *N*^1^-methylated duplexes showed NMR signatures of Hoogsteen bps (7), including upfield shifted T-H3 and downfield shifted m^1^A^+^-H6 protons for A-T Hoogsteen, and the downfield shifted C-H4 proton for m^1^G-C^+^ Hoogsteen bps, respectively (Fig. S9). The tandem *N*^1^-methylated duplexes also exhibited the expected downfield shifted purine-C8 resonances corresponding to the *N*^1^-methylated *syn* purine bases (7, 63), and downfield shifted m^1^A^+^-H6 resonances corresponding to the m^1^A^+^-T Hoogsteen bps. However, the T-H3 signal was not observable, and was likely broadened out of detection due to rapid solvent exchange kinetics.

We then measured the cooperativity of forming Hoogsteen bps using delta-melt experiments on the doubly and singly methylated duplexes (Fig. S1, Table S1). Indeed, across all four bp steps and different buffer conditions examined, tandem Hoogsteen bps formed cooperatively with sizeable 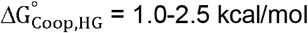 (Fig. 8b).

If tandem Hoogsteen bps do indeed form cooperatively in DNA as implied by delta-melt, and robustly in a manner independent of the methyl modification, the thermodynamic preferences to form a Hoogsteen bp at an unmodified bp should also increase when the adjacent site is a Hoogsteen bp stabilized by an *N*^1^-methylated purine modification. We tested this prediction using NMR RD experiments to directly measure the thermodynamic preferences to form G10-C15^+^ and A16-T9 Hoogsteen bps in A_6_-DNA when adjacent to pre-formed m^1^A16^+^-T9 or m^1^G10-C15^+^ Hoogsteen bps, respectively (Fig. S10a-d). Indeed, the thermodynamic preference to form Hoogsteen increased by ~0.2-1.0 kcal/mol when the neighboring bp was a modified pre-formed Hoogsteen. Similar increases were also observed recently when using T-T or C-T mismatches to mimic the Hoogsteen bp (19).

To further test these delta-melt predictions of tandem Hoogsteen cooperativity in an unmodified DNA duplex, we took advantage of the anti-tumor antibiotic echinomycin (79)(Fig. S10e-i). Echinomycin has previously been shown to bind duplex DNA containing two CG steps separated by a CA step, inducing tandem G-C^+^/A-T Hoogsteen bps at the CA step at low pH 5.3 (80). If the tandem Hoogsteen bps at the CA step form cooperatively, reducing the preference to form a Hoogsteen bp at one position should also reduce the preference to form the Hoogsteen bp at the second position, resulting in the loss of NMR signatures unique to tandem Hoogsteen bps. Indeed, reducing the thermodynamic preference to form G-C^+^ Hoogsteen either by increasing the pH to 6.9 or by substituting guanine with 7-deaza guanine at pH 5.3 (68, 81) resulted in the loss of NMR signals characteristic of tandem Hoogsteen bps (Fig S10e-i). Furthermore, the loss of tandem Hoogsteen bp signatures at pH 6.9 could be rescued by using m^1^G to stabilize the G-C^+^ Hoogsteen bp (Fig S10e-i). The above results suggest that Hoogsteen bps form cooperatively in echinomycin bound DNA, which may explain why multiple intercalating drugs such as echinomycin bind cooperatively to DNA (82).

Taken together, the above results strongly suggest that tandem Hoogsteen bps can form cooperatively in duplex DNA. Future experiments could independently verify tandem Hoogsteen cooperativity, for example, by examining the impact of single or double Hoogsteen mimicking substitutions on DNA-protein and DNA-drug binding affinities (4), or through the development of new approaches that can directly detect tandem Hoogsteen bps in unmodified DNA.

### Using delta-melt to measure sequence-specific signatures of Hoogsteen and base open states in duplex DNA

Many biochemical processes involving DNA including protein recognition (12), damage induction (83) and repair (84), as well as mutations including those related to cancer (85) are all strongly dependent on DNA sequence. When biochemical processes include steps that act on minor non-native conformational states of nucleic acids, such as the Hoogsteen and base open state, the sequence-specific thermodynamic preferences to form such conformations can shape the sequence-specificities of the biochemical processes as well. However, the ‘conformational signatures’ describing the sequence-specific preferences to form alternative conformational states have not been measured, primarily because of the difficulties in performing such a large number of NMR measurements.

Although the thermodynamic preferences to form minor non-native conformations such as Hoogsteen (8) and base open states (72) have been measured for various bp steps, these prior observations were confounded by additional variation in the global sequence context and position along the helix. We therefore used delta-melt to overcome these barriers and measured conformational signatures for A-T and G-C^+^ Hoogsteen bps, and the A-T base open state for all 16 trinucleotide sequence contexts, at a given positional context. These measurements were performed on a highly stable scaffold sequence (scaf2, Fig. 9a) enabling robust melting measurements across all 16 sequence contexts, including when destabilizing modifications (m^1^A^+^, m^1^G, and m^3^T) are present.

**Figure 9.**
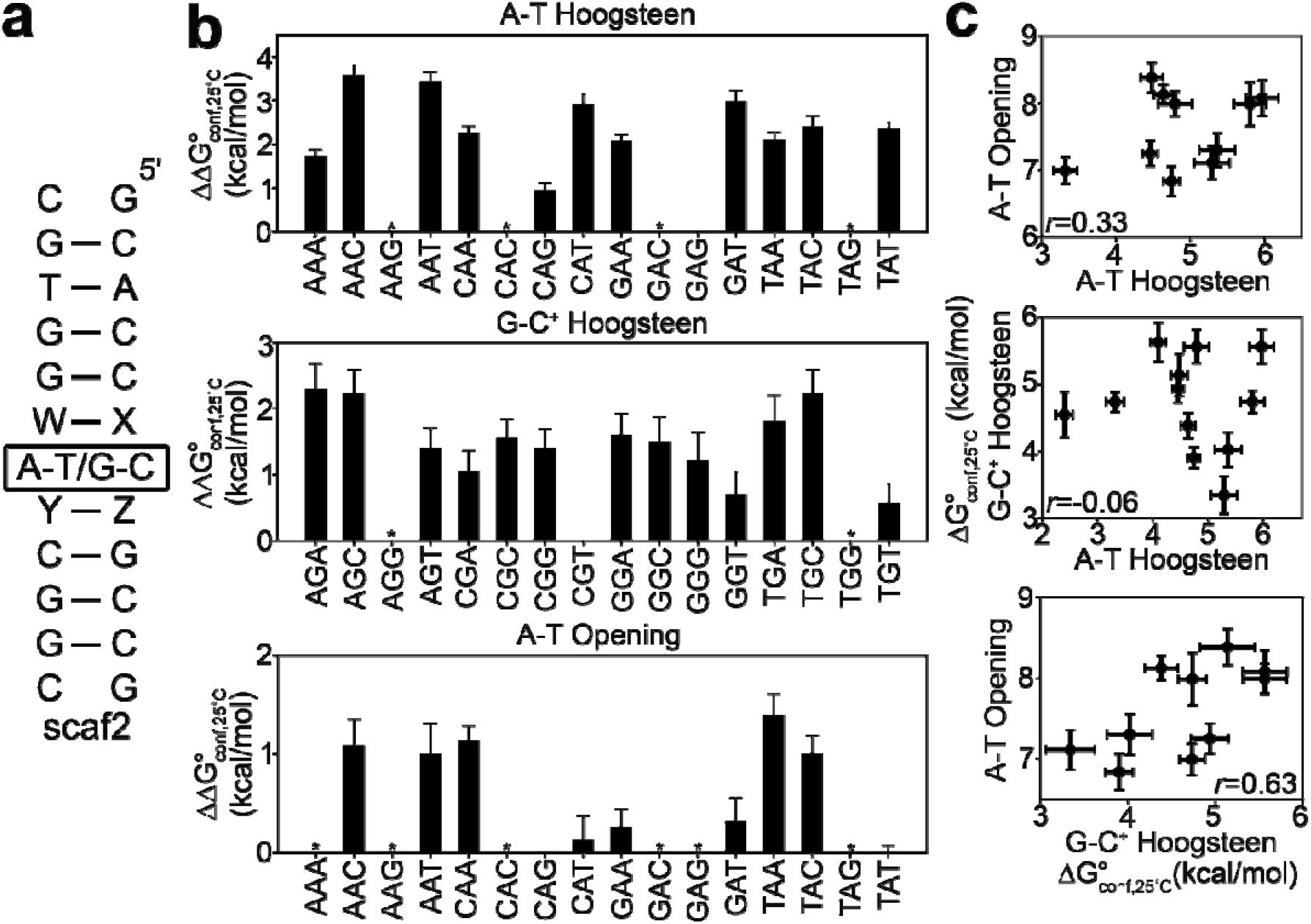
Measuring sequence-dependent thermodynamic preferences in DNA using delta-melt. (a) DNA duplex used to measure sequence-dependent thermodynamic preferences using delta-melt. W-X and Y-Z represent either of the four Watson-Crick bps (A-T/T-A/C-G/G-C). Thermodynamic preferences to form base open or Hoogsteen conformations were measured using delta-melt for the central bp indicated using a box. (b) Relative thermodynamic preferences () to form A-T and G-C^+^ Hoogsteen bps and the A-T base open states measured using delta-melt at 25 °C (Fig. S1-S2, Tables S1-S2). Sequence contexts are given in the 5’ to 3’ direction. Thermodynamic preferences were referenced to the sequence context with the most favorable preference (5’-GAG-3’, 5’-CGT-3’, and 5’-CAG-3’ for A-T Hoogsteen, G-C^+^ Hoogsteen and A-T base open states, respectively). Sequences for which reliable thermodynamic preferences could not be obtained due to non-two-state melting behavior are denoted using * (Methods). Buffer conditions for the measurements are described in Methods. (c) Correlation plots between the thermodynamic preferences for formation of A-T and G-C^+^ Hoogsteen bps, and open A-T bps at 25 °C. The Pearson correlation coefficient (*r*) is shown in the inset. Note that for correlating preferences for A-T and G-C^+^ Hoogsteen bps, the purine bases were aligned with each other, for example, the sequence context 5’-AAA-3’ for A-T Hoogsteen was correlated with 5’-AGA-3’ for G-C^+^ Hoogsteen). Error bars in panel b and c were obtained by propagating the error from the UV melts and, as described in Methods.

In measuring the conformational signatures using delta-melt, our analysis assumed that the offsets c(i) are independent of sequence context. Although this is supported by our benchmarks (Fig. 3–7), without additional delta-melt and NMR calibration experiments, we cannot rule out different c(i) values are applicable for different sequence contexts. In addition, for some sequence contexts, the melting curves deviated from a simple two-state behavior, and the data could not be used to extract reliable thermodynamic preferences (* symbols in Fig. 9b, Fig. S3, Methods). Additional studies are needed to understand the origins of the non-two-state behavior of the melting curves and how to extract robust thermodynamic parameters from them.

delta-melt revealed large sequence-specific variations in the thermodynamic preferences to form A-T (~1.7-3.5 kcal/mol) and G-C^+^ (0.5-2.2 kcal/mol) Hoogsteen bps, as well as the A-T base open state (0.3-1.4 kcal/mol), which are on the order of 2-3 kcal/mol, over the sequence contexts which could be measured reliably using delta-melt (Fig. 9b, Fig. S1-S2 and S11-S12, and Tables S1-S2). This corresponds to ~10-100-fold sequence-specific variations in the population of the minor conformation in the dynamic ensemble purely due to changes in the sequence of the immediate neighbor.

To verify the conformational preferences obtained using delta-melt, we used NMR RD to independently measure the energetic cost to form G-C^+^ Hoogsteen bps in the 5’-TGC-3’ and 5’-CGT-3’ sequence contexts (Fig. 10a), predicted by delta-melt to have low and high propensities (Fig. 9b) to form G-C^+^ Hoogsteen bps, respectively. Measurements were performed under two different buffer conditions and three different temperatures to test the robustness of the predictions. One of the contexts, 5’-TGC-3’, (data point 6, Fig. 10b) with low propensity is predicted by delta-melt to have little to no observable NMR RD even under low pH conditions (pH 5.4, 150 mM NaCl, 15 °C). Such behavior has never been observed previously, and therefore represents a strong prediction from the delta-melt approach. On the other side, the 5’-CGT-3’ context (data point 1, Fig. 10b) is predicted to have a relatively large population (~0.7 %) of the minor G-C^+^ Hoogsteen bp at low pH (pH 5.4, 25mM NaCl, 30 °C). Prospective comparison of six thermodynamic preferences measured using delta-melt (Fig. 9b) and NMR RD (Fig. 10b) reveals an average accuracy of ~0.5 kcal/mol, which is small relative to the range of 0.5-2.0 kcal/mol measured across different sequence contexts. delta-melt correctly predicted the lack of observable RD for the 5’-TGC-3’ sequence (data point 6, Fig. 10b), and enabled measurements of thermodynamic preferences at this position when it was not feasible using NMR RD. delta-melt also correctly predicted the 5’-CGT-3’ context having the highest propensity to form G-C^+^ Hoogsteen (data point 1, Fig. 10b). Nevertheless, we also observed a relatively large deviation of ~0.8 kcal/mol between the delta-melt and NMR-derived thermodynamic preferences for G6-C19^+^ Hoogsteen bp formation in scaf2_TGC^GC^ at pH 5.4 high salt (150mM NaCl) and 30 °C (data point 4, Fig. 10b). This sequence context may have a unique c(i) offset (Fig. 5c), as only 1 out of the 7 measurements used for establishing calibration curves contains the same 5’-TGC-3’ tri-nucleotide step (Fig. 5b-c). These results support the utility of delta-melt to determine sequence-specific conformational signatures and to identify potential conformational hotspots.

**Figure 10.**
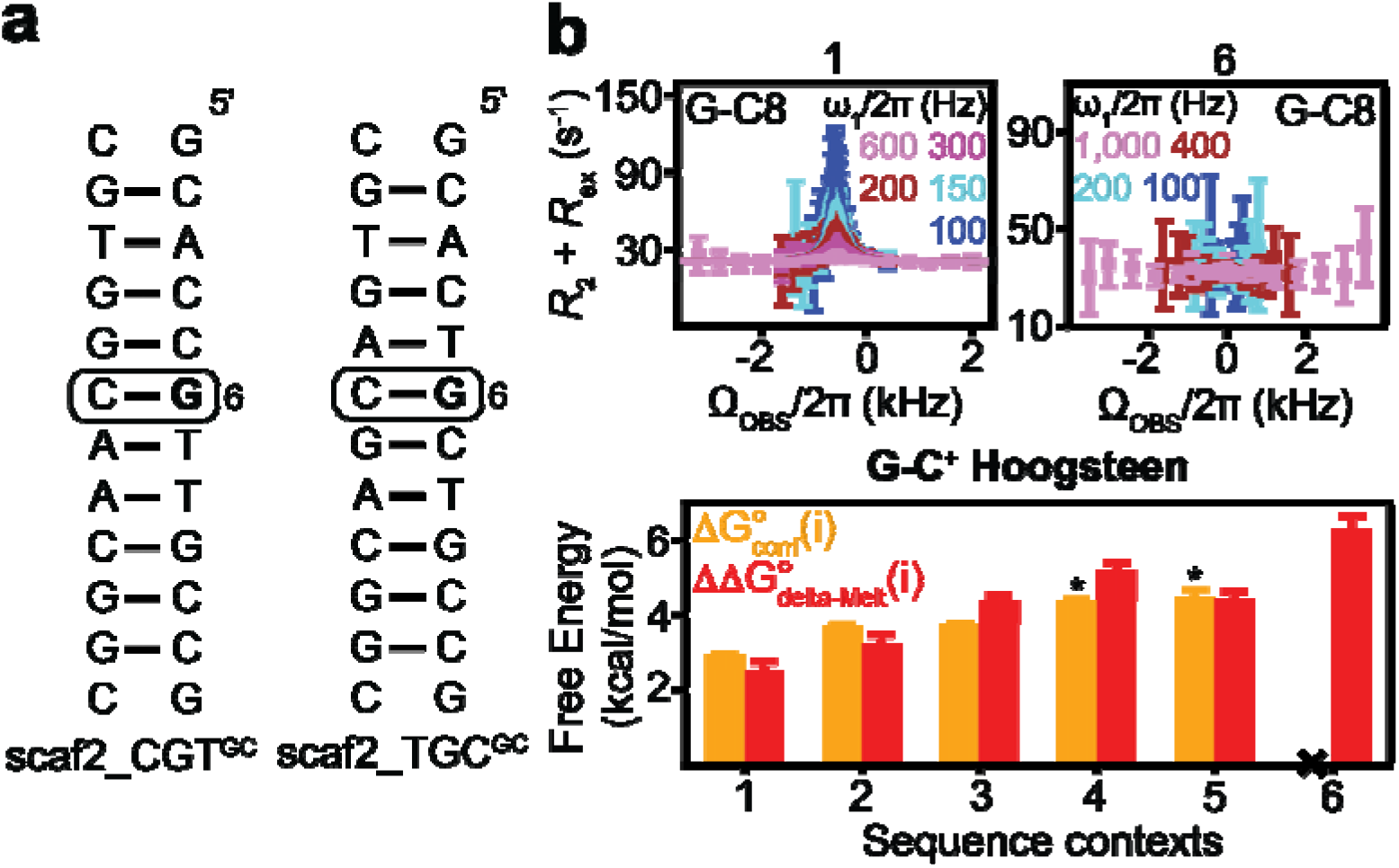
Testing the accuracy of delta-melt derived thermodynamic preferences to form G-C^+^ Hoogsteen bps using NMR. (a) Duplexes used to test delta-melt derived thermodynamic preferences. ^13^C/^15^N isotopically labeled residues are in bold. All remaining residues are unlabeled. (b) (Bottom) Comparison of measured thermodynamic preferences to form G-C^+^ Hoogsteen bps using NMR (orange, Table S5) and values measured using delta-melt (red, Table S1). Numbers 1-6 correspond to combinations of sequences and buffer/temperature conditions for the measurements and are given in Table S9. Lower-confidence data points (Methods) obtained from weak *R*_1ρ_ RD profiles are denoted using “*”. Data point 6 with a flat *R*_1ρ_ RD profile is denoted using “x”, indicating that the minor G-C^+^ Hoogsteen bp in this case falls outside the NMR RD detection limit. Error bars for free energies were obtained by propagating the uncertainty from UV melts (Methods). (Top) Representative examples of *R*_1ρ_ RD profiles showing the measured effective transverse relaxation rate (*R*_2_+*R*_ex_) as a function of offset (Ω_OBS_/2π) and color-coded spin-lock power (ω_1_/2π)(Methods). Errors bars were obtained by propagating the uncertainty in *R*_1ρ_ as described in Methods.

Given the robustness of the delta-melt derived thermodynamic preferences (Fig. 10b), we then proceeded to examine their sequence-dependencies. The largest sequence-dependence was observed for A-T Hoogsteen 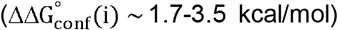 followed by G-C^+^ Hoogsteen 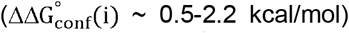 and with the open A-T state showing the weakest sequence-dependence 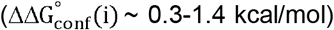 possibly because of its relatively increased flexibility. Interestingly, the delta-melt derived conformational signatures were very different (*r*~0-0.63) for these three conformational states (Fig. 9c). Therefore, those trinucleotide sequence contexts showing a preference for forming one state may not also favor the formation of another state. For example, the 5’-GAT-3’/5’-GGT-3’ trinucleotide context is the third most favorable context for forming the A-T base open state and G-C^+^ Hoogsteen bps but is the third least favorable context for A-T Hoogsteen bps. If different processes act on different conformational states of the DNA, these conformational signatures could translate into different sequence-specificities for the distinct DNA biochemical reactions.

The delta-melt data also reveals that the 5’ and 3’ neighboring bps of the purine base of a given bp influence thermodynamic preferences to form the three non-native conformational states to different extents. The 3’ neighboring bp dominates the preferences for A-T and G-C^+^ Hoogsteen bps, whereas for the A-T base open state both neighbors contribute equally (Fig. S13). In addition, substantial variations (~1.5-2.0 kcal/mol) in the sequence-specific thermodynamic preferences due to the third bp are seen for a given dinucleotide step, indicating that conformational signatures must at minimum be described by a triplet code (Fig. S13).

Another advantage of delta-melt is that it provides unique insights into the origins of these different sequence-specific conformational signatures. For example, the sequence-specific variations to form A-T Hoogsteen bps appear to be dominated by the sequence-specific variations in the melting stability of the Watson-Crick A-T bp (Fig. S13). Therefore, those sequences that are least stable in the unmodified DNA have the highest propensities to form A-T Hoogsteen bps. This is in line with a prior study showing that A-T Hoogsteen bps have high propensities to form in CA and TA steps, which are the least stable in the Watson-Crick form (8).

Interestingly, in contrast, the sequence-specific variations to form G-C^+^ Hoogsteen bps appears to be dominated by the sequence-specific variations in the melting stability of the m^1^G-C^+^ Hoogsteen bp (Fig. S13, Discussion S3). These variations could arise due to large sequence-specific variations in the pKa of the cytosine imino proton, possibly mediated by sequence-specific stacking. Finally, in the case of the base open state, the sequence-specific preferences are determined by both the stability of the A-T Watson-Crick and base open (or specifically A-m^3^T) states (Fig. S13).

## Discussion

Melting experiments have been used with great success to understand factors contributing to the stability of the major folded conformations of nucleic acids relative to the unfolded state, providing the basis for the nearest neighbor rules, which are widely used to predict nucleic acid secondary structure and other properties of interest (44, 47). These prior studies took advantage of the relatively high throughput and low sample requirements of melting experiments, which permit broad explorations of how folding thermodynamics varies across different sequence and structural motifs and experimental conditions. In this study, we showed that this well-established approach (43, 44, 47–49) can be extended to the thermodynamic stabilities of minor non-native conformational states in the ensemble, whose contribution to melting experiments are typically negligible (53).

The key to delta-melt is to use chemical modifications to render minor non-native conformational states in the ensemble the major state. This then makes it possible to access information regarding thermodynamic stabilities of minor conformations. A fundamental assumption in delta-melt is that the relative energetic cost for modifying the minor conformation and single strands does not vary with sequence contexts and conditions. Under this assumption, calibration with independent measurements of conformational preferences over different sequence contexts and conditions as done here using NMR, provides a basis for exploring the energetics of other conformations in the ensemble using melting experiments. While our study supports this assumption for the conformational states studied here, it needs to be further verified, especially for different sequence contexts through additional measurements of sequence-specific thermodynamic preferences using NMR. This might reveal sub-groups of sequence contexts that should be calibrated independently, thus improving the accuracy of the approach.

Prior studies established the utility of modifications as structural mimics for a variety of minor non-native conformations in nucleic acids (reviewed in (40)). Our results further establish chemical modifications as energetic mimics of minor conformational states. Thus, one can measure thermodynamic preferences to form non-native states in a facile manner using melting experiments, in contrast to NMR RD based approaches, which are comparatively more costly and sample intensive. In addition, our study introduced m^3^T as a mimic of the A-T base open state. Solving high-resolution structures and dynamic ensembles (62) for m^3^T modified DNA duplexes could shed light on the structure of this elusive state, in a manner analogous to how m^1^A^+^-T provided structural insights into transient Hoogsteen bps (60).

delta-melt made a number of unique applications possible. It allowed us to gain rare insights into the cooperativity of forming non-native bp conformations in nucleic acids, revealing that tandem Hoogsteen bps form cooperatively in duplex DNA. This implies that *N*^1^-methylation and possibly other forms of DNA damage can increase the Hoogsteen preferences at neighboring sites, and this could in turn be exploited by damage repair enzymes such as glycosylases during recognition and repair. delta-melt could be used in the future to more broadly examine how Hoogsteen preferences are altered with other perturbations such as mismatches, nicks, and other lesions (19). delta-melt can also potentially be used to measure cooperativity involving different types of bp conformational changes such Hoogsteen and base open states as well as to assess whether cooperativity can arise between bps that are not immediately adjacent to one another (86).

For the first time, delta-melt enabled us to measure the preferences to form non-native conformations in DNA as a function of the trinucleotide sequence context, revealing ~10-100-fold differences in the probabilities of forming A-T and G-C^+^ Hoogsteen and A-T base open states due to changes in sequence context alone. We observe rich behavior in the sequence-dependent preferences to form these three conformational states. Not only did the scale of the sequence-dependencies vary across the three states, being weakest for the base open conformation, we also observed differences in the extent to which these sequence preferences are driven by the stability of the major versus minor state. These results strongly suggest that hidden within the DNA double helix is a rich landscape of sequence-specific minor conformations. Future studies should verify these predictions from delta-melt and examine their biological implications.

In the future, delta-melt could be used to study how changes in DNA sequence beyond the trinucleotide context affect conformational preferences and examine how the conformational preferences are shaped by epigenetic modifications such as 5-methylcytosine. The sequence-dependent thermodynamic propensities obtained from delta-melt can also be cross-referenced with sequence-dependent biological processes, including over 60 mutational signatures linked to cancer (87), to potentially identify minor conformations that drive them (Fig. S16, Discussion S4). For example, we observe a strong correlation between the sequence-dependent probabilities for T>A mutations in mutational signature 17A and the thermodynamic propensities for A-T Hoogsteen bp formation obtained using delta-melt, suggesting potential biological roles for Hoogsteen bps that require future investigation (Discussion S4).

It should be possible to apply delta-melt to other minor conformational states not examined here. For example O^6^-methylguanine, a naturally occurring mutagen (88) could be used to measure thermodynamic preferences to form Watson-Crick like G•T/U mismatches in DNA, RNA and RNA-DNA hybrids, which have been proposed to evade fidelity checkpoints, contributing to misincorporation errors during DNA replication, translation, and transcription (20). Furthermore, *N*^3^-methylcytosine could be used to measure thermodynamic preferences for opening G-C bps, and 8-oxo guanine and adenine, which are commonly occurring form of oxidative damage (89), could be used to measure the thermodynamic preferences to form alternative conformations of A-G mismatches such as A(*anti*)^+^-G(*syn*) and A(*syn*)-G(*anti*). delta-melt could also be used to study the impact of experimental conditions on forming non-native conformations involving secondary structural rearrangements in RNA (90). It may be possible to extend delta-melt to proteins, for which mutants stabilizing minor conformations have been reported (91, 92). The throughput of delta-melt could also be increased in the future using high-throughput melting experiments (93). Through these applications, it may be feasible to extend the nearest neighbor model to predict the energetics of various alternative nucleic acid conformational states, an important step toward predicting the conformational ensembles of nucleic acids.

## Materials and Methods

### Sample preparation

#### Buffer preparation

With the exception of ^1^H proton exchange measurements (see below), the buffer used in NMR experiments consisted of 15 mM sodium phosphate, 25, 125, or 150 mM sodium chloride and 0.1 mM ethylenediaminetetraacetic acid (EDTA) in 90% H_2_O:10%D_2_O at pH between 4.4 and 6.8 (summarized in Table S3). The buffer used in imino ^1^H exchange experiments on G-DNA and A_6_-DNA consisted of 10 mM sodium phosphate, 100 mM sodium chloride, 1 mM EDTA and 1 mM triethanolamine (TEOA) with or without ammonia in 95% H_2_O:5% D_2_O at pH 8.8. Buffers with effective ammonia concentrations of 20, 40, 100 and 150 mM were prepared by titrating ammonium hydroxide solution (14.8 M, Millipore Sigma) to the NMR buffer, followed by adjusting pH to 8.8 by adding hydrochloric acid (HCl)(94). The effective concentration of ammonia [NH_3_] was computed using the buffer pH (8.8) and total ammonia concentration added to the buffer ([NH_3_]_0_) as follows:

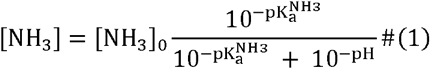

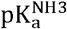 is the pKa for ammonia. The buffer used in imino ^1^H exchange experiments on TBP-DNA in the prior study by Chen et al. (72) consisted of 100 mM sodium chloride, 2 mM EDTA and 2 mM TEOA in 90% H_2_O:10% D_2_O at pH 8.0.

UV melting experiments for A-T and G-C^+^ Hoogsteen bps and methyl rotation in m^6^A, were performed in a buffer solution with the same composition as that used for NMR measurements, but in 100% H_2_O. UV melting measurements on A_2_-DNA show that the effect of adding 10% D_2_O on the melting free energies falls within the error of UV measurements (Fig. S1 and Table S1). Hence, the lack of D_2_O in the buffer used for UV melting is not expected to affect the delta-melt correlations (Fig. 3c, 4c, 4e, 5c and 7c). The UV melting experiments for A-T base opening were performed in buffers containing the same H_2_O:D_2_O composition used in the imino ^1^H exchange measurements, as defined above.

#### Annealing and buffer exchange

Duplex DNA and RNA samples were prepared by mixing the two complementary strands (~1 mM) in a 1:1 ratio, heating to 95 °C for 5-10 min, followed by slow annealing at room temperature. Hairpin DNA and RNA samples were prepared by diluting the samples to concentrations < 100 μM, heating to 95 °C for 5-10 min, followed by rapid annealing on ice. DNA and RNA samples used for NMR measurements were buffer exchanged into the desired NMR buffer with final concentration ~1 mM using Amicon Ultra-0.5/15 centrifugal concentrators (3-kDa cutoff, Millipore Sigma). The samples used for optical melting experiments were prepared by diluting NMR samples to ~3 μM using buffer. Extinction coefficients for all single-strands were estimated using the ADT Bio Oligo Calculator (https://www.atdbio.com/tools/oligo-calculator). Extinction coefficients for the modified single-strands were assumed to be equal to their unmodified counterparts (modified bases are estimated to affect the extinction coefficient for the oligos used here by <10% based on reference values in Basanta-Sanchez *et al*. (95)). The DNA-echinomycin complexes were prepared by mixing DNA duplexes in NMR buffer with 3x echinomycin (Sigma-Aldrich) dissolved in methanol, maintaining the NMR buffer:methanol ratio at 2:1 (volume:volume). The mixture was shaken and incubated at room temperature for 30 min, followed by slow solvent evaporation under an air stream overnight. The dried samples were redissolved in the appropriate amount of water, ensuring that the final concentration of each buffer component was identical to the NMR buffer. NMR samples in D_2_O were prepared by rapidly freezing and lyophilizing samples in water overnight and resuspending them into 100% D_2_O (Millipore Sigma).

#### Unlabeled oligonucleotides

In what follows we use “lb” inside a bracket following the construct name to refer to specific duplexes (see Fig. S4) in which one or both strands are isotopically labeled, and isotopically labeled hairpins. For example, A_6_-DNA^m1A16^(s2lb), is a version of the A_6_-DNA^m1A16^ duplex with a specific labelling scheme defined in Fig. S4. Constructs without “lb” in the name are unlabeled. Secondary structures and labeling schemes of all constructs used for NMR measurements are given in Fig. S4.

All the unmodified DNA strands were purchased from Integrated DNA Technologies (IDT) with standard desalting purification. The unlabeled m^1^A^+^ modified single-strands for the A_6_-DNA^m1A16^ and A_6_-DNA^m1A16^(s2lb) duplexes were purchased from Midland DNA Technologies with reverse-phase (RP) HPLC purification, while the m^1^rG modified single-strand comprising the A_6_-DNA^m1rG10^ duplex was purchased from GE Healthcare Dharmacon with HPLC purification. Modified single-strands for A_6_-DNA^m1rA16^, A_2_-DNA^m1G6^, A_2_-DNA^m1G20^, A_2_-DNA^m1G6,20^, hpAcDNA^m1G7^, A_2_-DNA^m1A7^, A_2_-DNA^m1A17^, A_2_-DNA^m1A7,17^, A_2_-DNA^m1A16,17^, A_6_-DNA^m3T5^, A_6_-DNA^m3T7^, A_6_-DNA^m3T9^, A_6_-DNA^m3T22^, TBP-DNA^m3T3^, TBP-DNA^m3T5^, TBP-DNA^m3T16^, TBP-DNA^m3T17^, TBP-DNA^m3T18^, TBP-DNA^m3T19^, TBP-DNA^m3T21^, E12DNA-HG^m1G13^ and E12DNA-HG^7deazaG13^ were purchased from Yale Keck Oligonucleotide Synthesis Facility with cartridge purification. The m^1^A^+^ modified single-strand in the A_6_-DNA^m1A21^ duplex was purchased from Yale Keck Oligonucleotide Synthesis Facility with HPLC purification. The m^1^A^+^ modified single-strand in the A_2_-DNA^m1A16^ duplex was purchased from Midland DNA Technologies with RP-HPLC purification and Yale Keck Oligonucleotide Synthesis Facility with cartridge purification for UV melting measurements in pH 6.8 at 25 mM NaCl and 150 mM NaCl, respectively. The A_6_-DNA^m1G10,m1A16^ duplex was comprised of an m^1^A^+^ modified single-strand purchased from Midland DNA Technologies with RP-HPLC purification, and an m^1^G modified single-strand purchased from Yale Keck Oligonucleotide Synthesis Facility with cartridge purification. To minimize Dimroth rearrangement (96) of m^1^A^+^ during oligonucleotide synthesis and purification, all the m^1^A^+^ modified DNA single-strands were synthesized and deprotected using UltraMild chemistry (https://www.glenresearch.com/reports/gr19-12). The m^1^G10 single-strand comprising the A_6_-DNA^m1G10^, A_6_-DNA^m1G10,C16^ and A_6_-DNA^m1G10,T16^ duplexes used for UV melting measurements was obtained from Yale Keck Oligonucleotide Synthesis Facility with cartridge purification, while that comprising the A_6_-DNA^m1G10^ (s1lb) duplex used for NMR measurements was purchased from Midland DNA Technologies with gel filtration purification.

The m^1^G and m^3^T modified strands in the scaf2 series of DNA duplexes and in the A_2_-DNA^m1G10^ and A_6_-DNA^m3T6^ duplexes, the rNTP modified single-strands in the A_6_-DNA^rA16^, A_6_-DNA^rC16^, A_6_-DNA^rT16^ duplexes, and the unlabeled RNA single-strands in A_6_-RNA^m6A16^(m^6^A16^C2,C8^lb), A_6_-RNA^m6A16^, A_6_-RNA^m62A16^, dsGGACU^m6A6^ and dsGGACU^m62A6^ were synthesized in-house using a MerMade 6 Oligo Synthesizer. Standard RNA phosphoramidites (n-ac-rA, n-ac-rG, n-ac-rC, rU, rT, rm^6^A and rm^6^_2_A, Chemgenes) and DNA phosphoramidites (n-ibu-G, n-bz-A, n-ac-C, T, m^3^T and n,n-dmf-m^1^G, Chemgenes), and columns (1000 Å from Bioautomation) were used with a coupling time of 6-12 min (RNA) and 1 min (DNA), with the final 5’-dimethoxy trityl (DMT) group retained during synthesis. The oligonucleotides were cleaved from the supports (1 μmol) using ~1 ml of AMA (1:1 ratio of ammonium hydroxide and methylamine) for 30 min and deprotected at room temperature for 2 hours. The m^1^G and m^3^T containing DNA samples were then purified using Glen-Pak DNA cartridges and ethanol precipitated, while all the other samples were dried under airflow to obtain oligonucleotide crystals. They were then dissolved in 115 μl DMSO, 60 μl TEA and 75 μl TEA.3HF and heated at 65°C for 2.5 h for 2’-O deprotection. The samples were then neutralized using 1.75 ml of Glen-Pak RNA quenching buffer, loaded onto Glen-Pak RNA cartridges for purification and were subsequently ethanol precipitated. The rG modified single-strand comprising the A_6_-DNA^rG10^ duplex used for UV melting measurements was synthesized above as described for the rA modified single-strand in A_6_-DNA^rA16^, while that comprising the A_6_-DNA^rG10^(s1lb) duplex used for NMR measurements was purchased from IDT.

The m^1^A^+^ modified strands in the scaf2 series of DNA duplexes were also synthesized in house using a Mermade-6 Oligo Synthesizer. UltraMild DNA phosphoramidites (T, n-pac-C, n-tbpac-G, n-pac-A, n-fmoc-m^1^A) and Ultramild Cap A (Glen Research), and columns (1000 Å from Bioautomation) were used with a coupling time of 1 min, with the final 5’-dimethoxy trityl (DMT) group retained during synthesis. The powdered supports were then incubated with ~2 mL of concentrated NH_4_OH for 1 hour and room temperature for cleavage of oligonucleotides from the support. The supports were spun down and the supernatant was collected in separate tubes. The remaining support was washed with ~0.5 mL of concentrated NH_4_OH, which was pooled together with the supernatant post spinning down, to result in a net volume of ~2 mL of the cleaved oligo in NH_4_OH. The oligonucleotides were then heated for 18 hours at 37 °C for base deprotection. The oligo solutions were then dried under air flow to bring the volume to ~1mL following which they were purified with Glen Pak DNA cartridges and ethanol precipitated.

#### ^13^C/^15^N-labeled oligonucleotides

Uniformly ^13^C/^15^N isotope labeled DNA strands comprising duplexes A_6_-DNA(ulb), A_6_-DNA(s1lb), A_6_-DNA(s2lb), A_6_-DNA^m1A16^(s2lb), A_6_-DNA^m1G10^(s1lb), A_6_-DNA^rG10^(s1lb) and AcDNA(ulb), and residue-type (G and T or C and A labeled on either strand) DNA strands comprising duplexes A_2_-DNA(ulb) and A_2_-DNA(s2lb) were synthesized using an *in vitro* primer extension approach (97) with a template DNA (IDT), Klenow fragment DNA polymerase (New England Biolabs), uniformly ^13^C, ^15^N-labeled deoxynucleotide triphosphates (dNTPs, Silantes) and/or unlabeled dNTPs (Thermo Fisher Scientific). The reaction mixture was centrifuged to remove excess pyrophosphate, and then subsequently concentrated to 1.5□mL using a 3□kDa molecular weight cutoff centrifugal concentrator (Millipore Sigma). 1.5□mL of a formamide based loading dye was then added to the reaction mixture, which was then heated at 95□°C for 5□min for denaturation. The mixture was then loaded onto a denaturing gel (20% 29:1 polyacrylamide/8M urea in TBE buffer) for resolution of the target oligonucleotide from other nucleic acid species. Gel bands corresponding to the pure target single-strands were identified by UV-shadowing and subject to electroelution (Whatman, GE Healthcare, in TAE buffer) followed by ethanol precipitation. Careful optimization of the NTP concentrations, in particular ATP, and that of Mg^2+^ during primer extension was seen to be necessary to avoid spurious NTP addition to form dangling ends in the resulting oligonucleotides.

The site-specifically labeled DNA strands comprising duplexes (A_6_-DNA(A16lb), scaf2_CGT^GC^(G6lb) and scaf2_TGC^GC^(G6lb)) were purchased from Yale Keck Oligonucleotide Synthesis Facility with cartridge purification.

The atom-specifically labeled DNA and RNA single-strands comprising constructs A_6_-DNA^rA16^(rA16^C8^,C15lb), A_6_-RNA^m6A16^(m^6^A16^C2,C8^lb), hpGGACUm^6^A6(m^6^A6^C2,C8^lb) and hpGGACUm^6^A6(m^6^A6^C10^lb) were synthesized in-house with a MerMade 6 Oligo Synthesizer as described above for the rA16 modified single-strand in A_6_-DNA^rA16^, using standard RNA (n-ac-rA, n-ac-rG, n-ac-rC, rU, Chemgenes) and DNA (n-ibu-G, n-bz-A, n-ac-C, T, Chemgenes) phosphoramidites, ^13^C C8 labeled rA (98), ^13^C/^15^N labeled C (Cambridge Isotope Laboratories), ^13^C C2 and C8 labeled rm^6^A (67), ^13^C C2 and C8 labeled rA (99), ^15^N N3 labeled rU (100) and ^13^C C10 labeled rm^6^A phosphoramidites (74).

### NMR experiments

The imino ^1^H exchange experiments were carried out on a Bruker Avance III 700 MHz spectrometer equipped with a HCN room temperature probe while the remaining NMR data was collected on Bruker Avance III 600 MHz or 700 MHz NMR spectrometers equipped with HCPN and HCN cryogenic probes, respectively.

#### Resonance assignment

Resonance assignments for A_6_-DNA, A_6_-DNA^m1A16^, A_6_-DNA^m1G10^, A_6_-DNA^rA16^, A_6_-DNA^rG10^, A_2_-DNA, AcDNA, hpGGACU^m6A6^, G-DNA, TAR, E12DNA-HG and E12DNA-WC were obtained from prior studies (7, 63, 80, 90, 101, 102)(Fig. S4). Assignments for A_6_-RNA^m6A16^ will be published in a subsequent study (74). These studies also show that the constructs fold into their expected secondary structures as given in Fig. S4. Note that the C1’-H1’ assignments for T5 and T6 were incorrectly swapped in assignments of A_6_-DNA, and consequently in those of A_6_-DNA^m1A16^, A_6_-DNA^m1G10^, A_6_-DNA^rA16^, A_6_-DNA^m1rA16^, A_6_-DNA^rG10^, A_6_-DNA^m1rG10^ in prior studies (7, 60, 62, 63, 68, 81). Nevertheless, the conclusions of these prior studies are not affected by the assignment swap. The assignment swap has been corrected in the assignments in this study and in the BMRB entries 30254 and 30255. Resonance assignments for A_6_-DNA^m3T9^ were obtained using 2D [^1^H, ^1^H] Nuclear Overhauser Effect Spectroscopy (NOESY), 2D [^1^H, ^1^H] Total Correlation Spectroscopy (TOCSY) and 2D [^13^C, ^1^H] Heteronuclear Single Quantum Correlation (HSQC) experiments. The assignments for scaf2_CGT^GC^ and scaf2_TGC^GC^ were easily obtained since they were site-specifically labeled. All the NMR spectra were processed using NMRpipe (103) and analyzed using SPARKY (T. D. Goddard and D. G. Kneller, SPARKY 3, University of California, San Francisco).

#### ^13^C/^15^N *R*_1ρ_ relaxation dispersion

Off-resonance ^13^C/^15^N *R*_1ρ_ relaxation dispersion (RD) experiments were implemented using a 1D selective excitation scheme as described in prior studies (39, 81, 104). The spin-lock power (ω_1_/2π) used ranged from 100 to 1000 Hz, while the offsets used ranged from ±10 and ±3.5 times the spin-lock power for C and N experiments, respectively (Table S4). For each resonance, six to ten delay times were selected during the relaxation period with maximum duration up to 60 ms and 150 ms for ^13^C and ^15^N, respectively. The experimental conditions for all the ^13^C/^15^N *R*_1ρ_ RD experiments (temperature, magnetic field, solvent) are summarized in Table S3.

#### Analysis of *R*_1ρ_ data

The *R*_1ρ_ data was analyzed as described previously (21). Briefly, 1D peak intensities as a function of delay times extracted using NMRPipe (103) were fitted to a mono-exponential decay to obtain the *R*_1ρ_ value for the different spin-lock power and offset combinations. The error in *R*_1ρ_ was estimated using a Monte Carlo procedure as described previously (40). Exchange parameters of interest, such as the population of the i^th^ minor conformation (*p*_i_), the exchange rate between the major and i^th^ minor conformation (*k*_ex_ = *k*_1_ + *k*_−1_, in which *k*_1_ and *k*_−1_ are the forward and backward rate constants, respectively), the chemical shift difference between the i^th^ minor and the major conformations (Δω_i-Major_ = ω_i_ – ω_Major_, in which ω_i_ and ω_Major_ are the chemical shifts of the nucleus in the i^th^ minor and the major conformations, respectively) were then extracted by fitting the *R*_1ρ_ data for a given nucleus to a two-state exchange model using the Bloch-McConnell equations. The validity of a two-state model to analyze the *R*_1ρ_ data for Hoogsteen bp formation and *N*^6^-methyl amino rotation has been extensively validated based on prior studies (7, 8, 62, 63, 68, 74, 81). Owing to the presence of an equilibration delay (5 ms) in the pulse sequences (81, 104), initial magnetization at the start of the Bloch-McConnell simulations was assumed to be equilibrated between the major and minor conformations. Combined fits of the *R*_1ρ_ data for multiple nuclei were performed by sharing *p*_i_ and *k*_ex_. Owing to the weak nature of the RD profiles (see below) for G6-C1’ and C8 in scaf2_TGC^GC^ at pH 5.4 150mM NaCl (high salt, >25mM, HS) and 40 °C, the values of Δω_i-Major_ obtained when fitting both data sets with shared *p*_i_ and *k*_ex_ were seen to deviate significantly from values determined under low salt (25 mM, 30 °C) conditions, where RD profiles were stronger and more well defined (Fig. S5). Thus, the high salt data set at 40 °C was fit while fixing Δω_i-Major_ for G6-C1’ and G6-C8 to the values obtained from RD measurements under low salt (25 mM, 30 °C) conditions (Table S5).

It should be noted that in addition to sensing conformational changes at the bp (active exchange), RD measurements at a given nucleus can also have contributions from conformational changes at neighboring bases (passive exchange) that need to be accounted for during fitting of the data (21, 62, 105). Such passive contributions are expected to be small (*R*_ex_ ~< 1 s^−1^) and not affect the extracted exchange parameters for most of the RD profiles measured in this study as they are strong (fitted *R*_ex_ values of > ~4 s^−1^), and given the small magnitude of the passive chemical shift changes (Δω_i-Major_ < ~0.5 ppm). Nevertheless, nuclei for which the RD profiles are intrinsically weak, with fitted *R*_ex_ values of < ~4 s^−1^ (G10-C1’ in A_6_-DNA at pH 6.8 25 °C, A16-C8 in A_2_-DNA at pH 6.8 25 °C, and G6-C1’ and G6-C8 in scaf2_TGC^GC^ at pH 5.4HS 30 °C and 40 °C) are likely to be influenced to a greater extent by such passive exchange contributions relative to nuclei with stronger RD profiles. These passive contributions could also cause deviations from the true exchange parameters for the active exchange when fitting the data assuming a two-state model. However, given the weak nature of the RD profiles, it is difficult to robustly fit them to a three-state exchange model to extract exchange parameters for both the active and passive exchange. Thus, we have fit these weak RD profiles (G10-C1’ in A_6_-DNA at pH 6.8 25 °C, A16-C8 in A_2_-DNA at pH 6.8 25 °C, and G6-C1’ and G6-C8 in scaf2_TGC^GC^ at pH 5.4HS 30 °C and 40 °C) assuming a two-state exchange model and suggest that the fitted exchange parameters be interpreted with caution, and have highlighted these weak data sets using open symbols and * in Fig. 3c, 4c, 4e, 5c and 10b.

The uncertainty in the exchange parameters was obtained using a Monte-Carlo scheme as described previously (106). All the fitting parameters have been summarized in Table S5. The sensitivity of the fit to the *R*_1ρ_ data to changes in *p*_i_ was examined by computing the reduced χ^2^ while fixing *p*_i_ to a range of different values (Fig. S6). Off-resonance *R*_1ρ_ profiles (Fig. 10b and Fig. S5) were generated by plotting (*R*_2_ + *R*_ex_) = (*R*_1ρ_ - *R*_1_cos^2^θ)/sin^2^θ, where θ is the angle between the effective field of the observed resonance and the z-axis, as a function of Ω_OBS_ = ω_OBS_ - ω_RF_, in which ω_OBS_ is the Larmor frequency of the observed resonance and ω_RF_ is the angular frequency of the applied spin-lock. Errors in (*R*_2_ + *R*_ex_) were determined by propagating the error in *R*_1ρ_ obtained as described above.

Alignment of initial magnetization during the Bloch-McConnell fitting was performed based on the *k*_ex_/Δω_i-Major_ ratio as described previously (40). Interestingly, RD profiles for A21-C1’ in A_6_-DNA at 25°C and 30 °C, G10-C1’ in A_6_-DNA at pH 6.8 25 °C, and G6-C1’ and C8 in scaf2_TGC^GC^ pH 5.4HS 30 °C and 40 °C could be better fit with a highly populated ES on the order of a few percent in slow exchange, while assuming alignment along the ground state effective field. Nevertheless, these highly populated solutions were excluded based on the lower populations measured previously for a wide range of sequence contexts in DNA (in the case of A21-C1’ in A_6_-DNA at 25 and 30 °C)(8), and alternative measurements at pH 5.4 (in the case of G10-C1’ in A_6_-DNA at pH 6.8 25 °C) and at low salt (G6-C1’ and C8 in scaf2_TGC^GC^ pH 5.4 30 and 40 °C) wherein the exchange parameters were robust (Fig. S6). Thus, the alignment of magnetization for A21-C1’ in A_6_-DNA at 25 and 30 °C, G6-C1’ and C8 in scaf2_TGC^GC^ pH 5.4HS 30 °C and 40 °C was fixed to be along the effective field of the population averaged state when plotting the reduced χ^2^ profiles as a function of *p*_i_ (Fig. S6).

#### ^13^C CEST

^13^C CEST experiments on the aromatic C8/C2 and C1’ carbons were carried out using a pulse sequence employing a selective excitation scheme in a 1D manner, as described previously (67, 107, 108). The ^13^C CEST experiments on the methyl carbon in hpGGACU^m6A6^(m^6^A6^C10^lb) were carried out in a 2D mode using a pulse sequence derived from prior ^13^C CEST pulse sequences (74, 107, 109).

The spin-lock power ranged from 10 to 70 Hz and from 10 to 50 Hz, respectively. The list of spin-lock power offset combinations used for CEST experiments can be found in Table S4. A relaxation time of 200 ms was used for all CEST experiments. The experimental conditions for all the ^13^C/^15^N CEST experiments (temperature, magnetic field, solvent) have been summarized in Table S3.

#### Analysis of CEST data

The analysis of CEST data was performed as described previously (67, 108). Briefly, 1D peak intensities were obtained in a manner similar to that for *R*_1ρ_. All intensities at a given radiofrequency (RF) power were normalized by the average of the intensities over the triplicate CEST measurements with zero relaxation delay using the same RF power, to obtain normalized CEST profiles. Normalized CEST profiles were plotted as a function of offset Ω = ω_RF_ - ω_OBS_, in which ω_OBS_ is the Larmor frequency of the observed resonance and ω_RF_ is the angular frequency of the applied spin-lock (Fig. S5). The error in the intensity for each RF power was obtained as the standard deviation of triplicate experiments with zero relaxation delay and same RF power. Exchange parameters of interest (*p*_i_, *k*_ex_, Δω_i-Major_) were then extracted by fitting the normalized CEST profiles to a two-state exchange model using the Bloch-McConnell equations. Combined fits of the CEST data for multiple nuclei were performed by sharing *p*_i_ and *k*_ex_.

Treatment of spin-lock inhomogeneity and alignment of the initial magnetization during CEST fitting was performed as described previously (108). Only GS magnetization was considered to be present at the start of the relaxation delay during CEST fitting for the C2/C8/C1’ and imino nitrogen spins, owing to the absence of the equilibration delay in the pulse sequence. GS and ES magnetization was considered to be equilibrated while fitting the methyl CEST data on hpGGACU^m6A6^(m^6^A6^C10^lb) as the pulse sequence employs hard pulses to excite spins (74). The sensitivity of the fit to the CEST data to changes in *p*_i_ was examined by computing the reduced χ^2^ while fixing *p*_i_ to a range of different values (Fig. S6). All the fitting parameters from CEST experiments have been summarized in Table S5. The errors of all the fitting parameters were estimated using 100 Monte Carlo iterations as described previously (67).

#### Imino ^1^H exchange

The kinetics of base opening were determined using a combination of experiments including a saturation recovery experiment to measure water proton *R*_1_ (*R*_1w_) and a magnetization transfer experiment to measure the exchange rate of the imino proton with water, as described previously (94). The relaxation delay times for measuring water proton *R*_1_ were 0.0, 0.4, 0.8, 1.2, 1.6, 2.0, 2.4, 2.8, 3.2, 3.6, 4.0, 4.4, 4.8, 5.2, 6.0, 7.0, 8.0, 9.0, 10.0, 12.0 and 15.0 sec. The series of relaxation delay times for all the imino ^1^H exchange measurements in this study are listed in Table S6.

#### Analysis of imino ^1^H exchange data

The net exchange rate (*k*_ex_) between imino and water protons can be measured by fitting the 1D imino ^1^H peak volume at each delay time (t) in the magnetization transfer experiment to Equation 2 shown below,

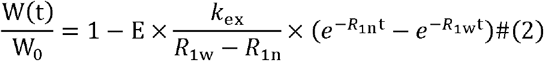

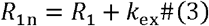

where W(t) is the imino ^1^H peak volume as a function of relaxation delay time t, W_0_ is the initial imino ^1^H peak volume at zero delay, W(t)/W_0_ is the normalized peak volume, E is the efficiency of the pulse for inverting water, *R*_1n_ represents the summation of imino ^1^H *R*_1_ and *k*_ex_. In the above equation, *R*_1w_ and E are fixed parameters while *k*_ex_ and *R*_1_ are fitting parameters. The error of all the fitting parameters was set to be equal to the standard fitting error, obtained as the square root of the diagonal elements of the covariance matrix of the fitting parameters. We also examined the sensitivity of the fitting to changes in the fitting parameter *k*_ex_ by fixing *k*_ex_ to a range of values and plotting the residual sum of squares (RSS) as a function of *k*_ex_ (Fig. S8b). All the fitting parameters have been summarized in Table S7. The efficiency of the inversion pulse was computed as follows,

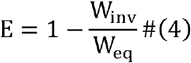

where W_inv_ and W_eq_ are the peak volumes of the water proton with and without applying the inversion pulse in the magnetization exchange experiment (with zero relaxation delay and no water suppression), respectively.

To obtain the population *p*_i_ of the base open state, the net *k*_ex_ between imino proton and water was measured in the presence of different effective ammonia concentrations (20 mM, 40 mM, 100 mM and 150 mM), and the inverse of *k*_ex_ (*τ_er_*=1/*k*_ex_) was linearly fit to the inverse of the effective ammonia concentration using Equation 5,

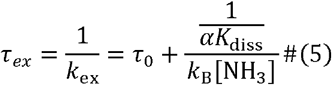

where *τ*_0_ is the inverse of the base opening rate (lifetime of the closed state), *α* is the base catalyst accessibility factor which is generally assumed to be 1 (110), [NH_3_] is the effective ammonia concentration, *K*_diss_ is the bp dissociation constant defined by,

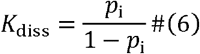

and *k*_B_ is the rate constant for exchange catalysis which is given by,

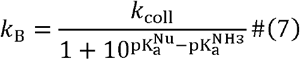

where *k*_coll_ is the bi-molecular collision rate constant between the imino proton and ammonia in the open state, and 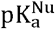 and 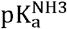 are the pKa for the imino proton in the open state and ammonia, respectively. Under the conditions used in our study *k*_B_ is equal to 2.5×10^8^ M^−1^s^−1^ at 25 °C (110).

Given the errors obtained for *k*_ex_ based on the covariance matrix as described above, Monte-Carlo simulations were performed to compute the errors in *K*_diss_. In particular, 1/*k*_ex_ at each ammonia concentration was sampled from a gaussian with mean equal to the average fit 1/*k*_ex_ value and standard deviation equal to the error in 1/*k*_ex_. The sampling was repeated for all ammonia concentrations for a given iteration, and the variation of 1/*k*_ex_ vs. 1/[NH_3_] was fit to compute a *K*_diss_ value. The procedure was repeated for 10,000 iterations and the mean and standard deviation of the resultant *K*_diss_ distribution were set to be equal to the mean and error of the fitted *K*_diss_. The region spanned by the family of straight lines thus obtained is colored blue in Fig. S8c. The error in *p*_i_ was determined by propagating the error in *K*_diss_ (Table S8).

To benchmark our implementation of the imino ^1^H exchange measurements in this study, we compared our measurements on T6 in G-DNA at 25 °C to a prior study (110) and obtained a consistent free energy for opening (Table S8). *p*_i_ values for T3, T5, T16, T17, T18, T19 and T21 in TBP-DNA at 15 °C were obtained from a prior study (72).

### UV melting

#### Experiments and sample conditions

Optical melting experiments were carried out on a PerkinElmer Lambda 25 UV/VIS spectrometer with an RTP 6 Peltier Temperature Programmer and a PCB 1500 Water Peltier System, and a Cary-100 UV-Vis spectrophotometer. At least three measurements were carried out for each DNA and RNA duplex using a sample volume of 400 μL in a Teflon-stoppered 1 cm path length quartz cell. The absorbance at 260 nm was monitored while the temperature was varied at a ramp rate of 1 °C/min.

#### Data analysis

The melting temperature (T_m_) and standard enthalpy change of melting (ΔH_melt,WT_ or ΔH_melt,Mod_) (Fig. 1d), respectively, were obtained by fitting the absorbance at 260 nm (A_260_) from the optical melting experiment to a two-state all-or-none model using Equation 8,

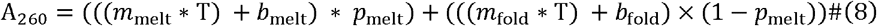

where *m*_melt_, *b*_melt_, *m*_fold_ and *b*_fold_ are coefficients describing the temperature-dependence of the extinction coefficients for the melted and folded species, respectively, and *p*_melt_ is the population of the melted duplex/hairpin species. *p*_melt_ for a WT duplex and hairpin are given by the following expressions (analogous expressions can also be written for melting of the modified nucleic acid Mod by replacing WT by Mod),

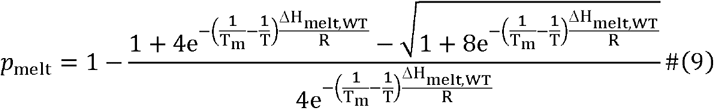

and

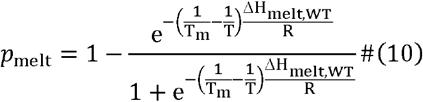

respectively, where T_m_ is the melting temperature (K), T is the temperature (K), and R is the gas constant (kcal/mol/K). It should be noted that *p*_melt_ is commonly denoted by the symbol α in the literature (64, 111).

The standard entropy change (ΔS_melt,WT_ or ΔS_melt,Mod_) and free energy change (ΔG_melt,WT_ or ΔG_melt,Mod_) of melting were computed using Equations 11 and 12 for a duplex and hairpin, respectively,

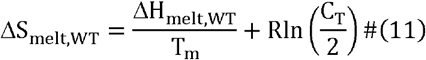

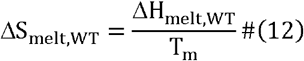

where C_T_ is total concentration of the duplex/hairpin. Using the obtained enthalpies and entropies, the free energy of melting was then computed as follows,

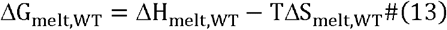

The mean values and uncertainties in T_m_, ΔH_melt,WT_, ΔS_melt,WT_ and ΔG_melt,WT_ were set to be equal to the average and standard deviation from multiple measurements (n >= 3).

For UV melting profiles of scaf2_AGG^m1GC^, scaf2_TGG^m1GC^, scaf2_AAG^AT^, scaf2_AAG^m1AT^, scaf2_AAG^Am3T^, scaf2_CAC^AT^, scaf2_GAC^AT^, scaf2_TAG^AT^, scaf2_TAG^m1AT^, scaf2_AAA^Am3T^ and scaf2_GAG^Am3T^ (Fig. S3), minor and reproducible deviations from two-state fits were observed in the lower/upper baselines which could potentially arise from the existence of multiple folded species in equilibrium with each other (52, 53). Due to the lack of knowledge about alternative folded species for the above samples, we have chosen not to fit the UV curves for them. Understanding the conformational dynamics that gives rise to such minor deviations and how that affects the interpretation of the melting data will be the subject of future studies.

### Comparison of thermodynamic preferences between NMR experiments and delta-melt

For a given equilibrium between the major and i^th^ minor conformation, Major ⇔ Minor, the thermodynamic preference for the conformational change (ΔG_conf_(i)) measured by NMR experiments was computed as follows,

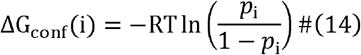

where T is the temperature (K), R is the gas constant (kcal/mol) and *p*_i_ is the population of the i^th^ minor conformation, respectively. Errors in ΔG_conf_(i) were obtained by propagating the error in *p*_i_ obtained from the NMR measurements.

The enthalpy (ΔH_conf_(i)) and entropy (ΔS_conf_(i)) differences between the major and i^th^ minor conformation were obtained by fitting ΔG_conf_(i) as a function of temperature to the following equation,

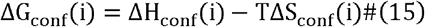

Errors in ΔH_conf_(i) and ΔS_conf_(i) were determined using a Monte-Carlo procedure. In particular, ΔG_conf_(i) for each point on the ΔG_conf_(i) vs. T plot was sampled from a Gaussian with mean and standard deviation equal to the mean value and error of the measured ΔG_conf_(i) values. Following gaussian sampling of all points on the plot, linear regression was performed to fit for ΔH_conf_(i) and ΔS_conf_(i). The procedure was repeated 10,000 times, and the mean and standard deviation of the resulting ΔH_conf_(i) and ΔS_conf_(i) distributions were set to be equal to the mean value and error of ΔH_conf_(i) and ΔS_conf_(i). The region spanned by the family of straight lines thus obtained is colored blue in Fig. S14.

Given conformational equilibria for the melting of WT and Mod nucleic acids, we will have the following equation according to Fig. 2c,

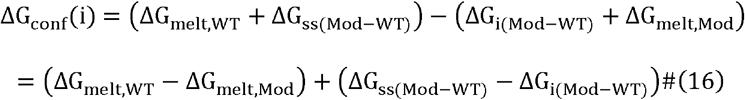

where ΔG_melt,WT_ and ΔG_melt,Mod_ are the melting energies of the folded WT and Mod species, ΔG_ss(Mod–WT)_ and ΔG_i(Mod–WT)_ are the free energies of modifying the melted single-stranded and the i^th^ folded minor conformation, respectively. We also define,

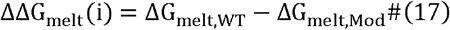

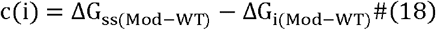

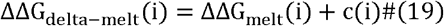

where ΔΔG_melt_(i) is the difference in melting energies of the WT and Mod species, and c(i) is the difference in free energies to modify the melted single-stranded (ss) and folded i^th^ minor species. Thus, ΔG_conf_(i) can be expressed as follows,

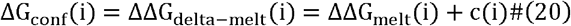

Errors in ΔΔG_melt_(i) were determined by propagating the errors in ΔG_melt,WT_ and ΔG_melt,Mod_ obtained from UV melting measurements. The error in c(i) was determined using a Monte-Carlo scheme as described below.

All the calibration curves of ΔG_conf_(i) versus ΔΔG_melt_(i) were linearly fit assuming that the slope was equal 1 using linear regression with Monte-Carlo iterations. In particular, each point on the ΔG_conf_(i) vs. ΔΔG_melt_(i) correlation plot was sampled from a gaussian with mean and standard deviation equal to the mean value and error of the measured ΔG_conf_(i) and ΔΔG_melt_(i) values along the y and x axes, respectively. Following gaussian sampling of all points on the plot, linear regression with a unit slope was performed to fit for c(i). The procedure was repeated 10,000 times, and the mean and standard deviation of the resulting c(i) distribution was set to be equal to the mean and error of c(i). The region spanned by the family of straight lines thus obtained is colored blue in Fig. 3c, 4c, 4e, 5c, 6c and 7c. The errors in ΔΔG_delta–melt_(i) were obtained by propagating the errors in ΔΔG_melt_(i) and c(i). The Pearson’s correlation coefficient (*r*) as well as the root mean square of error (RMSE) between ΔG_conf_(i) and ΔΔG_delta–melt_(i) = ΔΔG_melt_(i) + c(i) are reported for each comparison (Fig. 3c, 4c, 4e, 5c, 6c and 7c). A detailed discussion of the physical meaning of the intercept c(i) has been provided in Discussion S2.

Given that the enthalpy and entropy change are also state variables in a manner analogous to the free energy, equations analogous to Equation 20 above, can also be written for them.

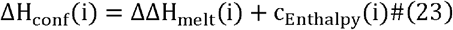

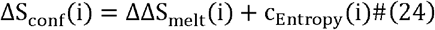

Errors in ΔΔH_melt_(i) and ΔΔS_melt_(i) were determined by propagating the errors in ΔH_melt,WT_ and ΔH_melt,Mod_, and ΔS_melt,WT_ and ΔS_melt,Mod_, obtained from UV melting measurements, as described above. A comparative analysis of enthalpy and entropy values obtained from NMR and delta-melt is provided in Fig. S14 and Discussion S5.

### Measurement of Hoogsteen bp cooperativity using delta-melt

Hoogsteen cooperativity can be quantified by the gained free energy (ΔG_coop,HG_) of forming tandem Hoogsteen bps at two adjacent sites (HG1+2) relative to forming Hoogsteen bps at each site (HG1 and HG2) independent of each other. ΔG_coop,HG_ is defined by the following equation (Fig. 8a),

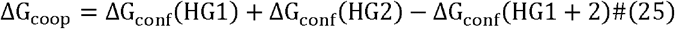

Substituting ΔG_conf_(i) = ΔΔG_melt_(i) + c(i) (Equation 20) to Equation 25 above yields,

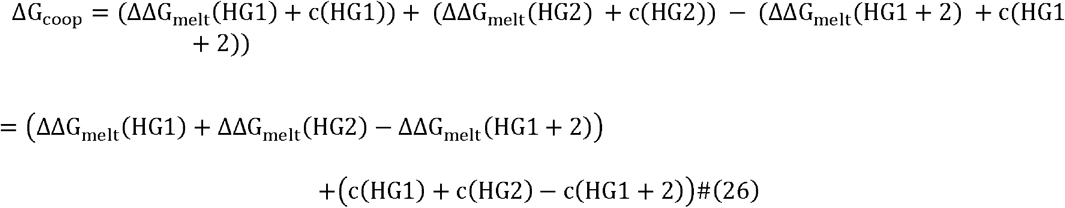

We also define,

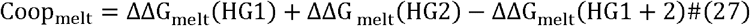

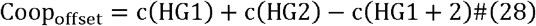

Therefore, we have

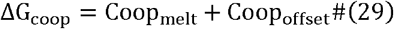

The Coop_melt_ term was determined using delta-melt by using duplexes containing *N*^1^-methylated purines at tandem bps, and their singly methylated counterparts (Fig. S9a). Errors in Coop_melt_ were obtained by propagating the errors in ΔΔG_melt_(i) values obtained as described above.

True estimation of Hoogsteen cooperativity requires knowledge of Coop_offset_. However, if we assume the offset term of double Hoogsteen c(HG1 + 2) and triple Hoogsteen c(HG1 + 2 + 3) is small, in a manner analogous to single Hoogsteen (Fig. 3c and 5c, Discussion S2), or that the offset for tandem Hoogsteen is the sum of offsets for single Hoogsteen bps, we will approximately have

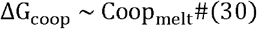

### Comparison of thermodynamic preferences with cancer signatures

This analysis assumes that all trinucleotide steps are equally populated in the genome. Given this assumption, the abundance of Hoogsteen bps at each step is given by the Boltzmann distribution over the thermodynamic preferences for formation of Hoogsteen bps obtained from delta-melt (Fig. 9b and Fig. S13). The free energies for the Hoogsteen bps thus obtained can be used to obtain their probabilities for formation, which were then compared to the probabilities of single base substitution mutations in cancer signatures obtained from the COSMIC mutation database v3.1 (87). The delta-melt derived values for this analysis (Fig. S16) were computed at 37 °C, and the probabilities of mutations in the cancer signatures were normalized that their sum was equal to 1.

## Acknowledgments

We would like to thank all members of the Al-Hashimi lab for critical input and Prof. Terry Oas for discussions with regards to UV melting experiments. We would also like to thank Dr. Richard Brennan (Duke University, USA) for providing access to the UV spectrophotometer for the UV melts performed in this study and Dr. Christopher Kreutz (University of Innsbruck, Austria) for providing us site-specifically labeled RNA phosphoramidites. The research in this study was funded by the US National Institutes of Health grants R01GM089846 and a Mathers Foundation grant to H.M.A.

## Code and Data Availability

All the raw data and scripts used for the analysis are available upon request to the authors.

