## Supplemental Information for "Measuring thermodynamic preferences to form non-native conformations in nucleic acids using melting experiments reveals a rich sequence-specific DNA conformational landscape"

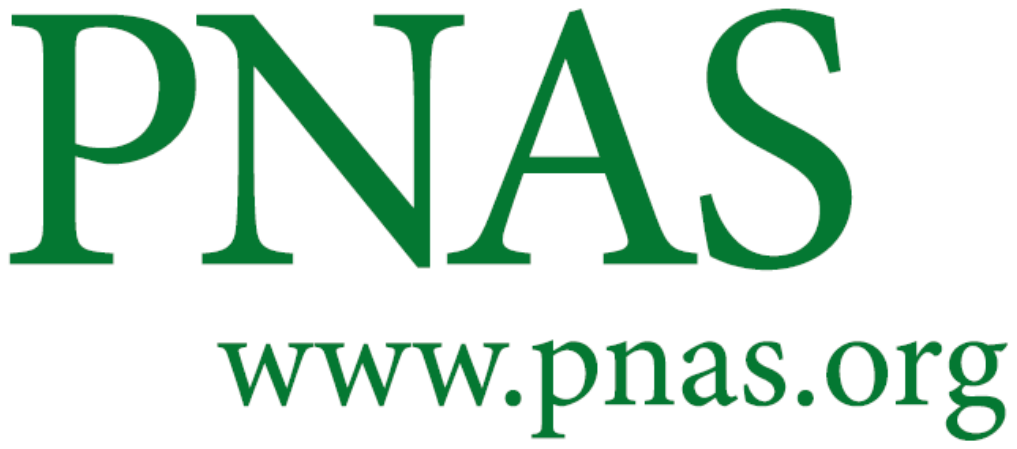

**Supplementary Information for**

Measuring thermodynamic preferences to form non-native conformations in nucleic acids using melting experiments reveals a rich sequence-specific DNA conformational landscape

Atul Rangadurai^1#^, Honglue Shi^2#^, Yu Xu^2#^, Bei Liu^1^, Hala Abou Assi^1^, John D. Boom^2,3^, Huiqing Zhou^1,4^, Isaac J. Kimsey^1,5^, and Hashim M. Al-Hashimi^1,2*^

1. Department of Biochemistry, Duke University School of Medicine, Durham, NC, USA

2. Department of Chemistry, Duke University, Durham, NC, USA

3. Department of Biomedical Engineering, Duke University, Durham, NC, USA

Present Address:

4. Department of Chemistry, Boston College, Boston, MA, USA

5. Nymirum, 4324 S. Alston Avenue, Durham, NC, USA

#These authors contributed equally to this work

*To whom correspondence should be addressed:

**This PDF file includes:**

Figures S1 to S14

Tables S1 to S9

Discussions S1 to S5

SI References

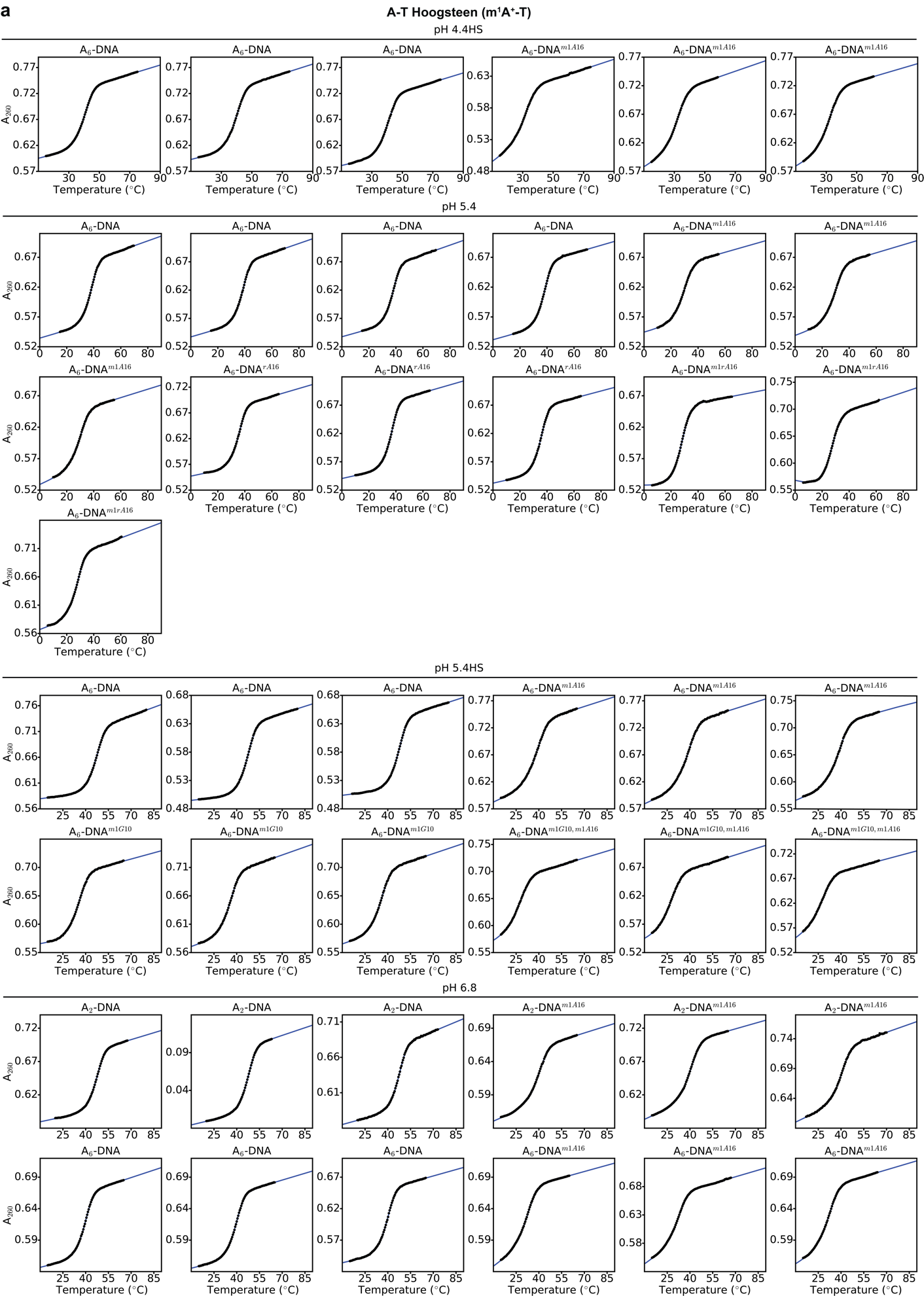

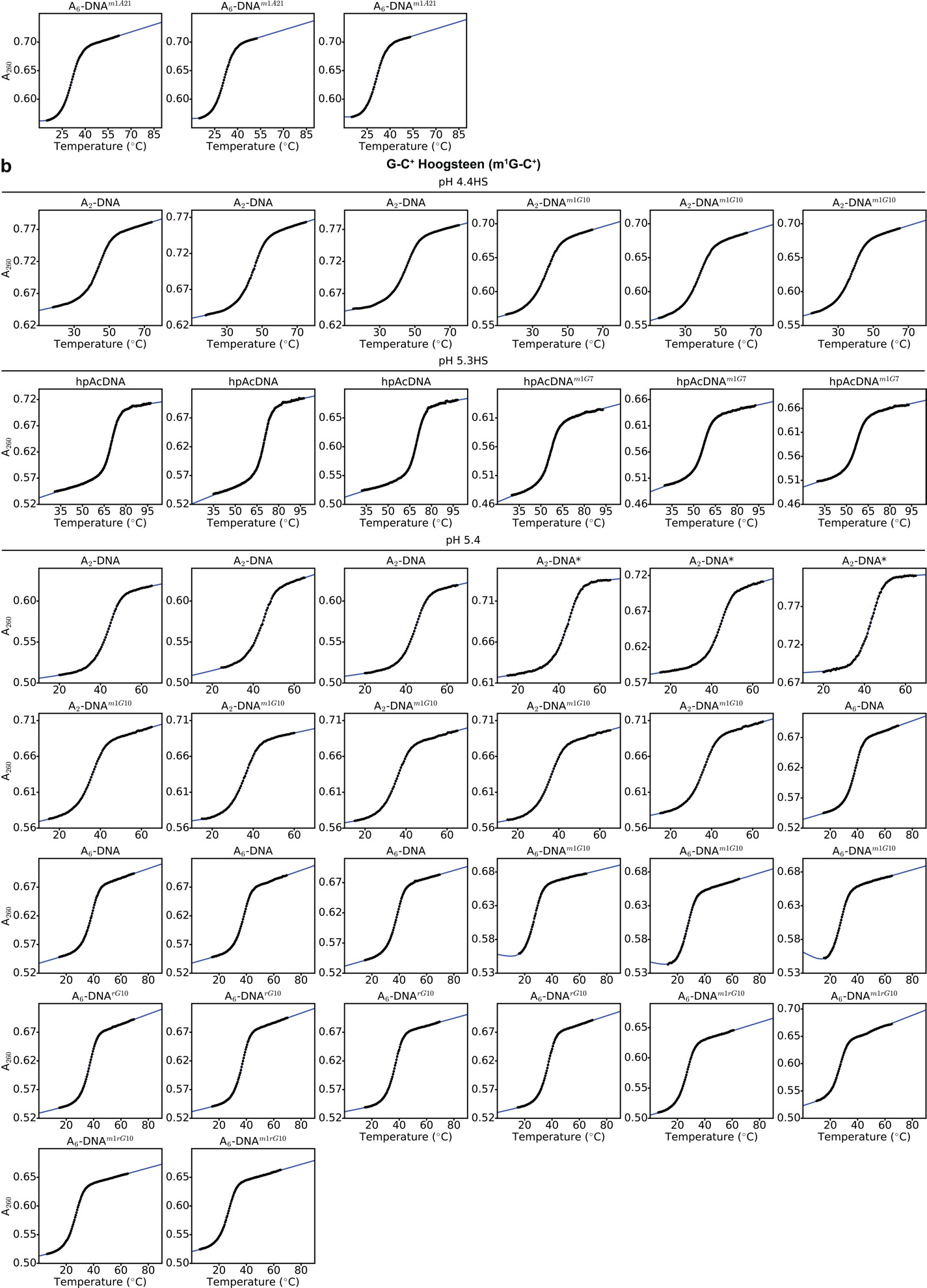

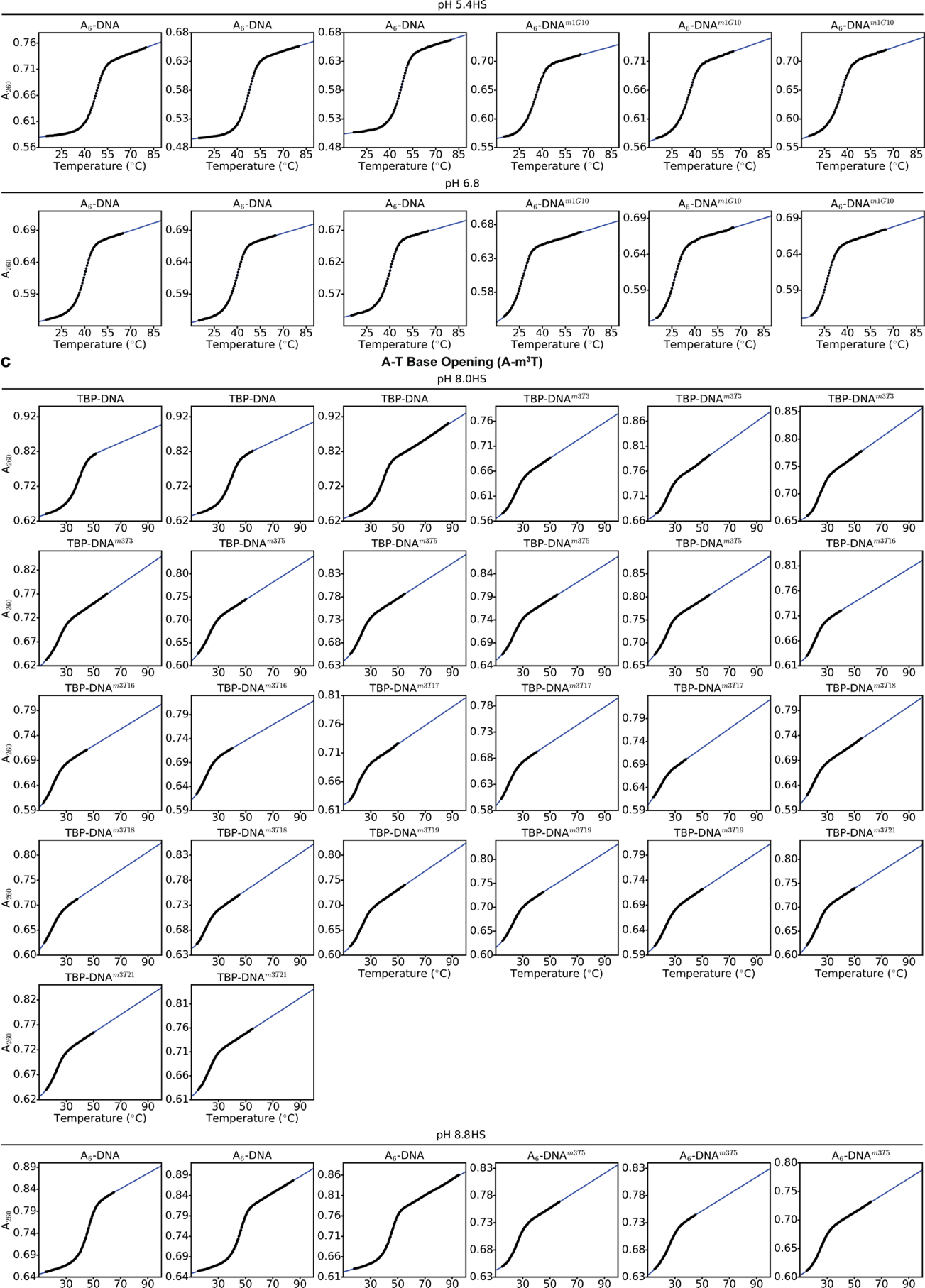

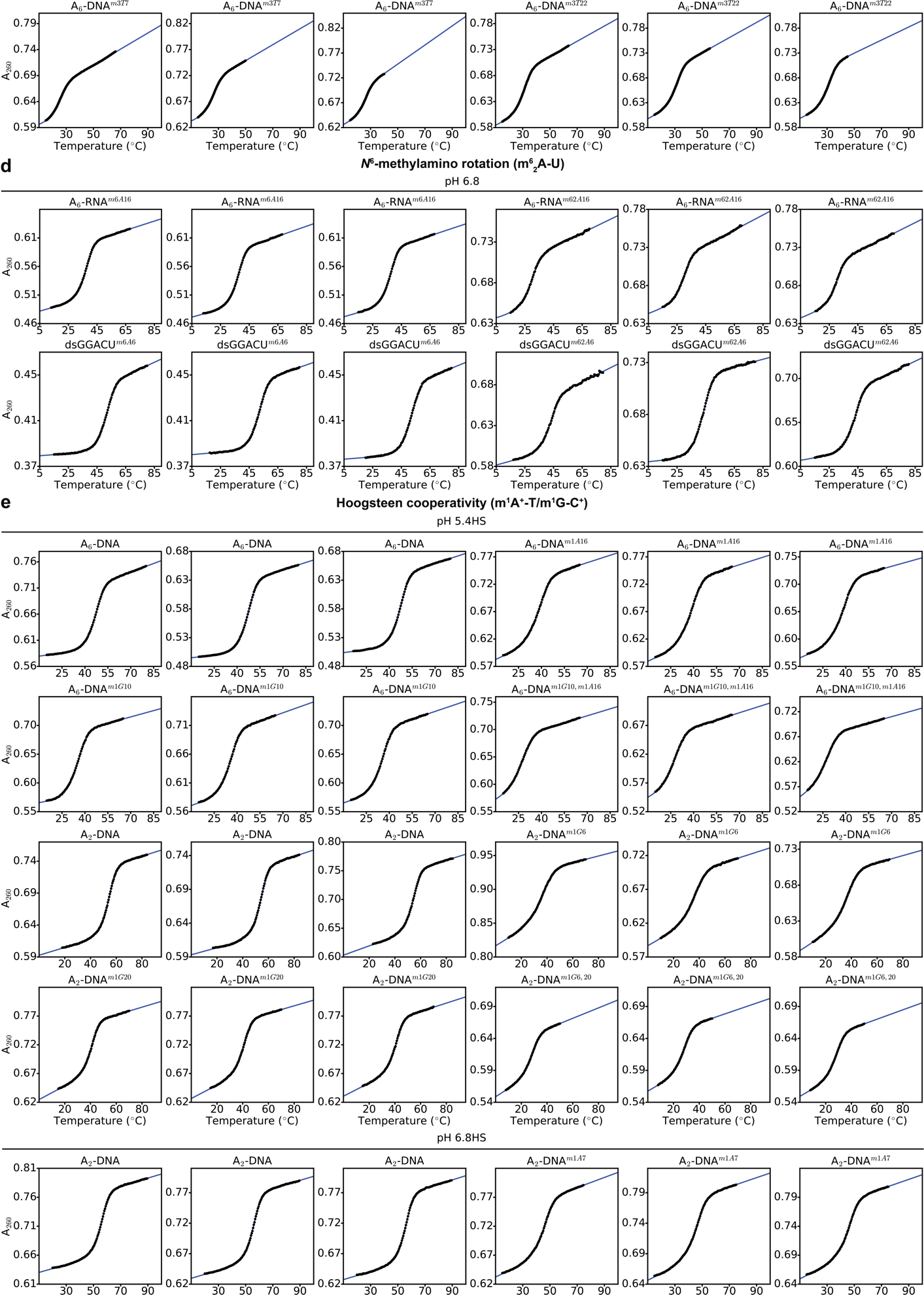

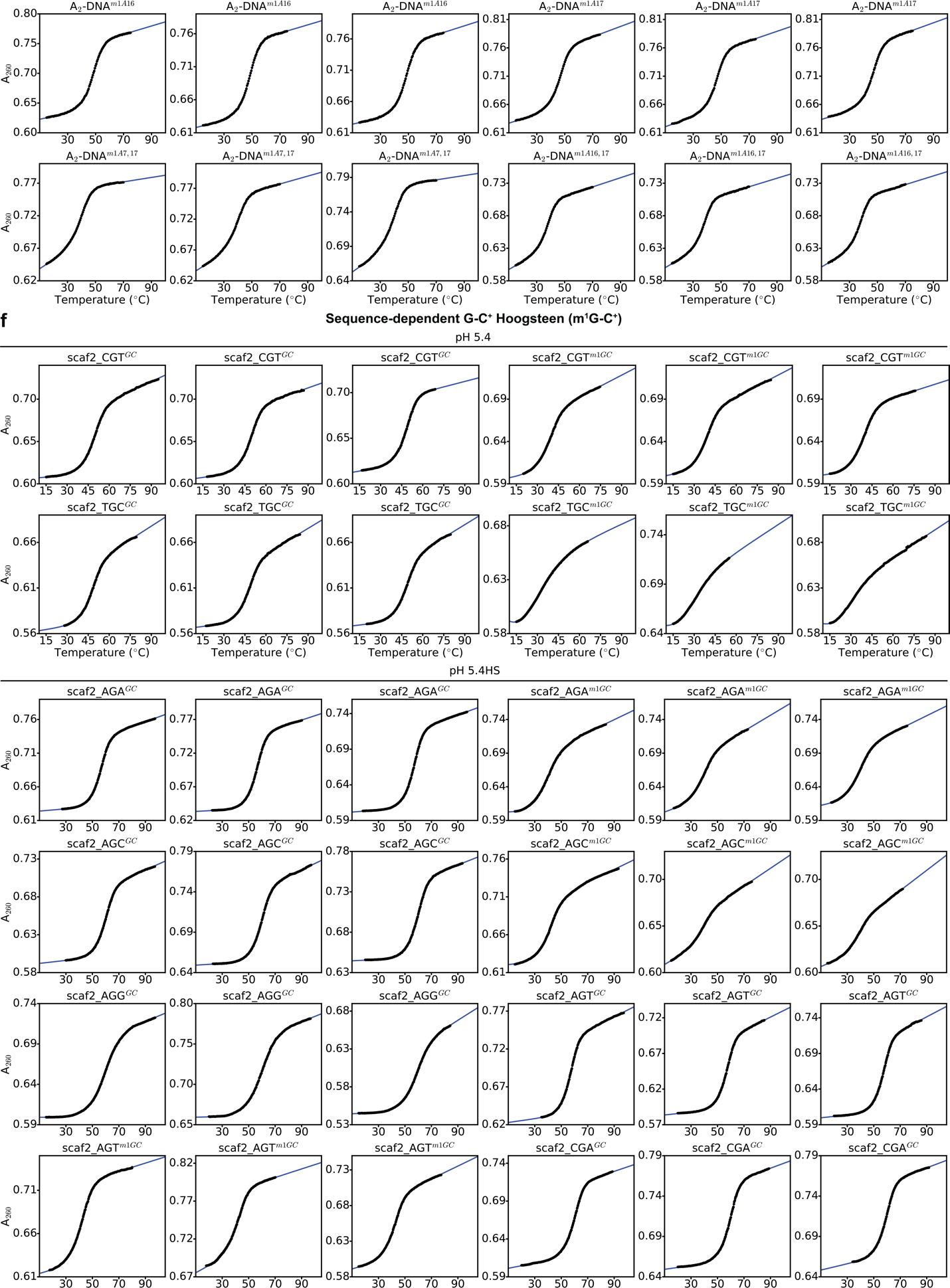

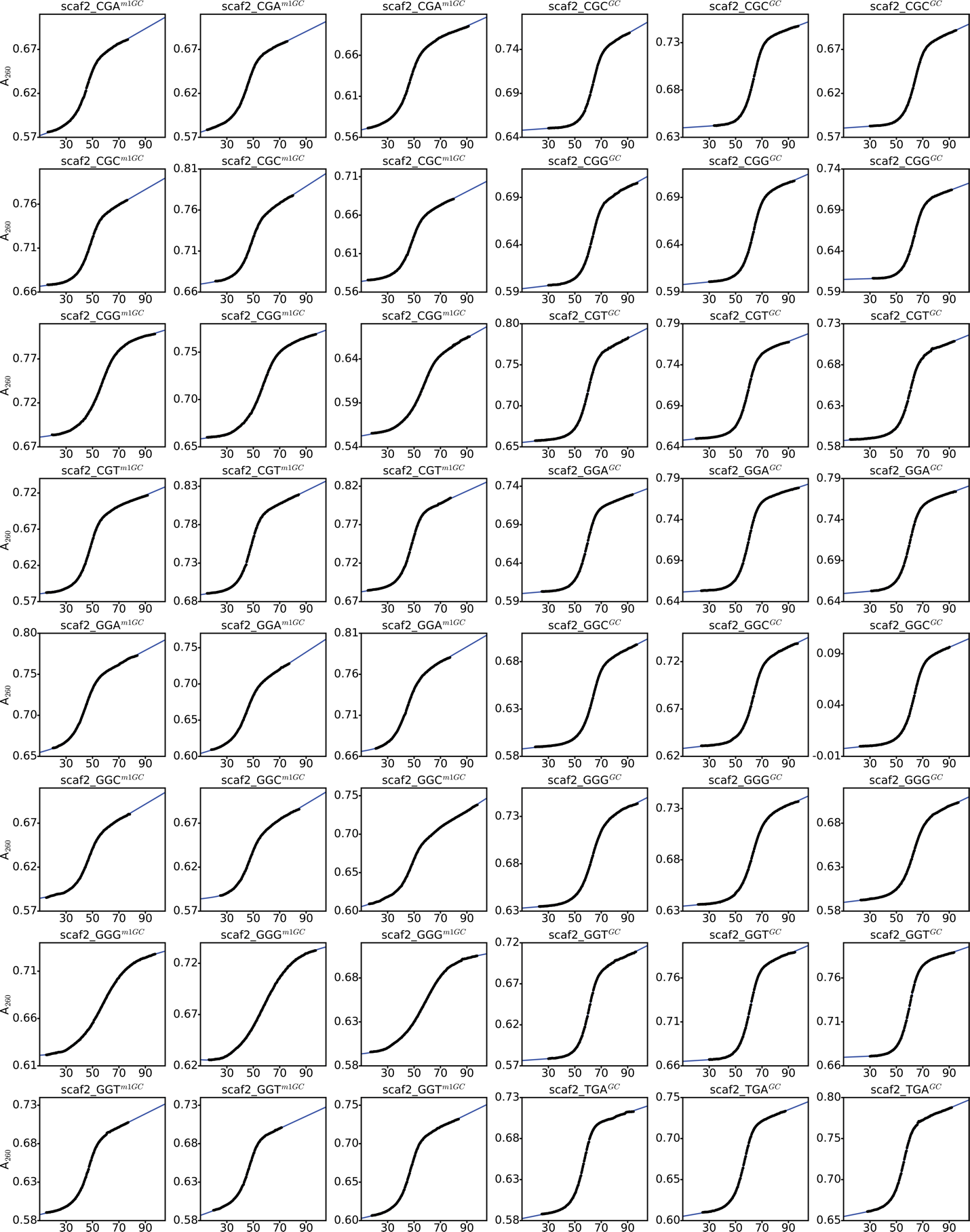

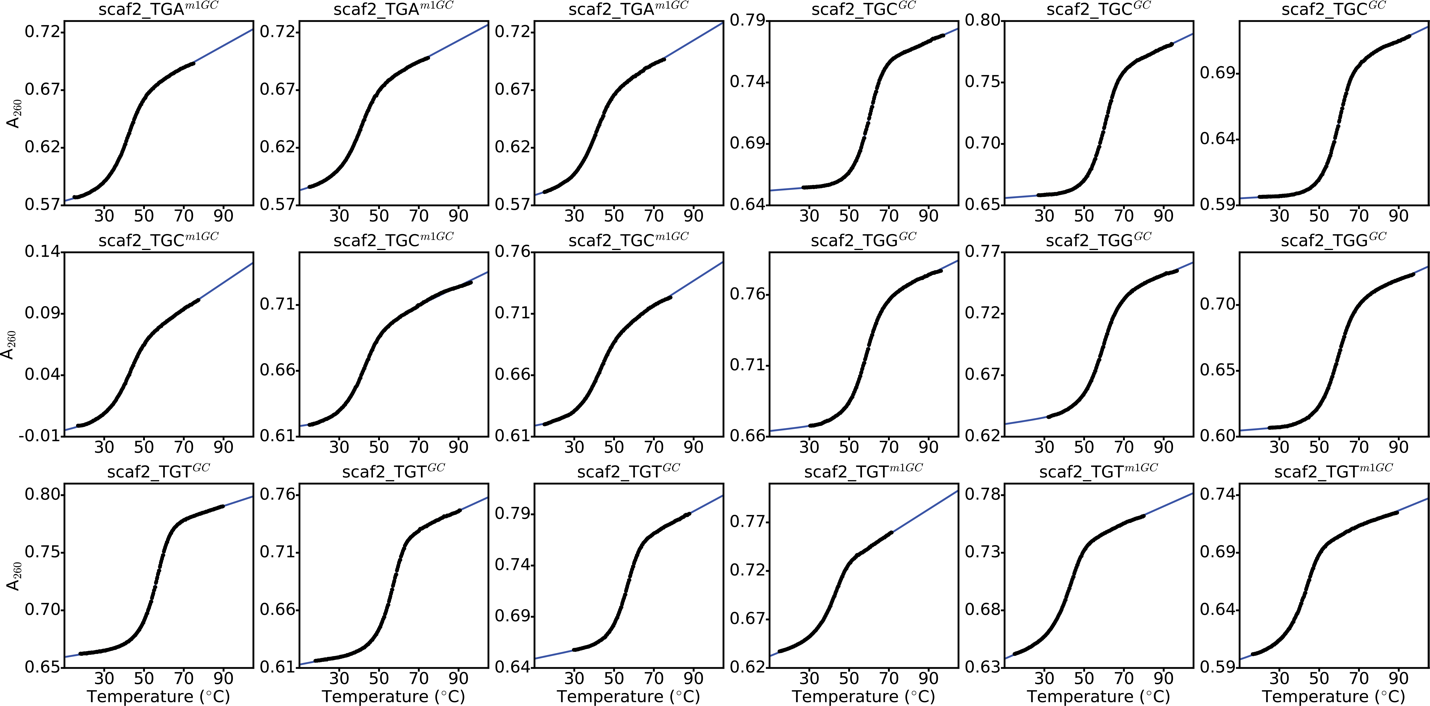

**Figure S1. UV-melting curves.** UV melting curves (black points) used to estimate thermodynamic preferences for (a) A-T and (b) G-C^+^ Hoogsteen bp formation, (c) A-T base opening, (d) *N*^6^-methylamino rotation in *N*^6^-methyl adenine (m^6^A), (e) cooperative formation of A-T/G-C^+^ Hoogsteen bps and (f) sequence-dependent formation of G-C^+^ Hoogsteen bps. Fits of the data to a two-state model are shown in blue lines (Methods). The modification used to measure the thermodynamic preferences is indicated in parentheses. The name of the constructs (secondary structures in Fig. S4) and buffer conditions are highlighted above the melts. A_260_ refers to the absorbance at 260nm. HS denotes high salt (> 25mM NaCl, Methods). The UV melting data for scaf2_AGG^m1GC^ and scaf2_TGG^m1GC^ are not shown above as they were not used to determine thermodynamic preferences, as they displayed non-two-state melting behavior (Fig. S3, Methods). All data were collected on a Perkin-Elmer UV-Vis spectrophotometer (Methods).

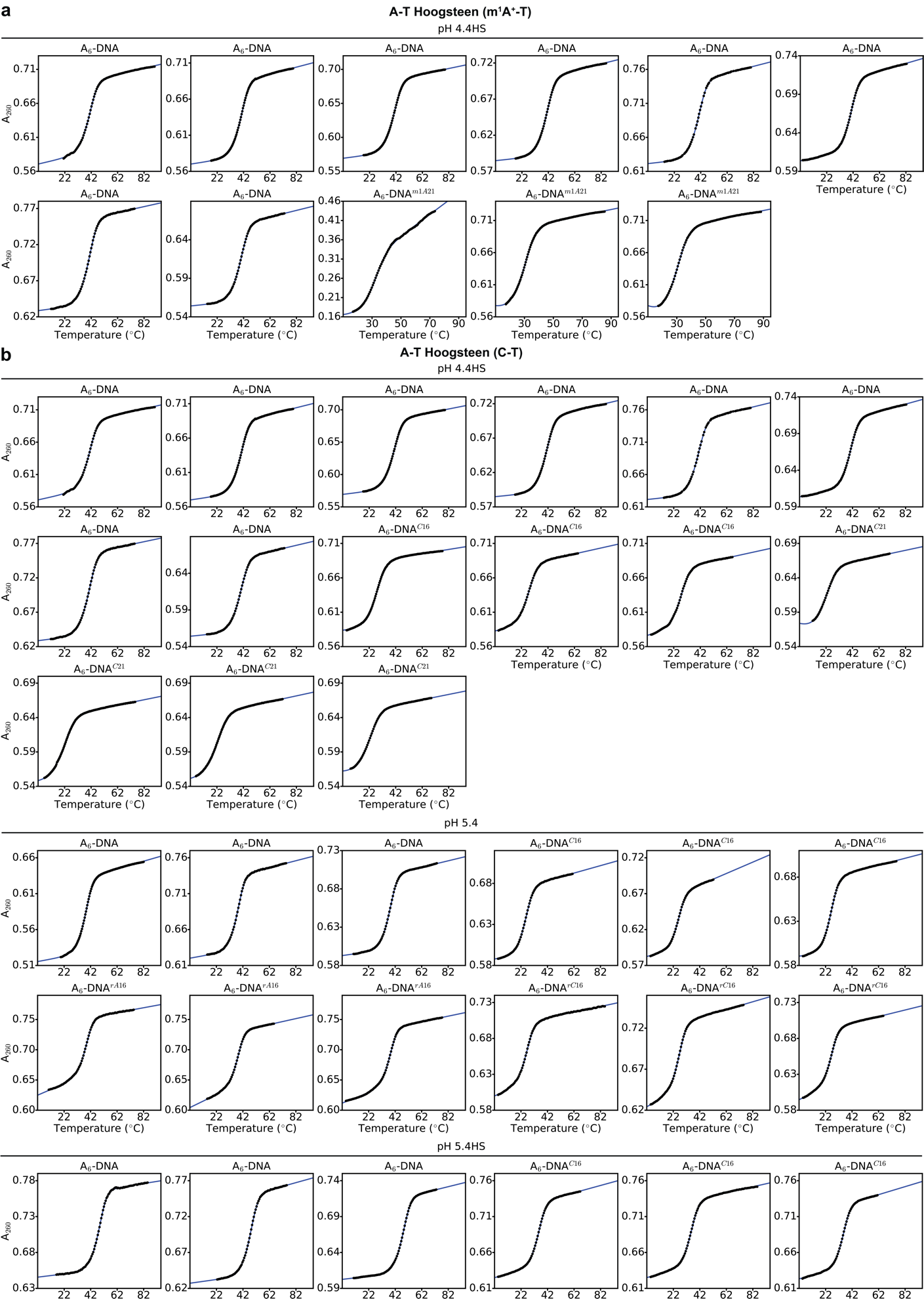

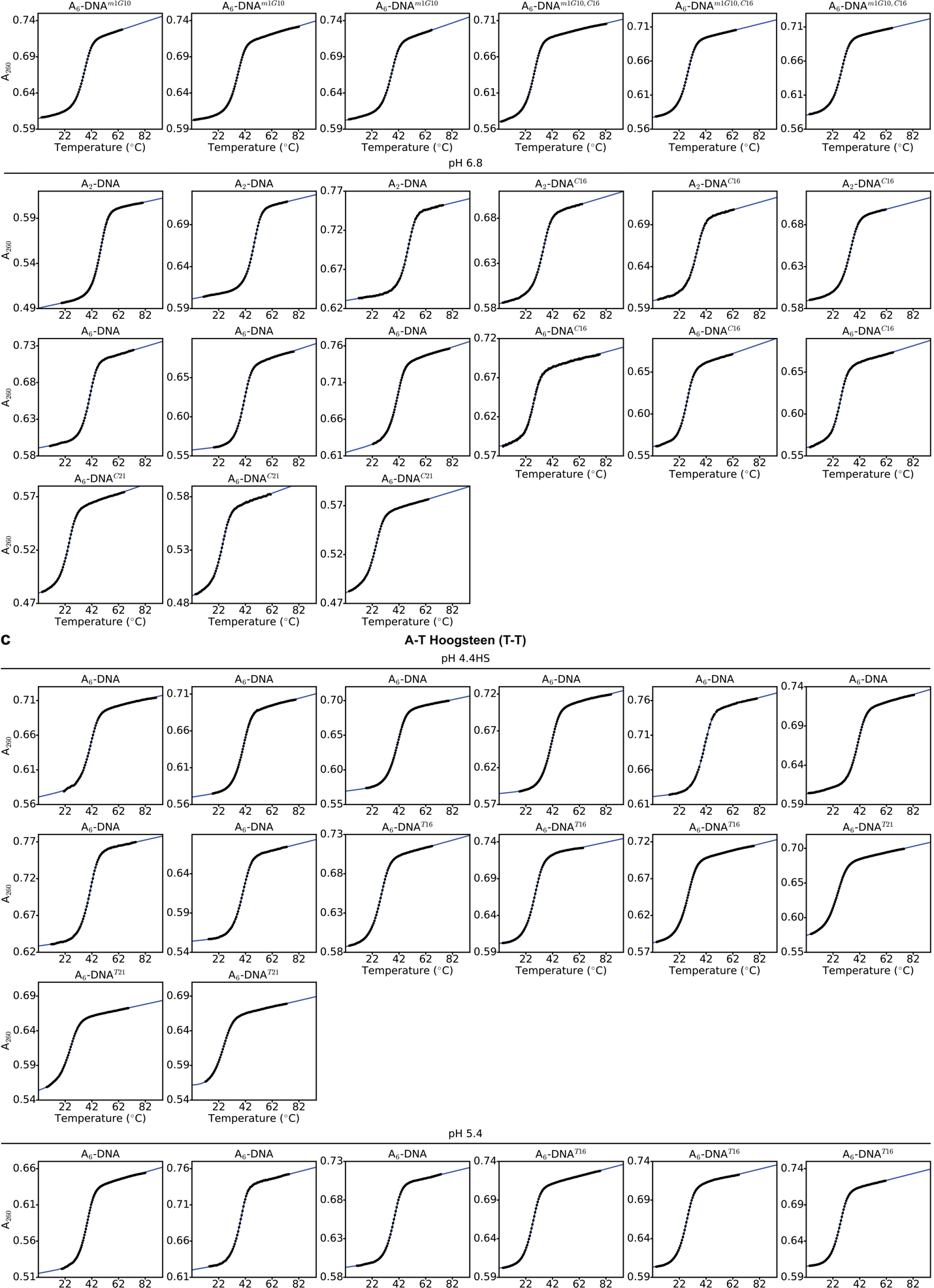

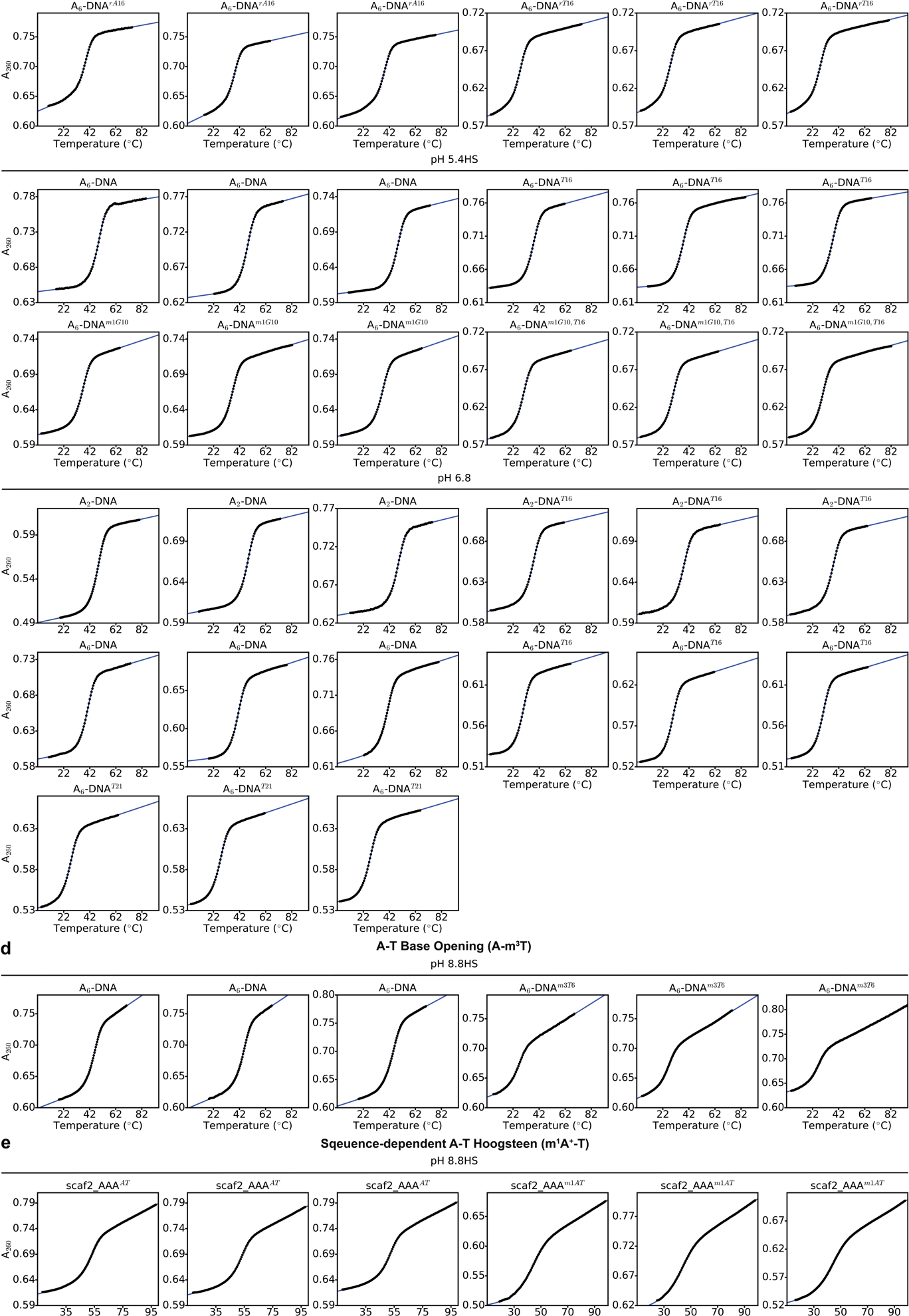

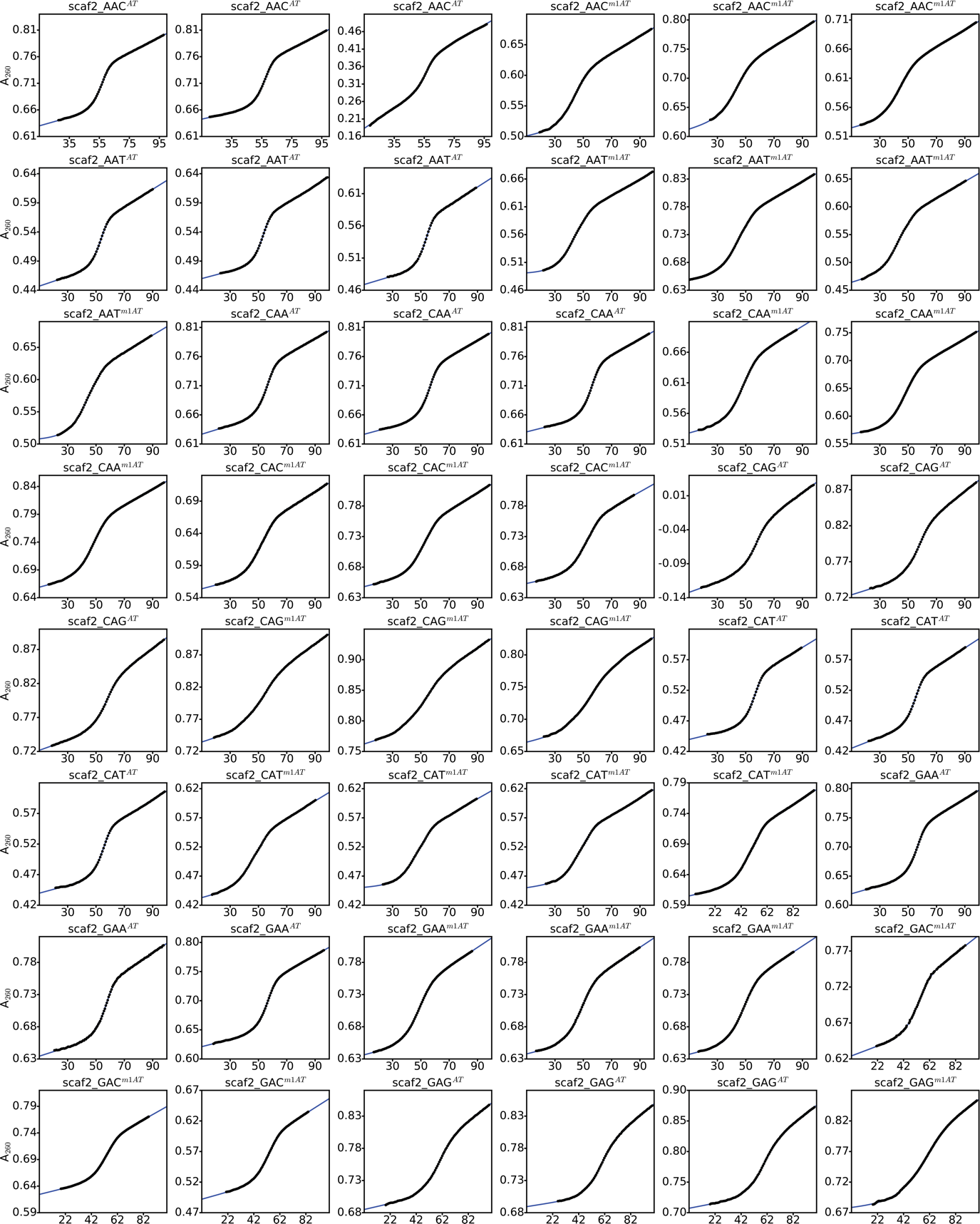

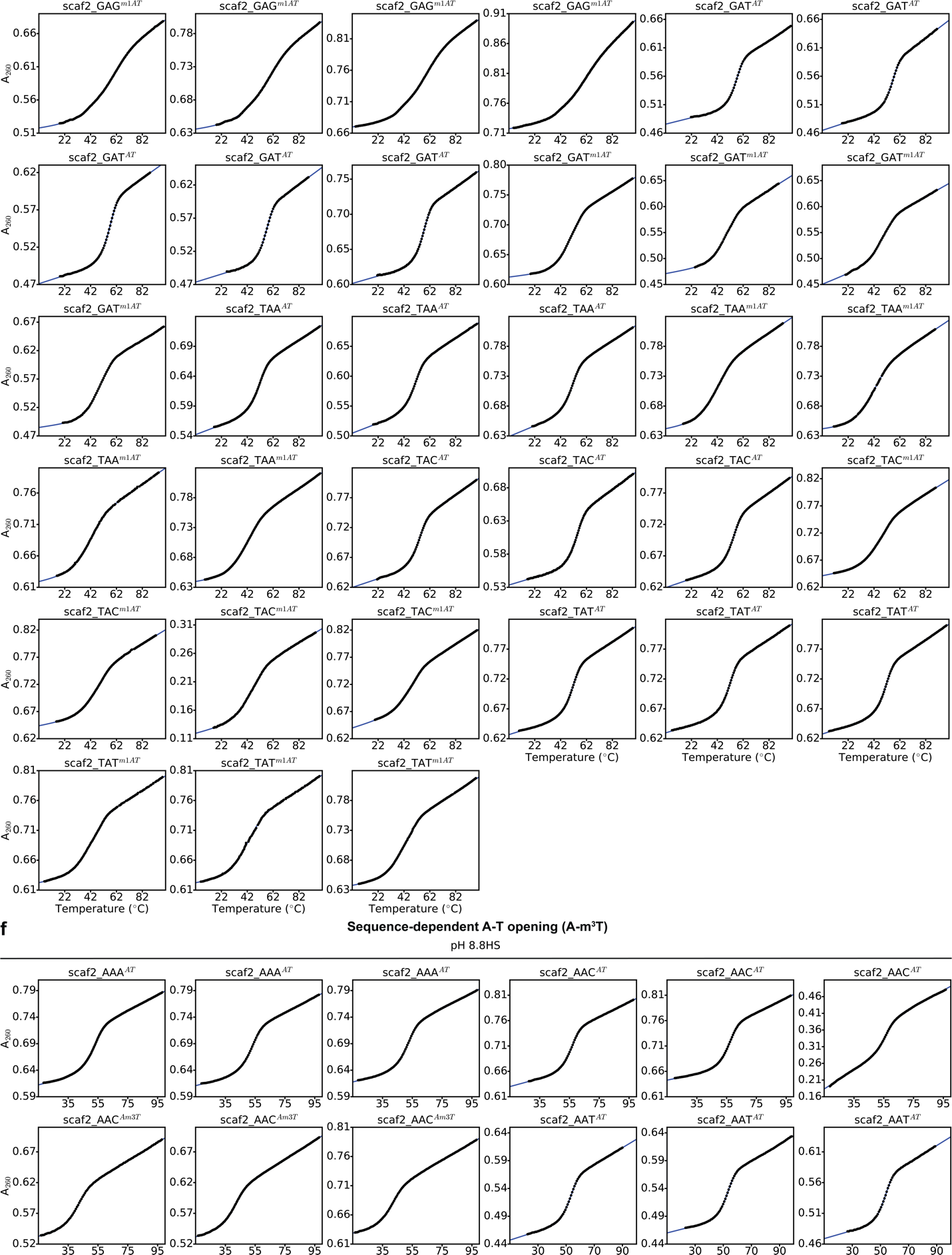

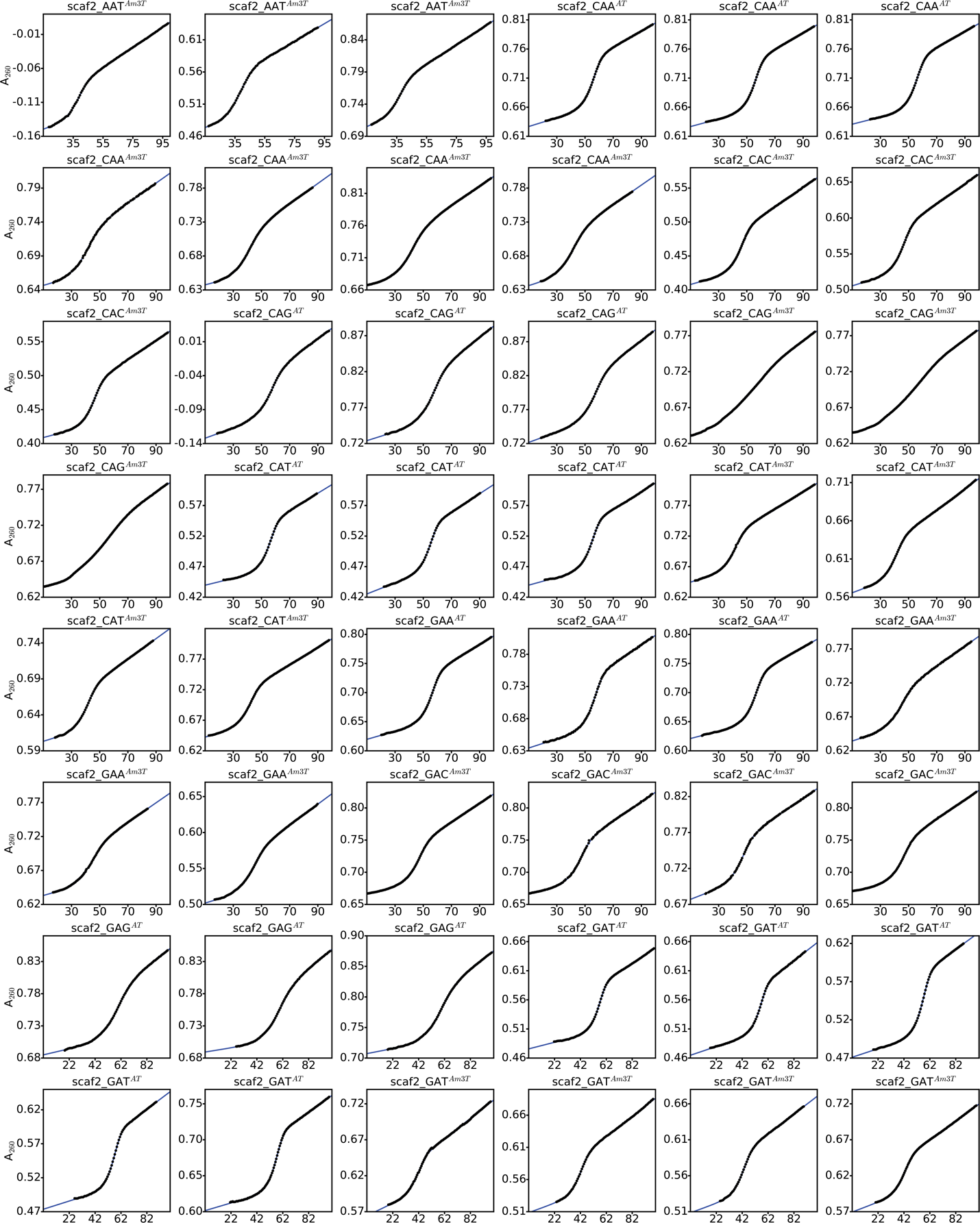

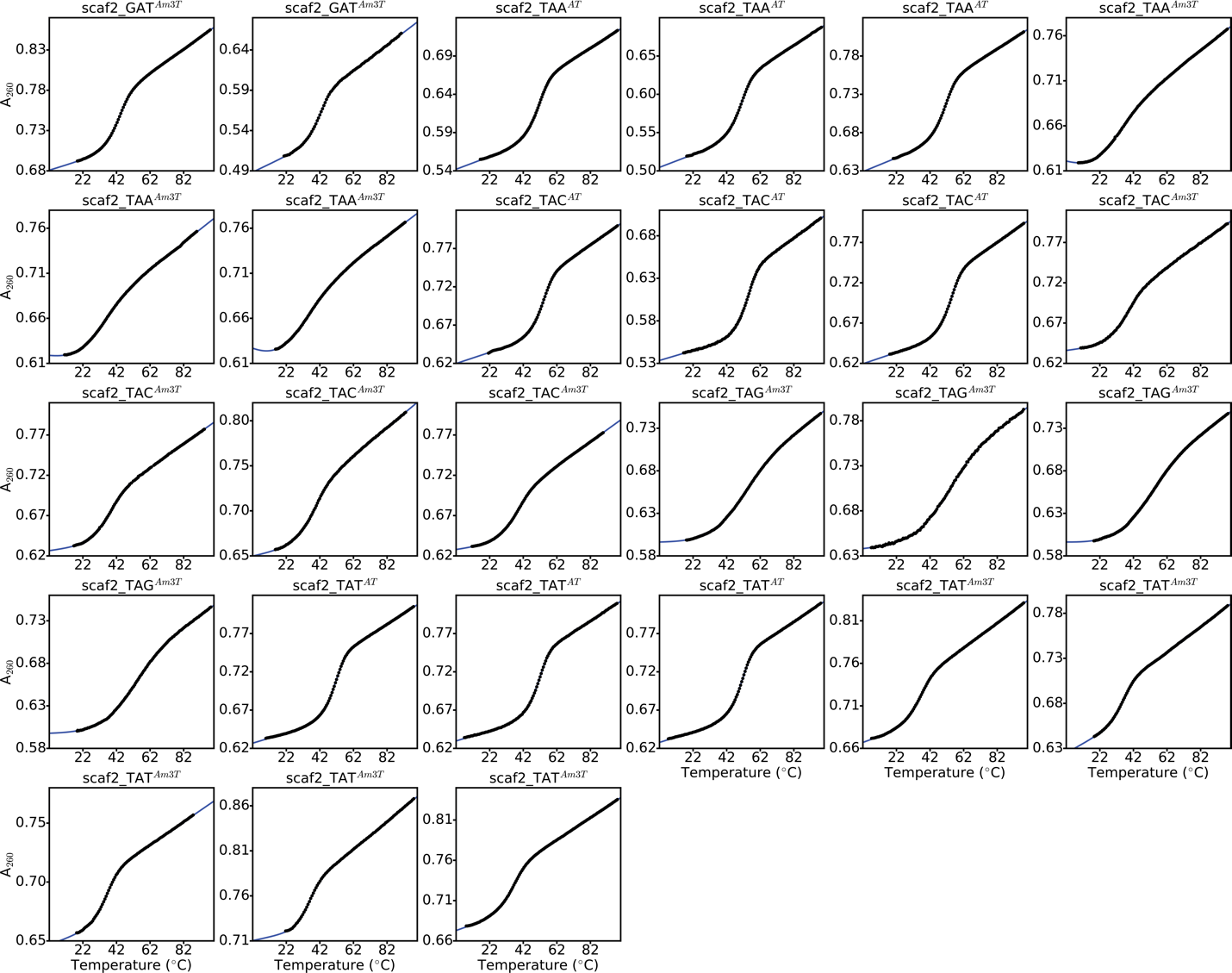

**Figure S2. UV-melting curves.** UV melting curves (black points) used to estimate thermodynamic preferences for (a, b and c) A-T Hoogsteen bp formation, (d) A-T base opening, and (e and f) sequence-dependent formation of A-T Hoogsteen bps and base opened states, respectively. Fits of the data to a two-state model are shown in blue lines (Methods). The modification used to measure the thermodynamic preferences is indicated in parentheses. The name of the constructs (secondary structures in Fig. S4) and buffer conditions are highlighted above the melts. A_260_ refers to the absorbance at 260nm. HS denotes high salt (> 25mM NaCl, Methods). The UV melting data for scaf2_AAG^AT^, scaf2_AAG^m1AT^, scaf2_AAG^Am3T^, scaf2_CAC^AT^, scaf2_GAC^AT^, scaf2_TAG^AT^, scaf2_TAG^m1AT^, scaf2_AAA^Am3T^ and scaf2_GAG^Am3T^ are not shown above as they were not used to determine thermodynamic preferences, as they displayed non-two-state melting behavior (Fig. S3, Methods). All data were collected on a Cary-100 UV-Vis spectrophotometer (Methods).

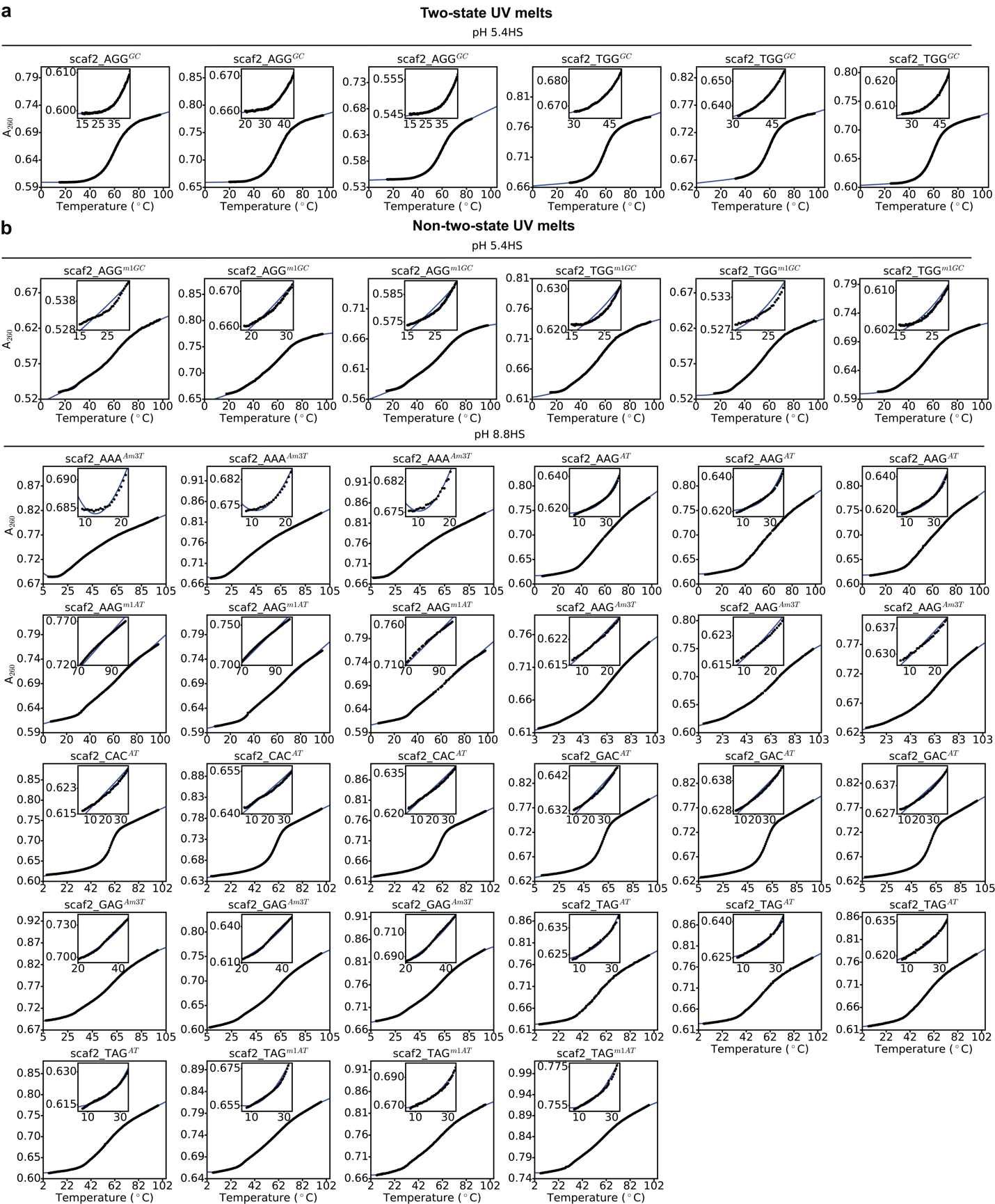

**Figure S3. UV melts showing two-state and non-two-state melting behavior.** Representative examples of UV melts (black points) showing (a) two-state melting behavior, and (b) all the UV melts in this study showing non-two-state melting behavior. Fits of the data to a two-state model are shown in blue lines (Methods). Also shown in the inset are zoomed in regions near the lower/upper baselines. The name of the constructs (secondary structures in Fig. S4) and buffer conditions are highlighted above the melts. A_260_ refers to the absorbance at 260nm. HS denotes high salt (> 25mM NaCl, Methods).

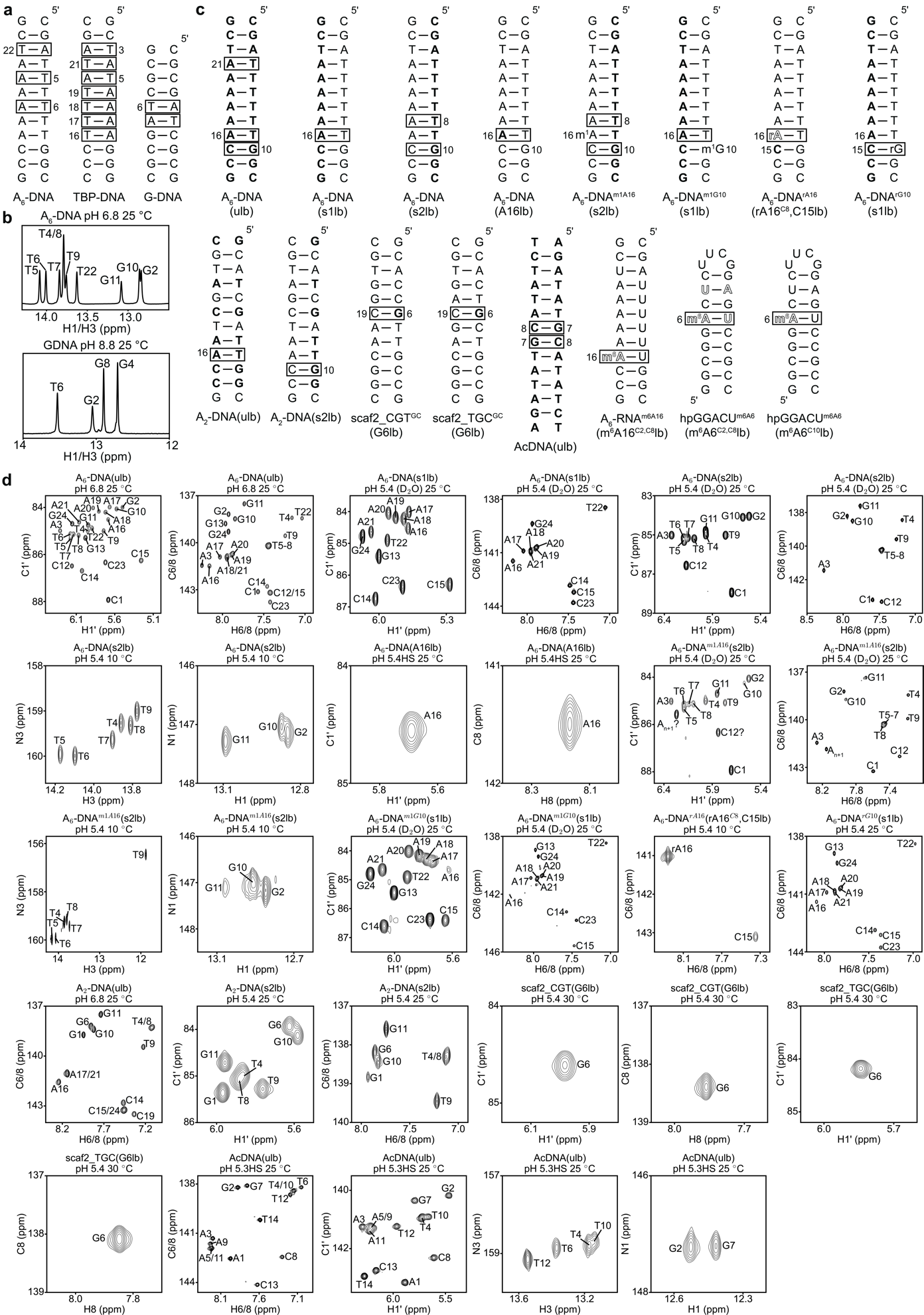

**Figure S4. Representative NMR spectra of the DNA and RNA constructs used for NMR measurements.** (a) DNA constructs used for imino proton exchange measurements. (b) Representative ^1^H 1D spectra of the imino region of A_6_-DNA and G-DNA. Spectra for TBP-DNA have been reported previously (1). (c) Isotope labeled DNA and RNA constructs used for *R*_1ρ_ and CEST measurements. Solid and dashed lines denote stable and labile Watson-Crick bps respectively. (d) Representative 2D [^13^C/^15^N, ^1^H] HSQC spectra of the isotope labeled DNA and RNA constructs. HS refers to high salt (> 25mM NaCl, Methods). Squares in panels a and c denote sites at which A-T base opening and A-T/G-C^+^ Hoogsteen bps were monitored using imino proton exchange and *R*_1ρ_/CEST, respectively. ^13^C, ^15^N uniformly labeled (A, C, T and G) nucleotides are denoted in bold while atom-specifically ^13^C/^15^N nucleotides are indicated using outlines. All other nucleotides are unlabeled. m^1^A, m^1^G, and m^6^A denote *N*^1^-methyl adenine, *N*^1^-methyl guanine and *N*^6^-methyl adenine, respectively. All nucleotides in DNA and RNA have deoxyribose and ribose sugars respectively, apart from rA/rG in DNA, which have a ribose sugar. The A and U nucleotides in hpGGACU^m6A6^(m^6^A6^C2,C8^lb) are site-specifically ^13^C labeled at C2 and C8, and ^15^N labeled at N3, respectively. Spectra for A_6_-RNA^m6A16^(m^6^A16^C2,C8^lb), hpGGACU^m6A6^(m^6^A6^C2,C8^lb) and hpGGACU^m6A6^(m^6^A6^C10^lb) were reported in another study (2).

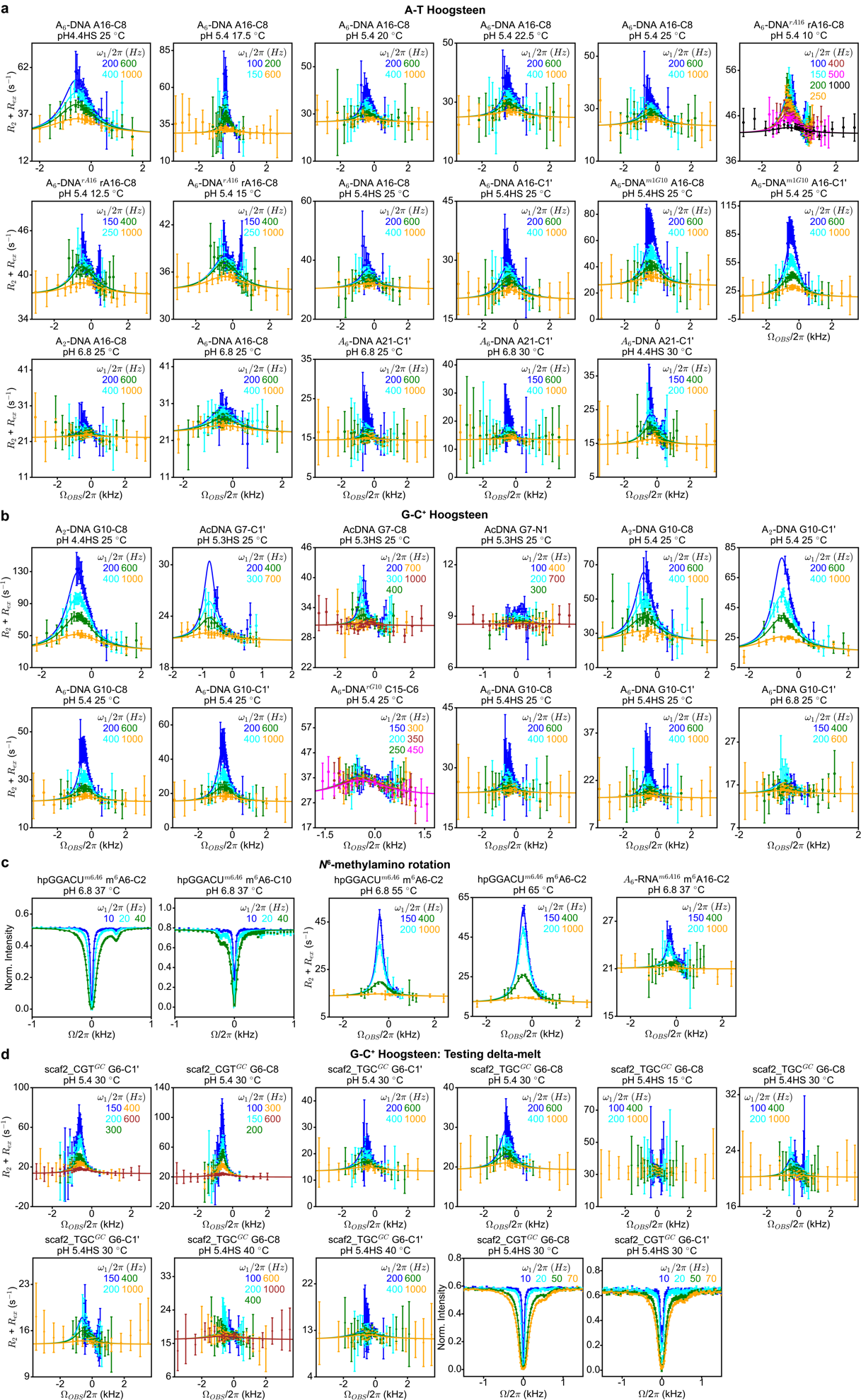

**Figure S5. Off-resonance *R*_1ρ_ relaxation dispersion and CEST profiles probing formation of minor conformational states.** *R*_1ρ_ relaxation dispersion and CEST profiles probing (a) A-T and (b, d) G-C^+^ Hoogsteen bp formation and (c) methylamino rotation in *N*^6^-methyl adenine (m^6^A). HS refers to high salt (> 25mM NaCl, Methods). Secondary structures and labeling schemes for the constructs used for the above measurements are provided in Fig. S4 and Table S3. RF powers for *R*_1ρ_ and CEST are color-coded. Two-state individual/shared fits (Table S5, Methods) of the data to the Bloch-McConnell equations are shown as solid lines. Error bars for the *R*_1ρ_ and CEST data were obtained using a Monte-Carlo based approach as described in the Methods. Error bars for the CEST intensities are smaller than the data points.

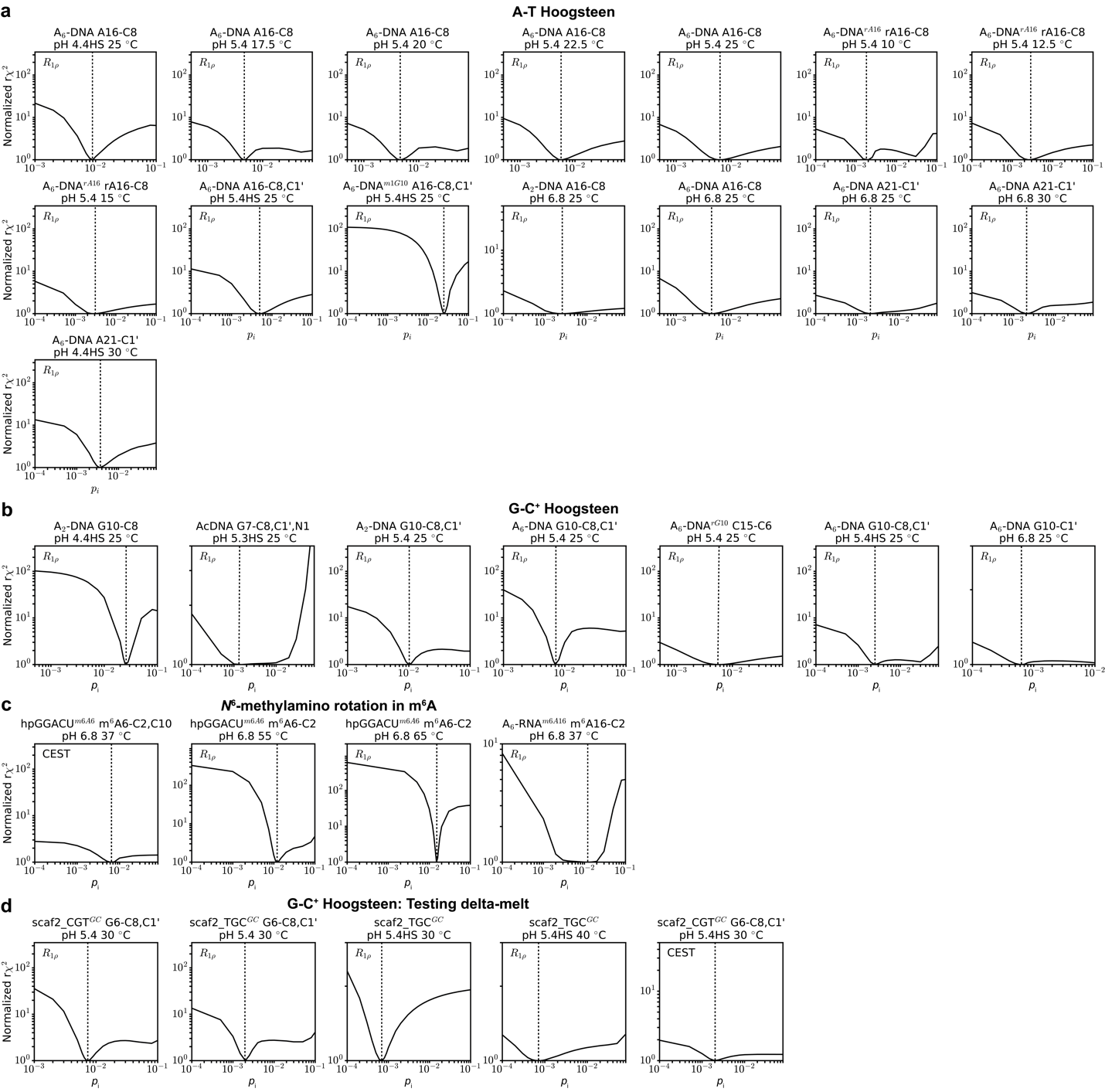

**Figure S6.** **Degeneracy analysis evaluating how well the population of the minor state (*p*_i_) is determined using the *R*_1ρ_ or CEST data.** Shown is the normalized reduced $\chi^{2}$ (${r\chi}^{2}$) as a function of fixing *p*_i_ to different values when fitting *R*_1ρ_ or CEST profiles probing (a) A-T and (b, d) G-C^+^ Hoogsteen bp formation and (d) methylamino rotation in *N*^6^-methyl adenine (m^6^A). HS refers to high salt (> 25mM NaCl, Methods). The absolute r$\chi^{2}$ values obtained from fitting the *R*_1ρ_ or CEST profiles were normalized such that r$\chi^{2}$= 1 for the best fit. The values of *p*_i_ obtained from free fitting the *R*_1ρ_ or CEST profiles are indicated using a dashed line. Differences in the shapes of the r$\chi^{2}$ profiles for different data sets could be related to the differences in experimental error (Methods) or sampling of the data points in the measurements (Fig. S5, Table S4).

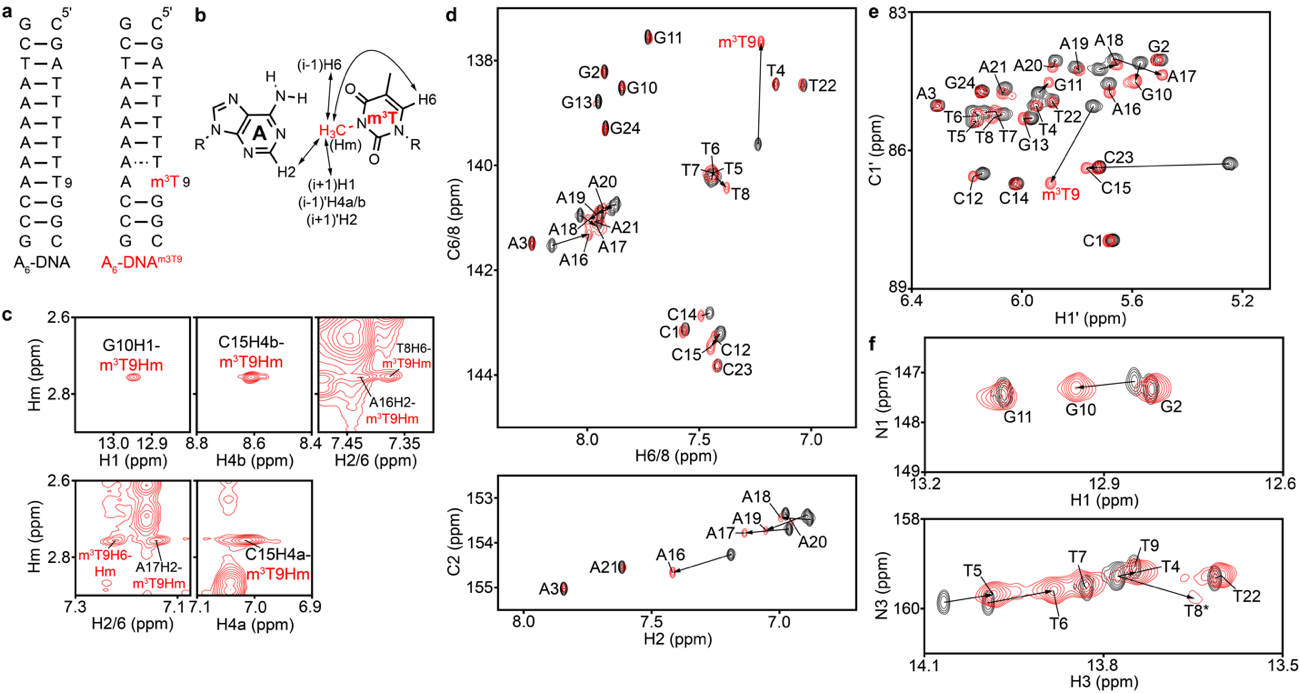

**Figure S7. NMR spectra of A_6_-DNA and A_6_-DNA^m3T9^.** (a) The A_6_-DNA and A_6_-DNA^m3T9^ duplexes. Solid and dashed lines denote stable and labile Watson-Crick bps, respectively. (b) Chemical structure of the A-m^3^T bp with observed NOE connectivities (see panel c) indicated using arrows. i-1, i and i+1 denote the residue preceding m^3^T (T8), the m^3^T residue itself and the residue after m^3^T (G10) in the 5' to 3' direction. (i-1)', i' and (i+1)' denote the residue preceding the adenine complementary to m^3^T (C15), the adenine itself, and the residue following it (A17) in the 5' to 3' direction. (c) Regions of the 2D [^1^H, ^1^H] NOESY spectra of A_6_-DNA^m3T9^ showing cross peaks between the *N*^3^-methyl protons (Hm) of m^3^T9 with imino, amino and aromatic protons of neighboring bases. (d-f) Overlays of 2D [^13^C, ^1^H] HSQC spectra of the aromatic (d) and C1' regions (e), and 2D [^15^N, ^1^H] HMQC spectra of the imino region (f) of A_6_-DNA (black) and A_6_-DNA^m3T9^ (red) in NMR buffer (15 mM sodium phosphate, 25 mM sodium chloride, 0.1 mM EDTA) at pH 6.8 and 25 °C.

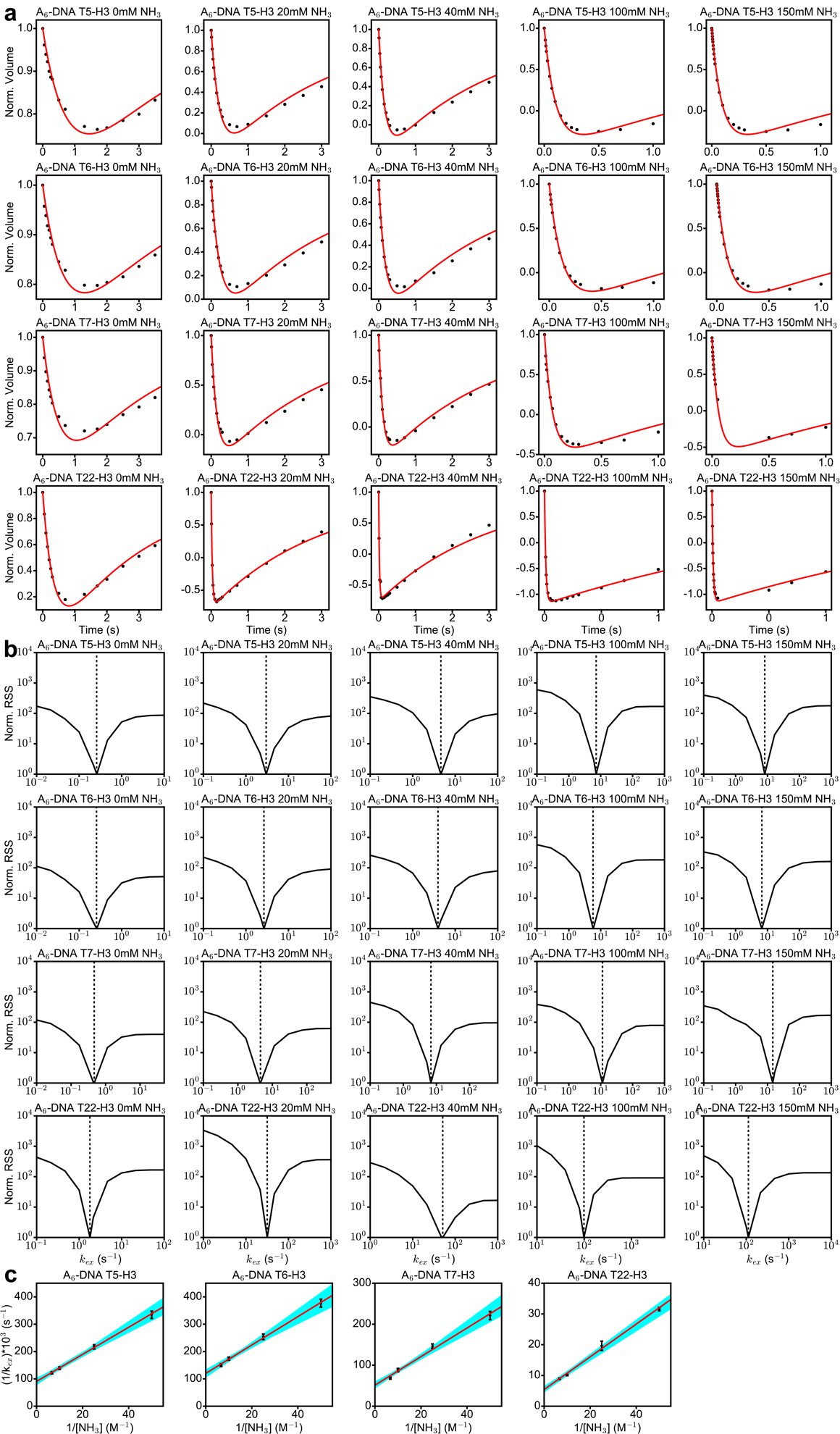

**Figure S8. Imino proton exchange measurements on DNA.** (a) Recovery of longitudinal imino proton magnetization post inversion. Shown is the normalized volume of the imino proton as a function of relaxation delay, computed as described in Methods. Experimental data points are in black and a fit to the data to Equation 2 is in red (Methods). (b) Quality of fit for the inversion recovery curves as a function of changing the net imino proton exchange rate *k*_ex_. RSS denotes the residual sum of squares and was computed as described in the Methods. Dashed lines denotes the value of *k*_ex_ obtained from the fitting. (c) Variation of the inverse of the exchange rate with the inverse of the effective ammonia concentration. Data points are in black and a linear fit of the data to Equation 5 (black line) was computed as described in the Methods. Error bars for *k*_ex_ were obtained from the fitting error of the inversion recovery curves in (a), as described in Methods. Blue shaded region denotes the family of lines obtained from iterative linear regression fits of the data using a Monte-Carlo sampling protocol (Methods).

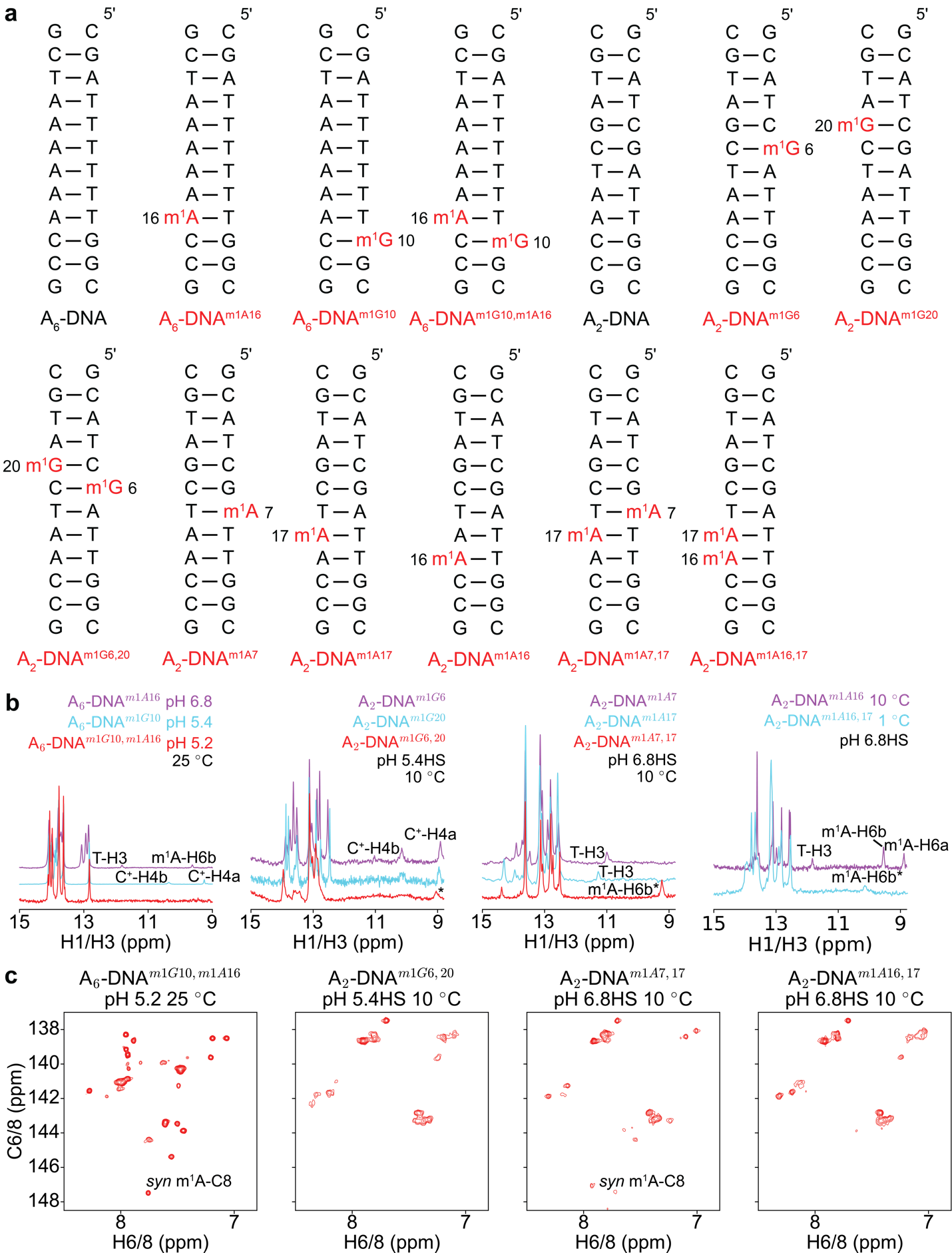

**Figure S9. NMR spectra of DNA duplexes with single and tandem *N*^1^-methylated purines.** (a) Duplexes used to examine cooperativity of Hoogsteen bp formation using delta-melt. m^1^A and m^1^G (red) denote *N*^1^-methyl adenine and guanine respectively. All nucleotides are unlabeled. (b) Overlay of ^1^H 1D spectra of the imino region of the duplexes with single and tandem *N*^1^-methylated purines. (c) 2D [^13^C, ^1^H] HSQC spectra of the aromatic region of DNA constructs with tandem *N*^1^-methylated purines. The downfield shifted m^1^A^+^-C8 resonance characteristic of a *syn* conformation of m^1^A^+^ in an m^1^A^+^-T Hoogsteen bp is indicated (3). HS corresponds to high salt (> 25 mM NaCl, Methods). In addition to NaCl, the buffers used for NMR measurements contained 15 mM sodium phosphate, 0.1 mM EDTA in a 90% H_2_O:10% D_2_O mixture. * denotes tentative or unknown assignments.

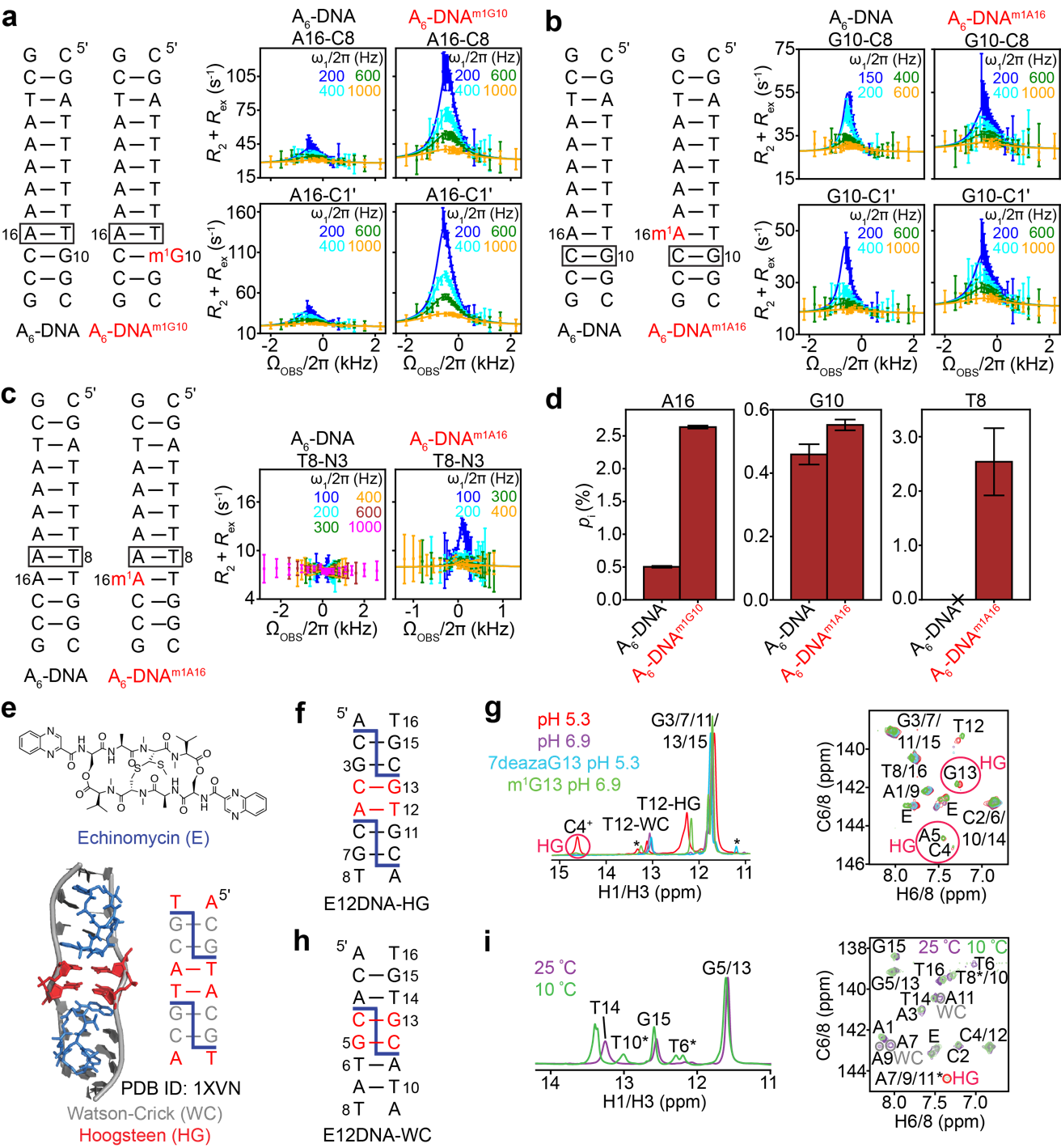

**Figure S10.** **NMR measurements probing cooperative formation of Hoogsteen bps.** Shown in panels a, b and c are the duplexes with and without preformed Hoogsteen bps induced by *N*^1^-methyl adenine/guanine (m^1^A^+^/m^1^G) (left) along with off-resonance *R*_1ρ_ relaxation dispersion profiles for the indicated nuclei (right). Sites which were probed for Hoogsteen bp formation using NMR are indicated with a box on the duplex. RF powers for *R*_1ρ_ profiles are color-coded. Two-state individual/shared fits (Table S5, Materials and Methods) of the data to the Bloch-McConnell equations is shown as solid lines. Error bars for the *R*_1ρ_ data were obtained from a Monte-Carlo based approach as described in the Materials and Methods. (d) Comparison of the population of the transient Hoogsteen bp (*p*_i_) obtained from NMR measurements in panels a, b and c, for samples with and without preformed Hoogsteen bps. Error bars for *p*_i_ were obtained using a Monte-Carlo scheme as described in Methods. ‘x’ denotes a flat *R*_1ρ_ profile from which *p*_i_ could not be obtained. (e) Chemical structure of echinomycin (top) and structure of echinomycin bound DNA (bottom, PDB ID: 1XVN). Watson-Crick and Hoogsteen bps in the echinomycin-DNA complex are shown in gray and red, respectively, while echinomycin is shown using blue sticks. Also shown is the secondary structure of echinomycin bound DNA, with echinomycin indicated using blue dashed lines. (f and h) Secondary structures of the duplexes used for NMR measurements in the presence of bound echinomycin (blue dashes). (g and i) Overlay of the imino region of the ^1^H 1D spectra (g, left) and ^1^H 1D imino SOFAST spectra (i, left), and overlay of the aromatic region of the 2D [^13^C, ^1^H] HSQC spectra (right). The E12DNA-WC echinomycin complex shows NMR signatures (red circles) of cooperative formation of tandem A11-T6/A7-T10/A9-T8 Hoogsteen bps at low temperature (10 °C), that are absent at high temperature (25 °C). * denotes tentative or unknown assignments. *R*_1ρ_ measurements in panels a, b and c were performed in a buffer containing 15 mM sodium phosphate, 25 mM sodium chloride, 0.1 mM EDTA at pH 5.4 (at 25 ºC in 100% D_2_O for panels a and b, and at 10 ºC 90% H_2_O:10% D_2_O for panel c). Echinomycin measurements were performed in a 90% H_2_O:10% D_2_O buffer containing 15 mM sodium phosphate, 125 mM sodium chloride, 0.1 mM EDTA, at the indicated pH.

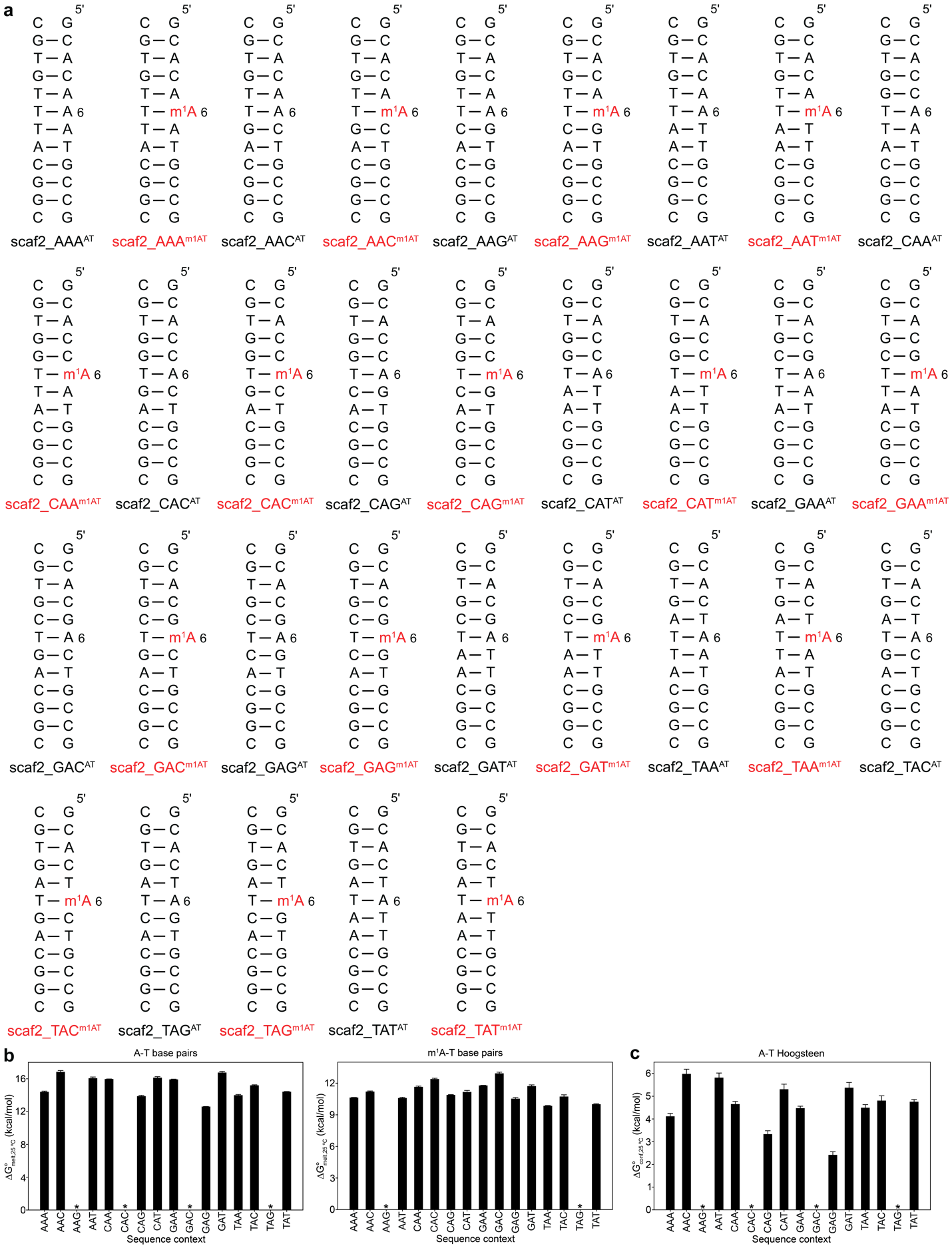

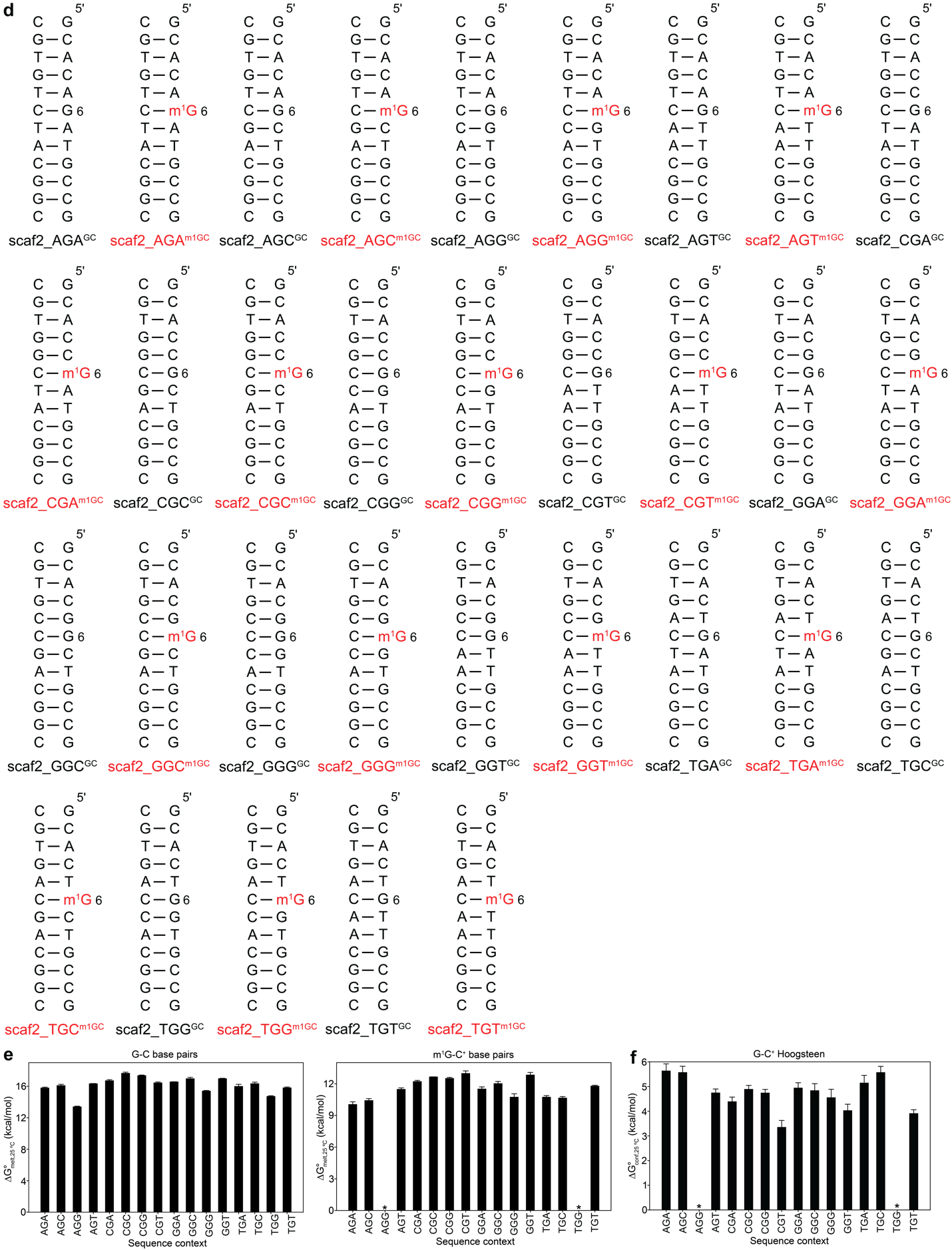

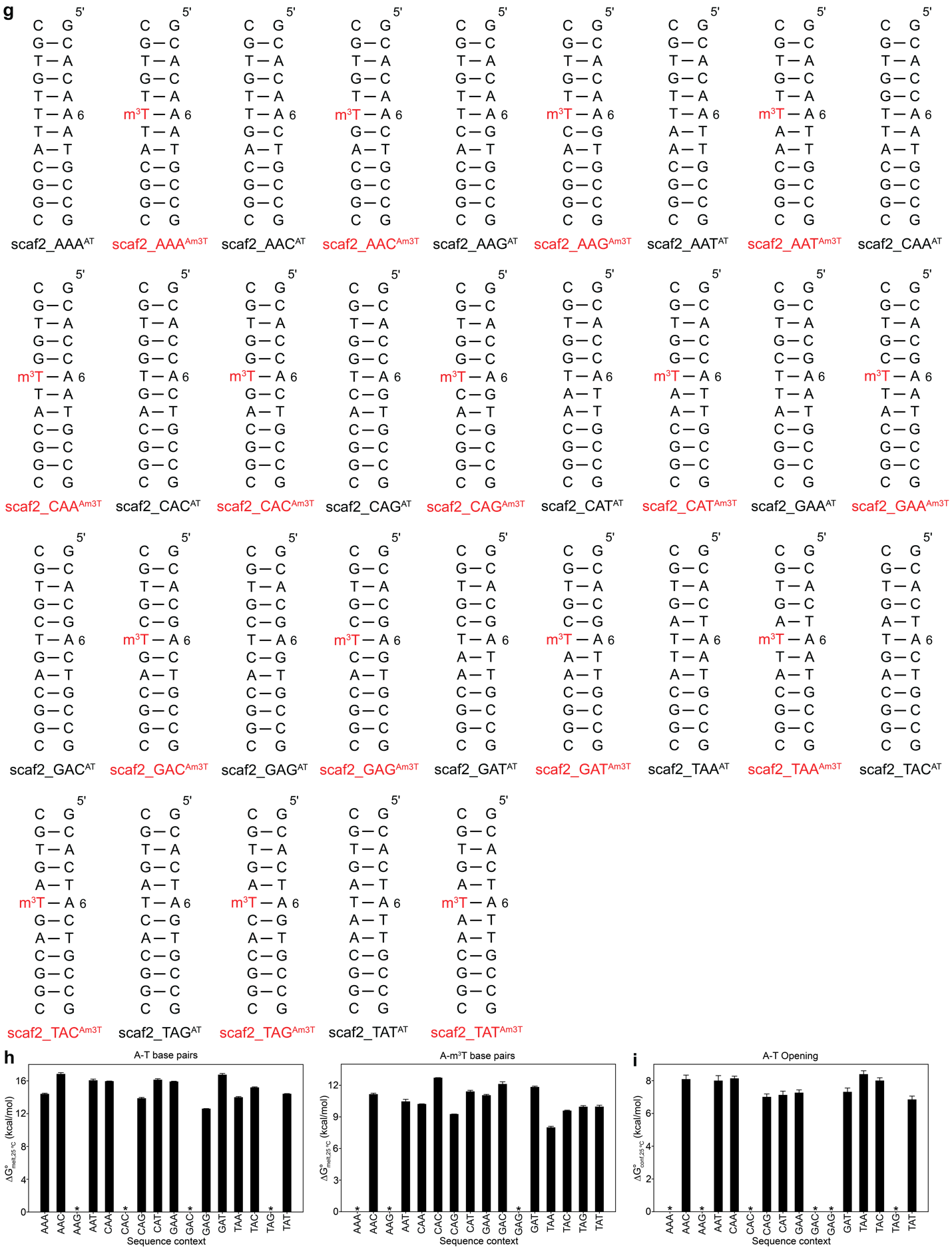

**Figure S11.** **Measuring sequence-dependent thermodynamic preferences for Hoogsteen bp formation and base opening using delta-melt.** (a, d, g) Duplexes used in the UV melting experiments for measuring sequence-dependent thermodynamic preferences. m^1^G, m^1^A and m^3^T (red) denote *N*^1^-methyl guanine, *N*^1^-methyl adenine, and *N*^3^-methyl thymine, respectively. All nucleotides are unlabeled and have deoxyribose sugars. (b, e, h) Bar plots of the energy for melting ($\Delta G_{melt,25 ^{\circ}C}^{^{\circ}}$ at 25 °C) of wild-type Watson-Crick A-T/G-C/A-T (b, e and h, left) and modified m^1^A^+^-T/m^1^G-C^+^/A-m^3^T (b, e and h, right) bp containing duplexes as a function of sequence. (c, f, i) Bar plot of the thermodynamic propensity ($\Delta\Delta G_{delta-melt,25 ^{\circ}C}^{^{\circ}}(i)$ at 25 °C) for the formation of A-T (c) and G-C^+^ (f) Hoogsteen, and A-T base opened states (i) obtained using delta-melt as a function of sequence. Buffer conditions are given in Tables S1-S2. Error bars for $\Delta G_{\mathrm{melt}}^{^{\circ}}$ and $\Delta\Delta G_{delta- melt}^{^{\circ}}(i)$ were obtained by propagating the error in UV melting measurements as described in Methods. * denotes samples for which the melts were non-two-state, preventing extraction of melting energetics (Methods, Fig. S3).

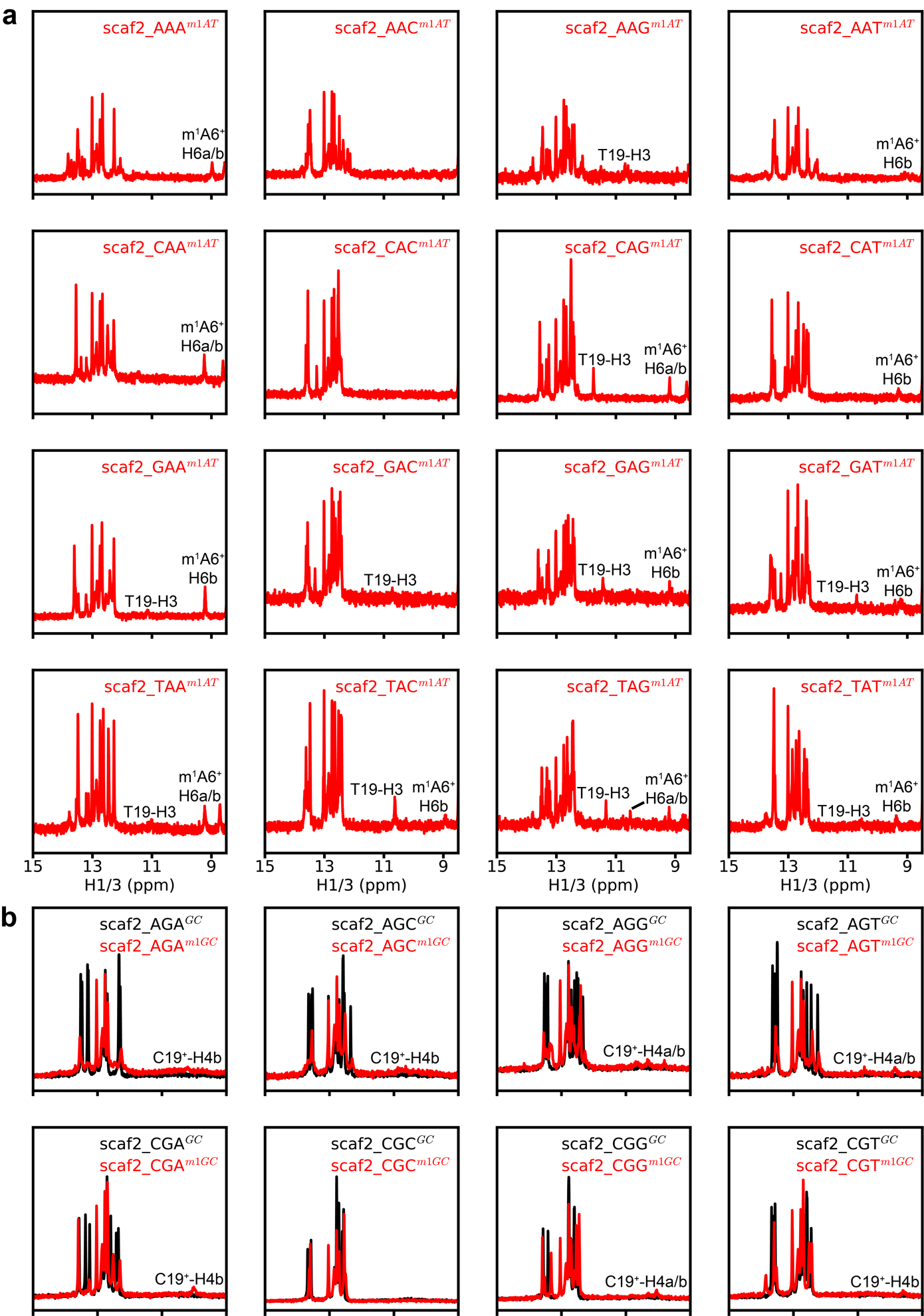

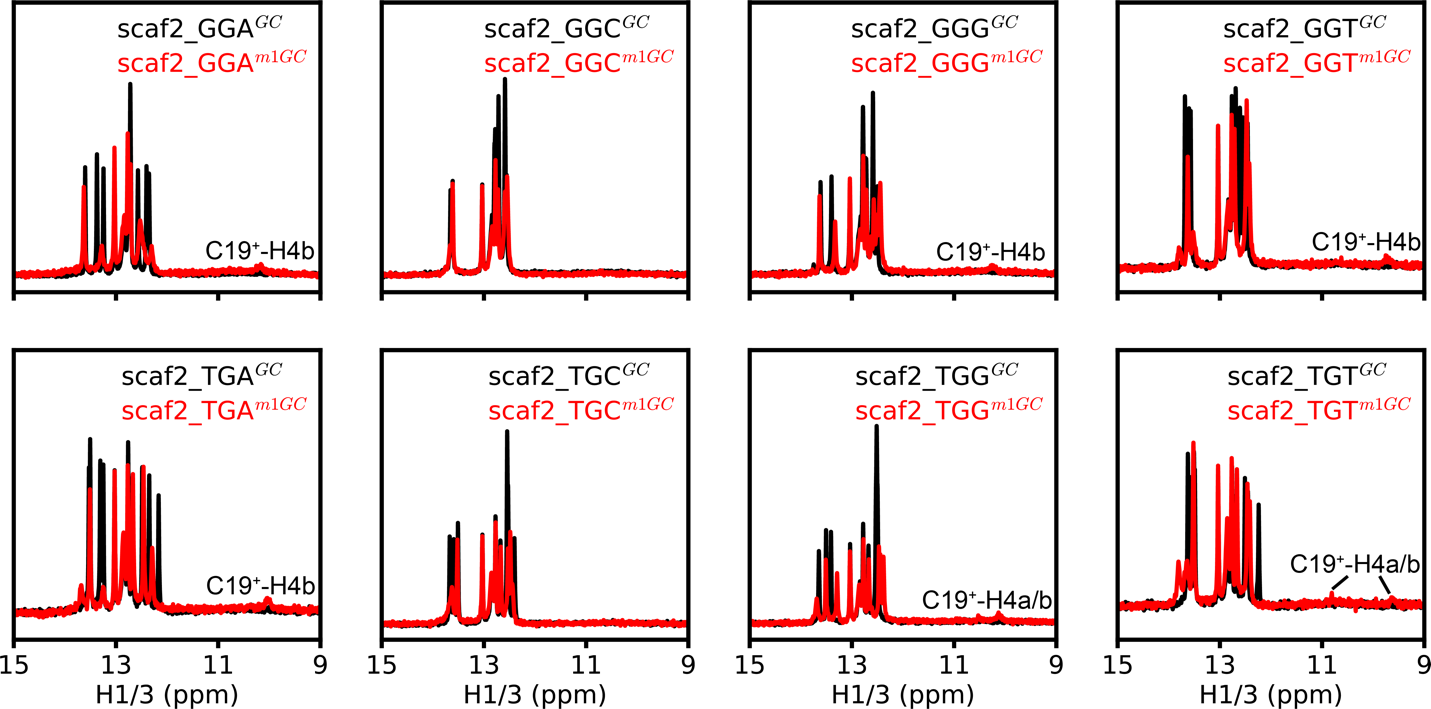

**Figure S12.** **^1^H 1D spectra of the constructs used to measure sequence-dependent thermodynamic propensities to form Hoogsteen bps.** Shown are overlays of the imino region of the ^1^H 1D spectra of the DNA duplexes used for measuring sequence-dependent thermodynamic preferences for formation of (a) A-T and (b) G-C^+^ Hoogsteen bps. Secondary structures of the constructs are given in Fig. S11d. All measurements were done at pH 5.4 in a buffer containing 15mM sodium phosphate, 150mM NaCl and 0.1mM EDTA at 10 °C. All the Watson-Crick G-C base pair containing constructs (panel b, black) display the number of imino peaks expected for duplex formation with expected secondary structures (Fig. S11d). The m^1^A^+^-T base pair containing constructs have proton signals around 9-11 ppm which correspond to T19-H3 and m^1^A6^+^-H6a/b in an m^1^A6^+^-T19 Hoogsteen bp. The m^1^G-C base pair containing constructs have proton signals around 9-11 ppm which correspond to downfield shifted amino protons for protonated cytosine, as expected for the formation of an m^1^G-C^+^ Hoogsteen base pair. The lack of Hoogsteen bp signatures for scaf2_AAA^m1AT^, scaf2_CAA^m1AT^, scaf2_AAC^m1AT^, scaf2_CAC^m1AT^, scaf2_AAT^m1AT^, scaf2_CAT^m1AT^, scaf2_CGC^m1GC^, scaf2_GGC^m1GC^ and scaf2_TGC^m1GC^ could be due to exchange broadening.

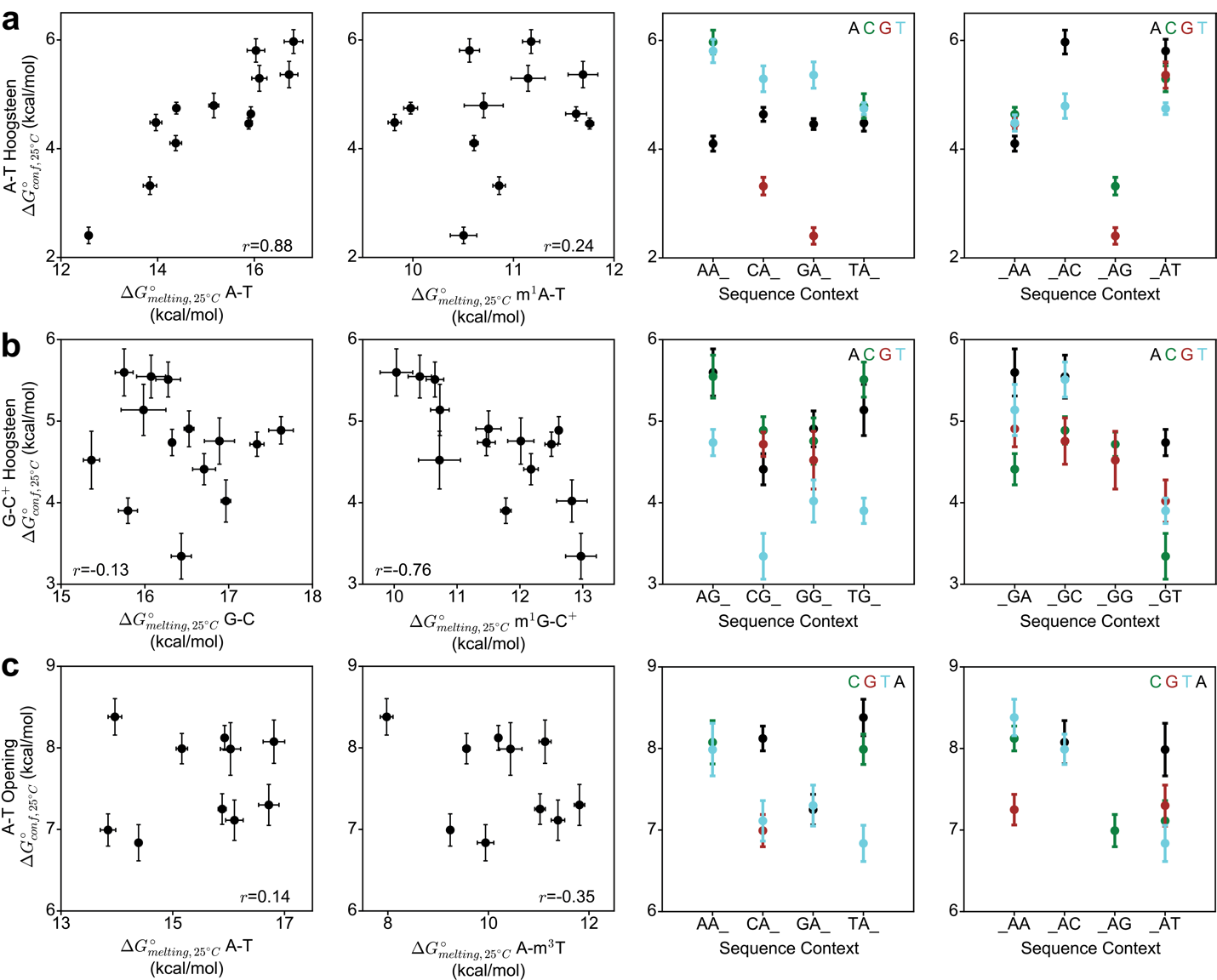

**Figure S13. Thermodynamic preferences for formation of minor conformational states from delta-melt.** Correlation plots (left) between the sequence-dependent thermodynamic preferences for A-T (a) and G-C^+^ Hoogsteen (b) bp formation, and A-T base opening (c), and the melting energetics for corresponding unmodified (A-T/G-C/A-T) and modified (m^1^A^+^-T/m^1^G-C^+^/A-m^3^T) duplexes used in delta-melt (left). The Pearson correlation coefficient (*r*) is indicated in inset. Plots (right) of the thermodynamic preferences for A-T (a) and G-C^+^ Hoogsteen (b) bp formation, and A-T base opening (c), as a function of sequence (5' to 3' direction). ‘_’ corresponds to either the A/T/C/G bases. Error bars for the conformational preferences and melting energies were obtained by propagating the error in melting energies, and by performing multiple UV melting experiments, respectively, as described in Methods.

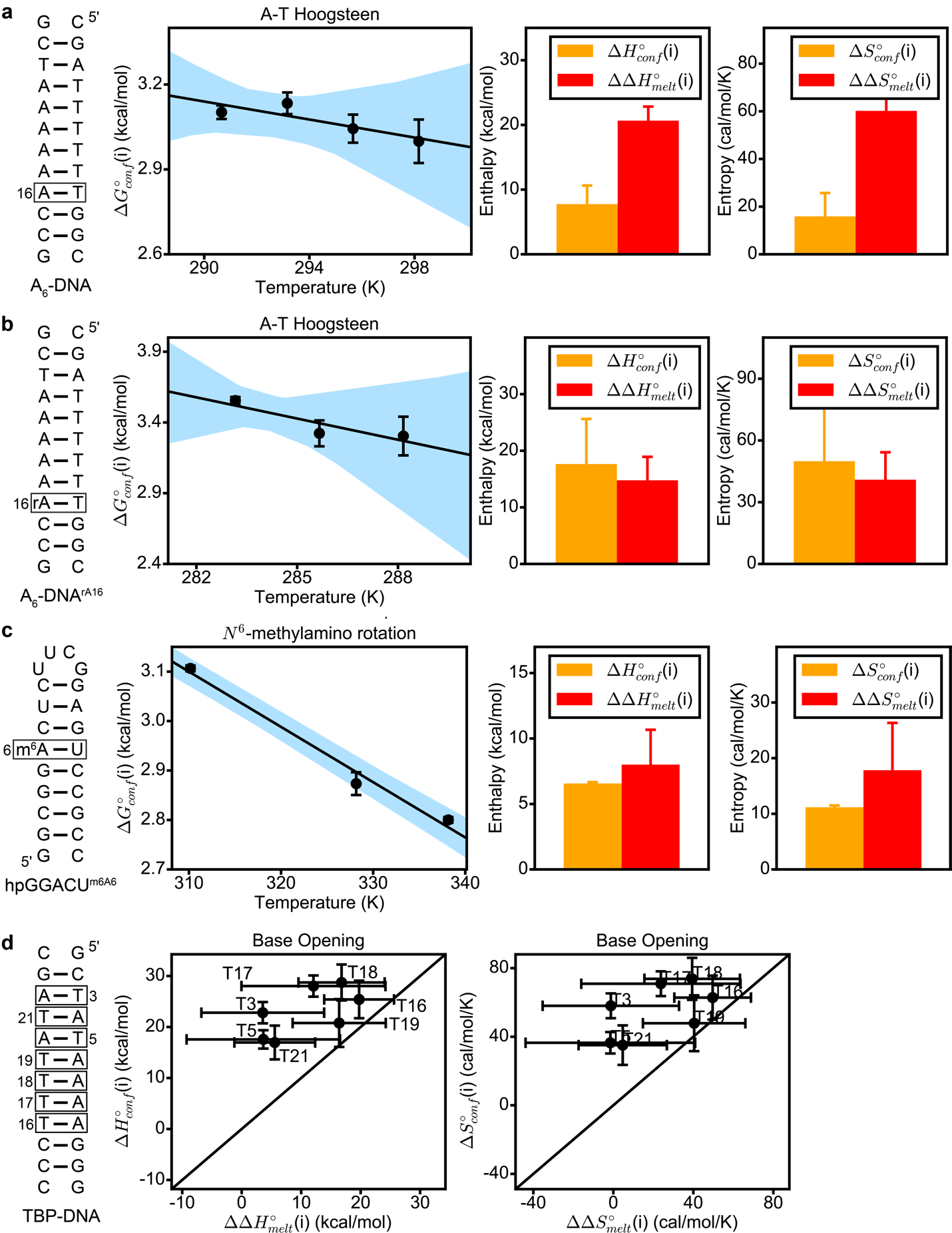

**Figure S14. Comparison of the entropy and enthalpy of conformational changes in nucleic acids from delta-melt and NMR.** Shown in panels a, b and c are the secondary structure of the construct used for NMR measurements with the position of interest enclosed by a box, temperature-dependence of the NMR derived free energies for the conformational change, and comparison of the enthalpy and entropy values obtained from NMR and that obtained from UV melting measurements (Methods). Shown in panel d is a correlation plot of the enthalpy and entropy values for base opening in TBP DNA obtained using imino proton exchange NMR measurements in Chen *et al.* (1), and UV melting measurements in this study. Black lines denotes y=x. Error bars for the thermodynamic parameters obtained from NMR and UV melting experiments were obtained as described in Methods. Blue shaded regions in panels a, b and c denote the estimate of error of linear regression obtained as described in Methods. Buffer conditions for panels a, b and c are summarized in Tables S1 and S3.

**Table S1.** Thermodynamic parameters obtained from UV melting experiments on a Perkin Elmer UV-Vis spectrophotometer

| **Construct** | **pH** | **[NaCl] (mM)** | **C_t_ (μm)** | **T_m_ (°C)** | $\boldsymbol{\Delta}\mathbf{H}_{\mathbf{melt}}^{\boldsymbol{^{\circ}}}$ **(kcal/mol)** | $\boldsymbol{\Delta}\mathbf{S}_{\mathbf{melt}}^{\boldsymbol{^{\circ}}}$ **(cal/mol/K)** | $\boldsymbol{\Delta}\mathbf{G}_{\mathbf{melt, 25}\boldsymbol{^{\circ}}\mathbf{C}}^{\boldsymbol{^{\circ}}}$ **(kcal/mol)** |
| --- | --- | --- | --- | --- | --- | --- | --- |
| **A-T Hoogsteen (m^1^A^+^-T)** | | | | | | | |
| A_6_-DNA | 4.4 | 150 | 3 | 39.7±0.3 | 85.0±0.9 | 244.9±2.6 | 12.0±0.1 |
| A_6_-DNA^m1A16^ | 4.4 | 150 | 3 | 31.9±0.1 | 71.8±1.0 | 208.6±3.3 | 9.6±0.1 |
| A_6_-DNA | 5.4 | 25 | 3 | 37.6±0.2 | 91.1±1.6 | 266.5±5.3 | 11.7±0.1 |
| A_6_-DNA^m1A16^ | 5.4 | 25 | 3 | 29.3±0.2 | 70.5±1.4 | 206.4±4.6 | 9.0±0.1 |
| A_6_-DNA^rA16^ | 5.4 | 25 | 3 | 35.5±0.6 | 89.8±1.4 | 264.2±3.9 | 11.0±0.2 |
| A_6_-DNA^m1rA16^ | 5.4 | 25 | 3 | 27.0±0.5 | 75.0±3.9 | 223.3±12.8 | 8.4±0.1 |
| A_6_-DNA | 5.4 | 150 | 3 | 46.6±0.1 | 93.2±0.2 | 264.8±0.6 | 14.3±0.1 |
| A_6_-DNA^m1A16^ | 5.4 | 150 | 3 | 38.3±0.1 | 73.5±0.7 | 209.4±2.3 | 11.1±0.1 |
| A_6_-DNA^m1G10^ | 5.4 | 150 | 3 | 35.1±0.2 | 80.4±2.4 | 234.0±7.6 | 10.6±0.1 |
| A_6_-DNA^m1G10,m1A16^ | 5.4 | 150 | 3 | 27.4±0.2 | 68.9±0.5 | 202.7±1.4 | 8.5±0.1 |
| A_2_-DNA | 6.8 | 25 | 3 | 47.0±0.1 | 102.4±1.2 | 293.1±3.9 | 15.0±0.1 |
| A_2_-DNA^m1A16^ | 6.8 | 25 | 3 | 40.0±0.2 | 81.9±1.7 | 234.8±5.3 | 11.9±0.1 |
| A_6_-DNA | 6.8 | 25 | 3 | 39.4±0.1 | 95.2±0.6 | 277.9±1.8 | 12.3±0.1 |
| A_6_-DNA^m1A16^ | 6.8 | 25 | 3 | 32.2±0.2 | 78.6±2.2 | 230.7±7.0 | 9.8±0.1 |
| A_6_-DNA^m1A21^ | 6.8 | 25 | 3 | 30.0±0.1 | 75.9±0.7 | 223.6±2.1 | 9.2±0.1 |
| **G-C^+^ Hoogsteen (m^1^G-C^+^)** | | | | | | | |
| A_2_-DNA | 4.4 | 150 | 3 | 44.2±0.4 | 76.9±2.7 | 215.6±8.6 | 12.6±0.1 |
| A_2_-DNA^m1G10^ | 4.4 | 150 | 3 | 37.3±0.2 | 70.1±1.1 | 199.1±3.6 | 10.7±0.1 |
| hpAcDNA | 5.3 | 125 | 3 | 70.1±0.1 | 83.4±1.6 | 242.9±4.6 | 10.9±0.2 |
| hpAcDNA^m1G7^ | 5.3 | 125 | 3 | 57.0±0.1 | 68.8±1.0 | 208.5±3.2 | 6.7±0.1 |
| A_2_-DNA | 5.4 | 25 | 3 | 43.8±0.3 | 90.6±1.6 | 259.2±4.8 | 13.3±0.2 |
| A_2_-DNA* | 5.4 | 25 | 3 | 43.6±0.5 | 90.0±1.5 | 257.5±4.9 | 13.2±0.2 |
| A_2_-DNA^m1G10^ | 5.4 | 25 | 3 | 35.3±0.2 | 81.1±1.8 | 236.4±6.0 | 10.7±0.1 |
| A_6_-DNA | 5.4 | 25 | 3 | 37.6±0.2 | 91.1±1.6 | 266.5±5.3 | 11.7±0.1 |
| A_6_-DNA^m1G10^ | 5.4 | 25 | 3 | 26.0±0.1 | 73.9±1.7 | 220.4±5.5 | 8.2±0.1 |
| A_6_-DNA^rG10^ | 5.4 | 25 | 3 | 36.5±0.1 | 92.8±1.8 | 273.1±5.8 | 11.4±0.1 |
| A_6_-DNA^m1rG10^ | 5.4 | 25 | 3 | 26.4±0.1 | 83.5±3.7 | 252.0±12.4 | 8.3±0.1 |
| A_6_-DNA | 5.4 | 150 | 3 | 46.6±0.1 | 93.2±0.2 | 264.8±0.6 | 14.3±0.1 |
| A_6_-DNA^m1G10^ | 5.4 | 150 | 3 | 35.1±0.2 | 80.4±2.4 | 234.0±7.6 | 10.6±0.1 |
| A_6_-DNA | 6.8 | 25 | 3 | 39.4±0.1 | 95.2±0.6 | 277.9±1.8 | 12.3±0.1 |
| A_6_-DNA^m1G10^ | 6.8 | 25 | 3 | 26.0±0.5 | 78.9±6.5 | 237.0±21.2 | 8.2±0.2 |
| **A-T Base Opening (A-m^3^T)** | | | | | | | |
| TBP-DNA | 8.0 | 100 | 3 | 38.0±0.2 | 84.0±5.6 | 243.2±18.2 | 11.5±0.2 |
| TBP-DNA^m3T3^ | 8.0 | 100 | 3 | 23.5±0.7 | 80.4±8.7 | 244.5±28.8 | 7.5±0.2 |
| TBP-DNA^m3T5^ | 8.0 | 100 | 3 | 22.8±0.9 | 80.3±11.6 | 244.7±38.3 | 7.4±0.2 |
| TBP-DNA^m3T16^ | 8.0 | 100 | 3 | 18.5±0.3 | 64.2±1.7 | 193.6±5.9 | 6.5±0.1 |
| TBP-DNA^m3T17^ | 8.0 | 100 | 3 | 18.8±1.5 | 71.9±10.7 | 219.5±35.4 | 6.5±0.2 |
| TBP-DNA^m3T18^ | 8.0 | 100 | 3 | 18.3±0.6 | 67.2±4.6 | 203.8±15.4 | 6.4±0.1 |
| TBP-DNA^m3T19^ | 8.0 | 100 | 3 | 21.3±0.7 | 67.6±5.5 | 202.8±18.0 | 7.1±0.1 |
| TBP-DNA^m3T21^ | 8.0 | 100 | 3 | 22.5±0.6 | 78.4±3.8 | 238.5±12.4 | 7.3±0.1 |
| A_6_-DNA | 8.8 | 100 | 3 | 44.9±0.4 | 94.5±1.7 | 270.6±5.4 | 13.9±0.2 |
| A_6_-DNA^m3T5^ | 8.8 | 100 | 3 | 24.6±0.6 | 74.5±2.8 | 223.4±9.2 | 7.9±0.2 |
| A_6_-DNA^m3T7^ | 8.8 | 100 | 3 | 25.3±0.6 | 80.8±6.9 | 244.2±22.8 | 8.0±0.2 |
| A_6_-DNA^m3T22^ | 8.8 | 100 | 3 | 30.6±0.1 | 81.7±3.2 | 242.4±10.3 | 9.5±0.1 |
| ***N*^6^-methylamino rotation in m^6^A (m^6^_2_A-U)** | | | | | | | |
| A_6_-RNA^m6A16^ | 6.8 | 25 | 3 | 37.0±0.1 | 94.0±0.6 | 276.4±2.1 | 11.6±0.1 |
| A_6_-RNA^m62A16^ | 6.8 | 25 | 3 | 29.4±0.3 | 80.9±3.9 | 240.6±12.7 | 9.1±0.1 |
| dsGGACU^m6A6^ | 6.8 | 25 | 3 | 51.2±0.2 | 88.2±1.2 | 245.1±3.7 | 15.1±0.1 |
| dsGGACU^m62A6^ | 6.8 | 25 | 3 | 42.5±0.2 | 80.2±2.4 | 227.3±7.6 | 12.4±0.1 |
| **Hoogsteen Cooperativity (m^1^A^+^-T/m^1^G-C^+^)** | | | | | | | |
| A_6_-DNA | 5.4 | 150 | 3 | 46.6±0.1 | 93.2±0.2 | 264.8±0.6 | 14.3±0.1 |
| A_6_-DNA^m1G10^ | 5.4 | 150 | 3 | 35.1±0.2 | 80.4±2.4 | 234.0±7.6 | 10.6±0.1 |
| A_6_-DNA^m1A16^ | 5.4 | 150 | 3 | 38.3±0.1 | 73.5±0.7 | 209.4±2.3 | 11.1±0.1 |
| A_6_-DNA^m1G10,m1A16^ | 5.4 | 150 | 3 | 27.4±0.2 | 68.9±0.5 | 202.7±1.4 | 8.5±0.1 |
| A_2_-DNA | 5.4 | 150 | 3 | 53.9±0.2 | 96.3±0.9 | 267.7±2.7 | 16.4±0.1 |
| A_2_-DNA^m1G6^ | 5.4 | 150 | 3 | 36.7±0.2 | 60.6±0.9 | 168.8±3.0 | 10.2±0.1 |
| A_2_-DNA^m1G20^ | 5.4 | 150 | 3 | 40.7±0.1 | 89.6±2.1 | 258.7±6.8 | 12.4±0.1 |
| A_2_-DNA^m1G6,20^ | 5.4 | 150 | 3 | 28.2±0.1 | 74.3±1.5 | 220.0±5.1 | 8.7±0.1 |
| A_2_-DNA | 6.8 | 150 | 3 | 55.6±0.1 | 99.8±0.6 | 277.0±1.8 | 17.2±0.1 |
| A_2_-DNA^m1A7^ | 6.8 | 150 | 3 | 45.9±0.2 | 76.3±0.9 | 212.5±2.8 | 12.9±0.1 |
| A_2_-DNA^m1A16^ | 6.8 | 150 | 3 | 48.0±0.1 | 80.4±0.5 | 223.6±1.6 | 13.7±0.1 |
| A_2_-DNA^m1A17^ | 6.8 | 150 | 3 | 46.5±0.1 | 82.2±1.9 | 230.3±5.9 | 13.5±0.1 |
| A_2_-DNA^m1A7,17^ | 6.8 | 150 | 3 | 40.2±0.1 | 71.8±0.7 | 202.4±2.4 | 11.4±0.1 |
| A_2_-DNA^m1A16,17^ | 6.8 | 150 | 3 | 38.0±0.1 | 86.4±0.6 | 251.1±1.9 | 11.6±0.1 |
| **Sequence-dependent G-C^+^ Hoogsteen (m^1^G-C^+^)** | | | | | | | |
| scaf2_CGT^GC^ | 5.4 | 25 | 3 | 48.6±0.2 | 72.4±3.7 | 198.5±11.5 | 13.2±0.3 |
| scaf2_CGT^m1GC^ | 5.4 | 25 | 3 | 38.0±0.2 | 59.0±2.2 | 163.0±7.0 | 10.4±0.1 |
| scaf2_TGC^GC^ | 5.4 | 25 | 3 | 47.1±0.2 | 68.4±1.3 | 187.1±4.3 | 12.7±0.1 |
| scaf2_TGC^m1GC^ | 5.4 | 25 | 3 | 20.6±2.0 | 27.7±3.4 | 67.5±11.0 | 7.6±0.1 |
| scaf2_AGA^GC^ | 5.4 | 150 | 3 | 55.9±0.2 | 83.4±0.7 | 226.8±1.9 | 15.8±0.1 |
| scaf2_AGA^m1GC^ | 5.4 | 150 | 3 | 37.8±0.5 | 50.7±4.6 | 136.4±14.4 | 10.0±0.3 |
| scaf2_AGC^GC^ | 5.4 | 150 | 3 | 59.1±0.2 | 79.4±1.2 | 212.3±3.5 | 16.1±0.2 |
| scaf2_AGC^m1GC^ | 5.4 | 150 | 3 | 38.5±1.6 | 57.4±2.6 | 157.5±9.2 | 10.4±0.2 |
| scaf2_AGG^GC^ | 5.4 | 150 | 3 | 57.9±0.5 | 54.8±1.6 | 139.0±4.8 | 13.4±0.1 |
| scaf2_AGT^GC^ | 5.4 | 150 | 3 | 56.5±0.1 | 87.7±0.5 | 239.4±1.5 | 16.3±0.1 |
| scaf2_AGT^m1GC^ | 5.4 | 150 | 3 | 41.8±0.1 | 66.1±2.5 | 183.3±7.9 | 11.5±0.1 |
| scaf2_CGA^GC^ | 5.4 | 150 | 3 | 59.3±0.2 | 85.0±1.4 | 229.1±4.3 | 16.7±0.1 |
| scaf2_CGA^m1GC^ | 5.4 | 150 | 3 | 45.0±0.6 | 67.8±3.4 | 186.4±10.9 | 12.2±0.1 |
| scaf2_CGC^GC^ | 5.4 | 150 | 3 | 62.7±0.1 | 86.4±1.5 | 230.4±4.6 | 17.6±0.1 |
| scaf2_CGC^m1GC^ | 5.4 | 150 | 3 | 46.7±0.1 | 69.4±0.4 | 190.2±1.3 | 12.6±0.1 |
| scaf2_CGG^GC^ | 5.4 | 150 | 3 | 62.3±0.2 | 84.6±1.0 | 225.6±3.1 | 17.4±0.1 |
| scaf2_CGG^m1GC^ | 5.4 | 150 | 3 | 54.8±0.5 | 50.2±1.6 | 126.5±5.0 | 12.5±0.1 |
| scaf2_CGT^GC^ | 5.4 | 150 | 3 | 58.9±0.3 | 83.2±1.8 | 224.1±5.8 | 16.4±1.1 |
| scaf2_CGT^m1GC^ | 5.4 | 150 | 3 | 46.9±0.1 | 73.4±3.6 | 202.7±11.3 | 13.0±0.2 |
| scaf2_GGA^GC^ | 5.4 | 150 | 3 | 58.8±0.4 | 84.7±1.1 | 228.4±3.7 | 16.6±0.1 |
| scaf2_GGA^m1GC^ | 5.4 | 150 | 3 | 43.3±0.2 | 61.4±2.8 | 167.5±8.7 | 11.5±0.2 |
| scaf2_GGC^GC^ | 5.4 | 150 | 3 | 62.2±0.1 | 81.2±1.5 | 215.6±4.4 | 17.0±0.2 |
| scaf2_GGC^m1GC^ | 5.4 | 150 | 3 | 45.7±0.7 | 62.7±2.1 | 170.0±6.4 | 12.0±0.2 |
| scaf2_GGG^GC^ | 5.4 | 150 | 3 | 61.5±0.1 | 68.3±0.7 | 177.3±2.2 | 15.4±0.1 |
| scaf2_GGG^m1GC^ | 5.4 | 150 | 3 | 54.3±1.1 | 31.0±2.6 | 67.9±7.7 | 10.7±0.3 |
| scaf2_GGT^GC^ | 5.4 | 150 | 3 | 59.6±0.3 | 86.7±0.4 | 234.0±1.5 | 17.0±0.1 |
| scaf2_GGT^m1GC^ | 5.4 | 150 | 3 | 46.0±0.1 | 74.2±3.4 | 206.0±10.7 | 12.8±0.2 |
| scaf2_TGA^GC^ | 5.4 | 150 | 3 | 55.5±0.5 | 86.6±1.8 | 236.7±5.3 | 16.0±0.3 |
| scaf2_TGA^m1GC^ | 5.4 | 150 | 3 | 39.9±0.5 | 58.4±1.3 | 159.9±3.8 | 10.7±0.1 |
| scaf2_TGC^GC^ | 5.4 | 150 | 3 | 59.1±0.2 | 81.7±1.5 | 219.4±4.5 | 16.3±0.2 |
| scaf2_TGC^m1GC^ | 5.4 | 150 | 3 | 40.7±0.2 | 54.1±2.6 | 145.7±8.2 | 10.6±0.1 |
| scaf2_TGG^GC^ | 5.4 | 150 | 3 | 58.0±0.3 | 68.0±1.4 | 178.6±4.5 | 14.7±0.1 |
| scaf2_TGT^GC^ | 5.4 | 150 | 3 | 55.7±0.1 | 84.1±1.3 | 229.0±4.1 | 15.8±0.1 |
| scaf2_TGT^m1GC^ | 5.4 | 150 | 3 | 42.1±0.5 | 70.8±3.3 | 197.8±10.9 | 11.8±0.1 |

[NaCl] denotes the concentration of sodium chloride used in the buffer, C_t_ denotes the concentration of the double stranded/hairpin species at the start of the melting measurement, T_m_ is the melting temperature,$\Delta H_{\mathrm{melt}}^{^{\circ}}$ and $\Delta S_{\mathrm{melt}}^{^{\circ}}$ denote the enthalpy and entropy of unfolding, respectively, $\Delta G_{melt,25 ^{\circ}C}^{^{\circ}}$ denotes the free energy of unfolding at 25 °C. The modification used to measure the thermodynamic preferences is indicated in parentheses. m^1^A, m^1^G, m^3^T, m^6^_2_A, m^6^A, denote *N*^1^-methyl adenine, *N*^1^-methyl guanine, *N*^3^-methyl thymine, *N*^6,6^-dimethyl adenine and *N*^6^-methyl adenine, respectively. * indicates melts conducted in 90% H_2_O:10% D_2_O solvent. Secondary structures of samples are given in Fig. 3b, 4b, 4d, 5b, 6b and 7b, and Fig. S11a, S11d and S11g. Buffer compositions for each sample are indicated under the ‘Buffer preparation’ section of the methods. Errors for thermodynamic quantities were obtained based on multiple UV melting experiments, as described in Methods. Melting parameters for scaf2_AGG^m1GC^ and scaf2_TGG^m1GC^ are not reported as these samples show deviations from 2-state melting behavior (Methods, Fig. S3).

**Table S2.** Thermodynamic parameters obtained from UV melting experiments on a Cary-100 UV-Vis spectrophotometer.

| **Construct** | **pH** | **[NaCl] (mM)** | **C_t_ (μm)** | **T_m_ (°C)** | $\boldsymbol{\Delta}\mathbf{H}_{\mathbf{melt}}^{\boldsymbol{^{\circ}}}$ **(kcal/mol)** | $\boldsymbol{\Delta}\mathbf{S}_{\mathbf{melt}}^{\boldsymbol{^{\circ}}}$ **(cal/mol/K)** | $\boldsymbol{\Delta}\mathbf{G}_{\mathbf{melt, 25}\boldsymbol{^{\circ}}\mathbf{C}}^{\boldsymbol{^{\circ}}}$ **(kcal/mol)** |
| --- | --- | --- | --- | --- | --- | --- | --- |
| **A-T Hoogsteen (m^1^A^+^-T)** | | | | | | | |
| A_6_-DNA | 4.4 | 150 | 3 | 39.9±0.3 | 80.8±1.9 | 231.5±6.0 | 11.8±0.1 |
| A_6_-DNA^m1A21^ | 4.4 | 150 | 3 | 29.1±1.0 | 56.4±1.9 | 159.9±5.9 | 8.7±0.2 |
| **A-T Hoogsteen (C-T)** | | | | | | | |
| A_6_-DNA | 4.4 | 150 | 3 | 39.9±0.3 | 80.8±1.9 | 231.5±6.0 | 11.8±0.1 |
| A_6_-DNA^C16^ | 4.4 | 150 | 3 | 27.1±0.1 | 69.0±2.1 | 203.3±7.0 | 8.4±0.1 |
| A_6_-DNA^C21^ | 4.4 | 150 | 3 | 21.0±0.3 | 54.0±0.8 | 156.6±2.9 | 7.2±0.1 |
| A_6_-DNA | 5.4 | 25 | 3 | 37.8±0.1 | 90.5±1.6 | 264.5±5.2 | 11.7±0.1 |
| A_6_-DNA^C16^ | 5.4 | 25 | 3 | 25.3±0.2 | 80.6±3.0 | 243.3±10.2 | 8.0±0.1 |
| A_6_-DNA^rA16^ | 5.4 | 25 | 3 | 38.1±0.3 | 95.6±3.0 | 280.5±9.7 | 12.0±0.1 |
| A_6_-DNA^rC16^ | 5.4 | 25 | 3 | 26.2±0.1 | 83.7±3.2 | 253.1±10.7 | 8.3±0.1 |
| A_6_-DNA | 5.4 | 150 | 3 | 47.7±0.3 | 93.5±0.6 | 264.9±1.7 | 14.5±0.1 |
| A_6_-DNA^C16^ | 5.4 | 150 | 3 | 35.0±0.1 | 88.8±2.1 | 261.6±6.8 | 10.8±0.1 |
| A_6_-DNA^m1G10^ | 5.4 | 150 | 3 | 36.0±0.1 | 85.5±0.5 | 250.1±1.6 | 11.0±0.1 |
| A_6_-DNA^m1G10,C16^ | 5.4 | 150 | 3 | 27.9±0.1 | 80.4±1.0 | 240.3±3.3 | 8.7±0.1 |
| A_2_-DNA | 6.8 | 25 | 3 | 48.0±0.2 | 96.3±0.8 | 273.4±2.5 | 14.8±0.1 |
| A_2_-DNA^C16^ | 6.8 | 25 | 3 | 34.3±0.1 | 80.8±0.5 | 236.3±1.6 | 10.4±0.1 |
| A_6_-DNA | 6.8 | 25 | 3 | 40.0±0.2 | 91.2±1.7 | 264.4±5.5 | 12.3±0.1 |
| A_6_-DNA^C16^ | 6.8 | 25 | 3 | 27.1±0.1 | 86.2±1.9 | 260.4±6.3 | 8.6±0.1 |
| A_6_-DNA^C21^ | 6.8 | 25 | 3 | 24.2±0.1 | 78.9±0.7 | 238.6±2.2 | 7.7±0.1 |
| **A-T Hoogsteen (T-T)** | | | | | | | |
| A_6_-DNA | 4.4 | 150 | 3 | 39.9±0.3 | 80.8±1.9 | 231.5±6.0 | 11.8±0.1 |
| A_6_-DNA^T16^ | 4.4 | 150 | 3 | 27.9±0.1 | 71.9±0.5 | 212.1±1.6 | 8.6±0.1 |
| A_6_-DNA^T21^ | 4.4 | 150 | 3 | 24.7±0.4 | 64.4±1.9 | 189.6±6.2 | 7.9±0.1 |
| A_6_-DNA | 5.4 | 25 | 3 | 37.8±0.1 | 90.5±1.6 | 264.5±5.2 | 11.7±0.1 |
| A_6_-DNA^T16^ | 5.4 | 25 | 3 | 26.6±0.1 | 78.8±1.5 | 236.4±4.9 | 8.4±0.1 |
| A_6_-DNA^rA16^ | 5.4 | 25 | 3 | 38.1±0.3 | 95.6±3.0 | 280.5±9.7 | 12.0±0.1 |
| A_6_-DNA^rT16^ | 5.4 | 25 | 3 | 27.3±0.1 | 85.7±1.5 | 258.5±5.2 | 8.6±0.1 |
| A_6_-DNA | 5.4 | 150 | 3 | 47.7±0.3 | 93.5±0.6 | 264.9±1.7 | 14.5±0.1 |
| A_6_-DNA^T16^ | 5.4 | 150 | 3 | 36.7±0.1 | 84.5±2.5 | 246.0±7.8 | 11.1±0.1 |
| A_6_-DNA^m1G10^ | 5.4 | 150 | 3 | 36.0±0.1 | 85.5±0.5 | 250.1±1.6 | 11.0±0.1 |
| A_6_-DNA^m1G10,T16^ | 5.4 | 150 | 3 | 29.2±0.2 | 84.3±2.0 | 252.0±6.4 | 9.1±0.1 |
| A_2_-DNA | 6.8 | 25 | 3 | 48.0±0.2 | 96.3±0.8 | 273.4±2.5 | 14.8±0.1 |
| A_2_-DNA^T16^ | 6.8 | 25 | 3 | 37.1±0.2 | 86.6±0.9 | 252.3±3.1 | 11.3±0.1 |
| A_6_-DNA | 6.8 | 25 | 3 | 40.0±0.2 | 91.2±1.7 | 264.4±5.5 | 12.3±0.1 |
| A_6_-DNA^T16^ | 6.8 | 25 | 3 | 29.2±0.2 | 86.2±0.8 | 258.3±2.6 | 9.1±0.1 |
| A_6_-DNA^T21^ | 6.8 | 25 | 3 | 26.7±0.1 | 80.4±1.7 | 241.5±5.6 | 8.4±0.1 |
| **A-T Opening (A-m^3^T)** | | | | | | | |
| A_6_-DNA | 8.8 | 100 | 3 | 44.4±0.3 | 91.7±4.6 | 262.2±14.3 | 13.6±0.4 |
| A_6_-DNA^m3T6^ | 8.8 | 100 | 3 | 25.3±0.3 | 79.6±6.0 | 240.0±20.3 | 8.0±0.1 |
| **Sequence-dependent A-T Hoogsteen (m^1^A^+^-T)** | | | | | | | |
| scaf2_AAA^AT^ | 8.8 | 100 | 3 | 51.2±0.3 | 79.6±1.2 | 218.7±3.6 | 14.4±0.1 |
| scaf2_AAA^m1AT^ | 8.8 | 100 | 3 | 41.6±0.2 | 50.6±0.5 | 134.0±1.7 | 10.6±0.1 |
| scaf2_AAC^AT^ | 8.8 | 100 | 3 | 55.8±0.3 | 94.6±2.0 | 261.1±6.3 | 16.8±0.2 |
| scaf2_AAC^m1AT^ | 8.8 | 100 | 3 | 44.8±0.1 | 51.8±1.2 | 136.1±3.6 | 11.2±0.1 |
| scaf2_AAT^AT^ | 8.8 | 100 | 3 | 52.2±0.2 | 96.8±1.6 | 270.8±4.9 | 16.0±0.2 |
| scaf2_AAT^m1AT^ | 8.8 | 100 | 3 | 41.7±0.6 | 49.3±0.9 | 130.1±2.9 | 10.6±0.1 |
| scaf2_CAA^AT^ | 8.8 | 100 | 3 | 54.8±0.1 | 88.0±0.3 | 241.4±0.9 | 15.9±0.1 |
| scaf2_CAA^m1AT^ | 8.8 | 100 | 3 | 46.1±0.3 | 55.7±0.8 | 147.7±2.3 | 11.6±0.1 |
| scaf2_CAC^m1AT^ | 8.8 | 100 | 3 | 49.9±0.1 | 57.3±1.5 | 150.6±4.6 | 12.4±0.1 |
| scaf2_CAG^AT^ | 8.8 | 100 | 3 | 55.4±0.1 | 63.7±1.4 | 167.1±4.1 | 13.8±0.1 |
| scaf2_CAG^m1AT^ | 8.8 | 100 | 3 | 49.4±0.5 | 38.5±1.5 | 92.6±4.9 | 10.9±0.1 |
| scaf2_CAT^AT^ | 8.8 | 100 | 3 | 54.2±0.5 | 91.5±2.6 | 252.7±8.3 | 16.1±0.2 |
| scaf2_CAT^m1AT^ | 8.8 | 100 | 3 | 45.2±0.9 | 50.4±2.0 | 131.6±6.3 | 11.1±0.2 |
| scaf2_GAA^AT^ | 8.8 | 100 | 3 | 55.3±0.3 | 86.0±0.4 | 235.3±1.4 | 15.9±0.1 |
| scaf2_GAA^m1AT^ | 8.8 | 100 | 3 | 46.7±0.1 | 56.2±0.5 | 149.1±1.6 | 11.8±0.1 |
| scaf2_GAC^m1AT^ | 8.8 | 100 | 3 | 51.5±0.3 | 60.8±1.1 | 160.8±3.3 | 12.9±0.1 |
| scaf2_GAG^AT^ | 8.8 | 100 | 3 | 56.2±0.3 | 48.9±0.4 | 121.9±1.4 | 12.6±0.1 |
| scaf2_GAG^m1AT^ | 8.8 | 100 | 3 | 52.6±0.8 | 30.2±0.9 | 66.0±2.7 | 10.5±0.1 |
| scaf2_GAT^AT^ | 8.8 | 100 | 3 | 55.6±0.3 | 94.2±1.8 | 259.8±5.6 | 16.7±0.2 |
| scaf2_GAT^m1AT^ | 8.8 | 100 | 3 | 47.0±0.4 | 54.6±1.6 | 143.8±5.0 | 11.7±0.1 |
| scaf2_TAA^AT^ | 8.8 | 100 | 3 | 49.8±0.3 | 78.4±1.3 | 216.2±3.9 | 14.0±0.1 |
| scaf2_TAA^m1AT^ | 8.8 | 100 | 3 | 38.8±0.2 | 42.3±1.1 | 108.9±3.4 | 9.8±0.1 |
| scaf2_TAC^AT^ | 8.8 | 100 | 3 | 53.0±0.3 | 84.2±0.5 | 231.4±1.3 | 15.2±0.1 |
| scaf2_TAC^m1AT^ | 8.8 | 100 | 3 | 44.0±0.6 | 46.0±2.2 | 118.4±6.9 | 10.7±0.2 |
| scaf2_TAT^AT^ | 8.8 | 100 | 3 | 50.0±0.1 | 83.4±1.2 | 231.5±3.8 | 14.4±0.1 |
| scaf2_TAT^m1AT^ | 8.8 | 100 | 3 | 39.6±0.3 | 43.5±0.8 | 112.3±2.3 | 10.0±0.1 |
| **Sequence-dependent A-T opening (A-m^3^T)** | | | | | | | |
| scaf2_AAA^AT^ | 8.8 | 100 | 3 | 51.2±0.3 | 79.6±1.2 | 218.7±3.6 | 14.4±0.1 |
| scaf2_AAC^AT^ | 8.8 | 100 | 3 | 55.8±0.3 | 94.6±2.0 | 261.1±6.3 | 16.8±0.2 |
| scaf2_AAC^Am3T^ | 8.8 | 100 | 3 | 40.0±0.1 | 66.5±2.2 | 185.8±7.0 | 11.1±0.1 |
| scaf2_AAT^AT^ | 8.8 | 100 | 3 | 52.2±0.2 | 96.8±1.6 | 270.8±4.9 | 16.0±0.2 |
| scaf2_AAT^Am3T^ | 8.8 | 100 | 3 | 36.7±0.7 | 66.1±5.0 | 186.8±16.1 | 10.4±0.2 |
| scaf2_CAA^AT^ | 8.8 | 100 | 3 | 54.8±0.1 | 88.0±0.3 | 241.4±0.9 | 15.9±0.1 |
| scaf2_CAA^Am3T^ | 8.8 | 100 | 3 | 39.3±0.2 | 49.3±0.9 | 131.2±2.9 | 10.2±0.1 |
| scaf2_CAC^Am3T^ | 8.8 | 100 | 3 | 44.8±0.2 | 76.1±0.3 | 212.8±0.9 | 12.7±0.1 |
| scaf2_CAG^AT^ | 8.8 | 100 | 3 | 55.4±0.1 | 63.7±1.4 | 167.1±4.1 | 13.8±0.1 |
| scaf2_CAG^Am3T^ | 8.8 | 100 | 3 | 44.9±0.4 | 20.7±0.3 | 38.5±1.1 | 9.2±0.1 |
| scaf2_CAT^AT^ | 8.8 | 100 | 3 | 54.2±0.5 | 91.5±2.6 | 252.7±8.3 | 16.1±0.2 |
| scaf2_CAT^Am3T^ | 8.8 | 100 | 3 | 40.2±0.4 | 71.1±2.4 | 200.3±7.7 | 11.4±0.1 |
| scaf2_GAA^AT^ | 8.8 | 100 | 3 | 55.3±0.3 | 86.0±0.4 | 235.3±1.4 | 15.9±0.1 |
| scaf2_GAA^Am3T^ | 8.8 | 100 | 3 | 42.6±0.2 | 55.4±1.4 | 148.8±4.2 | 11.0±0.1 |
| scaf2_GAC^Am3T^ | 8.8 | 100 | 3 | 45.0±0.6 | 65.8±2.1 | 180.1±6.1 | 12.1±0.2 |
| scaf2_GAG^AT^ | 8.8 | 100 | 3 | 56.2±0.3 | 48.9±0.4 | 121.9±1.4 | 12.6±0.1 |
| scaf2_GAT^AT^ | 8.8 | 100 | 3 | 55.6±0.3 | 94.2±1.8 | 259.8±5.6 | 16.7±0.2 |
| scaf2_GAT^Am3T^ | 8.8 | 100 | 3 | 41.2±0.6 | 75.1±3.7 | 212.2±12.2 | 11.8±0.1 |
| scaf2_TAA^AT^ | 8.8 | 100 | 3 | 49.8±0.3 | 78.4±1.3 | 216.2±3.9 | 14.0±0.1 |
| scaf2_TAA^Am3T^ | 8.8 | 100 | 3 | 25.2±1.3 | 28.6±2.3 | 69.1±7.5 | 8.0±0.1 |
| scaf2_TAC^AT^ | 8.8 | 100 | 3 | 53.0±0.3 | 84.2±0.5 | 231.4±1.3 | 15.2±0.1 |
| scaf2_TAC^Am3T^ | 8.8 | 100 | 3 | 35.3±0.5 | 48.5±0.8 | 130.5±2.6 | 9.6±0.1 |
| scaf2_TAG^Am3T^ | 8.8 | 100 | 3 | 46.0±0.9 | 30.2±0.9 | 68.1±2.5 | 9.9±0.1 |
| scaf2_TAT^AT^ | 8.8 | 100 | 3 | 50.0±0.1 | 83.4±1.2 | 231.5±3.8 | 14.4±0.1 |
| scaf2_TAT^Am3T^ | 8.8 | 100 | 3 | 34.3±0.4 | 65.8±4.9 | 187.5±15.9 | 9.9±0.2 |

[NaCl] denotes the concentration of sodium and magnesium chloride used in the buffers, respectively, C_t_ denotes the concentration of the double stranded/hairpin species at the start of the melting measurement, T_m_ is the melting temperature,$\Delta H_{\mathrm{melt}}^{^{\circ}}$ and $\Delta S_{\mathrm{melt}}^{^{\circ}}$ denote the enthalpy and entropy of unfolding, respectively, $\Delta G_{melt,25 ^{\circ}C}^{^{\circ}}$ denotes the free energy of unfolding at 25 °C and m^6^A denotes *N*^6^-methyl adenine. The modification used to measure the thermodynamic preferences is indicated in parentheses. m^1^A, rA, rC, rT, m^1^G, m^3^T, m^6^_2_A, m^6^A, denote *N*^1^-methyl adenine, *N*^1^-methyl guanine, ribo adenine, ribo cytosine, ribo thymidine, *N*^3^-methyl thymine, *N*^6,6^-dimethyl adenine and *N*^6^-methyl adenine, respectively. * indicates melts conducted in 90% H_2_O:10% D_2_O solvent. Secondary structures of samples are given in Fig. 3b, 4b, 4d, 5b, 6b and 7b, and Fig. S11a, S11d and S11g. Buffer compositions for each sample are indicated under the ‘Buffer preparation’ section of the methods. Errors for thermodynamic quantities were obtained based on multiple UV melting experiments, as described in Methods. The melting parameters for scaf2_AAA^Am3T^, scaf2_AAG^AT^, scaf2_AAG^m1AT^, scaf2_AAG^Am3T^, scaf2_CAC^AT^, scaf2_GAC^AT^, scaf2_GAG^Am3T^, scaf2_TAG^AT^ and scaf2_TAG^m1AT^ are not reported as they show deviations from two-state melting behavior (Methods, Fig. S3).

**Table S3.** Summary of the experimental conditions and labeling schemes used for collecting the *R*_1ρ_ and CEST data in this study

| **Construct** | **pH** | **Temp**  **(°C)** | **[NaCl]**  **(mM)** | **H_2_O:**  **D_2_O** | **Nucleus** | **Spectrom-eter**  **(MHz)** | **Expt.** | **Labeling** |
| --- | --- | --- | --- | --- | --- | --- | --- | --- |
| **A-T Hoogsteen** | | | | | | | | |
| A_6_-DNA | 4.4 | 25 | 150 | 90:10 | A16-C8 | 700 | *R*_1ρ_ | A_6_-DNA(s1lb) |
| A_6_-DNA | 4.4 | 30 | 150 | 90:10 | A21-C1' | 600 | *R*_1ρ_ | A_6_-DNA(ulb) |
| A_6_-DNA | 5.4 | 17.5 | 25 | 90:10 | A16-C8 | 600 | *R*_1ρ_ | A_6_-DNA(A16lb) |
| A_6_-DNA | 5.4 | 20 | 25 | 90:10 | A16-C8 | 600 | *R*_1ρ_ | A_6_-DNA(A16lb) |
| A_6_-DNA | 5.4 | 22.5 | 25 | 90:10 | A16-C8 | 600 | *R*_1ρ_ | A_6_-DNA(A16lb) |
| A_6_-DNA | 5.4 | 25 | 25 | 90:10 | A16-C8 | 600 | *R*_1ρ_ | A_6_-DNA(A16lb) |
| A_6_-DNA^rA16^ | 5.4 | 10 | 25 | 90:10 | rA16-C8 | 600 | *R*_1ρ_ | A_6_-DNA^rA16^  (rA16^C8^,C15lb) |
| A_6_-DNA^rA16^ | 5.4 | 12.5 | 25 | 90:10 | rA16-C8 | 600 | *R*_1ρ_ | A_6_-DNA^rA16^ (rA16^C8^,C15lb) |
| A_6_-DNA^rA16^ | 5.4 | 15 | 25 | 90:10 | rA16-C8 | 600 | *R*_1ρ_ | A_6_-DNA^rA16^ (rA16^C8^,C15lb) |
| A_6_-DNA | 5.4 | 25 | 150 | 90:10 | A16-C8,C1' | 600 | *R*_1ρ_ | A_6_-DNA(A16lb) |
| A_6_-DNA^m1G10^ | 5.4 | 25 | 150 | 90:10 | A16-C8,C1' | 600 | *R*_1ρ_ | A_6_-DNA^m1G10^(s1lb) |
| A_2_-DNA | 6.8 | 25 | 25 | 90:10 | A16-C8 | 600 | *R*_1ρ_ | A_2_-DNA(ulb) |
| A_6_-DNA | 6.8 | 25 | 25 | 90:10 | A16-C8 | 700 | *R*_1ρ_ | A_6_-DNA(ulb) |
| A_6_-DNA | 6.8 | 25 | 25 | 90:10 | A21-C1' | 600 | *R*_1ρ_ | A_6_-DNA(ulb) |
| A_6_-DNA | 6.8 | 30 | 25 | 90:10 | A21-C1' | 600 | *R*_1ρ_ | A_6_-DNA(ulb) |
| **G-C^+^ Hoogsteen** | | | | | | | | |
| A_2_-DNA | 4.4 | 25 | 150 | 90:10 | G10-C8 | 700 | *R*_1ρ_ | A_2_-DNA(s2lb) |
| AcDNA | 5.3 | 25 | 125 | 90:10 | G7-C1',C8,N1 | 700 | *R*_1ρ_ | AcDNA(ulb) |
| A_2_-DNA | 5.4 | 25 | 25 | 90:10 | G10-C8,C1' | 700 | *R*_1ρ_ | A_2_-DNA(s2lb) |
| A_6_-DNA | 5.4 | 25 | 25 | 90:10 | G10-C8,C1' | 600 | *R*_1ρ_ | A_6_-DNA(s2lb) |
| A_6_-DNA^rG10^ | 5.4 | 25 | 25 | 90:10 | C15-C6 | 700 | *R*_1ρ_ | A_6_-DNA^rG10^ (s1lb) |
| A_6_-DNA | 5.4 | 25 | 150 | 90:10 | G10-C8,C1' | 600 | *R*_1ρ_ | A_6_-DNA(s2lb) |
| A_6_-DNA | 6.8 | 25 | 25 | 90:10 | G10-C1' | 700 | *R*_1ρ_ | A_6_-DNA(ulb) |
| ***N*^6^-methylamino rotation in m^6^A** | | | | | | | | |
| hpGGACU^m6A6^ | 6.8 | 37 | 25 | 90:10 | m^6^A6-C2 | 600 | CEST | hpGGACU^m6A6^ (m^6^A6^C2,C8^lb) |
| hpGGACU^m6A6^ | 6.8 | 37 | 25 | 90:10 | m^6^A6-C10 | 600 | CEST | hpGGACU^m6A6^  (m^6^A6^C10^lb) |
| hpGGACU^m6A6^ | 6.8 | 55 | 25 | 90:10 | m^6^A6-C2 | 600 | *R*_1ρ_ | hpGGACU^m6A6^  (m^6^A6^C2,C8^lb) |
| hpGGACU^m6A6^ | 6.8 | 65 | 25 | 90:10 | m^6^A6-C2 | 600 | *R*_1ρ_ | hpGGACU^m6A6^  (m^6^A6^C2,C8^lb) |
| A_6_-RNA^m6A16^ | 6.8 | 37 | 25 | 90:10 | m^6^A16-C2 | 600 | *R*_1ρ_ | A_6_-RNA^m6A16^  (m^6^A16^C2,C8^lb) |
| **Hoogsteen Cooperativity** | | | | | | | | |
| A_6_-DNA | 5.4 | 25 | 25 | 0:100 | A16-C8,C1' | 700 | *R*_1ρ_ | A_6_-DNA(s1lb) |
| A_6_-DNA | 5.4 | 25 | 25 | 0:100 | G10-C8,C1' | 700 | *R*_1ρ_ | A_6_-DNA(s2lb) |
| A_6_-DNA^m1A16^ | 5.4 | 25 | 25 | 0:100 | G10-C8,C1' | 700 | *R*_1ρ_ | A_6_-DNA^m1A16^(s2lb) |
| A_6_-DNA^m1G10^ | 5.4 | 25 | 25 | 0:100 | A16-C8,C1' | 700 | *R*_1ρ_ | A_6_-DNA^m1G10^(s1lb) |
| A_6_-DNA | 5.4 | 10 | 25 | 90:10 | T8-N3 | 700 | *R*_1ρ_ | A_6_-DNA(s2lb) |
| A_6_-DNA^m1A16^ | 5.4 | 10 | 25 | 90:10 | T8-N3 | 700 | *R*_1ρ_ | A_6_-DNA^m1A16^(s2lb) |
| **G-C^+^ Hoogsteen: Testing delta-melt** | | | | | | | | |
| Scaf2_  CGT^GC^ | 5.4 | 30 | 25 | 90:10 | G6-C8,C1' | 600 | *R*_1ρ_ | Scaf2_CGT^GC^  (G6lb) |
| Scaf2_  TGC^GC^ | 5.4 | 30 | 25 | 90:10 | G6-C8,C1' | 600 | *R*_1ρ_ | Scaf2_TGC^GC^  (G6lb) |
| Scaf2_  TGC^GC^ | 5.4 | 15 | 150 | 90:10 | G6-C8 | 600 | *R*_1ρ_ | Scaf2_TGC^GC^  (G6lb) |
| Scaf2_  TGC^GC^ | 5.4 | 30 | 150 | 90:10 | G6-C8,C1' | 600 | *R*_1ρ_ | Scaf2_TGC^GC^  (G6lb) |
| Scaf2_  TGC^GC^ | 5.4 | 40 | 150 | 90:10 | G6-C8,C1' | 600 | *R*_1ρ_ | Scaf2_TGC^GC^  (G6lb) |
| Scaf2_  CGT^GC^ | 5.4 | 30 | 150 | 90:10 | G6-C8,C1' | 600 | CEST | Scaf2_CGT^GC^  (G6lb) |

[NaCl] denotes the concentration of sodium chloride used in the buffers, respectively, H_2_O:D_2_O denotes the ratio of H_2_O and D_2_O in the buffer. rA, m^1^G, rG, m^6^A, m^1^A denote ribo adenosine, *N*^1^-methyl guanine, ribo guanine, *N*^6^-methyl adenine, *N*^1^-methyl adenosine, respectively. Secondary structures of the constructs are given in Fig. S4.

**Table S4.** List of spin-lock powers (ω_1_/2π, in Hz) and offsets (Ω/2π, in Hz) used in off-resonance ^13^C/^15^N *R*_1ρ_ and CEST experiments

| **Nucleus** | **[ω_1_/2π (Hz)] {Ω/2π (Hz)}** |
| --- | --- |
| **A-T Hoogsteen** | |
| A_6_-DNA  A16-C8  pH 4.4HS  25 °C  (*R*_1ρ_) | [150, 200, 250, 300, 400, 500, 600, 700, 900, 1000, 1200, 1400, 1600, 2000, 2500] {0}  [200], {-600.0, -400.0, -300.0, -200.0, -150.0, -100.0, -50.0, 50.0, 100.0, 150.0, 200.0, 200.0, 250.0, 300.0, 350.0, 400.0, 450.0, 500.0, 550.0, 600.0}  [400], {-1200.0, -1000.0, -850.0, -700.0, -550.0, -400.0, -250.0, -100.0, -50.0, 50.0, 100.0, 150.0, 200.0, 200.0, 250.0, 300.0, 350.0, 400.0, 450.0, 500.0, 650.0, 800.0}  [600], {-1600.0, -1200.0, -800.0, -600.0, -400.0, -200.0, -100.0, 50.0, 100.0, 150.0, 200.0, 200.0, 250.0, 300.0, 350.0, 400.0, 500.0, 600.0, 800.0, 1000.0, 1200.0, 1600.0}  [1000], {-1600.0, -1000.0, -700.0, -400.0, -100.0, 50.0, 150.0, 200.0, 200.0, 250.0, 350.0, 500.0, 800.0, 1100.0, 1400.0, 2000.0} |
| A_6_-DNA  A21-C1'  pH 4.4HS  30 °C  (*R*_1ρ_) | [150], {-500.0, -450.0, -400.0, -350.0, -300.0, -250.0, -200.0, -150.0, -100.0, -60.0, -30.0, 30.0, 60.0, 100.0, 150.0, 200.0, 250.0, 300.0, 350.0, 400.0, 450.0, 500.0}  [200], {-600.0, -500.0, -400.0, -350.0, -300.0, -250.0, -200.0, -150.0, -100.0, -60.0, -30.0, 30.0, 60.0, 100.0, 150.0, 200.0, 250.0, 300.0, 350.0, 400.0, 500.0, 600.0}  [400], {-1200.0, -1000.0, -800.0, -600.0, -500.0, -400.0, -300.0, -200.0, -150.0, -100.0, -50.0, 50.0, 100.0, 150.0, 200.0, 300.0, 400.0, 500.0, 600.0, 800.0, 1000.0, 1200.0}  [1000], {-3500.0, -3000.0, -2400.0, -1800.0, -1200.0, -900.0, -600.0, -300.0, -150.0, -50.0, 50.0, 150.0, 300.0, 600.0, 900.0, 1200.0, 1800.0, 2400.0, 3000.0, 3500.0}  [150, 200, 250, 300, 400, 500, 600, 700, 900, 1000, 1200, 1400, 1600, 2000, 2500, 3000] {0} |
| A_6_-DNA  A16-C8  pH 5.4  17.5 °C  (*R*_1ρ_) | [100, 150, 200, 250, 300, 400, 500, 600, 700, 900, 1000, 1200, 1600, 2000, 2500, 3000] {0}  [100], {-375.0, -225.0, -125.0, -25.0, 75.0, 125.0, 175.0, 225.0, 275.0, 295.0, 315.0, 335.0, 355.0, 375.0, 395.0, 435.0, 455.0, 475.0, 525.0, 575.0}  [150], {-625.0, -375.0, -225.0, -125.0, -25.0, 75.0, 125.0, 175.0, 225.0, 275.0, 295.0, 315.0, 335.0, 355.0, 375.0, 395.0, 415.0, 435.0, 455.0, 475.0, 525.0, 575.0, 675.0, 775.0, 875.0}  [200], {-625.0, -425.0, -225.0, -125.0, -25.0, 75.0, 125.0, 175.0, 225.0, 275.0, 295.0, 315.0, 335.0, 355.0, 375.0, 395.0, 415.0, 435.0, 455.0, 475.0, 525.0, 575.0, 625.0, 675.0, 775.0, 875.0, 975.0, 1175.0}  [600], {-2625.0, -2225.0, -1825.0, -1425.0, -1025.0, -625.0, -425.0, -225.0, -25.0, 75.0, 175.0, 275.0, 305.0, 335.0, 355.0, 375.0, 395.0, 415.0, 445.0, 475.0, 575.0, 675.0, 775.0, 975.0, 1175.0, 1375.0, 1775.0, 2175.0, 2575.0, 2975.0, 3375.0} |
| A_6_-DNA  A16-C8  pH 5.4  20.0 °C  (*R*_1ρ_) | [150, 200, 250, 300, 400, 500, 600, 700, 900, 1000, 1200, 1600, 2000, 2500, 3000] {0}  [200], {-625.0, -425.0, -225.0, -125.0, -25.0, 75.0, 125.0, 175.0, 225.0, 275.0, 315.0, 345.0, 375.0, 405.0, 435.0, 475.0, 525.0, 575.0, 625.0, 675.0, 775.0}  [400], {-1225.0, -825.0, -625.0, -425.0, -225.0, -75.0, 75.0, 175.0, 225.0, 275.0, 315.0, 345.0, 375.0, 405.0, 435.0, 475.0, 525.0, 575.0, 675.0, 825.0, 975.0, 1175.0, 1375.0, 1575.0}  [600], {-2225.0, -1825.0, -1425.0, -1025.0, -625.0, -425.0, -225.0, -25.0, 75.0, 175.0, 275.0, 305.0, 335.0, 375.0, 415.0, 445.0, 475.0, 575.0, 675.0, 775.0, 975.0, 1175.0, 1375.0, 1775.0, 2175.0}  [1000], {-3125.0, -2625.0, -2025.0, -1425.0, -825.0, -525.0, -225.0, 75.0, 225.0, 325.0, 375.0, 425.0, 525.0, 675.0, 975.0, 1275.0, 1575.0, 2175.0, 2775.0, 3375.0} |
| A_6_-DNA  A16-C8  pH 5.4  22.5 °C  (*R*_1ρ_) | [100, 150, 200, 250, 300, 400, 500, 600, 700, 900, 1000, 1200, 1600, 2000, 2500, 3000] {0}  [200], {-625.0, -425.0, -225.0, -125.0, -25.0, 75.0, 125.0, 175.0, 225.0, 275.0, 315.0, 345.0, 375.0, 405.0, 435.0, 475.0, 525.0, 575.0, 625.0, 675.0, 775.0}  [400], {-1225.0, -825.0, -625.0, -425.0, -225.0, -75.0, 75.0, 175.0, 225.0, 275.0, 315.0, 345.0, 375.0, 405.0, 435.0, 475.0, 525.0, 575.0, 675.0, 825.0, 975.0, 1175.0, 1375.0, 1575.0}  [600], {-2225.0, -1825.0, -1425.0, -1025.0, -625.0, -425.0, -225.0, -25.0, 75.0, 175.0, 275.0, 305.0, 335.0, 375.0, 415.0, 445.0, 475.0, 575.0, 675.0, 775.0, 975.0, 1175.0, 1375.0, 1775.0, 2175.0}  [1000], {-3125.0, -2625.0, -2025.0, -1425.0, -825.0, -525.0, -225.0, 75.0, 225.0, 325.0, 375.0, 425.0, 525.0, 675.0, 975.0, 1275.0, 1575.0, 2175.0, 2775.0, 3375.0} |
| A_6_-DNA  A16-C8  pH 5.4  25 °C  (*R*_1ρ_) | [150, 200, 250, 300, 400, 500, 600, 700, 900, 1000, 1200, 1600, 2000, 2500, 3000] {0}  [200], {-625.0, -225.0, -125.0, -25.0, 75.0, 125.0, 175.0, 225.0, 275.0, 315.0, 345.0, 375.0, 405.0, 435.0, 475.0, 525.0, 575.0, 625.0, 675.0, 775.0}  [400], {-1225.0, -825.0, -625.0, -425.0, -225.0, -75.0, 75.0, 175.0, 225.0, 275.0, 315.0, 345.0, 375.0, 405.0, 435.0, 475.0, 525.0, 575.0, 675.0, 825.0, 975.0, 1175.0, 1375.0, 1575.0}  [600], {-2225.0, -1825.0, -1425.0, -1025.0, -625.0, -425.0, -225.0, -25.0, 75.0, 175.0, 275.0, 305.0, 335.0, 375.0, 415.0, 445.0, 475.0, 575.0, 675.0, 775.0, 975.0, 1175.0, 1375.0, 1775.0, 2175.0}  [1000], {-3125.0, -2625.0, -2025.0, -1425.0, -825.0, -525.0, -225.0, 75.0, 225.0, 325.0, 375.0, 425.0, 525.0, 675.0, 975.0, 1275.0, 1575.0, 2175.0, 2775.0, 3375.0} |
| A_6_-DNA^rA16^  rA16-C8  pH 5.4  10 °C  (*R*_1ρ_) | [100, 150, 200, 250, 300, 350, 400, 500, 600, 700, 800, 1000, 1200, 1400, 1600, 1800, 2000, 2500, 3000, 3500] {0}  [100], {-360.0, -300.0, -240.0, -180.0, -120.0, -90.0, -60.0, -30.0, 30.0, 60.0, 90.0, 120.0, 150.0, 180.0, 210.0, 240.0, 270.0, 300.0, 330.0, 360.0}  [150], {-500.0, -480.0, -450.0, -420.0, -400.0, -360.0, -350.0, -300.0, -300.0, -250.0, -240.0, -200.0, -180.0, -150.0, -120.0, -100.0, -60.0, -50.0, 50.0, 60.0, 100.0, 120.0, 150.0, 180.0, 200.0, 240.0, 250.0, 300.0, 300.0, 350.0, 360.0, 390.0, 400.0, 420.0, 450.0, 450.0, 480.0, 500.0, 510.0, 540.0}  [200], {-600.0, -540.0, -480.0, -420.0, -360.0, -300.0, -270.0, -240.0, -180.0, -120.0, -60.0, 60.0, 120.0, 180.0, 240.0, 270.0, 300.0, 360.0, 390.0, 420.0, 450.0, 480.0, 510.0, 540.0, 570.0, 600.0, 630.0, 660.0}  [250], {-720.0, -720.0, -660.0, -600.0, -600.0, -540.0, -480.0, -450.0, -420.0, -400.0, -360.0, -350.0, -300.0, -300.0, -240.0, -200.0, -180.0, -150.0, -120.0, -100.0, -60.0, -50.0, 50.0, 60.0, 100.0, 120.0, 150.0, 180.0, 200.0, 240.0, 300.0, 300.0, 350.0, 360.0, 400.0, 420.0, 450.0, 450.0, 480.0, 510.0, 540.0, 570.0, 600.0, 600.0, 630.0, 660.0, 720.0, 720.0}  [400], {-1200.0, -1000.0, -800.0, -700.0, -600.0, -500.0, -400.0, -300.0, -200.0, -100.0, 100.0, 200.0, 300.0, 400.0, 500.0, 600.0, 700.0, 800.0, 1000.0, 1200.0}  [500], {-1600.0, -1400.0, -1200.0, -1000.0, -800.0, -600.0, -500.0, -400.0, -300.0, -200.0, -100.0, 100.0, 200.0, 300.0, 400.0, 440.0, 480.0, 520.0, 560.0, 600.0, 640.0, 800.0, 1000.0, 1200.0, 1400.0, 1600.0}  [1000], {-3000.0, -2400.0, -1800.0, -1400.0, -1100.0, -800.0, -600.0, -400.0, -200.0, -100.0, 100.0, 200.0, 400.0, 600.0, 800.0, 1100.0, 1400.0, 1800.0, 2400.0, 3000.0} |
| A_6_-DNA^rA16^  rA16-C8  pH 5.4  12.5 °C  (*R*_1ρ_) | [100, 150, 200, 250, 300, 350, 400, 500, 600, 700, 800, 1000, 1200, 1400, 1600, 1800, 2000, 2500, 3000, 3500] {0}  [150], {-500.0, -450.0, -400.0, -350.0, -300.0, -250.0, -200.0, -150.0, -100.0, -50.0, 50.0, 100.0, 150.0, 200.0, 250.0, 300.0, 350.0, 400.0, 450.0, 500.0}  [250], {-720.0, -600.0, -450.0, -400.0, -350.0, -300.0, -200.0, -150.0, -100.0, -50.0, 50.0, 100.0, 150.0, 200.0, 300.0, 350.0, 400.0, 450.0, 600.0, 720.0}  [400], {-1200.0, -1000.0, -800.0, -700.0, -600.0, -500.0, -400.0, -300.0, -200.0, -100.0, 100.0, 200.0, 300.0, 400.0, 500.0, 600.0, 700.0, 800.0, 1000.0, 1200.0}  [1000], {-3000.0, -2400.0, -1800.0, -1400.0, -1100.0, -800.0, -600.0, -400.0, -200.0, -100.0, 100.0, 200.0, 400.0, 600.0, 800.0, 1100.0, 1400.0, 1800.0, 2400.0, 3000.0} |
| A_6_-DNA^rA16^  rA16-C8  pH 5.4  15 °C  (*R*_1ρ_) | [100, 150, 200, 250, 300, 350, 400, 500, 600, 700, 800, 1000, 1200, 1400, 1600, 1800, 2000, 2500, 3000, 3500] {0}  [150], {-500.0, -450.0, -400.0, -350.0, -300.0, -250.0, -200.0, -150.0, -100.0, -50.0, 50.0, 100.0, 150.0, 200.0, 250.0, 300.0, 350.0, 400.0, 450.0, 500.0}  [250], {-720.0, -600.0, -450.0, -400.0, -350.0, -300.0, -200.0, -150.0, -100.0, -50.0, 50.0, 100.0, 150.0, 200.0, 300.0, 350.0, 400.0, 450.0, 600.0, 720.0}  [400], {-1200.0, -1000.0, -800.0, -700.0, -600.0, -500.0, -400.0, -300.0, -200.0, -100.0, 100.0, 200.0, 300.0, 400.0, 500.0, 600.0, 700.0, 800.0, 1000.0, 1200.0}  [1000], {-3000.0, -2400.0, -1800.0, -1400.0, -1100.0, -800.0, -600.0, -400.0, -200.0, -100.0, 100.0, 200.0, 400.0, 600.0, 800.0, 1100.0, 1400.0, 1800.0, 2400.0, 3000.0} |
| A_6_-DNA  A16-C8  pH 5.4HS  25 °C  (*R*_1ρ_) | [150, 200, 250, 300, 400, 500, 600, 700, 900, 1000, 1200, 1400, 1600, 2000, 2500, 3000] {0}  [200], {-500.0, -300.0, -200.0, -100.0, 50.0, 100.0, 150.0, 200.0, 220.0, 240.0, 260.0, 280.0, 300.0, 320.0, 340.0, 360.0, 380.0, 450.0, 500.0, 600.0, 700.0}  [400], {-900.0, -700.0, -500.0, -300.0, -150.0, 100.0, 150.0, 200.0, 220.0, 240.0, 260.0, 280.0, 300.0, 320.0, 340.0, 360.0, 380.0, 400.0, 450.0, 500.0, 600.0, 750.0, 900.0, 1100.0}  [600], {-1500.0, -1100.0, -700.0, -500.0, -300.0, -100.0, 100.0, 200.0, 230.0, 260.0, 280.0, 300.0, 320.0, 340.0, 370.0, 400.0, 500.0, 600.0, 700.0, 900.0, 1100.0, 1300.0, 1700.0, 2100.0}  [1000], {-3200.0, -2100.0, -1500.0, -900.0, -600.0, -300.0, 150.0, 250.0, 300.0, 350.0, 450.0, 600.0, 900.0, 1200.0, 1500.0, 2100.0} |
| A_6_-DNA  A16-C1'  pH 5.4HS  25 °C  (*R*_1ρ_) | [150, 200, 250, 300, 400, 500, 600, 700, 900, 1000, 1200, 1400, 1600, 2000, 2500, 3000] {0}  [200], {-600.0, -400.0, -200.0, -100.0, 100.0, 150.0, 200.0, 250.0, 300.0, 320.0, 340.0, 360.0, 380.0, 400.0, 420.0, 440.0, 460.0, 480.0, 500.0, 550.0, 600.0, 650.0, 700.0}  [400], {-800.0, -600.0, -400.0, -200.0, -50.0, 100.0, 200.0, 250.0, 300.0, 320.0, 340.0, 360.0, 380.0, 400.0, 420.0, 440.0, 460.0, 480.0, 500.0, 550.0, 600.0, 700.0, 850.0, 1000.0, 1200.0, 1400.0}  [600], {-1400.0, -1000.0, -600.0, -400.0, -200.0, 100.0, 200.0, 300.0, 330.0, 360.0, 380.0, 400.0, 420.0, 440.0, 470.0, 500.0, 600.0, 700.0, 800.0, 1000.0, 1200.0, 1400.0, 1800.0}  [1000], {-3100.0, -2600.0, -2000.0, -1400.0, -800.0, -500.0, -200.0, 100.0, 250.0, 350.0, 400.0, 450.0, 550.0, 700.0, 1000.0, 1300.0, 1600.0, 2200.0, 2800.0, 3400.0} |
| A_6_-DNA^m1G10^  A16-C8  pH 5.4HS  25 °C  (*R*_1ρ_) | [150, 200, 250, 300, 400, 500, 600, 700, 900, 1000, 1200, 1400, 1600, 2000, 2500, 3000] {0}  [200], {-600.0, -400.0, -200.0, -100.0, 100.0, 150.0, 200.0, 250.0, 300.0, 320.0, 340.0, 360.0, 380.0, 400.0, 420.0, 440.0, 460.0, 480.0, 500.0, 550.0, 600.0, 650.0, 700.0}  [400], {-1200.0, -800.0, -600.0, -400.0, -200.0, -50.0, 100.0, 200.0, 250.0, 300.0, 320.0, 340.0, 360.0, 380.0, 400.0, 420.0, 440.0, 460.0, 480.0, 500.0, 550.0, 600.0, 700.0, 850.0, 1000.0, 1200.0, 1400.0}  [600], {-1800.0, -1400.0, -1000.0, -600.0, -400.0, -200.0, 100.0, 200.0, 300.0, 330.0, 360.0, 380.0, 400.0, 420.0, 440.0, 470.0, 500.0, 600.0, 700.0, 800.0, 1000.0, 1200.0, 1400.0, 1800.0}  [1000], {-3100.0, -2600.0, -2000.0, -1400.0, -800.0, -500.0, -200.0, 100.0, 250.0, 350.0, 400.0, 450.0, 550.0, 700.0, 1000.0, 1300.0, 1600.0, 2200.0, 2800.0, 3400.0} |
| A_6_-DNA^m1G10^  A16-C1'  pH 5.4HS  25 °C  (*R*_1ρ_) | [200, 250, 300, 400, 500, 600, 700, 900, 1000, 1200, 1400, 1600, 2000, 2500, 3000] {0}  [200], {-600.0, -400.0, -200.0, -100.0, 100.0, 150.0, 200.0, 250.0, 300.0, 320.0, 340.0, 360.0, 380.0, 400.0, 420.0, 440.0, 460.0, 480.0, 500.0, 550.0, 600.0, 650.0, 700.0}  [400], {-1200.0, -800.0, -600.0, -400.0, -200.0, -50.0, 100.0, 200.0, 250.0, 300.0, 320.0, 340.0, 360.0, 380.0, 400.0, 420.0, 440.0, 460.0, 480.0, 500.0, 550.0, 600.0, 700.0, 850.0, 1000.0, 1200.0, 1400.0}  [600], {-1800.0, -1400.0, -1000.0, -600.0, -400.0, -200.0, 100.0, 200.0, 300.0, 330.0, 360.0, 380.0, 400.0, 420.0, 440.0, 470.0, 500.0, 600.0, 700.0, 800.0, 1000.0, 1200.0, 1400.0, 1800.0}  [1000], {-3100.0, -2600.0, -2000.0, -1400.0, -800.0, -500.0, -200.0, 100.0, 250.0, 350.0, 400.0, 450.0, 550.0, 700.0, 1000.0, 1300.0, 1600.0, 2200.0, 2800.0, 3400.0} |
| A_2_-DNA  A16-C8  pH 6.8  25 °C  (*R*_1ρ_) | [100, 150, 200, 250, 300, 400, 500, 600, 700, 900, 1000, 1200, 1400, 1600, 2000, 2500, 3000] {0}  [200], {-700.0, -500.0, -300.0, -200.0, -100.0, 50.0, 100.0, 150.0, 200.0, 220.0, 240.0, 260.0, 280.0, 300.0, 320.0, 340.0, 360.0, 380.0, 400.0, 450.0, 500.0, 550.0, 600.0, 700.0}  [400], {-1300.0, -900.0, -700.0, -500.0, -300.0, -150.0, 100.0, 150.0, 200.0, 220.0, 240.0, 260.0, 280.0, 300.0, 320.0, 340.0, 360.0, 380.0, 400.0, 450.0, 500.0, 600.0, 750.0, 900.0, 1100.0, 1300.0}  [600], {-1900.0, -1500.0, -1100.0, -700.0, -500.0, -300.0, -100.0, 100.0, 200.0, 230.0, 260.0, 280.0, 300.0, 320.0, 340.0, 370.0, 400.0, 500.0, 600.0, 700.0, 900.0, 1100.0, 1300.0, 1700.0, 2100.0}  [1000], {-3200.0, -2700.0, -2100.0, -1500.0, -900.0, -600.0, -300.0, 150.0, 250.0, 300.0, 350.0, 450.0, 600.0, 900.0, 1200.0, 1500.0, 2100.0, 2700.0, 3300.0} |
| A_6_-DNA  A16-C8  pH 6.8  25 °C  (*R*_1ρ_) | [150, 200, 250, 300, 400, 500, 600, 700, 900, 1000, 1200, 1400, 1600, 2000, 2500] {0}  [200], {-600.0, -400.0, -200.0, -100.0, 50.0, 100.0, 150.0, 200.0, 250.0, 300.0, 400.0, 400.0, 450.0, 500.0, 550.0}  [400], {-1200.0, -1000.0, -800.0, -650.0, -500.0, -350.0, -200.0, -50.0, 100.0, 150.0, 200.0, 250.0, 300.0, 350.0, 400.0, 400.0, 450.0, 500.0, 550.0, 600.0, 650.0, 700.0, 850.0, 1000.0, 1150.0}  [600], {-1800.0, -1400.0, -600.0, -400.0, -200.0, 100.0, 200.0, 250.0, 300.0, 350.0, 400.0, 400.0, 450.0, 500.0, 550.0, 600.0, 700.0, 800.0, 1000.0, 1400.0, 1800.0}  [1000], {-2000.0, -1400.0, -800.0, -500.0, -200.0, 100.0, 250.0, 350.0, 400.0, 400.0, 450.0, 550.0, 700.0, 1000.0, 1300.0, 1600.0, 2200.0} |
| A_6_-DNA  A21-C1'  pH 6.8  25 °C  (*R*_1ρ_) | [200, 250, 300, 400, 500, 600, 700, 900, 1000, 1200, 1400, 1600, 2000, 2500, 3000] {0}  [200], {-600.0, -400.0, -200.0, -100.0, 100.0, 150.0, 200.0, 250.0, 300.0, 320.0, 340.0, 360.0, 380.0, 400.0, 420.0, 440.0, 460.0, 480.0, 500.0, 550.0, 600.0, 650.0, 700.0}  [400], {-1200.0, -800.0, -600.0, -400.0, -200.0, -50.0, 100.0, 200.0, 250.0, 300.0, 320.0, 340.0, 360.0, 380.0, 400.0, 420.0, 440.0, 460.0, 480.0, 500.0, 550.0, 600.0, 700.0, 850.0, 1000.0, 1200.0, 1400.0}  [600], {-1800.0, -1400.0, -1000.0, -600.0, -400.0, -200.0, 100.0, 200.0, 300.0, 330.0, 360.0, 380.0, 400.0, 420.0, 440.0, 470.0, 500.0, 600.0, 700.0, 800.0, 1000.0, 1200.0, 1400.0, 1800.0}  [1000], {-3100.0, -2600.0, -2000.0, -1400.0, -800.0, -500.0, -200.0, 100.0, 250.0, 350.0, 400.0, 450.0, 550.0, 700.0, 1000.0, 1300.0, 1600.0, 2200.0, 2800.0, 3400.0} |
| A_6_-DNA  A21-C1'  pH 6.8  30 °C  (*R*_1ρ_) | [200, 250, 300, 400, 500, 600, 700, 900, 1000, 1200, 1400, 1600, 2000, 2500, 3000] {0}  [150], {-625.0, -375.0, -225.0, -125.0, -25.0, 175.0, 225.0, 275.0, 295.0, 315.0, 335.0, 355.0, 375.0, 395.0, 415.0, 435.0, 455.0, 475.0, 525.0, 575.0, 625.0, 675.0}  [400], {-1625.0, -1225.0, -825.0, -625.0, -425.0, -225.0, -75.0, 75.0, 175.0, 225.0, 275.0, 295.0, 315.0, 335.0, 355.0, 375.0, 395.0, 415.0, 435.0, 455.0, 475.0, 525.0, 575.0, 675.0, 825.0, 975.0, 1175.0, 1375.0, 1575.0, 1975.0}  [600], {-2625.0, -2225.0, -1825.0, -1425.0, -1025.0, -625.0, -425.0, -225.0, -25.0, 75.0, 175.0, 275.0, 305.0, 335.0, 355.0, 375.0, 395.0, 415.0, 445.0, 475.0, 575.0, 675.0, 775.0, 975.0, 1175.0, 1375.0, 1775.0, 2175.0, 2575.0, 2975.0}  [1000], {-3125.0, -2625.0, -2025.0, -1425.0, -825.0, -525.0, -225.0, 75.0, 225.0, 325.0, 375.0, 425.0, 525.0, 675.0, 975.0, 1275.0, 1575.0, 2175.0, 2775.0, 3375.0} |
| **G-C^+^ Hoogsteen** | |
| A_2_-DNA  G10-C8  pH 4.4HS  25 °C  (*R*_1ρ_) | [150, 200, 250, 300, 400, 500, 600, 700, 900, 1000, 1200, 1400, 1600, 2000, 2500] {0}  [200], {-600.0, -400.0, -200.0, -100.0, 50.0, 100.0, 150.0, 200.0, 250.0, 300.0, 350.0, 400.0, 400.0, 450.0, 500.0, 550.0, 600.0}  [400], {-1200.0, -1000.0, -800.0, -650.0, -500.0, -350.0, -200.0, -50.0, 100.0, 150.0, 200.0, 250.0, 300.0, 350.0, 400.0, 400.0, 450.0, 500.0, 550.0, 600.0, 650.0, 700.0, 850.0, 1000.0, 1150.0}  [600], {-1800.0, -1400.0, -1000.0, -600.0, -400.0, -200.0, 100.0, 200.0, 250.0, 300.0, 350.0, 400.0, 400.0, 450.0, 500.0, 550.0, 600.0, 700.0, 800.0, 1000.0, 1200.0, 1400.0, 1800.0}  [1000], {-2000.0, -1400.0, -800.0, -500.0, -200.0, 100.0, 250.0, 350.0, 400.0, 400.0, 450.0, 550.0, 700.0, 1000.0, 1300.0, 1600.0, 2200.0} |
| AcDNA  G7-C1'  pH 5.3HS  25 °C  (*R*_1ρ_) | [150, 200, 250, 300, 400, 500, 700, 900, 1200, 1600, 2000, 2400] {0}  [200], {-600.0, -500.0, -400.0, -300.0, -200.0, -100.0, -50.0, 50.0, 100.0, 150.0, 200.0, 250.0, 300.0, 350.0, 400.0, 500.0, 550.0, 600.0}  [300], {-800.0, -600.0, -400.0, -300.0, -200.0, -100.0, 100.0, 200.0, 300.0, 400.0, 500.0, 600.0, 700.0, 800.0, 900.0}  [400], {-800.0, -600.0, -400.0, -300.0, -200.0, -100.0, 100.0, 200.0, 300.0, 400.0, 500.0, 600.0, 700.0, 800.0, 900.0, 1100.0}  [700], {-900.0, -700.0, -500.0, -300.0, -200.0, -100.0, 100.0, 200.0, 300.0, 400.0, 500.0, 600.0, 700.0, 800.0, 900.0, 1100.0, 1300.0, 1500.0, 1700.0} |
| AcDNA  G7-C8  pH 5.3HS  25 °C  (*R*_1ρ_) | [200, 250, 300, 400, 500, 600, 700, 800, 900, 1000, 1200, 1400, 1800, 2200, 2600, 3000] {0}  [200], {-600.0, -500.0, -400.0, -300.0, -200.0, -100.0, 50.0, 100.0, 200.0, 250.0, 300.0, 350.0, 400.0, 450.0, 500.0, 600.0}  [300], {-900.0, -800.0, -700.0, -600.0, -500.0, -400.0, -300.0, -200.0, -100.0, 100.0, 200.0, 300.0, 400.0, 500.0, 600.0, 700.0, 800.0, 900.0}  [400], {-1000.0, -800.0, -500.0, -300.0, -200.0, -100.0, 100.0, 200.0, 300.0, 400.0, 500.0, 600.0, 700.0, 800.0, 900.0, 1000.0, 1200.0}  [700], {-1700.0, -1200.0, -800.0, -600.0, -400.0, -200.0, 100.0, 200.0, 300.0, 400.0, 500.0, 600.0, 700.0, 800.0, 900.0, 1000.0, 1200.0, 1400.0, 1600.0, 2000.0}  [1000], {-2500.0, -2000.0, -1600.0, -1000.0, -600.0, -400.0, -200.0, 100.0, 200.0, 300.0, 400.0, 500.0, 600.0, 700.0, 800.0, 900.0, 1100.0, 1300.0, 1500.0, 1800.0, 2000.0, 2600.0, 3000.0} |
| AcDNA  G7-N1  pH 5.3HS  25 °C  (*R*_1ρ_) | [100, 150, 200, 250, 300, 400, 500, 600, 700, 900, 1200, 1600] {0}  [100], {-350.0, -300.0, -250.0, -200.0, -150.0, -100.0, -50.0, 50.0, 100.0, 150.0, 200.0, 250.0, 300.0}  [200], {-600.0, -500.0, -400.0, -350.0, -300.0, -200.0, -150.0, -100.0, -50.0, 50.0, 100.0, 150.0, 200.0, 300.0, 400.0}  [300], {-900.0, -800.0, -700.0, -600.0, -500.0, -400.0, -300.0, -200.0, -100.0, 100.0, 200.0, 300.0, 400.0, 500.0, 600.0, 800.0}  [400], {-1100.0, -900.0, -800.0, -700.0, -600.0, -500.0, -400.0, -300.0, -200.0, -100.0, 100.0, 200.0, 300.0, 400.0, 500.0, 700.0}  [700], {-1600.0, -1300.0, -1000.0, -800.0, -600.0, -500.0, -400.0, -300.0, -200.0, -100.0, 100.0, 200.0, 300.0, 400.0, 600.0, 800.0, 1000.0, 1300.0} |
| A_2_-DNA  G10-C8  pH 5.4  25 °C  (*R*_1ρ_) | [150, 200, 250, 300, 400, 500, 600, 700, 900, 1000, 1400, 1600, 2000, 2500] {0}  [200], {-600.0, -400.0, -200.0, -100.0, 50.0, 100.0, 150.0, 200.0, 250.0, 300.0, 350.0, 400.0, 400.0, 450.0, 550.0}  [400], {-1200.0, -1000.0, -800.0, -650.0, -500.0, -350.0, -200.0, -50.0, 100.0, 150.0, 250.0, 300.0, 350.0, 400.0, 400.0, 450.0, 500.0, 550.0, 600.0, 650.0, 700.0, 850.0, 1150.0}  [600], {-1800.0, -600.0, -400.0, -200.0, 100.0, 200.0, 250.0, 300.0, 350.0, 400.0, 400.0, 450.0, 500.0, 550.0, 600.0, 700.0, 800.0, 1000.0, 1200.0, 1400.0, 1800.0}  [1000], {-2000.0, -1400.0, -800.0, -500.0, -200.0, 100.0, 250.0, 350.0, 400.0, 400.0, 450.0, 550.0, 700.0, 1000.0, 1300.0, 2200.0} |
| A_2_-DNA  G10-C1'  pH 5.4  25 °C  (*R*_1ρ_) | [150, 200, 250, 300, 400, 500, 600, 700, 900, 1000, 1200, 1400, 1600, 2000, 2500] {0}  [200], {-600.0, -400.0, -200.0, -100.0, 50.0, 100.0, 150.0, 200.0, 250.0, 300.0, 350.0, 400.0, 400.0, 450.0, 550.0}  [400], {-1200.0, -1000.0, -800.0, -650.0, -500.0, -350.0, -200.0, -50.0, 100.0, 150.0, 200.0, 250.0, 300.0, 350.0, 400.0, 400.0, 450.0, 500.0, 550.0, 600.0, 650.0, 700.0, 850.0, 1000.0, 1150.0}  [600], {-1800.0, -1400.0, -600.0, -400.0, -200.0, 100.0, 200.0, 250.0, 300.0, 350.0, 400.0, 400.0, 450.0, 500.0, 550.0, 600.0, 700.0, 800.0, 1000.0, 1400.0}  [1000], {-2000.0, -1400.0, -800.0, -500.0, 100.0, 250.0, 350.0, 400.0, 400.0, 450.0, 550.0, 700.0, 1000.0, 1300.0, 1600.0, 2200.0} |
| A_6_-DNA  G10-C8  pH 5.4  25 °C  (*R*_1ρ_) | [100, 150, 200, 250, 300, 400, 500, 600, 700, 900, 1000, 1200, 1400, 1600, 2000, 2500, 3000] {0}  [200], {-600.0, -400.0, -200.0, -100.0, 100.0, 150.0, 200.0, 250.0, 300.0, 320.0, 340.0, 360.0, 380.0, 400.0, 420.0, 440.0, 460.0, 480.0, 500.0, 550.0, 600.0, 650.0, 700.0}  [400], {-1200.0, -800.0, -600.0, -400.0, -200.0, -50.0, 100.0, 200.0, 250.0, 300.0, 320.0, 340.0, 360.0, 380.0, 400.0, 420.0, 440.0, 460.0, 480.0, 500.0, 550.0, 600.0, 700.0, 850.0, 1200.0, 1400.0}  [600], {-1800.0, -1400.0, -1000.0, -600.0, -400.0, -200.0, 100.0, 200.0, 300.0, 330.0, 360.0, 380.0, 400.0, 420.0, 440.0, 470.0, 500.0, 600.0, 700.0, 800.0, 1000.0, 1200.0, 1400.0, 1800.0}  [1000], {-3100.0, -2600.0, -2000.0, -1400.0, -800.0, -500.0, -200.0, 100.0, 250.0, 350.0, 400.0, 450.0, 550.0, 700.0, 1000.0, 1300.0, 1600.0, 2200.0, 2800.0, 3400.0} |
| A_6_-DNA  G10-C1'  pH 5.4  25 °C  (*R*_1ρ_) | [150, 200, 250, 300, 400, 500, 600, 700, 900, 1000, 1200, 1400, 1600, 2000, 2500, 3000] {0}  [200], {-600.0, -400.0, -200.0, -100.0, 100.0, 150.0, 200.0, 250.0, 300.0, 320.0, 340.0, 360.0, 380.0, 400.0, 420.0, 440.0, 460.0, 480.0, 500.0, 550.0, 600.0, 650.0, 700.0}  [400], {-1200.0, -800.0, -600.0, -400.0, -200.0, -50.0, 100.0, 200.0, 250.0, 300.0, 320.0, 340.0, 360.0, 380.0, 400.0, 420.0, 440.0, 460.0, 480.0, 500.0, 550.0, 600.0, 700.0, 850.0, 1000.0, 1200.0, 1400.0}  [600], {-1800.0, -1400.0, -1000.0, -600.0, -400.0, -200.0, 100.0, 200.0, 300.0, 330.0, 360.0, 380.0, 400.0, 420.0, 440.0, 470.0, 500.0, 600.0, 700.0, 800.0, 1000.0, 1200.0, 1400.0, 1800.0}  [1000], {-3100.0, -2600.0, -2000.0, -1400.0, -800.0, -500.0, -200.0, 100.0, 250.0, 350.0, 400.0, 450.0, 550.0, 700.0, 1000.0, 1300.0, 1600.0, 2200.0, 2800.0, 3400.0} |
| A_6_-DNA^rG10^  C15-C6  pH 5.4  25 °C  (*R*_1ρ_) | [200, 250, 300, 350, 400, 500, 600, 700, 800, 900, 1000, 1200, 1400, 1600, 1800, 2000, 2500, 3000, 3500] {0}  [150], {-522.0, -464.0, -406.0, -348.0, -290.0, -232.0, -174.0, -116.0, -58.0, 10.0, 58.0, 116.0, 174.0, 232.0, 290.0, 348.0, 406.0, 464.0, 522.0}  [200], {-702.0, -624.0, -546.0, -468.0, -390.0, -312.0, -234.0, -156.0, -78.0, -10.0, 10.0, 78.0, 156.0, 234.0, 312.0, 390.0, 468.0, 546.0, 624.0, 702.0}  [250], {-873.0, -776.0, -679.0, -582.0, -485.0, -388.0, -291.0, -194.0, -97.0, -10.0, 10.0, 97.0, 194.0, 291.0, 388.0, 485.0, 582.0, 679.0, 776.0, 873.0}  [300], {-1053.0, -936.0, -819.0, -702.0, -585.0, -468.0, -351.0, -234.0, -117.0, -10.0, 10.0, 117.0, 234.0, 351.0, 468.0, 585.0, 702.0, 819.0, 936.0, 1053.0}  [350], {-1224.0, -1088.0, -952.0, -816.0, -680.0, -544.0, -408.0, -272.0, -136.0, -10.0, 10.0, 136.0, 272.0, 408.0, 544.0, 680.0, 816.0, 952.0, 1088.0, 1224.0}  [450], {-1575.0, -1400.0, -1225.0, -1050.0, -875.0, -700.0, -525.0, -350.0, -175.0, -10.0, 10.0, 175.0, 350.0, 525.0, 700.0, 875.0, 1050.0, 1225.0, 1400.0, 1575.0} |
| A_6_-DNA  G10-C8  pH 5.4HS  25 °C  (*R*_1ρ_) | [100, 150, 200, 250, 300, 400, 500, 600, 700, 900, 1000, 1200, 1400, 1600, 2000, 2500, 3000] {0}  [200], {-600.0, -400.0, -200.0, -100.0, 100.0, 150.0, 200.0, 250.0, 300.0, 320.0, 340.0, 360.0, 380.0, 400.0, 420.0, 440.0, 480.0, 500.0, 550.0, 600.0, 650.0, 700.0}  [400], {-1200.0, -800.0, -600.0, -400.0, -200.0, -50.0, 100.0, 200.0, 250.0, 300.0, 320.0, 340.0, 360.0, 380.0, 400.0, 420.0, 440.0, 460.0, 480.0, 500.0, 550.0, 600.0, 700.0, 850.0, 1000.0, 1400.0}  [600], {-1800.0, -1400.0, -1000.0, -600.0, -400.0, -200.0, 100.0, 200.0, 300.0, 330.0, 360.0, 380.0, 400.0, 420.0, 440.0, 470.0, 500.0, 600.0, 700.0, 800.0, 1000.0, 1200.0, 1400.0, 1800.0}  [1000], {-3100.0, -2600.0, -2000.0, -1400.0, -800.0, -500.0, -200.0, 100.0, 250.0, 350.0, 400.0, 450.0, 550.0, 700.0, 1000.0, 1300.0, 1600.0, 2200.0, 2800.0, 3400.0} |
| A_6_-DNA  G10-C1'  pH 5.4HS  25 °C  (*R*_1ρ_) | [150, 200, 250, 300, 400, 500, 600, 700, 900, 1000, 1200, 1400, 1600, 2000, 2500, 3000] {0}  [200], {-600.0, -400.0, -200.0, -100.0, 100.0, 150.0, 200.0, 250.0, 300.0, 320.0, 340.0, 360.0, 380.0, 400.0, 420.0, 440.0, 460.0, 480.0, 500.0, 550.0, 600.0, 650.0, 700.0}  [400], {-1200.0, -800.0, -600.0, -400.0, -200.0, -50.0, 100.0, 200.0, 250.0, 300.0, 320.0, 340.0, 360.0, 380.0, 400.0, 420.0, 440.0, 460.0, 480.0, 500.0, 550.0, 600.0, 700.0, 850.0, 1200.0, 1400.0}  [600], {-1400.0, -600.0, -400.0, -200.0, 100.0, 200.0, 300.0, 330.0, 360.0, 380.0, 400.0, 420.0, 440.0, 470.0, 500.0, 600.0, 700.0, 800.0, 1000.0, 1200.0, 1400.0, 1800.0}  [1000], {-3100.0, -2600.0, -2000.0, -1400.0, -800.0, -500.0, -200.0, 100.0, 250.0, 350.0, 400.0, 450.0, 550.0, 700.0, 1000.0, 1300.0, 1600.0, 2200.0, 2800.0, 3400.0} |
| A_6_-DNA  G10-C1'  pH 6.8  25 °C  (*R*_1ρ_) | [150, 200, 250, 300, 400, 500, 600, 700, 900, 1000, 1200, 1400, 1600, 2000, 2500] {0}  [150], {-300.0, -100.0, 60.0, 120.0, 160.0, 200.0, 240.0, 280.0, 320.0, 400.0, 400.0}  [200], {-200.0, -100.0, 50.0, 150.0, 250.0, 300.0, 350.0, 400.0, 400.0, 450.0, 500.0, 550.0, 600.0}  [400], {-1200.0, -1000.0, -800.0, -650.0, -500.0, -350.0, -200.0, -50.0, 100.0, 150.0, 200.0, 250.0, 300.0, 350.0, 400.0, 400.0, 450.0, 500.0, 550.0, 600.0, 650.0, 700.0, 850.0, 1000.0, 1150.0}  [600], {-1800.0, -1400.0, -1000.0, -600.0, -400.0, -200.0, 100.0, 200.0, 250.0, 300.0, 350.0, 400.0, 400.0, 450.0, 500.0, 550.0, 600.0, 700.0, 800.0, 1000.0, 1200.0, 1400.0, 1800.0} |
| ***N*^6^-methylamino rotation in m^6^A** | |
| hpGGACU^m6A6^  m^6^A6-C2  pH 6.8  37 °C  (CEST) | [10.0], {-1000.0, -922.2, -844.4, -766.7, -688.9, -611.1, -533.3, -455.6, -377.8, -300.0, -300.0, -290.7, -281.4, -272.1, -262.8, -253.4, -244.1, -234.8, -225.5, -216.2, -206.9, -197.6, -188.3, -179.0, -169.7, -160.3, -151.0, -141.7, -132.4, -123.1, -113.8, -104.5, -95.2, -85.9, -76.6, -67.2, -57.9, -48.6, -39.3, -30.0, 30.0, 39.6, 49.2, 58.8, 68.4, 78.0, 87.6, 97.1, 106.7, 116.3, 125.9, 135.5, 145.1, 154.7, 164.3, 173.9, 183.5, 193.1, 202.7, 212.2, 221.8, 231.4, 241.0, 250.6, 260.2, 269.8, 279.4, 289.0, 298.6, 308.2, 317.8, 327.3, 336.9, 346.5, 356.1, 365.7, 375.3, 384.9, 394.5, 404.1, 413.7, 423.3, 432.9, 442.4, 452.0, 461.6, 471.2, 480.8, 490.4, 500.0, 500.0, 555.6, 611.1, 666.7, 722.2, 777.8, 833.3, 888.9, 944.4, 1000.0}  [20.0], {-1000.0, -922.2, -844.4, -766.7, -688.9, -611.1, -533.3, -455.6, -377.8, -300.0, -300.0, -290.7, -281.4, -272.1, -262.8, -253.4, -244.1, -234.8, -225.5, -216.2, -206.9, -197.6, -188.3, -179.0, -169.7, -160.3, -151.0, -141.7, -132.4, -123.1, -113.8, -104.5, -95.2, -85.9, -76.6, -67.2, -57.9, -48.6, -39.3, -30.0, 30.0, 39.6, 49.2, 58.8, 68.4, 78.0, 87.6, 97.1, 106.7, 116.3, 125.9, 135.5, 145.1, 154.7, 164.3, 173.9, 183.5, 193.1, 202.7, 212.2, 221.8, 231.4, 241.0, 250.6, 260.2, 269.8, 279.4, 289.0, 298.6, 308.2, 317.8, 327.3, 336.9, 346.5, 356.1, 365.7, 375.3, 384.9, 394.5, 404.1, 413.7, 423.3, 432.9, 442.4, 452.0, 461.6, 471.2, 480.8, 490.4, 500.0, 500.0, 555.6, 611.1, 666.7, 722.2, 777.8, 833.3, 888.9, 944.4, 1000.0}  [40.0], {-1000.0, -922.2, -844.4, -766.7, -688.9, -611.1, -533.3, -455.6, -377.8, -300.0, -300.0, -290.7, -281.4, -272.1, -262.8, -253.4, -244.1, -234.8, -225.5, -216.2, -206.9, -197.6, -188.3, -179.0, -169.7, -160.3, -151.0, -141.7, -132.4, -123.1, -113.8, -104.5, -95.2, -85.9, -76.6, -67.2, -57.9, -48.6, -39.3, -30.0, 30.0, 39.6, 49.2, 58.8, 68.4, 78.0, 87.6, 97.1, 106.7, 116.3, 125.9, 135.5, 145.1, 154.7, 164.3, 173.9, 183.5, 193.1, 202.7, 212.2, 221.8, 231.4, 241.0, 250.6, 260.2, 269.8, 279.4, 289.0, 298.6, 308.2, 317.8, 327.3, 336.9, 346.5, 356.1, 365.7, 375.3, 384.9, 394.5, 404.1, 413.7, 423.3, 432.9, 442.4, 452.0, 461.6, 471.2, 480.8, 490.4, 500.0, 500.0, 555.6, 611.1, 666.7, 722.2, 777.8, 833.3, 888.9, 944.4, 1000.0} |
| hpGGACU^m6A6^  m^6^A6-C10  pH 6.8  37 °C  (CEST) | [10.0], {-1000.0, -922.2, -844.4, -766.7, -688.9, -611.1, -533.3, -455.6, -377.8, -300.0, -300.0, -290.7, -281.4, -272.1, -262.8, -253.4, -244.1, -234.8, -225.5, -216.2, -206.9, -197.6, -188.3, -179.0, -169.7, -160.3, -151.0, -141.7, -132.4, -123.1, -113.8, -104.5, -95.2, -85.9, -76.6, -67.2, -57.9, -48.6, -39.3, -30.0, 30.0, 39.6, 49.2, 58.8, 68.4, 78.0, 87.6, 97.1, 106.7, 116.3, 125.9, 135.5, 145.1, 154.7, 164.3, 173.9, 183.5, 193.1, 202.7, 212.2, 221.8, 231.4, 241.0, 250.6, 260.2, 269.8, 279.4, 289.0, 298.6, 308.2, 317.8, 327.3, 336.9, 346.5, 356.1, 365.7, 375.3, 384.9, 394.5, 404.1, 413.7, 423.3, 432.9, 442.4, 452.0, 461.6, 471.2, 480.8, 490.4, 500.0, 500.0, 555.6, 611.1, 666.7, 722.2, 777.8, 833.3, 888.9, 944.4, 1000.0}  [20.0], {-1000.0, -922.2, -844.4, -766.7, -688.9, -611.1, -533.3, -455.6, -377.8, -300.0, -300.0, -290.7, -281.4, -272.1, -262.8, -253.4, -244.1, -234.8, -225.5, -216.2, -206.9, -197.6, -188.3, -179.0, -169.7, -160.3, -151.0, -141.7, -132.4, -123.1, -113.8, -104.5, -95.2, -85.9, -76.6, -67.2, -57.9, -48.6, -39.3, -30.0, 30.0, 39.6, 49.2, 58.8, 68.4, 78.0, 87.6, 97.1, 106.7, 116.3, 125.9, 135.5, 145.1, 154.7, 164.3, 173.9, 183.5, 193.1, 202.7, 212.2, 221.8, 231.4, 241.0, 250.6, 260.2, 269.8, 279.4, 289.0, 298.6, 308.2, 317.8, 327.3, 336.9, 346.5, 356.1, 365.7, 375.3, 384.9, 394.5, 404.1, 413.7, 423.3, 432.9, 442.4, 452.0, 461.6, 471.2, 480.8, 490.4, 500.0, 500.0, 555.6, 611.1, 666.7, 722.2, 777.8, 833.3, 888.9, 944.4, 1000.0}  [40.0], {-1000.0, -922.2, -844.4, -766.7, -688.9, -611.1, -533.3, -455.6, -377.8, -300.0, -300.0, -290.7, -281.4, -272.1, -262.8, -253.4, -244.1, -234.8, -225.5, -216.2, -206.9, -197.6, -188.3, -179.0, -169.7, -160.3, -151.0, -141.7, -132.4, -123.1, -113.8, -104.5, -95.2, -85.9, -76.6, -67.2, -57.9, -48.6, -39.3, -30.0, 30.0, 39.6, 49.2, 58.8, 68.4, 78.0, 87.6, 97.1, 106.7, 116.3, 125.9, 135.5, 145.1, 154.7, 164.3, 173.9, 183.5, 193.1, 202.7, 212.2, 221.8, 231.4, 241.0, 250.6, 260.2, 269.8, 279.4, 289.0, 298.6, 308.2, 317.8, 327.3, 336.9, 346.5, 356.1, 365.7, 375.3, 384.9, 394.5, 404.1, 413.7, 423.3, 432.9, 442.4, 452.0, 461.6, 471.2, 480.8, 490.4, 500.0, 500.0, 555.6, 611.1, 666.7, 722.2, 777.8, 833.3, 888.9, 944.4, 1000.0} |
| hpGGACU^m6A6^  m^6^A6-C2  pH 6.8  55 °C  (*R*_1ρ_) | [150, 200, 250, 300, 400, 500, 600, 700, 900, 1000, 1200, 1600, 2000, 2500] {0}  [150], {-400.0, -340.0, -280.0, -240.0, -200.0, -160.0, -120.0, -80.0, -40.0, 40.0, 80.0, 120.0, 160.0, 200.0, 240.0, 280.0, 340.0, 400.0}  [200], {-600.0, -500.0, -400.0, -350.0, -300.0, -250.0, -200.0, -150.0, -100.0, -50.0, 50.0, 100.0, 150.0, 200.0, 250.0, 300.0, 350.0, 400.0, 500.0, 600.0}  [400], {-1200.0, -1050.0, -900.0, -600.0, -450.0, -300.0, -250.0, -200.0, -150.0, -100.0, -50.0, 50.0, 100.0, 150.0, 200.0, 250.0, 300.0, 450.0, 600.0, 750.0, 900.0, 1050.0, 1200.0}  [1000], {-2400.0, -1800.0, -1200.0, -900.0, -600.0, -300.0, -150.0, -50.0, 50.0, 150.0, 300.0, 600.0, 900.0, 1200.0, 1800.0, 2400.0} |
| hpGGACU^m6A6^  m^6^A6-C2  pH 6.8  65 °C  (*R*_1ρ_) | [150, 200, 250, 400, 500, 600, 700, 900, 1000, 1200, 1400, 1600, 2000, 2500] {0}  [150], {-400.0, -340.0, -280.0, -240.0, -200.0, -160.0, -120.0, -80.0, -40.0, 40.0, 80.0, 120.0, 200.0, 240.0, 280.0, 340.0, 400.0}  [200], {-600.0, -500.0, -400.0, -350.0, -300.0, -250.0, -200.0, -150.0, -100.0, -50.0, 50.0, 100.0, 150.0, 200.0, 250.0, 300.0, 350.0, 400.0, 500.0, 600.0}  [400], {-1200.0, -1050.0, -900.0, -750.0, -600.0, -450.0, -300.0, -250.0, -200.0, -150.0, -100.0, -50.0, 50.0, 100.0, 150.0, 200.0, 250.0, 300.0, 450.0, 600.0, 750.0, 900.0, 1050.0, 1200.0}  [1000], {-2400.0, -1800.0, -1200.0, -900.0, -600.0, -300.0, -150.0, -50.0, 50.0, 150.0, 300.0, 600.0, 900.0, 1200.0, 1800.0, 2400.0} |
| A_6_-RNA^m6A16^  m^6^A16-C2  pH 6.8  37 °C  (*R*_1ρ_) | [150, 200, 250, 300, 400, 500, 600, 700, 900, 1000, 1200, 1400, 1600, 2000, 2500] {0}  [150], {-400.0, -340.0, -280.0, -240.0, -200.0, -160.0, -120.0, -80.0, -40.0, 40.0, 80.0, 120.0, 160.0, 200.0, 240.0, 280.0, 340.0, 400.0}  [200], {-600.0, -500.0, -400.0, -350.0, -300.0, -250.0, -200.0, -150.0, -100.0, -50.0, 50.0, 100.0, 150.0, 200.0, 250.0, 300.0, 350.0, 400.0, 500.0, 600.0}  [400], {-1200.0, -1050.0, -900.0, -750.0, -600.0, -450.0, -300.0, -250.0, -200.0, -150.0, -100.0, -50.0, 50.0, 100.0, 150.0, 200.0, 250.0, 300.0, 450.0, 600.0, 750.0, 900.0, 1050.0, 1200.0}  [1000], {-2400.0, -1800.0, -1200.0, -900.0, -600.0, -300.0, -150.0, -50.0, 50.0, 150.0, 300.0, 600.0, 900.0, 1200.0, 1800.0, 2400.0} |
| **Hoogsteen Cooperativity** | |
| A_6_-DNA  A16-C8  pH 5.4 (D_2_O) 25 °C  (*R*_1ρ_) | [150, 200, 250, 300, 400, 500, 600, 700, 900, 1000, 1200, 1400, 1600, 2000, 2500] {0}  [200], {-600.0, -400.0, -300.0, -200.0, -150.0, -100.0, -50.0, 50.0, 100.0, 150.0, 200.0, 200.0, 250.0, 300.0, 350.0, 400.0, 450.0, 500.0, 550.0, 600.0}  [400], {-1200.0, -1000.0, -850.0, -700.0, -550.0, -400.0, -250.0, -100.0, -50.0, 50.0, 100.0, 150.0, 200.0, 200.0, 250.0, 300.0, 350.0, 400.0, 450.0, 500.0, 650.0, 800.0, 950.0, 1100.0}  [600], {-1600.0, -1200.0, -800.0, -600.0, -400.0, -200.0, -100.0, 50.0, 100.0, 150.0, 200.0, 200.0, 250.0, 300.0, 350.0, 400.0, 500.0, 600.0, 800.0, 1000.0, 1200.0, 1600.0}  [1000], {-2200.0, -1600.0, -1000.0, -700.0, -400.0, -100.0, 50.0, 150.0, 200.0, 200.0, 250.0, 350.0, 500.0, 800.0, 1100.0, 1400.0, 2000.0} |
| A_6_-DNA  A16-C1'  pH 5.4 (D_2_O) 25 °C  (*R*_1ρ_) | [150, 200, 250, 300, 400, 500, 600, 700, 900, 1000, 1200, 1400, 1600, 2000, 2500] {0}  [200], {-600.0, -400.0, -300.0, -200.0, -150.0, -100.0, -50.0, 50.0, 100.0, 150.0, 200.0, 200.0, 250.0, 300.0, 350.0, 400.0, 450.0, 500.0, 550.0, 600.0}  [400], {-1200.0, -1000.0, -850.0, -700.0, -550.0, -400.0, -250.0, -100.0, -50.0, 50.0, 100.0, 150.0, 200.0, 200.0, 250.0, 300.0, 350.0, 400.0, 450.0, 500.0, 650.0, 800.0, 950.0, 1100.0}  [600], {-1600.0, -1200.0, -800.0, -600.0, -400.0, -200.0, -100.0, 50.0, 100.0, 150.0, 200.0, 200.0, 250.0, 300.0, 350.0, 400.0, 500.0, 600.0, 800.0, 1000.0, 1200.0, 1600.0}  [1000], {-2200.0, -1600.0, -1000.0, -700.0, -400.0, -100.0, 50.0, 150.0, 200.0, 200.0, 250.0, 350.0, 500.0, 800.0, 1100.0, 1400.0, 2000.0} |
| A_6_-DNA  G10-C8  pH 5.4 (D_2_O) 25 °C  (*R*_1ρ_) | [150, 200, 250, 300, 400, 500, 600, 700, 900, 1000, 1200, 1400, 1600, 2000, 2500] {0}  [150], {-300.0, -100.0, 60.0, 120.0, 160.0, 200.0, 240.0, 280.0, 320.0, 360.0, 400.0, 400.0, 440.0}  [200], {-600.0, -400.0, -200.0, -100.0, 50.0, 100.0, 150.0, 200.0, 250.0, 300.0, 350.0, 400.0, 400.0, 450.0, 500.0, 550.0, 600.0}  [400], {-1200.0, -1000.0, -800.0, -650.0, -500.0, -350.0, -200.0, -50.0, 100.0, 150.0, 200.0, 250.0, 300.0, 350.0, 400.0, 400.0, 450.0, 500.0, 550.0, 600.0, 650.0, 700.0, 850.0, 1000.0, 1150.0}  [600], {-1800.0, -1400.0, -1000.0, -600.0, -400.0, -200.0, 100.0, 200.0, 250.0, 300.0, 350.0, 400.0, 400.0, 450.0, 500.0, 550.0, 600.0, 700.0, 800.0, 1000.0, 1200.0, 1400.0, 1800.0} |
| A_6_-DNA  G10-C1'  pH 5.4 (D_2_O) 25 °C  (*R*_1ρ_) | [150, 200, 250, 300, 400, 500, 600, 700, 900, 1000, 1200, 1400, 1600, 2000, 2500] {0}  [200], {-600.0, -400.0, -200.0, -100.0, 50.0, 100.0, 150.0, 200.0, 250.0, 300.0, 350.0, 400.0, 400.0, 450.0, 500.0, 550.0, 600.0}  [400], {-1200.0, -1000.0, -800.0, -650.0, -500.0, -350.0, -200.0, -50.0, 100.0, 150.0, 200.0, 250.0, 300.0, 350.0, 400.0, 400.0, 450.0, 500.0, 550.0, 600.0, 650.0, 700.0, 850.0, 1000.0, 1150.0}  [600], {-1800.0, -1400.0, -1000.0, -600.0, -400.0, -200.0, 100.0, 200.0, 250.0, 300.0, 350.0, 400.0, 400.0, 450.0, 500.0, 550.0, 600.0, 700.0, 800.0, 1000.0, 1200.0, 1400.0, 1800.0}  [1000], {-2000.0, -1400.0, -800.0, -500.0, -200.0, 100.0, 250.0, 350.0, 400.0, 400.0, 450.0, 550.0, 700.0, 1000.0, 1300.0, 1600.0, 2200.0} |
| A_6_-DNA^m1A16^  G10-C8  pH 5.4 (D_2_O)  25 °C  (*R*_1ρ_) | [150, 200, 250, 300, 400, 500, 600, 700, 900, 1000, 1200, 1400, 1600, 2000, 2500] {0}  [200], {-600.0, -400.0, -300.0, -200.0, -150.0, -100.0, -50.0, 50.0, 100.0, 150.0, 200.0, 200.0, 250.0, 300.0, 350.0, 400.0, 450.0, 500.0, 550.0, 600.0}  [400], {-1200.0, -1000.0, -850.0, -700.0, -550.0, -400.0, -250.0, -100.0, -50.0, 50.0, 100.0, 150.0, 200.0, 200.0, 250.0, 300.0, 350.0, 400.0, 450.0, 500.0, 650.0, 800.0, 950.0, 1100.0}  [600], {-1600.0, -1200.0, -800.0, -600.0, -400.0, -200.0, -100.0, 50.0, 100.0, 150.0, 200.0, 200.0, 250.0, 300.0, 350.0, 400.0, 500.0, 600.0, 800.0, 1000.0, 1200.0, 1600.0}  [1000], {-2200.0, -1600.0, -1000.0, -700.0, -400.0, -100.0, 50.0, 150.0, 200.0, 200.0, 250.0, 350.0, 500.0, 800.0, 1100.0, 1400.0, 2000.0} |
| A_6_-DNA^m1A16^  G10-C1'  pH 5.4 (D_2_O)  25 °C  (*R*_1ρ_) | [150, 200, 250, 300, 400, 500, 600, 700, 900, 1000, 1200, 1400, 1600, 2000, 2500] {0}  [200], {-600.0, -400.0, -300.0, -200.0, -150.0, -100.0, -50.0, 50.0, 100.0, 150.0, 200.0, 200.0, 250.0, 300.0, 350.0, 400.0, 450.0, 500.0, 550.0, 600.0}  [400], {-1200.0, -1000.0, -850.0, -700.0, -550.0, -400.0, -250.0, -100.0, -50.0, 50.0, 100.0, 150.0, 200.0, 200.0, 250.0, 300.0, 350.0, 400.0, 450.0, 500.0, 650.0, 800.0, 950.0, 1100.0}  [600], {-1600.0, -1200.0, -800.0, -600.0, -400.0, -200.0, -100.0, 50.0, 100.0, 150.0, 200.0, 200.0, 250.0, 300.0, 350.0, 400.0, 500.0, 600.0, 800.0, 1000.0, 1200.0, 1600.0}  [1000], {-2200.0, -1600.0, -1000.0, -700.0, -400.0, -100.0, 50.0, 150.0, 200.0, 200.0, 250.0, 350.0, 500.0, 800.0, 1100.0, 1400.0, 2000.0} |
| A_6_-DNA^m1G10^  A16-C8  pH 5.4 (D_2_O)  25 °C  (*R*_1ρ_) | [150, 200, 250, 300, 400, 500, 600, 700, 900, 1000, 1200, 1400, 1600, 2000, 2500] {0}  [200], {-600.0, -300.0, -200.0, -150.0, -100.0, -50.0, 50.0, 100.0, 150.0, 200.0, 200.0, 250.0, 300.0, 350.0, 400.0, 450.0, 500.0, 550.0, 600.0}  [400], {-1200.0, -1000.0, -850.0, -700.0, -550.0, -400.0, -250.0, -100.0, -50.0, 50.0, 100.0, 150.0, 200.0, 200.0, 250.0, 300.0, 350.0, 400.0, 450.0, 500.0, 650.0, 800.0, 950.0, 1100.0}  [600], {-1600.0, -1200.0, -800.0, -600.0, -400.0, -200.0, -100.0, 50.0, 100.0, 150.0, 200.0, 200.0, 250.0, 300.0, 350.0, 400.0, 500.0, 600.0, 800.0, 1000.0, 1200.0, 1600.0}  [1000], {-2200.0, -1600.0, -1000.0, -700.0, -400.0, -100.0, 50.0, 150.0, 200.0, 200.0, 250.0, 350.0, 500.0, 800.0, 1100.0, 1400.0, 2000.0} |
| A_6_-DNA^m1G10^  A16-C1'  pH 5.4 (D_2_O)  25 °C  (*R*_1ρ_) | [150, 200, 250, 300, 400, 500, 600, 700, 900, 1000, 1200, 1400, 1600, 2000, 2500] {0}  [200], {-600.0, -400.0, -300.0, -200.0, -150.0, -100.0, -50.0, 50.0, 100.0, 150.0, 200.0, 200.0, 250.0, 300.0, 350.0, 400.0, 450.0, 500.0, 550.0, 600.0}  [400], {-1200.0, -1000.0, -850.0, -700.0, -550.0, -400.0, -250.0, -100.0, -50.0, 50.0, 100.0, 150.0, 200.0, 200.0, 250.0, 300.0, 350.0, 400.0, 450.0, 500.0, 650.0, 800.0, 950.0, 1100.0}  [600], {-1600.0, -1200.0, -800.0, -600.0, -400.0, -200.0, -100.0, 50.0, 100.0, 150.0, 200.0, 200.0, 250.0, 300.0, 350.0, 400.0, 500.0, 600.0, 800.0, 1000.0, 1200.0, 1600.0}  [1000], {-2200.0, -1600.0, -1000.0, -700.0, -400.0, -100.0, 50.0, 150.0, 200.0, 200.0, 250.0, 350.0, 500.0, 800.0, 1100.0, 1400.0, 2000.0} |
| A_6_-DNA  T8-N3  pH 5.4 10 °C  (*R*_1ρ_) | [100, 150, 200, 250, 300, 400, 500, 600, 700, 900, 1000, 1200, 1400, 1600, 2000, 2500] {0} [100, 150, 200, 250, 300, 400, 500, 600, 700, 900, 1000, 1200, 1400, 1600, 2000, 2500] {0}  [100], {-290.0, -260.0, -230.0, -200.0, -200.0, -170.0, -140.0, -110.0, -80.0, -50.0, 50.0, 100.0, 150.0, 200.0, 250.0, 300.0}  [200], {-600.0, -550.0, -500.0, -450.0, -400.0, -350.0, -300.0, -250.0, -200.0, -200.0, -150.0, -100.0, -50.0, 50.0, 100.0, 150.0, 200.0, 300.0, 400.0, 600.0}  [300], {-800.0, -700.0, -600.0, -550.0, -500.0, -450.0, -400.0, -350.0, -300.0, -250.0, -200.0, -200.0, -150.0, -100.0, -50.0, 50.0, 100.0, 150.0, 200.0, 300.0, 400.0, 600.0, 800.0}  [400], {-1100.0, -950.0, -800.0, -650.0, -500.0, -450.0, -400.0, -350.0, -300.0, -250.0, -200.0, -200.0, -150.0, -100.0, -50.0, 50.0, 100.0, 250.0, 400.0, 550.0, 700.0, 850.0, 1000.0, 1200.0}  [600], {-1600.0, -1200.0, -1000.0, -800.0, -600.0, -500.0, -400.0, -350.0, -300.0, -250.0, -200.0, -200.0, -150.0, -100.0, -50.0, 100.0, 200.0, 400.0, 600.0, 800.0, 1200.0, 1600.0}  [1000], {-2600.0, -2000.0, -1400.0, -1100.0, -800.0, -500.0, -350.0, -250.0, -200.0, -200.0, -150.0, -50.0, 100.0, 400.0, 700.0, 1000.0, 1600.0, 2200.0, 2800.0} |
| A_6_-DNA^m1A16^  T8-N3  pH 5.4 10 °C  (*R*_1ρ_) | [100, 150, 200, 250, 300, 400, 500, 600, 700, 900, 1000, 1200, 1400, 1600, 2000, 2500] {0}  [100], {-290.0, -260.0, -230.0, -200.0, -200.0, -170.0, -140.0, -110.0, -80.0, -50.0, 50.0, 100.0, 150.0, 200.0, 250.0, 300.0}  [200], {-600.0, -550.0, -500.0, -450.0, -400.0, -350.0, -300.0, -250.0, -200.0, -200.0, -150.0, -100.0, -50.0, 50.0, 100.0, 150.0, 200.0, 300.0, 400.0, 600.0}  [300], {-800.0, -700.0, -600.0, -550.0, -500.0, -450.0, -400.0, -350.0, -300.0, -250.0, -200.0, -200.0, -150.0, -100.0, -50.0, 50.0, 100.0, 150.0, 200.0, 300.0, 400.0, 600.0, 800.0}  [400], {-800.0, -650.0, -500.0, -450.0, -400.0, -350.0, -300.0, -250.0, -200.0, -200.0, -150.0, -100.0, -50.0, 50.0, 100.0, 250.0, 400.0, 550.0, 700.0, 850.0, 1000.0, 1200.0} |
| **G-C^+^ Hoogsteen: Testing delta-melt** | |
| scaf2_CGT^GC^  G6-C1'  pH 5.4  30 °C  (*R*_1ρ_) | [150, 200, 250, 300, 400, 500, 600, 700, 900, 1000, 1200, 1400, 1600, 2000, 2500, 3000] {0}  [150], {-400.0, -150.0, 100.0, 200.0, 300.0, 350.0, 400.0, 450.0, 500.0, 520.0, 540.0, 560.0, 580.0, 600.0, 620.0, 640.0, 660.0, 680.0, 700.0, 750.0, 800.0, 850.0, 900.0, 1000.0, 1100.0, 1200.0, 1350.0}  [200], {-400.0, -200.0, 100.0, 200.0, 300.0, 350.0, 400.0, 450.0, 500.0, 520.0, 540.0, 560.0, 580.0, 600.0, 620.0, 640.0, 660.0, 680.0, 700.0, 750.0, 800.0, 850.0, 900.0, 1000.0, 1100.0, 1200.0, 1400.0, 1600.0}  [300], {-400.0, -200.0, 100.0, 200.0, 300.0, 350.0, 400.0, 450.0, 500.0, 520.0, 540.0, 560.0, 580.0, 600.0, 620.0, 640.0, 660.0, 680.0, 700.0, 750.0, 800.0, 850.0, 900.0, 1000.0, 1100.0, 1200.0, 1600.0}  [400], {-1400.0, -1000.0, -600.0, -400.0, -200.0, 150.0, 300.0, 400.0, 450.0, 500.0, 520.0, 540.0, 560.0, 580.0, 600.0, 620.0, 640.0, 660.0, 680.0, 700.0, 750.0, 800.0, 900.0, 1050.0, 1200.0, 1400.0, 1600.0, 1800.0, 2200.0, 2600.0}  [600], {-2400.0, -2000.0, -1600.0, -1200.0, -800.0, -400.0, -200.0, 200.0, 300.0, 400.0, 500.0, 530.0, 560.0, 580.0, 600.0, 620.0, 640.0, 670.0, 700.0, 800.0, 900.0, 1000.0, 1200.0, 1400.0, 1600.0, 2000.0, 2400.0, 2800.0} |
| scaf2_CGT^GC^  G6-C8  pH 5.4  30 °C  (*R*_1ρ_) | [100, 150, 200, 250, 300, 400, 500, 600, 700, 900, 1000, 1200, 1400, 1600, 2000, 2500, 3000] {0}  [100], {-400.0, -150.0, 100.0, 200.0, 300.0, 350.0, 400.0, 450.0, 500.0, 520.0, 540.0, 560.0, 580.0, 600.0, 620.0, 640.0, 660.0, 680.0, 700.0, 750.0, 800.0, 850.0, 900.0, 1000.0}  [150], {-400.0, -150.0, 100.0, 200.0, 300.0, 350.0, 400.0, 450.0, 500.0, 520.0, 540.0, 560.0, 580.0, 600.0, 620.0, 640.0, 660.0, 680.0, 700.0, 750.0, 800.0, 850.0, 900.0, 1000.0, 1100.0, 1200.0, 1350.0}  [200], {-400.0, -200.0, 100.0, 200.0, 300.0, 350.0, 400.0, 450.0, 500.0, 520.0, 540.0, 560.0, 580.0, 600.0, 620.0, 640.0, 660.0, 680.0, 700.0, 750.0, 800.0, 850.0, 900.0, 1000.0, 1100.0, 1200.0, 1400.0, 1600.0}  [300], {-400.0, -200.0, 100.0, 200.0, 300.0, 350.0, 400.0, 450.0, 500.0, 520.0, 540.0, 560.0, 580.0, 600.0, 620.0, 640.0, 660.0, 680.0, 700.0, 750.0, 800.0, 850.0, 900.0, 1000.0, 1100.0, 1200.0, 1400.0, 1600.0}  [600], {-2000.0, -1600.0, -1200.0, -800.0, -400.0, -200.0, 200.0, 300.0, 400.0, 500.0, 530.0, 560.0, 580.0, 600.0, 620.0, 640.0, 670.0, 700.0, 800.0, 900.0, 1000.0, 1200.0, 1400.0, 1600.0, 2000.0, 2400.0, 2800.0, 3200.0} |
| scaf2_TGC^GC^  G6-C1'  pH 5.4  30 °C  (*R*_1ρ_) | [200, 250, 300, 400, 500, 600, 700, 900, 1000, 1200, 1400, 1600, 2000, 2500, 3000] {0}  [200], {-600.0, -400.0, -200.0, -100.0, 100.0, 150.0, 200.0, 250.0, 300.0, 320.0, 340.0, 360.0, 380.0, 400.0, 420.0, 440.0, 460.0, 480.0, 500.0, 550.0, 600.0, 650.0, 700.0}  [400], {-1200.0, -800.0, -600.0, -400.0, -200.0, -50.0, 100.0, 200.0, 250.0, 300.0, 320.0, 340.0, 360.0, 380.0, 400.0, 420.0, 440.0, 460.0, 480.0, 500.0, 550.0, 600.0, 700.0, 850.0, 1000.0, 1200.0, 1400.0}  [600], {-1800.0, -1400.0, -1000.0, -600.0, -400.0, -200.0, 100.0, 200.0, 300.0, 330.0, 360.0, 380.0, 400.0, 420.0, 440.0, 470.0, 500.0, 600.0, 700.0, 800.0, 1000.0, 1200.0, 1800.0}  [1000], {-2600.0, -2000.0, -1400.0, -800.0, -500.0, -200.0, 100.0, 250.0, 350.0, 400.0, 450.0, 550.0, 700.0, 1000.0, 1300.0, 1600.0, 2200.0, 2800.0, 3400.0} |
| scaf2_TGC^GC^  G6-C8  pH 5.4  30 °C  (*R*_1ρ_) | [100, 150, 200, 250, 300, 400, 500, 600, 700, 900, 1000, 1200, 1400, 1600, 2000, 2500, 3000] {0}  [200], {-600.0, -400.0, -200.0, -100.0, 100.0, 150.0, 200.0, 250.0, 300.0, 320.0, 360.0, 380.0, 400.0, 420.0, 440.0, 460.0, 480.0, 500.0, 550.0, 600.0, 650.0, 700.0}  [400], {-800.0, -600.0, -400.0, -200.0, -50.0, 100.0, 200.0, 250.0, 300.0, 320.0, 340.0, 360.0, 380.0, 400.0, 420.0, 440.0, 460.0, 480.0, 500.0, 550.0, 600.0, 700.0, 850.0, 1000.0}  [600], {-1000.0, -600.0, -400.0, -200.0, 100.0, 200.0, 300.0, 330.0, 360.0, 380.0, 400.0, 420.0, 440.0, 470.0, 500.0, 600.0, 700.0, 800.0, 1000.0, 1200.0, 1400.0, 1800.0}  [1000], {-3100.0, -2600.0, -2000.0, -1400.0, -800.0, -500.0, -200.0, 100.0, 250.0, 350.0, 400.0, 450.0, 550.0, 700.0, 1000.0, 1300.0, 1600.0, 2200.0, 2800.0, 3400.0} |
| scaf2_TGC^GC^  G6-C8  pH 5.4HS  15 °C  (*R*_1ρ_) | [150, 200, 250, 300, 400, 500, 600, 700, 900, 1000, 1200, 1400, 1600, 2000, 2500, 3000] {0}  [100], {-400.0, -300.0, -250.0, -200.0, -150.0, -100.0, -60.0, -30.0, 30.0, 60.0, 100.0, 150.0, 200.0, 250.0, 300.0, 400.0}  [200], {-800.0, -600.0, -500.0, -400.0, -300.0, -250.0, -200.0, -150.0, -100.0, -60.0, -30.0, 30.0, 60.0, 100.0, 150.0, 200.0, 250.0, 300.0, 400.0, 500.0, 600.0, 800.0}  [400], {-1600.0, -1200.0, -1000.0, -800.0, -600.0, -450.0, -300.0, -200.0, -150.0, -100.0, -60.0, -30.0, 30.0, 60.0, 100.0, 150.0, 200.0, 300.0, 450.0, 600.0, 800.0, 1000.0, 1200.0, 1600.0}  [1000], {-3500.0, -3000.0, -2400.0, -1800.0, -1200.0, -900.0, -600.0, -300.0, -150.0, -50.0, 50.0, 150.0, 300.0, 600.0, 900.0, 1200.0, 1800.0, 2400.0, 3000.0, 3500.0} |
| scaf2_TGC^GC^  G6-C8  pH 5.4HS  30 °C  (*R*_1ρ_) | [100, 150, 200, 250, 300, 400, 500, 600, 700, 900, 1000, 1200, 1600, 2000, 2500, 3000] {0}  [100], {-350.0, -300.0, -250.0, -200.0, -150.0, -120.0, -90.0, -60.0, -30.0, 30.0, 60.0, 90.0, 120.0, 150.0, 200.0, 250.0, 300.0}  [200], {-600.0, -500.0, -400.0, -350.0, -300.0, -250.0, -200.0, -150.0, -100.0, -60.0, -30.0, 30.0, 60.0, 100.0, 150.0, 200.0, 250.0, 300.0, 350.0, 400.0, 500.0, 600.0}  [400], {-1200.0, -1000.0, -800.0, -600.0, -500.0, -400.0, -300.0, -200.0, -150.0, -100.0, -50.0, 50.0, 100.0, 150.0, 200.0, 300.0, 400.0, 500.0, 600.0, 800.0, 1000.0, 1200.0}  [1000], {-3500.0, -3000.0, -2400.0, -1800.0, -1200.0, -900.0, -600.0, -300.0, -150.0, -50.0, 50.0, 150.0, 300.0, 600.0, 900.0, 1200.0, 1800.0, 2400.0, 3000.0, 3500.0} |
| scaf2_TGC^GC^  G6-C1'  pH 5.4HS  30 °C  (*R*_1ρ_) | [150, 200, 250, 300, 400, 500, 600, 700, 900, 1000, 1200, 1400, 1600, 2000, 2500, 3000] {0}  [150], {-500.0, -450.0, -400.0, -350.0, -300.0, -250.0, -200.0, -150.0, -100.0, -60.0, -30.0, 30.0, 60.0, 100.0, 150.0, 200.0, 250.0, 300.0, 350.0, 400.0, 450.0, 500.0}  [200], {-600.0, -500.0, -400.0, -350.0, -300.0, -250.0, -200.0, -150.0, -100.0, -60.0, -30.0, 30.0, 60.0, 100.0, 150.0, 200.0, 250.0, 300.0, 350.0, 400.0, 500.0, 600.0}  [400], {-1200.0, -1000.0, -800.0, -600.0, -500.0, -400.0, -300.0, -200.0, -150.0, -100.0, -50.0, 50.0, 100.0, 150.0, 200.0, 300.0, 400.0, 500.0, 600.0, 800.0, 1000.0, 1200.0}  [1000], {-3500.0, -3000.0, -2400.0, -1800.0, -1200.0, -900.0, -600.0, -300.0, -150.0, -50.0, 50.0, 150.0, 300.0, 600.0, 900.0, 1200.0, 1800.0, 2400.0, 3000.0, 3500.0} |
| scaf2_TGC^GC^  G6-C8  pH 5.4HS  40 °C  (*R*_1ρ_) | [100, 150, 200, 250, 300, 400, 500, 600, 700, 900, 1000, 1200, 1400, 1600, 2000, 2500, 3000] {0}  [100], {-300.0, -200.0, -150.0, -100.0, -60.0, -30.0, 30.0, 60.0, 100.0, 150.0, 200.0, 250.0, 300.0}  [200], {-600.0, -500.0, -400.0, -300.0, -250.0, -200.0, -150.0, -100.0, -60.0, -30.0, 30.0, 60.0, 100.0, 150.0, 200.0, 250.0, 300.0, 350.0, 400.0, 450.0, 500.0, 520.0, 540.0, 560.0, 580.0, 600.0, 620.0, 640.0, 660.0, 680.0, 700.0}  [400], {-1200.0, -1000.0, -800.0, -600.0, -450.0, -300.0, -200.0, -150.0, -100.0, -60.0, -30.0, 30.0, 60.0, 100.0, 150.0, 200.0, 300.0, 400.0, 450.0, 500.0, 520.0, 540.0, 560.0, 580.0, 600.0, 620.0, 640.0, 660.0, 680.0, 700.0, 750.0, 800.0, 900.0, 1000.0, 1050.0, 1200.0, 1400.0}  [600], {-2000.0, -1600.0, -1200.0, -800.0, -400.0, -200.0, 200.0, 300.0, 400.0, 500.0, 530.0, 560.0, 580.0, 600.0, 620.0, 640.0, 670.0, 700.0, 800.0, 900.0, 1000.0, 1200.0, 1400.0, 1600.0, 2000.0}  [1000], {-3500.0, -3000.0, -2400.0, -1800.0, -1200.0, -900.0, -600.0, -300.0, -150.0, -50.0, 50.0, 150.0, 300.0, 450.0, 550.0, 600.0, 650.0, 750.0, 900.0, 1200.0, 1500.0, 1800.0, 2400.0, 3000.0, 3500.0} |
| scaf2_TGC^GC^  G6-C1'  pH 5.4HS  40 °C  (*R*_1ρ_) | [150, 200, 250, 300, 400, 500, 600, 700, 900, 1000, 1200, 1400, 1600, 2000, 2500, 3000] {0}  [200], {-500.0, -100.0, 100.0, 200.0, 250.0, 300.0, 400.0, 420.0, 440.0, 460.0, 480.0, 500.0, 540.0, 560.0, 580.0, 650.0, 700.0}  [400], {-1100.0, -700.0, -500.0, -300.0, -100.0, 50.0, 200.0, 300.0, 350.0, 400.0, 420.0, 440.0, 460.0, 480.0, 500.0, 520.0, 540.0, 560.0, 580.0, 600.0, 650.0, 700.0, 800.0, 950.0, 1100.0, 1300.0}  [600], {-2100.0, -1700.0, -1300.0, -900.0, -500.0, -300.0, -100.0, 100.0, 200.0, 300.0, 400.0, 430.0, 460.0, 480.0, 500.0, 520.0, 540.0, 570.0, 600.0, 700.0, 800.0, 900.0, 1100.0, 1300.0, 1500.0, 1900.0}  [1000], {-3500.0, -3000.0, -2500.0, -1900.0, -1300.0, -700.0, -400.0, -100.0, 200.0, 350.0, 450.0, 500.0, 550.0, 650.0, 800.0, 1100.0, 1400.0, 1700.0, 2300.0, 3500.0} |
| scaf2_CGT^GC^  G6-C8  pH 5.4HS  30 °C  (CEST) | [10.0], {-1500.0, -1442.1, -1384.2, -1326.3, -1268.4, -1210.5, -1152.6, -1094.7, -1036.8, -978.9, -921.1, -863.2, -805.3, -747.4, -689.5, -631.6, -573.7, -515.8, -457.9, -400.0, -400.0, -383.5, -367.1, -350.6, -334.2, -317.7, -301.3, -284.8, -268.4, -251.9, -235.4, -219.0, -202.5, -186.1, -169.6, -153.2, -136.7, -120.3, -103.8, -87.3, -70.9, -54.4, -38.0, -21.5, -5.1, 11.4, 27.8, 44.3, 60.8, 77.2, 93.7, 110.1, 126.6, 143.0, 159.5, 175.9, 192.4, 208.9, 225.3, 241.8, 258.2, 274.7, 291.1, 307.6, 324.1, 340.5, 357.0, 373.4, 389.9, 406.3, 422.8, 439.2, 455.7, 472.2, 488.6, 505.1, 521.5, 538.0, 554.4, 570.9, 587.3, 603.8, 620.3, 636.7, 653.2, 669.6, 686.1, 702.5, 719.0, 735.4, 751.9, 768.4, 784.8, 800.0, 801.3, 817.7, 834.2, 836.8, 850.6, 867.1, 873.7, 883.5, 900.0, 910.5, 947.4, 984.2, 1021.1, 1057.9, 1094.7, 1131.6, 1168.4, 1205.3, 1242.1, 1278.9, 1315.8, 1352.6, 1389.5, 1426.3, 1463.2, 1500.0}  [20.0], {-1500.0, -1442.1, -1384.2, -1326.3, -1268.4, -1210.5, -1152.6, -1094.7, -1036.8, -978.9, -921.1, -863.2, -805.3, -747.4, -689.5, -631.6, -573.7, -515.8, -457.9, -400.0, -400.0, -383.5, -367.1, -350.6, -334.2, -317.7, -301.3, -284.8, -268.4, -251.9, -235.4, -219.0, -202.5, -186.1, -169.6, -153.2, -136.7, -120.3, -103.8, -87.3, -70.9, -54.4, -38.0, -21.5, -5.1, 11.4, 27.8, 44.3, 60.8, 77.2, 93.7, 110.1, 126.6, 143.0, 159.5, 175.9, 192.4, 208.9, 225.3, 241.8, 258.2, 274.7, 291.1, 307.6, 324.1, 340.5, 357.0, 373.4, 389.9, 406.3, 422.8, 439.2, 455.7, 472.2, 488.6, 505.1, 521.5, 538.0, 554.4, 570.9, 587.3, 603.8, 620.3, 636.7, 653.2, 669.6, 686.1, 702.5, 719.0, 735.4, 751.9, 768.4, 784.8, 800.0, 801.3, 817.7, 834.2, 836.8, 850.6, 867.1, 873.7, 883.5, 900.0, 910.5, 947.4, 984.2, 1021.1, 1057.9, 1094.7, 1131.6, 1168.4, 1205.3, 1242.1, 1278.9, 1315.8, 1352.6, 1389.5, 1426.3, 1463.2, 1500.0}  [50.0], {-1500.0, -1442.1, -1384.2, -1326.3, -1268.4, -1210.5, -1152.6, -1094.7, -1036.8, -978.9, -921.1, -863.2, -805.3, -747.4, -689.5, -631.6, -573.7, -515.8, -457.9, -400.0, -400.0, -383.5, -367.1, -350.6, -334.2, -317.7, -301.3, -284.8, -268.4, -251.9, -235.4, -219.0, -202.5, -186.1, -169.6, -153.2, -136.7, -120.3, -103.8, -87.3, -70.9, -54.4, -38.0, -21.5, -5.1, 11.4, 27.8, 44.3, 60.8, 77.2, 93.7, 110.1, 126.6, 143.0, 159.5, 175.9, 192.4, 208.9, 225.3, 241.8, 258.2, 274.7, 291.1, 307.6, 324.1, 340.5, 357.0, 373.4, 389.9, 406.3, 422.8, 439.2, 455.7, 472.2, 488.6, 505.1, 521.5, 538.0, 554.4, 570.9, 587.3, 603.8, 620.3, 636.7, 653.2, 669.6, 686.1, 702.5, 719.0, 735.4, 751.9, 768.4, 784.8, 800.0, 801.3, 817.7, 834.2, 836.8, 850.6, 867.1, 873.7, 883.5, 900.0, 910.5, 947.4, 984.2, 1021.1, 1057.9, 1094.7, 1131.6, 1168.4, 1205.3, 1242.1, 1278.9, 1315.8, 1352.6, 1389.5, 1426.3, 1463.2, 1500.0}  [70.0], {-1500.0, -1442.1, -1384.2, -1326.3, -1268.4, -1210.5, -1152.6, -1094.7, -1036.8, -978.9, -921.1, -863.2, -805.3, -747.4, -689.5, -631.6, -573.7, -515.8, -457.9, -400.0, -400.0, -383.5, -367.1, -350.6, -334.2, -317.7, -301.3, -284.8, -268.4, -251.9, -235.4, -219.0, -202.5, -186.1, -169.6, -153.2, -136.7, -120.3, -103.8, -87.3, -70.9, -54.4, -38.0, -21.5, -5.1, 11.4, 27.8, 44.3, 60.8, 77.2, 93.7, 110.1, 126.6, 143.0, 159.5, 175.9, 192.4, 208.9, 225.3, 241.8, 258.2, 274.7, 291.1, 307.6, 324.1, 340.5, 357.0, 373.4, 389.9, 406.3, 422.8, 439.2, 455.7, 472.2, 488.6, 505.1, 521.5, 538.0, 554.4, 570.9, 587.3, 603.8, 620.3, 636.7, 653.2, 669.6, 686.1, 702.5, 719.0, 735.4, 751.9, 768.4, 784.8, 800.0, 801.3, 817.7, 834.2, 836.8, 850.6, 867.1, 873.7, 883.5, 900.0, 910.5, 947.4, 984.2, 1021.1, 1057.9, 1094.7, 1131.6, 1168.4, 1205.3, 1242.1, 1278.9, 1315.8, 1352.6, 1389.5, 1426.3, 1463.2, 1500.0} |
| scaf2_CGT^GC^  G6-C1'  pH 5.4HS  30 °C  (CEST) | [10.0], {-1500.0, -1442.1, -1384.2, -1326.3, -1268.4, -1210.5, -1152.6, -1094.7, -1036.8, -978.9, -921.1, -863.2, -805.3, -747.4, -689.5, -631.6, -573.7, -515.8, -457.9, -400.0, -400.0, -383.5, -367.1, -350.6, -334.2, -317.7, -301.3, -284.8, -268.4, -251.9, -235.4, -219.0, -202.5, -186.1, -169.6, -153.2, -136.7, -120.3, -103.8, -87.3, -70.9, -54.4, -38.0, -21.5, -5.1, 11.4, 27.8, 44.3, 60.8, 77.2, 93.7, 110.1, 126.6, 143.0, 159.5, 175.9, 192.4, 208.9, 225.3, 241.8, 258.2, 274.7, 291.1, 307.6, 324.1, 340.5, 357.0, 373.4, 389.9, 406.3, 422.8, 439.2, 455.7, 472.2, 488.6, 505.1, 521.5, 538.0, 554.4, 570.9, 587.3, 603.8, 620.3, 636.7, 653.2, 669.6, 686.1, 702.5, 719.0, 735.4, 751.9, 768.4, 784.8, 800.0, 801.3, 817.7, 834.2, 836.8, 850.6, 867.1, 873.7, 883.5, 900.0, 910.5, 947.4, 984.2, 1021.1, 1057.9, 1094.7, 1131.6, 1168.4, 1205.3, 1242.1, 1278.9, 1315.8, 1352.6, 1389.5, 1426.3, 1463.2, 1500.0}  [20.0], {-1500.0, -1442.1, -1384.2, -1326.3, -1268.4, -1210.5, -1152.6, -1094.7, -1036.8, -978.9, -921.1, -863.2, -805.3, -747.4, -689.5, -631.6, -573.7, -515.8, -457.9, -400.0, -400.0, -383.5, -367.1, -350.6, -334.2, -317.7, -301.3, -284.8, -268.4, -251.9, -235.4, -219.0, -202.5, -186.1, -169.6, -153.2, -136.7, -120.3, -103.8, -87.3, -70.9, -54.4, -38.0, -21.5, -5.1, 11.4, 27.8, 44.3, 60.8, 77.2, 93.7, 110.1, 126.6, 143.0, 159.5, 175.9, 192.4, 208.9, 225.3, 241.8, 258.2, 274.7, 291.1, 307.6, 324.1, 340.5, 357.0, 373.4, 389.9, 406.3, 422.8, 439.2, 455.7, 472.2, 488.6, 505.1, 521.5, 538.0, 554.4, 570.9, 587.3, 603.8, 620.3, 636.7, 653.2, 669.6, 686.1, 702.5, 719.0, 735.4, 751.9, 768.4, 784.8, 800.0, 801.3, 817.7, 834.2, 836.8, 850.6, 867.1, 873.7, 883.5, 900.0, 910.5, 947.4, 984.2, 1021.1, 1057.9, 1094.7, 1131.6, 1168.4, 1205.3, 1242.1, 1278.9, 1315.8, 1352.6, 1389.5, 1426.3, 1463.2, 1500.0}  [50.0], {-1500.0, -1442.1, -1384.2, -1326.3, -1268.4, -1210.5, -1152.6, -1094.7, -1036.8, -978.9, -921.1, -863.2, -805.3, -747.4, -689.5, -631.6, -573.7, -515.8, -457.9, -400.0, -400.0, -383.5, -367.1, -350.6, -334.2, -317.7, -301.3, -284.8, -268.4, -251.9, -235.4, -219.0, -202.5, -186.1, -169.6, -153.2, -136.7, -120.3, -103.8, -87.3, -70.9, -54.4, -38.0, -21.5, -5.1, 11.4, 27.8, 44.3, 60.8, 77.2, 93.7, 110.1, 126.6, 143.0, 159.5, 175.9, 192.4, 208.9, 225.3, 241.8, 258.2, 274.7, 291.1, 307.6, 324.1, 340.5, 357.0, 373.4, 389.9, 406.3, 422.8, 439.2, 455.7, 472.2, 488.6, 505.1, 521.5, 538.0, 554.4, 570.9, 587.3, 603.8, 620.3, 636.7, 653.2, 669.6, 686.1, 702.5, 719.0, 735.4, 751.9, 768.4, 784.8, 800.0, 801.3, 817.7, 834.2, 836.8, 850.6, 867.1, 873.7, 883.5, 900.0, 910.5, 947.4, 984.2, 1021.1, 1057.9, 1094.7, 1131.6, 1168.4, 1205.3, 1242.1, 1278.9, 1315.8, 1352.6, 1389.5, 1426.3, 1463.2, 1500.0}  [70.0], {-1500.0, -1442.1, -1384.2, -1326.3, -1268.4, -1210.5, -1152.6, -1094.7, -1036.8, -978.9, -921.1, -863.2, -805.3, -747.4, -689.5, -631.6, -573.7, -515.8, -457.9, -400.0, -400.0, -383.5, -367.1, -350.6, -334.2, -317.7, -301.3, -284.8, -268.4, -251.9, -235.4, -219.0, -202.5, -186.1, -169.6, -153.2, -136.7, -120.3, -103.8, -87.3, -70.9, -54.4, -38.0, -21.5, -5.1, 11.4, 27.8, 44.3, 60.8, 77.2, 93.7, 110.1, 126.6, 143.0, 159.5, 175.9, 192.4, 208.9, 225.3, 241.8, 258.2, 274.7, 291.1, 307.6, 324.1, 340.5, 357.0, 373.4, 389.9, 406.3, 422.8, 439.2, 455.7, 472.2, 488.6, 505.1, 521.5, 538.0, 554.4, 570.9, 587.3, 603.8, 620.3, 636.7, 653.2, 669.6, 686.1, 702.5, 719.0, 735.4, 751.9, 768.4, 784.8, 800.0, 801.3, 817.7, 834.2, 836.8, 850.6, 867.1, 873.7, 883.5, 900.0, 910.5, 947.4, 984.2, 1021.1, 1057.9, 1094.7, 1131.6, 1168.4, 1205.3, 1242.1, 1278.9, 1315.8, 1352.6, 1389.5, 1426.3, 1463.2, 1500.0} |

Ω = ω_RF_ - ω_OBS_, where ω_OBS_ is the Larmor frequency of the observed resonance and ω_RF_ is the angular frequency of the RF field. Secondary structures of the constructs are given in Fig. S4. rA, m^1^G, rG, m^6^A, m^1^A denote ribo adenosine, *N*^1^-methyl guanine, ribo guanine, *N*^6^-methyl adenine, *N*^1^-methyl adenosine, respectively.

**Table S5.** Exchange parameters obtained from fitting *R*_1ρ_/CEST relaxation dispersion data

| **Nucleus** | ***p*_i_ (%)** | ***k*_ex_ (s^-1^)** | **Δω_i-Major_ (ppm)** | ***R*_1_ (s^-1^)** | ***R*_2_ (s^-1^)** | **Red. χ^2^** |
| --- | --- | --- | --- | --- | --- | --- |
| **A-T Hoogsteen** | | | | | | |
| A_6_-DNA A16-C8  pH 4.4HS 25 °C (*R*_1ρ_) | 0.888±0.043 | 3709±218 | 3.35±0.11 | 0.73±0.16 | 26.23±0.47 | 0.19 |
| A_6_-DNA A21-C1'  pH 4.4HS 30 °C (*R*_1ρ_) | 0.369±0.026 | 2813±218 | 2.92±0.16 | 1.97±0.05 | 14.52±0.18 | 0.14 |
| A_6_-DNA A16-C8  pH 5.4 17.5 °C (*R*_1ρ_) | 0.463±0.019 | 1515±120 | 2.83±0.10 | 2.66±0.07 | 28.83±0.19 | 0.24 |
| A_6_-DNA A16-C8  pH 5.4 20 °C (*R*_1ρ_) | 0.459±0.030 | 2025±171 | 2.89±0.13 | 2.30±0.08 | 26.49±0.19 | 0.33 |
| A_6_-DNA A16-C8  pH 5.4 22.5 °C (*R*_1ρ_) | 0.559±0.047 | 3068±219 | 2.77±0.13 | 2.96±0.07 | 24.64±0.22 | 0.22 |
| A_6_-DNA A16-C8  pH 5.4 25 °C (*R*_1ρ_) | 0.629±0.081 | 3652±302 | 2.62±0.17 | 2.98±0.08 | 23.33±0.28 | 0.30 |
| A_6_-DNA^rA16^  rA16-C8  pH 5.4 10 °C  (*R*_1ρ_) | 0.180±0.008 | 2547±213 | 3.79±0.14 | 1.64±0.03 | 42.08±0.13 | 0.43 |
| A_6_-DNA^rA16^  rA16-C8  pH 5.4 12.5 °C (*R*_1ρ_) | 0.286±0.046 | 4398±438 | 2.96±0.28 | 1.65±0.04 | 37.25±0.17 | 0.37 |
| A_6_-DNA^rA16^  rA16-C8  pH 5.4 15 °C (*R*_1ρ_) | 0.311±0.074 | 5173±577 | 2.91±0.37 | 1.84±0.04 | 33.59±0.21 | 0.54 |
| A_6_-DNA  A16-C8,C1' pH 5.4HS 25 °C (*R*_1ρ_) | 0.498±0.038 | 3430±170 | 2.44±0.11  2.88±0.12 | 2.28±0.08  1.93±0.06 | 30.26±0.17  19.99±0.16 | 0.17 |
| A_6_-DNA^m1G10^  A16-C8,C1'  pH 5.4HS 25 °C (*R*_1ρ_) | 2.460±0.041 | 2357±37 | 2.48±0.03  3.00±0.03 | 2.51±0.12  1.27±0.11 | 25.49±0.23  18.15±0.20 | 0.08 |
| A_2_-DNA A16-C8  pH 6.8 25 °C (*R*_1ρ_) | 0.252±0.086 | 3169±554 | 2.09±0.36 | 2.17±0.06 | 22.12±0.17 | 0.21 |
| A_6_-DNA A16-C8  pH 6.8 25 °C (*R*_1ρ_) | 0.441±0.058 | 3770±315 | 2.32±0.15 | 1.78±0.06 | 23.03±0.21 | 0.13 |
| A_6_-DNA A21-C1'  pH 6.8 25 °C (*R*_1ρ_) | 0.204±0.029 | 1099±263 | 2.86±0.20 | 1.67±0.07 | 14.44±0.12 | 0.20 |
| A_6_-DNA A21-C1'  pH 6.8 30 °C (*R*_1ρ_) | 0.205±0.019 | 1484±199 | 2.94±0.20 | 2.21±0.06 | 13.34±0.12 | 0.21 |
| **G-C^+^ Hoogsteen** | | | | | | |
| A_2_-DNA G10-C8  pH 4.4HS 25 °C (*R*_1ρ_) | 2.586±0.052 | 3003±91 | 3.26±0.04 | 0.56±0.21 | 30.87±0.56 | 0.17 |
| AcDNA  G7-C1',C8,N1  pH 5.3HS 25 °C (*R*_1ρ_) | 0.137±0.036 | 549±166 | 4.36±0.17  3.88±0.35  -2.39±0.26 | 1.23±0.04  1.35±0.06  2.03±0.02 | 21.17±0.06  30.42±0.08  8.52±0.03 | 0.79 |
| A_2_-DNA  G10-C8,C1'  pH 5.4 25 °C (*R*_1ρ_) | 1.014±0.024 | 2340±134 | 3.11±0.11  3.87±0.07 | 1.14±0.37  1.19±0.19 | 25.18±0.53  15.78±0.39 | 0.90 |
| A_6_-DNA  G10-C8,C1'  pH 5.4 25 °C (*R*_1ρ_) | 0.717±0.010 | 1766±44 | 2.97±0.03  3.66±0.05 | 1.98±0.05  1.87±0.06 | 20.85±0.09  15.12±0.12 | 0.11 |
| A_6_-DNA^rG10^  C15-C6  pH 5.4 25 °C (*R*_1ρ_) | 0.577±0.105 | 4949±526 | 2.44±0.28 | 1.88±0.06 | 29.71±0.37 | 0.42 |
| A_6_-DNA G10-C8,C1' pH 5.4HS  25 °C (*R*_1ρ_) | 0.251±0.012 | 1321±108 | 2.91±0.09  3.49±0.12 | 2.22±0.05  1.72±0.06 | 23.69±0.08  16.45±0.09 | 0.11 |
| A_6_-DNA G10-C1'  pH 6.8 25 °C (*R*_1ρ_) | 0.063±0.015 | 2069±1036 | 3.04±0.53 | 1.55±0.10 | 14.89±0.21 | 0.32 |
| ***N*^6^-methylamino rotation in m^6^A** | | | | | | |
| hpGGACU^m6A6^  m^6^A6-C2,C10  pH 6.8 37 °C (CEST) | 0.643±0.006 | 192±5 | 2.78±0.01  -1.56±0.01 | 3.30±0.01  1.24±0.01 | 18.38±0.04  4.55±0.05 | 25.91 |
| hpGGACU^m6A6^  m^6^A6-C2  pH 6.8 55 °C (*R*_1ρ_) | 1.205±0.042 | 534±24 | 2.63±0.02 | 4.45±0.02 | 13.71±0.04 | 0.23 |
| hpGGACU^m6A6^  m^6^A6-C2  pH 6.8 65 °C (*R*_1ρ_) | 1.526±0.013 | 1187±16 | 2.56±0.01 | 4.90±0.03 | 11.91±0.05 | 0.23 |
| A_6_-RNA^m6A16^  m^6^A16-C2  pH 6.8 37 °C (*R*_1ρ_) | 1.203±0.580 | 99±56 | 1.84±0.08 | 4.55±0.03 | 21.02±0.03 | 0.38 |
| **Hoogsteen Cooperativity** | | | | | | |
| A_6_-DNA A16-C8,C1' pH 5.4  (D_2_O) 25 °C (*R*_1ρ_) | 0.503±0.014 | 2909±117 | 2.72±0.06  3.29±0.07 | 1.48±0.08  1.41±0.07 | 27.86±0.16  18.17±0.16 | 0.11 |
| A_6_-DNA G10-C8,C1' pH 5.4  (D_2_O) 25 °C (*R*_1ρ_) | 0.459±0.032 | 794±75 | 3.04±0.05  3.57±0.07 | 1.37±0.06  1.19±0.06 | 27.53±0.09  18.22±0.08 | 0.15 |
| A_6_-DNA^m1A16^  G10-C8,C1'  pH 5.4 (D_2_O)  25 °C (*R*_1ρ_) | 0.552±0.017 | 2388±122 | 2.93±0.09  3.03±0.08 | 1.25±0.12  1.08±0.09 | 28.74±0.22  20.62±0.19 | 0.17 |
| A_6_-DNA^m1G10^  A16-C8, C1'  pH 5.4 (D_2_O)  25 °C (*R*_1ρ_) | 2.634±0.023 | 1950±33 | 2.59±0.02  3.16±0.02 | 1.00±0.16  0.75±0.14 | 30.43±0.25  20.02±0.22 | 0.14 |
| A_6_-DNA^m1A16^  T8-N3 pH 5.4  10 °C (*R*_1ρ_) | 2.539±0.619 | 282±76 | -1.27±0.08 | 1.47±0.03 | 7.98±0.04 | 0.14 |
| **G-C^+^ Hoogsteen: Testing delta-melt** | | | | | | |
| scaf2_CGT^GC^  G6-C8,C1'  pH 5.4 30 °C  (*R*_1ρ_) | 0.736±0.015 | 680±19 | 3.76±0.02  3.91±0.02 | 2.84±0.03  2.23±0.03 | 19.73±0.06  13.36±0.06 | 0.16 |
| scaf2_TGC^GC^  G6-C8,C1'  pH 5.4 30 °C  (*R*_1ρ_) | 0.196±0.005 | 2528±137 | 4.27±0.11  4.12±0.12 | 2.82±0.05  2.39±0.06 | 19.25±0.08  13.37±0.11 | 0.16 |
| scaf2_TGC^GC^  G6-C8,C1'  pH 5.4HS 30 °C  (*R*_1ρ_) | 0.069±0.005 | 2177±355 | 3.20±0.33  4.11±0.32 | 2.49±0.04  2.16±0.03 | 20.21±0.08  13.93±0.08 | 0.23 |
| scaf2_TGC^GC^  G6-C8,C1'  pH 5.4HS 40 °C  (*R*_1ρ_)* | 0.071±0.019 | 6571±1566 | 4.27  4.12 | 2.68±0.04  2.38±0.05 | 16.02±0.22  11.27±0.20 | 0.25 |
| scaf2_CGT^GC^  G6-C8,C1'  pH 5.4HS 30 °C  (CEST) | 0.203±0.002 | 490±12 | 3.78±0.01  3.93±0.02 | 2.68±0.01  2.27±0.01 | 19.16±0.03  12.43±0.07 | 13.63 |

*p*_i_ denotes the population of the i^th^ minor conformation, *k*_ex_ the exchange rate (sum of forward and backward rate constants) between the major and i^th^ minor conformation, Δω_i-Major_ = ω_i_ - ω_Major_, where ω_i_ and ω_Major_ are the Larmor frequencies of the spin in the i^th^ minor conformation and the major conformation, *R*_1_ and *R*_2_ are the longitudinal and transverse spin relaxation rate constants, respectively, and m^6^A denotes *N*^6^-methyl adenine. * denotes data for which Δω was fixed during fitting (Methods). Errors in the exchange parameters were obtained using a Monte-Carlo scheme as described in Methods.

**Table S6.** Summary of the relaxation delay times used in the imino proton exchange experiments

| **Nucleus** | **[NH_3_]**  **(M)** | **Delay (s)** |
| --- | --- | --- |
| GDNA  T6-H3  pH 8.8, 25 °C | 0 | 0.0, 0.05, 0.1, 0.15, 0.2, 0.25, 0.3, 0.4, 0.5, 0.6, 0.8, 1.0, 1.2, 1.5, 1.8, 2.1, 2.4, 3.0, 3.6, 4.2, 4.8 |
|  | 0.02 | 0.0, 0.005, 0.01, 0.02, 0.03, 0.04, 0.05, 0.1, 0.2, 0.3, 0.5, 0.7, 1.0, 1.3, 1.6, 2.0, 2.5, 3.0, 3.5 |
|  | 0.04 | 0.0, 0.002, 0.004, 0.006, 0.008, 0.01, 0.015, 0.02, 0.025, 0.03, 0.04, 0.05, 0.1, 0.2, 0.3, 0.5, 0.7, 1.0, 1.3 |
|  | 0.1 | 0.0, 0.001, 0.002, 0.003, 0.004, 0.005, 0.006, 0.008, 0.01, 0.015, 0.02, 0.03, 0.04, 0.05, 0.1, 0.2, 0.3 |
|  | 0.15 | 0.0, 0.001, 0.002, 0.003, 0.004, 0.005, 0.006, 0.008, 0.01, 0.02, 0.05 |
| A_6_-DNA  T5-H3  pH 8.8, 25 °C | 0 | 0.0, 0.05, 0.1, 0.15, 0.2, 0.25, 0.3, 0.5, 0.7, 1.3, 1.7, 2.0, 2.5, 3.0, 3.5 |
|  | 0.02 | 0.0, 0.01, 0.03, 0.05, 0.07, 0.1, 0.15, 0.2, 0.25, 0.3, 0.5, 0.7, 1.0, 1.5, 2.0, 2.5, 3.0 |
|  | 0.04 | 0.0, 0.01, 0.03, 0.05, 0.07, 0.1, 0.15, 0.2, 0.25, 0.3, 0.5, 0.7, 1.0, 1.5, 2.0, 2.5, 3.0 |
|  | 0.1 | 0.0, 0.01, 0.015, 0.02, 0.03, 0.05, 0.07, 0.1, 0.15, 0.2, 0.25, 0.3, 0.5, 0.7, 1.0 |
|  | 0.15 | 0.0, 0.002, 0.004, 0.006, 0.008, 0.01, 0.013, 0.016, 0.02, 0.025, 0.03, 0.05, 0.07, 0.1, 0.15, 0.2, 0.25, 0.3, 0.5, 0.7, 1.0 |
| A_6_-DNA  T6-H3  pH 8.8, 25 °C | 0 | 0, 0.05, 0.1, 0.15, 0.2, 0.25, 0.3, 0.5, 0.7, 1.3, 1.7, 2, 2.5, 3, 3.5 |
|  | 0.02 | 0, 0.01, 0.03, 0.05, 0.07, 0.1, 0.15, 0.2, 0.25, 0.3, 0.5, 0.7, 1.0, 1.5, 2.0, 2.5, 3.0 |
|  | 0.04 | 0.0, 0.01, 0.03, 0.05, 0.07, 0.1, 0.15, 0.20, 0.25, 0.30, 0.50, 0.70, 1.0, 1.5, 2.0, 2.5, 3.0 |
|  | 0.1 | 0.0, 0.01, 0.015, 0.02, 0.03, 0.05, 0.07, 0.1, 0.15, 0.2, 0.25, 0.3, 0.5, 0.7, 1.0 |
|  | 0.15 | 0.0, 0.002, 0.004, 0.006, 0.008, 0.01, 0.013, 0.016, 0.02, 0.025, 0.03, 0.05, 0.07, 0.1, 0.15, 0.2, 0.25, 0.3, 0.5, 0.7, 1.0 |
| A_6_-DNA  T7-H3  pH 8.8, 25 °C | 0 | 0.0, 0.05, 0.1, 0.15, 0.2, 0.25, 0.3, 0.5, 0.7, 1.3, 1.7, 2.0, 2.5, 3.0, 3.5 |
|  | 0.02 | 0.0, 0.01, 0.03, 0.05, 0.07, 0.1, 0.15, 0.2, 0.25, 0.3, 0.5, 0.7, 1.0, 1.5, 2.0, 2.5, 3.0 |
|  | 0.04 | 0.0, 0.01, 0.03, 0.05, 0.07, 0.1, 0.15, 0.2, 0.25, 0.3, 0.5, 0.7, 1.0, 1.5, 2.0, 2.5, 3.0 |
|  | 0.1 | 0.0, 0.01, 0.015, 0.02, 0.03, 0.05, 0.07, 0.1, 0.15, 0.2, 0.25, 0.3, 0.5, 0.7, 1.0 |
|  | 0.15 | 0.0, 0.002, 0.004, 0.006, 0.008, 0.01, 0.013, 0.016, 0.02, 0.025, 0.03, 0.05, 0.5, 0.7, 1.0 |
| A_6_-DNA  T22-H3  pH 8.8, 25 °C | 0 | 0.0, 0.05, 0.1, 0.15, 0.2, 0.25, 0.3, 0.5, 0.7, 1.3, 1.7, 2.0, 2.5, 3.0, 3.5 |
|  | 0.02 | 0.0, 0.01, 0.03, 0.05, 0.07, 0.1, 0.15, 0.2, 0.25, 0.3, 0.5, 0.7, 1.0, 1.5, 2.0, 2.5, 3.0 |
|  | 0.04 | 0.0, 0.01, 0.03, 0.05, 0.07, 0.1, 0.15, 0.2, 0.25, 0.3, 0.5, 0.7, 1.0, 1.5, 2.0, 2.5, 3.0 |
|  | 0.1 | 0.0, 0.01, 0.015, 0.02, 0.03, 0.05, 0.07, 0.1, 0.15, 0.2, 0.25, 0.3, 0.5, 0.7, 1.0 |
|  | 0.15 | 0.0, 0.002, 0.004, 0.006, 0.008, 0.01, 0.013, 0.016, 0.02, 0.025, 0.03, 0.05, 0.5, 0.7, 1.0 |

Summary of the relaxation delay times used in the imino proton exchange experiments. [NH_3_] denotes the effective ammonia concentration in the buffer. Secondary structures of the DNA constructs used for the imino proton exchange measurements are given in Fig. S4.

**Table S7.** Kinetic parameters obtained from imino proton exchange experiments.

| **Construct** | **pH** | **Temperature**  **(°C)** | **Residue** | **[NH_3_]**  **(M)** | ***R*_1w_**  **(s^-1^)** | **E** | ***R*_1n_**  **(s^-1^)** | ***k*_ex_**  **(s^-1^)** |
| --- | --- | --- | --- | --- | --- | --- | --- | --- |
| GDNA | 8.8 | 25 | 6 | 0.00 | 0.31 | 1.88 | 4.71±0.07 | 3.81±0.05 |
| GDNA | 8.8 | 25 | 6 | 0.02 | 0.34 | 1.86 | 49.68±2.11 | 52.25±2.00 |
| GDNA | 8.8 | 25 | 6 | 0.04 | 0.34 | 1.82 | 85.31±3.57 | 102.16±3.72 |
| GDNA | 8.8 | 25 | 6 | 0.10 | 0.33 | 1.96 | 140.76±8.43 | 232.07±11.90 |
| GDNA | 8.8 | 25 | 6 | 0.15 | 0.33 | 1.93 | 172.90±19.87 | 389.55±32.27 |
| A_6_-DNA | 8.8 | 25 | 5 | 0.00 | 0.35 | 1.90 | 1.19±0.07 | 0.26±0.01 |
| A_6_-DNA | 8.8 | 25 | 5 | 0.02 | 0.33 | 1.85 | 4.53±0.23 | 2.99±0.12 |
| A_6_-DNA | 8.8 | 25 | 5 | 0.04 | 0.33 | 1.78 | 6.28±0.28 | 4.60±0.17 |
| A_6_-DNA | 8.8 | 25 | 5 | 0.10 | 0.33 | 1.95 | 9.69±0.42 | 7.19±0.25 |
| A_6_-DNA | 8.8 | 25 | 5 | 0.15 | 0.32 | 1.93 | 11.07±0.47 | 8.19±0.28 |
| A_6_-DNA | 8.8 | 25 | 6 | 0.00 | 0.35 | 1.90 | 1.42±0.10 | 0.26±0.01 |
| A_6_-DNA | 8.8 | 25 | 6 | 0.02 | 0.33 | 1.85 | 4.16±0.20 | 2.65±0.10 |
| A_6_-DNA | 8.8 | 25 | 6 | 0.04 | 0.33 | 1.78 | 5.63±0.28 | 3.94±0.16 |
| A_6_-DNA | 8.8 | 25 | 6 | 0.10 | 0.33 | 1.95 | 8.08±0.34 | 5.75±0.19 |
| A_6_-DNA | 8.8 | 25 | 6 | 0.15 | 0.32 | 1.93 | 9.32±0.42 | 6.66±0.24 |
| A_6_-DNA | 8.8 | 25 | 7 | 0.00 | 0.35 | 1.90 | 1.99±0.14 | 0.47±0.03 |
| A_6_-DNA | 8.8 | 25 | 7 | 0.02 | 0.33 | 1.85 | 6.43±0.36 | 4.53±0.21 |
| A_6_-DNA | 8.8 | 25 | 7 | 0.04 | 0.33 | 1.78 | 8.96±0.40 | 6.83±0.25 |
| A_6_-DNA | 8.8 | 25 | 7 | 0.10 | 0.33 | 1.95 | 14.38±0.81 | 11.35±0.54 |
| A_6_-DNA | 8.8 | 25 | 7 | 0.15 | 0.32 | 1.93 | 17.60±0.68 | 14.66±0.49 |
| A_6_-DNA | 8.8 | 25 | 22 | 0.00 | 0.35 | 1.90 | 2.93±0.09 | 1.79±0.05 |
| A_6_-DNA | 8.8 | 25 | 22 | 0.02 | 0.33 | 1.85 | 33.43±0.63 | 31.67±0.56 |
| A_6_-DNA | 8.8 | 25 | 22 | 0.04 | 0.33 | 1.78 | 51.76±4.16 | 50.86±3.87 |
| A_6_-DNA | 8.8 | 25 | 22 | 0.10 | 0.33 | 1.95 | 88.40±2.11 | 98.78±2.19 |
| A_6_-DNA | 8.8 | 25 | 22 | 0.15 | 0.32 | 1.93 | 100.14±3.65 | 112.38±3.27 |

Kinetic parameters obtained from imino proton exchange experiments. [NH_3_] denotes the effective ammonia concentration in the buffer, *R*_1w_ and *R*_1n_ are the longitudinal relaxation rate constants of water and the thymine imino proton, respectively, E denotes the efficiency of inversion of the shaped pulse in the NMR experiment (Methods) and *k*_ex_ denotes the exchange rate of the thymine imino proton with water. Errors for *R*_1n_ and *k*_ex_ were set to be equal to the standard error of the fit and were computed as described in Methods. Secondary structures of the constructs are given in Fig. S4.

**Table S8.** Summary of exchange parameters obtained from fitting the imino proton exchange data as a function of ammonia concentration in GDNA and A_6_-DNA

| **Construct** | **pH** | **Temperature**  **(°C)** | **Residue** | ***p*_i_** | **ΔGº_conf,25ºC_(i)**  **(kcal/mol)** | **τ_open_ (s)** |
| --- | --- | --- | --- | --- | --- | --- |
| GDNA | 8.8 | 25 | 6 | (1.06±0.05)*10^-5^ | 6.79±0.03 | (2.96±2.73)*10^-4^ |
| A_6_-DNA | 8.8 | 25 | 5 | (8.14±0.54)*10^-7^ | 8.31±0.04 | (9.10±0.53)*10^-2^ |
| A_6_-DNA | 8.8 | 25 | 6 | (7.68±0.48)*10^-7^ | 8.34±0.04 | (11.97±0.65)*10^-2^ |
| A_6_-DNA | 8.8 | 25 | 7 | (1.15±0.08)*10^-6^ | 8.10±0.04 | (5.16±0.39)*10^-2^ |
| A_6_-DNA | 8.8 | 25 | 22 | (7.53±0.20)*10^-6^ | 7.00±0.02 | (5.40±0.40)*10^-3^ |

*p*_i_ denotes the population of the base open state. ΔGº_conf,25 ºC_(i) denotes the conformational penalty for opening the bp at 25 °C. τ_open_ denotes the inverse of the rate consant for opening of the bp. Errors of the exchange parameters were computed using a Monte-Carlo based approach as described in Methods. Buffer used for the measurements was composed of 10 mM sodium phosphate, 100 mM sodium chloride, 1 mM EDTA and 1 mM triethanolamine (TEOA) in 95% H_2_O:5% D_2_O.

**Table S9.** Sequence contexts used for testing the predictive power of delta-melt

| **Sequence**  **Context** | **Sample** | **Nuclei** | **pH** | **[NaCl]**  **(mM)** | **Temperature**  **(°C)** |
| --- | --- | --- | --- | --- | --- |
| 1 | scaf2_CGT^GC^ | G6-C1',C8 | 5.4 | 25 | 30 |
| 2 | scaf2_CGT^GC^ | G6-C1',C8 | 5.4 | 150 | 30 |
| 3 | scaf2_TGC^GC^ | G6-C1',C8 | 5.4 | 25 | 30 |
| 4 | scaf2_TGC^GC^ | G6-C1',C8 | 5.4 | 150 | 30 |
| 5 | scaf2_TGC^GC^ | G6-C1',C8 | 5.4 | 150 | 40 |
| 6 | scaf2_TGC^GC^ | G6-C8 | 5.4 | 150 | 15 |

Sequence context refers to the context numbers in Fig. 10b in the main text. Secondary structure of samples are given in Fig. S4. Nuclei denotes the atoms on which *R*_1ρ_ experiments were performed and [NaCl] denotes the concentration of sodium chloride used in the buffer for the measurements.

**Supplementary Discussions**

**Discussion S1: Extracting melting energetics from UV melting experiments**

In a typical UV melting experiment, a nucleic acid sample at a given concentration is slowly heated in a controlled manner to ensure the different conformational states are at equilibrium throughout the process (4). During the experiment, the absorbance of the sample, typically at 260 nm (A_260_), is recorded to obtain a melting curve (Fig. S15a). The baseline at low temperatures in the melting curve reflects temperature-dependent changes in the extinction coefficient of the folded (duplex/hairpin) species. Following the lower baseline, the melting curve (Fig. S15a) has a sigmoidal part during which the folded species gradually transitions into the unfolded single-stranded species. The baseline at high temperatures (Fig. S15a) then reflects temperature-dependent changes in the extinction coefficient of the unfolded single-stranded species.

The melting curves can be fit to a two-state model assuming only folded and unfolded single-stranded species are present in solution by least-squares minimization, to obtain the enthalpy (${\Delta H}_{\mathrm{melt}}^{^{\circ}}$) and entropy (${\Delta S}_{\mathrm{melt}}^{^{\circ}}$), and consequently the free energy (${\Delta G}_{\mathrm{melt}}^{^{\circ}}$=${\Delta H}_{\mathrm{melt}}^{^{\circ}}$-${\Delta S}_{\mathrm{melt}}^{^{\circ}}$) of melting, in addition to the extinction coefficients for the unfolded single-stranded and folded species (5, 6). It is important to note that accurate extraction of thermodynamic parameters from this ‘curve fitting’ approach requires well defined lower and upper baselines (~10-15 °C) (4). Indeed, samples for which we do not have well defined baselines (for example, A_6_-DNA^m1A16^ at pH 6.8, A_6_-DNA^m1G10^ at pH 6.8, data for A-T base opening in Fig. S1) do have relatively larger error bars in the delta-melt correlation plots (Fig. 3c, 5c, 6c). Deviations from fitting to a two-state model (Fig. S3 and S15b) can indicate presence of alternative species in solution such as intra or inter-molecularly folded single-strands (7-9).

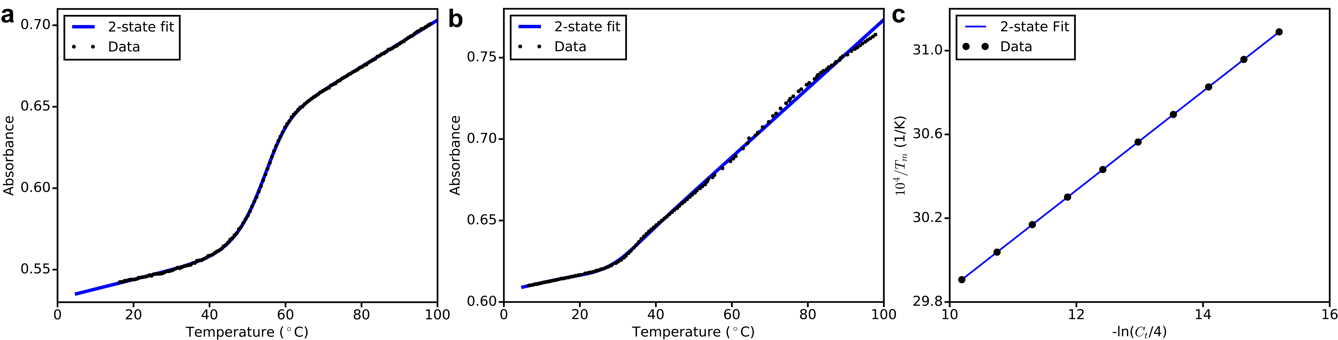

**Figure S15. Analysis of optical melting data.** Representative example of a melting curve (black dots) with two-state fit to the data (blue line) for (a) two and (b) non-two-state melting of a duplex. (c) Concentration-dependent melting data for duplex in (a)(black points).

As an independent test of the two-state model to fit the data for nucleic acid duplexes, UV melting experiments typically are also performed as a function of sample concentration (Fig. S15c)(4, 10, 11). The variation of the melting temperature (T_m_) of the duplex sample, the temperature at which melting is half complete, with sample concentration can also be fit to obtain ${\Delta H}_{\mathrm{melt}}^{^{\circ}}$ and ${\Delta S}_{\mathrm{melt}}^{^{\circ}}$. For duplex samples that melt in a two-state manner, the ${\Delta H}_{\mathrm{melt}}^{^{\circ}}$ and ${\Delta S}_{\mathrm{melt}}^{^{\circ}}$ values obtained from curve fitting at a given concentration or through concentration-dependent T_m_ measurements typically agree to within 15% (4, 5, 12).

Here it is important to note that agreement between the concentration-dependent and curve fitting approaches is a necessary but insufficient condition for two-state melting for duplexes (13). Validity of a two-state melting approximation can only be completely ascertained by supplementing melting curves with information from other techniques such as calorimetry, which can independently measure melting thermodynamics (12, 14, 15), or NMR (9, 16, 17) and analytical centrifugation (18, 19), which can monitor structural changes during melting. In this study, we used the curve fitting approach to extract thermodynamic parameters for all samples, given its simplicity and high throughput, and the fact that two-state melting behavior has been independently validated for some of the duplexes in this study using NMR (A_2_-DNA and A_6_-DNA)(20).

Lastly, it should also be noted that fitting of the data using both curve fitting or concentration-dependent measurements as described above typically assumes a temperature invariant enthalpy and entropy of melting i.e., a heat capacity change of zero during melting (5, 12). However, nucleic acid melting is well known to have a non-zero heat capacity change, which is generally attributed to the temperature-dependent unstacking when forming the single-stranded species (5, 21, 22). Interestingly, studies from the Turner lab have shown that the errors in ${\Delta H}_{\mathrm{melt}}^{^{\circ}}$ and ${\Delta S}_{\mathrm{melt}}^{^{\circ}}$ obtained by neglecting heat capacity changes mutually cancel one another, resulting in ${\Delta G}_{\mathrm{melt}}^{^{\circ}}$ values that are most accurate near the sample T_m_ (11, 21), even for samples that do not melt in a two-state manner (7, 23). In other studies, they have also shown that the errors in ${\Delta G}_{\mathrm{melt}}^{^{\circ}}$ on account of assuming a heat capacity of zero that accumulate on moving away from the T_m_ cancel one other when taking the difference in ${\Delta G}_{\mathrm{melt}}^{^{\circ}}$ between two samples, resulting in accurate values for ${\Delta\Delta G}_{\mathrm{melt}}^{^{\circ}}$ between two samples (23). Thus, even though our curve fitting based melting experiments cannot completely rule out non two-state behavior for samples for which the fit to the data is good, it is not expected to significantly affect the ${\Delta\Delta G}_{\mathrm{melt}}^{^{\circ}}$ values and the delta-melt correlations.

**Discussion S2: Offsets** $\mathbf{c(i)}$**from delta-melt experiments**

The offset $c(i)$ is the difference in energetic cost to modify the single-stranded (ss) versus the i^th^ minor conformational state i(WT), typically with a methyl group in the cases studied here. In this way, $c(i)$ carries interesting information concerning the difference in thermodynamic reactivity between the single-stranded and the minor conformation. This could for example find utility in understanding reactivities of nucleic acids structures to chemical probing reagents (24) or in defining the damage susceptibility of different nucleic acid conformations. Note that the $c(i)$ provides no information regarding kinetics, which could be important for the above processes.

Interestingly, for m^1^A^+^-T bps, $c(i)$ ~0.3 kcal/mol (Fig. 3c), suggesting that the free energy of *N*^1^-methylation of a single-stranded adenine, and an adenine in an A-T Hoogsteen bp are similar to each other. This supports findings from a recent study indicating that the A-T Hoogsteen bps increase the susceptibility of A-N1 to alkylation damage relative to Watson-Crick bps (25). Similar results are obtained for G-C^+^ Hoogsteen bps, with $c(i)$ ~-0.1 kcal/mol (Fig. 5c), suggesting that the free energy of *N*^1^-methylation of a single-stranded guanine, and a guanine in a G-C^+^ Hoogsteen bp are similar to each other.

Interestingly, for C-T mismatches $c(i)$ ~ -0.6 kcal/mol (Fig. 4c) is slightly negative, suggesting that it is more energetically favorable to replace an A to a C in single-stranded DNA, relative to an A-T Hoogsteen bp in duplex DNA, while that for T-T mismatches $c(i)$ ~-0.2 kcal/mol (Fig. 4e) is close to zero, suggesting that A to T substitutions in A-T Hoogsteen bps and single-strands are iso-energetic.

For A-T base opening, $c(i)$ ~2.4 kcal/mol, suggesting that the free energy of *N*^3^-methylation of T in an open A-T bp is much more favorable than the methylation of T when it is single-stranded, presumably due to favorable stacking interactions of the methyl within the context of the open bp in the duplex.

However, we cannot rule out that the large magnitude of $c(i)$ arises from over-estimation of the base catalyst accessibility factor $\alpha$. Although $\alpha$ is typically assumed to be 1 when interpreting imino proton exchange data (26), if $\alpha$ was truly a smaller number than one, it would lead to a reduction in $\Delta G_{\mathrm{conf}}^{○}(i)$ measured by NMR imino exchange, leading to a decrease in $c(i)$(Methods). Thus, given that m^1^A^+^, m^1^G and m^3^T occur naturally as forms of damage (27), delta-melt provides insights into the relative thermodynamic preferences of single- versus double-stranded DNA to undergo damage.

For isomerization of the m^6^A *N*^6^-methyl amino group, $c(i)$ ~ 0.7 kcal/mol, implying that *N*^6^-methylation of *syn* m^6^A in single-strand RNA is less thermodynamically favorable relative to *N*^6^-methylation of an m^6^A(*syn*)-U bp in duplex RNA, potentially due to stacking interactions of the methyl in the duplex context.

**Discussion S3: Rationalizing sequence-dependent stabilities of Watson-Crick and Hoogsteen bps obtained from delta-melt with prior literature**

The delta-melt data (Fig. 9b, Fig. S13) suggests that sequence-dependencies of the energies of the A-T Watson-Crick and G-C^+^ Hoogsteen bps dominate variations in the thermodynamic preferences for formation of A-T and G-C^+^ Hoogsteen bps, respectively. This is in contrast to earlier observations by Alvey *et al* suggesting that preferences for both A-T and G-C^+^ Hoogsteen bp formation are dominated by variations in the stability of the Watson-Crick bps. This observation was based on an inverse correlation of the thermodynamic propensities for forming Hoogsteen bps with the stability of the Watson-Crick bp computed using nearest neighbor thermodynamic parameters.

This discrepancy with Alvey *et al* for the results for G-C^+^ Hoogsteen bps can be rationalized based on the observation that the preferences in the prior study were averaged over different trinucleotide steps to compute an averaged preference for a dinucleotide step, while such averaging was not performed in this case, and because of differences in buffer conditions between nearest neighbor measurements and those used in this study.

**Discussion S4: Correlating cancer signatures with thermodynamic preferences from delta-melt**

Analysis of cancer genomes has shown that it is possible to decompose the set of all mutations in a cancerous genome into contributions from distinct mutational fingerprints or signatures, which are proposed to arise from different mutational processes acting on DNA (28). For many of these signatures, the underlying causative biological processes remain unknown.

Interestingly, we found a good correlation between the probabilities for formation of A-T Hoogsteen bps obtained from the thermodynamic preferences measured using delta-melt (Fig. 9b, Methods), and probabilities of T>A mutations corresponding to signature 17A (SBS17A, Fig. S16b, Methods), whose etiology is unknown, though it has been proposed that it could arise from 5-fluoro-uracil exposure (29) and/or oxidative damage to DNA (30). Transient A-T Hoogsteen bps which expose the Watson-Crick face of adenine to the solvent could be damaged by a damaging agent, resulting in a DNA lesion (Fig. S16c). If left unrepaired, this lesion can be replicated by translesion synthesis polymerases such as pol iota, which can misincorporate an adenine opposite the lesion, leading to a T>A mutation (Fig. S16c). Thus, the extent of this mutation that occurs would be proportional to the amount of lesion formed, which in turn would be proportional to the population of the transient Hoogsteen bp, resulting in the correlation we observe (Fig. S16b). Indeed, it has been shown that pol iota can misincorporate A opposite lesions on the Watson-Crick face such as m^1^A^+^ (31) *in-vitro*, leading to a T>A mutation. Additional experiments are required to test this hypothesis for the origin of T>A mutations in Signature 17A. As a control, a weaker correlation is observed for T>A mutations in Signature 8 (SBS8), whose etiology is also unknown (Fig. S16b).

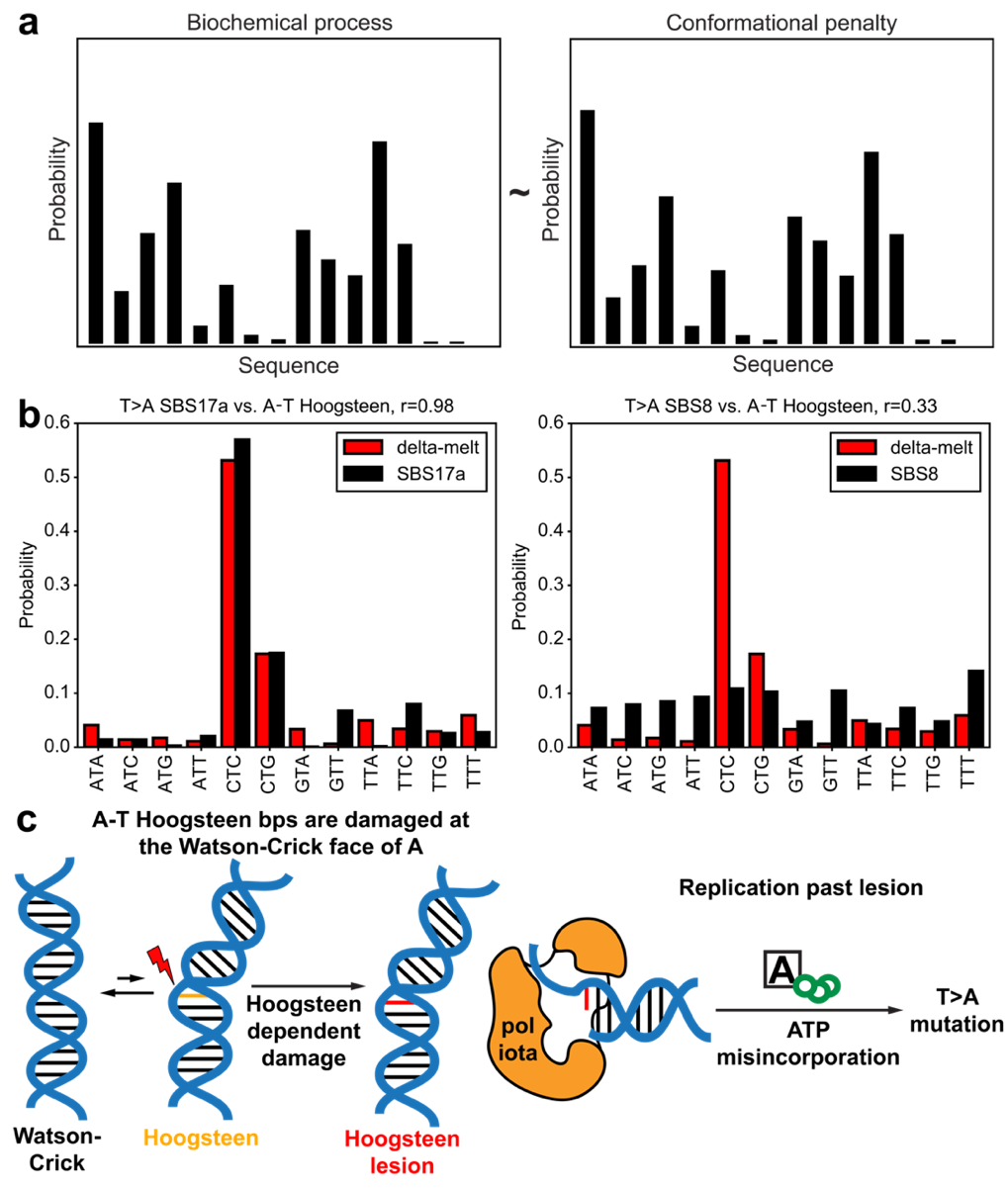

**Figure S16. Using delta-melt to identify dynamic drivers of biochemical processes.** (a) Correlating sequence-dependent biochemical processes to thermodynamic preferences from delta-melt may offer insights into dynamic drivers of these processes. (b) Correlation of the probabilities of A-T Hoogsteen bp formation from delta-melt (Methods) with the probabilities for T>A mutations in Signature 17A (SBS17A, left) and Signature 8 (SBS8, right). (c) Proposed mechanism for the origins of Signature 17A. Transient A-T Hoogsteen bps undergo damage on the Watson-Crick face of A. The resulting DNA lesion if left unrepaired, can be replicated by pol iota, which can misincorporate an Adenine, leading to a T>A mutation.

**Discussion S5: Comparison of enthalpy and entropy from NMR and delta-melt**

Comparison of $\Delta H_{\mathrm{conf}}^{○}(i)$ and $\Delta S_{\mathrm{conf}}^{○}(i)$ values obtained from temperature-dependent NMR experiments with $\Delta\Delta H_{\mathrm{melt}}^{○}(i)$ and $\Delta\Delta S_{\mathrm{melt}}^{○}(i)$ values from delta-melt show that in some cases, the agreement is better than others (Fig. S14). This could be due to systematic errors in estimation of enthalpies and entropies in UV melting measurements due to the assumption of a heat capacity of zero i.e., neglecting the temperature-dependence of the enthalpy and entropy (11). These errors are compensatory in nature such that estimate of free energies from UV melting is accurate (11). Nevertheless, we cannot rule out that deviations in enthalpies and entropies from delta-melt and NMR are also due to process and/or sequence specific non-zero $c_{\mathrm{Enthalpy}}(i)$ and $c_{\mathrm{Entropy}}(i)$ values (Methods, Equation 23-24).
